# Expansion and Adaptive Evolution of the *mTERF* Gene Family in Plants

**DOI:** 10.1101/2022.01.18.476766

**Authors:** Yanxin Zhao, Manjun Cai, Meijie Luo, Jianhua Zhang, Yurong Li, Xiaobo Zhang, Bing Yue, Hailin Xiao, Jiuran Zhao, Yonglian Zheng, Fazhan Qiu

## Abstract

Mitochondrial transcription termination factor (*mTERF*) genes are encoded in the nucleus and bind to nucleic acids to regulate the replication, transcription and translation of mitochondrial genomes. Plants possess a large family of *mTERF* genes that play important roles in regulating organellar gene expression and stress response. However, their origin and expansion in land plants has not been examined. Here, we conducted a comprehensive molecular evolution analysis of 611 *mTERF* genes identified in 18 plant species, including algae, moss, fern, gymnosperm and flowering plants. Higher plants have more *mTERF* genes compared to lower plants, forming a huge higher plant-specific clade (M-class *mTERF* genes). M-class *mTERF* genes occur in clusters, suggesting that tandem duplication contributed to their expansion. Compared to other *mTERF* genes, M-class *mTERF* genes have undergone rapid evolution, and several significant positively selected sites were located in nucleic acid-binding sites. The strong correlation between the number of M-class *mTERF* genes and corresponding mitochondrial genome variation suggests that the rapid evolution of M-class *mTERF* genes might account for the changes in the complex machinery for expression regulation of plant mitochondrial genomes, providing molecular evidence for the host-parasite interaction hypothesis between the nucleus and mitochondria.

## Introduction

Eukaryotes have two semi-autonomous organelles that harbor extranuclear genetic materials, namely, mitochondria (in fungi, plants and animals) and chloroplasts (in plants), which were respectively derived from the endosymbiosis of α-proteobacterial and cyanobacterial ancestors (Gray et al. 1999; Howe et al. 2008). During endosymbiotic evolution, most organellar genes were either lost or transferred to the nucleus, leaving only a small part encoded in organellar genomes (Timmis et al. 2004). Most proteins involved in biogenesis, development, energy synthesis and metabolism within these organelles are encoded by the nuclear genome, such that growth and division of mitochondria and chloroplasts are regulated by host–cellular machinery (Osteryoung and Nunnari 2003). Inversely, functional changes in organelles regulate the expression of nuclear genes in a retrograde signaling pathway that coordinates nuclear and organellar activities (Kleine and Leister 2016; Woodson and Chory 2008).

Unlike their bacterial ancestors, organellar gene expression is regulated by hundreds of nuclear gene products (Weihe et al. 2012; Woodson and Chory 2008). Of these plant nuclear genes, several gene families and subfamilies encoding mitochondria- or chloroplast-targeted proteins have been formed, such as the *MORF/RIP* gene family for organellar RNA editing (Sun et al. 2016a), sigma factor gene family for chloroplast gene transcription (Ueda et al. 2013), and pentatricopeptide repeat (PPR) gene family for organellar RNA processing (O’Toole et al. 2008). PPR proteins are composed of multiple tandem repeats of a degenerate 35-amino acid motif, and based on the type and combination of PPR motifs can be further divided into 3 subfamilies, namely, the PLS-PPR genes, P-PPR genes and *Rf-PPR-like* (*RFL*) genes (Fujii et al. 2011; Lurin et al. 2004). The expansion and evolution of the huge PPR gene family in higher plants has been associated with diverse posttranscriptional processes in organelles, including RNA cleavage, editing, stability and translation (Schmitz-Linneweber and Small 2008; Stern et al. 2010). The PLS-PPR genes have undergone purifying selection of RNA edited sites in the mitochondria, and conversely, chloroplast genome evolution is constrained by the PPR genes (Fujii and Small 2011; Hayes and Mulligan 2011; Hayes et al. 2012; O’Toole et al. 2008). Diversifying selection of *RFL* genes is driven by chimeric genes in the mitochondria, which have been strongly associated with cytoplasmic male sterility (CMS) in flowering plants. Adaptive selection of the *RFL* genes provides molecular evidence for the “arms-race” between the nuclear and mitochondrial genomes, similar to a host-parasite relationship (Fujii et al. 2011).

Another nuclear-encoded gene family encoding organelle-targeted proteins was recently identified in plants, the *mitochondrial Transcription tERmination Factor* (*mTERF*) gene family. To date, *mTERF* genes have only been identified in animals and plants and are missing in fungi and prokaryotes (Kleine 2012; Linder et al. 2005). In animals, mTERF proteins are characterized by tandem repeats of a degenerate 32-amino acid mTERF motif and play important roles in replication, transcription, translation of mitochondrial DNA (mtDNA), and mitochondrial ribosomal biogenesis by directly binding to DNA or RNA (Kleine and Leister 2015). Human mTERF1 promotes the termination of mitochondrial transcripts from the first transcription initiation site (H1) at the 3’-end of the 16S rRNA gene by binding specifically to mtDNA (Fernandez-Silva et al. 1997; Kruse et al. 1989). Crystal structure analysis has revealed that human mTERF1 can form a positively charged groove that can bind to mtDNA or mtRNA (Yakubovskaya et al. 2010).

In plants, the *Chlamydomonas reinhardtii MOC1* gene has the evolutionarily conserved transcription termination activity similar to that of human *mTERF1*. MOC1 can specifically bind to an octanucleotide sequence within the mitochondrial rRNA-coding module S3 to suppress the read-through transcription of mitochondrial RNA (Wobbe and Nixon, 2013). Nonetheless, higher plants have more *mTERF* genes than lower plants and animals. Of these, seven *mTERF* genes documented in *Arabidopsis* and maize play roles in the expression of organelle genes and chloroplast or mitochondria biogenesis (Quesada 2016). Only one mTERF protein has been reported in higher plants, namely, *Arabidopsis mTERF6*, which promotes the termination of transcription within *trnI.2* by binding specifically to an RNA sequence within the chloroplast rRNA operon *in vitro* (Romani et al. 2015). *Arabidopsis BSM/RUG2/mTERF4* and maize *Zm-mTERF4* are homologs and are essential to RNA splicing of group II introns in chloroplasts and their loss impedes chloroplast development and plant growth (Babiychuk et al. 2011; Hammani and Barkan 2014; Sun et al. 2016b). *ArabidopsismTERF15* is required for mitochondrial *nad2* intron 3 splicing (Hsu et al. 2014). Additionally, alterations in the expression of organelle genes due to mutations in *mTERFs* regulates nuclear gene expression through retrograde signaling to alleviate abiotic stress (Kim et al. 2012; Meskauskiene et al. 2009; Robles et al. 2010a; Robles et al. 2015; Sun et al. 2016b). However, only a few *mTERF* genes have been extensively studied and our understanding of their expansion, evolutionary pattern and significance is limited. Here, we performed a systematic genome-wide identification and comparison of *mTERF* genes in 18 organisms that encompass diverse plant lineages. Our results suggest that plant *mTERF* genes are similar in terms of function and evolutionary patterns to PPR genes. One group of *mTERF* genes (M-class *mTERF*) that specifically occurs in higher plants has undergone strong positive evolution and its origin and expansion may be related to variations in the mitochondrial genome. The characterization of these genes may facilitate the development of molecular tools to elucidate co-ordination between the nucleus and mitochondria. Moreover, this study provides insight into the regulatory roles of *mTERF* gene expression in organelles.

## Results

### Higher plants have more *mTERF* genes than lower plants

*mTERF* genes were identified in 18 plant species that represent different branches of the evolutionary tree of green plants (fig. 1). *C. reinhardtii* and *Volvox carteri* are unicellular and multicellular aquatic green algae, respectively, belonging to the Chlorophyta. The two algae are members of lineages that diverged before the evolution of land plants. *Physcomitrella patens* (a moss) and *Selaginella moellendorffii* (spikemoss, a lycophyte) are representatives of two early-divergent lineages of land plants. Moss diverged prior to the emergence of vascular plants, whereas spikemoss diverged before the origin of seed plants. *Picea abies* (Norway spruce) was included in the present study as a representative gymnosperm. The remaining species examined in this study were angiosperms, which included monocots, eudicots, and one basal angiosperm, *Amborella trichopoda*. The monocots investigated in this study consisted of *Brachypodium distachyon* (purple false brome), *Oryza sativa ssp.* japonica (japonica rice), *Zea mays* (maize), and *Sorghum bicolor* (sorghum),whereas the eudicots comprised *Vitis vinifera* (grape), *Carica papaya* (papaya), *Arabidopsis thaliana* (thale cress), *Arabidopsis lyrata* (rockcress), *Cucumiss ativus* (cucumber), *Glycine max* (soybean), *Medicago truncatula* (barrel medic, a close relative of alfalfa), and *Populus trichocarpa* (poplar). Finally, 7 *mTERF* genes in *C. reinhardtii*, 9 in *V. carteri*, 15 in *P. patens*, 13 in *S. moellendorffii*, 60 in *P. abies*, 51 in *A. trichopoda*, 41 in *B. distachyon*, 34 in rice, 28 in maize, 35 in sorghum, 30 in *V. vinifera*, 26 in *C. papaya*, 35 in *A. thaliana*, 46 in *A. lyrata*, 50 in *C. sativus*, 57 in *G. max*, 23 in *M. truncatula*, and 52 in *P. trichocarpa* were identified (fig. 1). Three maize *mTERF* genes, *ZmTERF4*, *ZmTERF7* and *ZmTERF20*, were excluded from the analysis because these were assumed to be pseudogenes (Zhao et al. 2014). In this study, several *mTERF* genes were re-annotated, such as *LOC_Os06g12060.1* in rice and *GSVIVT01028382001* in grape. The former was divided into two independent intronless *mTERF* genes, *LOC_Os06g12060.1a* and *LOC_Os06g12060.1b*, whereas four new grape *mTERF* genes were detected around the *GSVIVT01028382001* gene (supplementary table S1). A total of 611 *mTERF* genes were identified in 18 plants and detailed information is presented in supplementary table S1.

**Figure 1.**
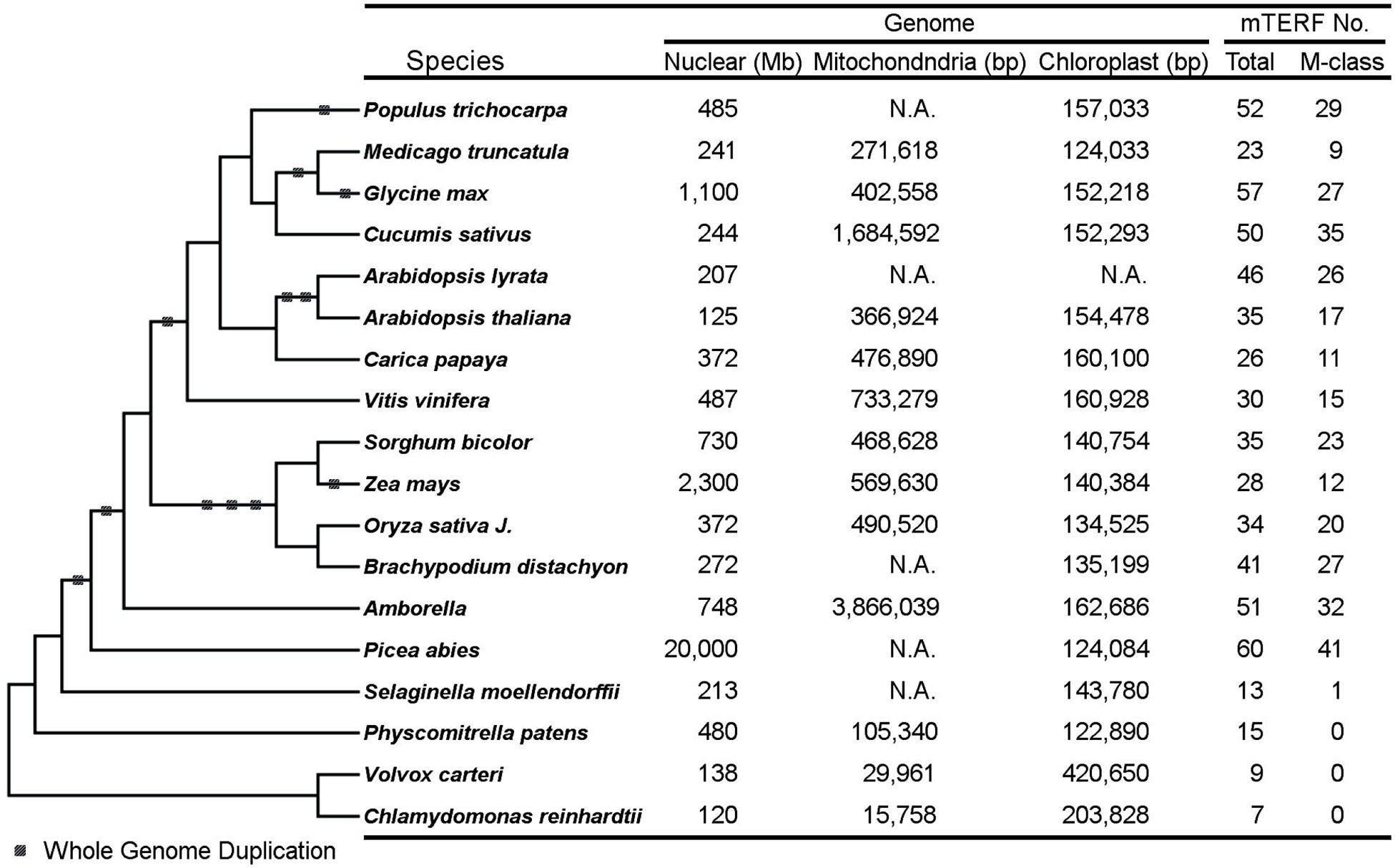
Summary of 18 plant genomes used and *mTERF* genes identified in this study. The tree on the left shows the relative evolutionary position of the plant species that was built on the base of the plant trees proposed in the PGDD database (http://chibba.agtec.uga.edu/duplication/). The information on the plant nuclear and organellar genomes in the table on the right were obtained from their genome database and NCBI Organelle Genome Resources (http://www.ncbi.nlm.nih.gov/genome/organelle/), respectively.

Compared to mammals that harbor four *mTERF* genes, plants have a higher number. Furthermore, higher plants have more *mTERF* genes than lower plants (fig. 1). Protein sequence analysis predicted that most plant *mTERF* proteins target the mitochondria or chloroplasts. In addition, intronless mTERF genes were predominant (76% in rice and 69% in *Arabidopsis*) (supplementary table S1).

### Classification and evolution analysis of *mTERF* genes

To investigate the phylogenetic relationship of plant *mTERF* genes, a distance tree consisting of 611 *mTERF* proteins was generated using the neighbor-joining (NJ) method. As shown in fig. 2, there were seven groups (I–VII) divided based on the resemblance of the NJ tree topology to previously constructed *Arabidopsis* and maize phylogenetic trees (Babiychuk et al. 2011; Zhao et al. 2014). Group VII is the largest cluster, of which 53% (325/611) consisted of *mTERF* genes that form species-specific paralogous clusters, indicating that these genes had undergone extensive evolution after divergence of these lineages. In addition, all of the group VII genes were from higher plants, except for *Sm444400*, which was identified in *S. moellendorffii*. In group VII, more than 68% (222/325) of the mTERF proteins were predicted to target mitochondria and were thus designated as M-class *mTERF* genes. Accordingly, other *mTERF* genes, including those in groups I–VI, were non-M-class *mTERF* genes. Most non-M-class *mTERF* genes formed orthologous clades, shared by all plant species examined in this study (fig. 2), suggesting that they evolved before these plants diverged and might play conserved roles. For instance, clade IV consists of *mTERF* genes from different plants and its topology resembles the evolutionary tree, indicative of plant evolution history (supplementary figure S1).

**Figure 2.**
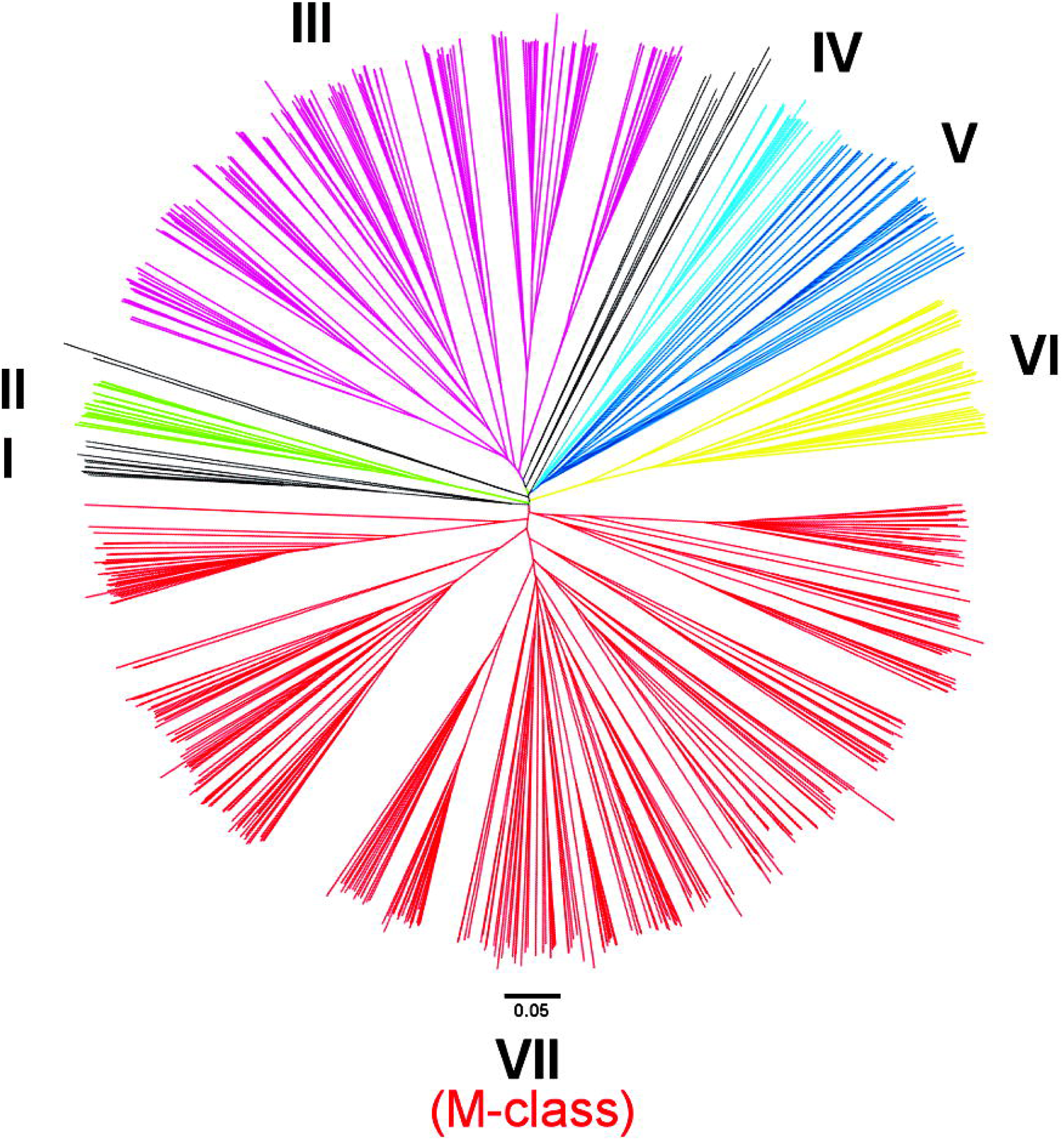
The phylogenetic relationships of all plant *mTERF* genes identified in this study. The neighbor-joining tree containing all plant *mTERF* genes was constructed by ClustalW 2.0 and displayed with FigTree v1.4.0. Groups I-VII outside the tree indicate the *mTERF* groups divided based on their phylogenetic relationship. The red clade consists of land plant-specific M-class *mTERF* genes.

### Tandem duplication of M-class *mTERF* genes account for the increase in the number of *mTERF* genes in higher plants

Seed plant genomes encode an average of 17 non-M-class *mTERF* genes, which is comparable to that in *P. patens* (15) and *S. moellendorffii* (13) and all together are included in the non M-class clades (fig. 3). Each of the higher plants has about 23 M-class *mTERF* genes that forms pecies-specific clades. Different plants have variable numbers of M-class genes. *P. abies* harbors 41 M-class *mTERF* genes, whereas *M. truncatula*has only 9. In higher plants, the number of M-class *mTERF* genes is independent of the number of non-M-class genes. However, there are two exceptions, namely in *G. max* and *P. trichocarpa*, which have more M-class and non-M-class *mTERF* genes (fig. 2) due to a recent whole genome duplication (WGD) event (Schmutz et al. 2010; Tuskan et al. 2006). These results demonstrate that the emergence and expansion of M-class *mTERF* genes in higher plants is responsible for the expansion of *mTERF* genes (r = 0.968, *P* < 0.01).

**Figure 3.**
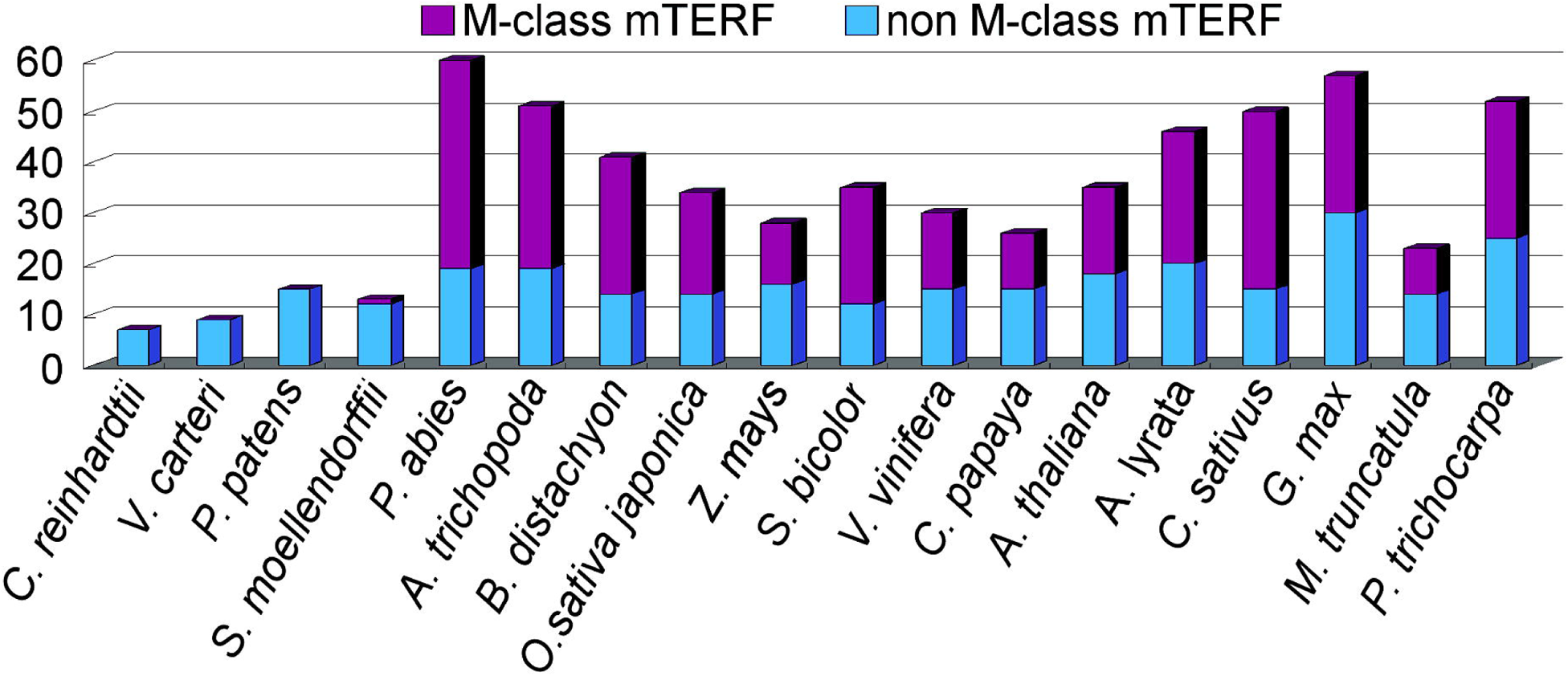
Comparison of the M-class and non-M-class *mTERF* genes.

To determine the causes of gene duplication of *mTERF* genes in higher plants, block duplication and tandem duplication events involved in the paralogous generation of *mTERF* genes were predicted using WGMapping, as provided by PLAZA v2.5 (Van Bel et al. 2012). As described in fig. 4, tandem duplication is the main driving force for the amplification of the members of the *mTERF* gene family, especially the M-class *mTERF* subfamily. Over 50% of the M-class *mTERF* genes in most of plants are tandemly duplicated except for those in maize (Zhao et al. 2014) and sorghum (fig. 4). Moreover, in *O. sativa* and *P. trichocarpa*, more than 70% of the M class genes are attributable to tandem duplication (fig. 4). Since tandemly duplicated genes often occur as clusters on chromosomes, we mapped the M-class *mTERF* genes onto the corresponding chromosomes or scaffold contigs of the plants that were queried in the present study, including *A. trichopoda*, rice, *Arabidopsis* and *C. sativus*. The classical gene clusters were apparently found on these genomes and all of the genes in the clusters were tandemly duplicated (supplementary figure S2). In rice, eight M-class genes were clustered on chromosome 6 within a 33-kilobasepair (kb) region (nucleotides 6,452,737–6,485,677) and in the 33-kb region, only one gene (*LOC_Os06g12090*) was not identified as an *mTERF* gene.

**Figure 4.**
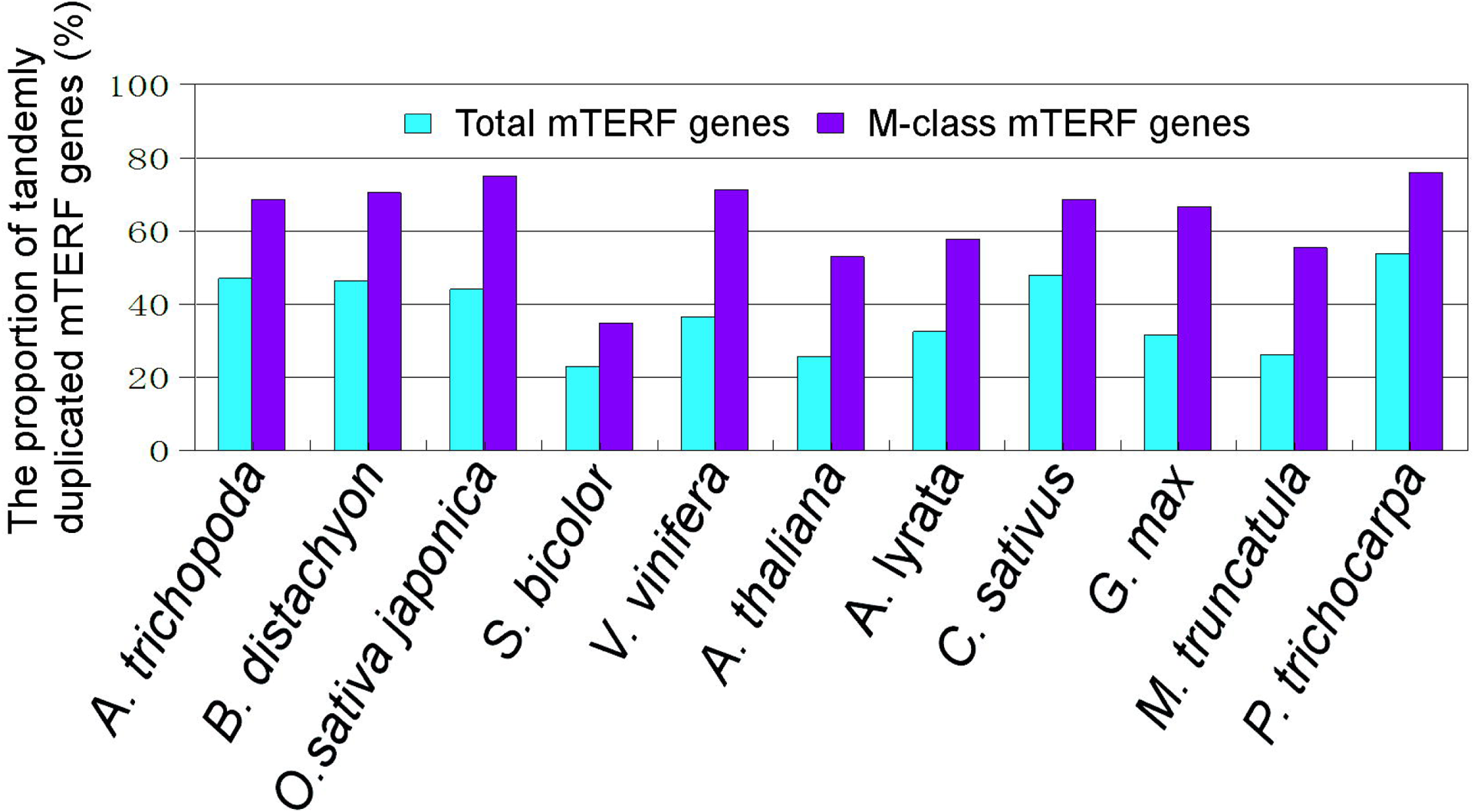
The proportion of tandemly duplicated *mTERF* genes in all *mTERF* genes. The tandem duplication event was estimated by using WGMapping in PLAZA v2.5 (Van Bel et al. 2012).

In *Arabidopsis*, a 9-M-class gene cluster was mapped to chromosome 1, within a < 70-kb region (nucleotides 22,902,150–22,971,764) (supplementary figure S2). Most chromosomal regions harboring M-class gene clusters were < 100 kb in length, thus again proving that tandem duplication plays an important role in the distribution and expansion of M-class *mTERF* genes in higher plants.

### Rapid evolution of M-class *mTERF* genes in plants

To estimate when M-class *mTERF* gene duplication had occurred, all 325 M-class mTERF proteins were aligned and used to build a maximum-likelihood (ML) inferred phylogenetic tree using RaxML (Stamatakis 2006) (fig. 5). A tree file in newick format is provided in supplementary data S1. Among the 14 higher plants examined in the present study, four monocots (rice, sorghum, maize and *B. distachyon*) and three dicots (*A. thaliana*, *A. lyrata*and *M. truncatula*) did not form species-specific clades. *A. thaliana* and *A. lyrata* M-class genes were grouped into a large clade, although two *A. lyrata* M-class genes matched only one *A. thaliana* gene in most of the tiny branches (supplementary figure S3). *M. truncatula* M-class genes and some of the *G. max* genes together generated one clade (G.max_2). Most of the

**Figure 5.**
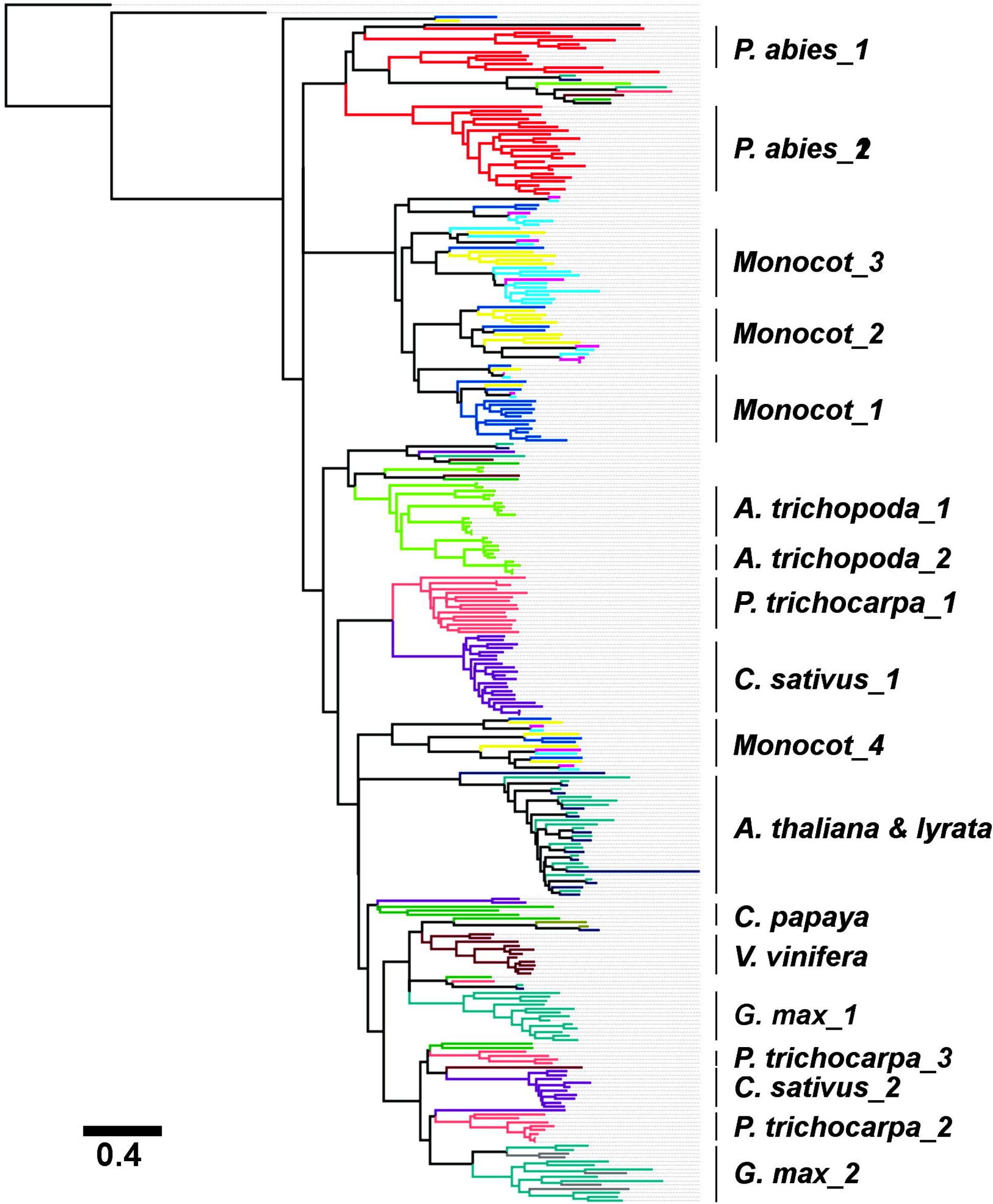
Maximum-likelihood method-based phylogeny of M-class *mTERF* genes in higher plants. The ML tree was built using RAxML (v7.2.8) with PROTGAMMABLOSUM62 as amino acid substitution model, 500 resampling for bootstrap test, and *A. thaliana At1g74120* as the outgroup. The monocots include *B. distachyon*, *O. sativa*, *S. bicolor* and *Z. mays*. The branches of the same species are marked by the same colors.

M-class genes in the rest of the higher plants formed at least one species-specific clade in the ML tree (fig. 5), indicating that the duplication of M-class genes occurred after these species diverged and the M-class genes underwent rapid evolution. In *A. trichopoda*, 24 out of 27 M-class genes were grouped into a single clade within the ML tree. The M-class genes of *P. trichocarpa* form three independent clades. Previous studies have estimated that the *P. trichocarpa* genome underwent a single WGD event approximately 8–13 million years ago (mya) (Sterck et al. 2005; Tuskan et al. 2006). Over 8,000 pairs of duplicated genes from this WGD event were reserved in *P. trichocarpa* genome, resulting in *Populus* having an average of 1.4–1.6 putative homologs for each *Arabidopsis* gene (Tuskan et al. 2006). The WGD event also led to an increase in non-M-class genes in *Populus* (fig. 3). There are three clusters of tandemly duplicated M-class genes on chromosomes 1 (PtC1_1 stands for the first cluster on chromosome 1 of *P. trichocarpa*), 3 (PtC3_1) and 4 (PtC4_1), respectively (supplementary figure S2). The PtC1_1 and PtC3_1 gene clusters were mapped to a homologous region that was formed during a WGD event that involved chromosomes 1 and 3. Two distinct subclades were formed by the PtC1_1 and PtC3_1 genes, although these belong to the same clade in the ML tree of *Populus* M-class genes (supplementary figure S4). Only one M-class gene in the region on chromosome 11 showed homology to the PtC4_1 cluster-containing region on chromosome 4, suggesting that the PtC4_1 gene cluster was generated after the WGD event (< 8 mya).

Another example of the rapid evolution of M-class *mTERF* genes was observed by comparing various monocots. Currently, monocots are thought to have undergone three rounds of WGD events after their divergence from dicots; the earliest one occurred before the Poaceae diverged from Zingiberales, whereas the latter two WGD events took place prior to the divergence of rice from maize and sorghum, approximately 120 mya and 70 mya, respectively (Tang et al. 2010; Wang et al. 2011). In addition, another recent WGD event involving the maize genome occurred approximately 5–12 mya (Schnable et al. 2009) (fig. 6). Rice, sorghum, maize and *B. distachyon* have similar numbers of *mTERF* genes, thus indicating that the WGD event involving maize did not result in an increase in the number of *mTERF* genes. Each monocot genome has at least one gene cluster comprising more than two M-class *mTERF* genes. Only one cluster in the maize genome was identified on chromosome 5, which we designated as ZmC5_1 (standing for the first cluster on chromosome 5 of *Zea mays*). Two M-class gene clusters (SbC1_1 and SbC4_1) were detected in sorghum, two (OsC6_1 and OsC11_1) in rice and four (BdC1_1, BdC3_1, BdC3_2) in *B. distachyon* (supplementary figure S5). As demonstrated in fig. 5, monocot M-class genes belong to two separated clades, with one clade subdivided into three subclades (Monocot_1, Monocot_2 and Monocot_3). In the ML tree that only contains monocot M-class genes, however, some M-class clusters in *B. distachyon*, rice, and sorghum also consisted of minor species-specific clades. For example, Monocot_1 consisted of BdC3_1 and BdC4_1, Monocot_2 comprised OsC6_1 and Monocot_3 included SbC1_1 (fig. 7), suggesting that these clusters underwent extensive evolution after species divergence. ZmC5_1, SbC4_1, OsC6_1, BdC1_1 and BdC3_2 were all mapped to inter- or intra-genomic homologous regions, whereas the variation in the number of M-class genes in the BdC1_1, SbC4_1 and OsC6_1 clusters occurred after these species had diverged (fig. 7). We did not detect any *mTERF* genes in the homologous regions of the SbC1_1, BdC3_1, BdC4_1, and OsC11_1 clusters, indicating that these genes emerged after sorghum had diverged from maize at around *<*11.9 mya (fig. 6) (Swigonová et al. 2004). Here, it is worth noting that the opportunity of tandem duplication to each of the clustered M-class genes is unequal when they contributed to the generation of gene clusters. For example, *LOC_Os06g12100.1* from the OsC6_1 cluster located within a 33-kb region does not belong to the same clade with others (fig. 7). The proposed history of tandem duplication events involving the OsC6_1 genes based on their phylogenetic relationship is presented in supplementary figure S6.

**Figure 6.**
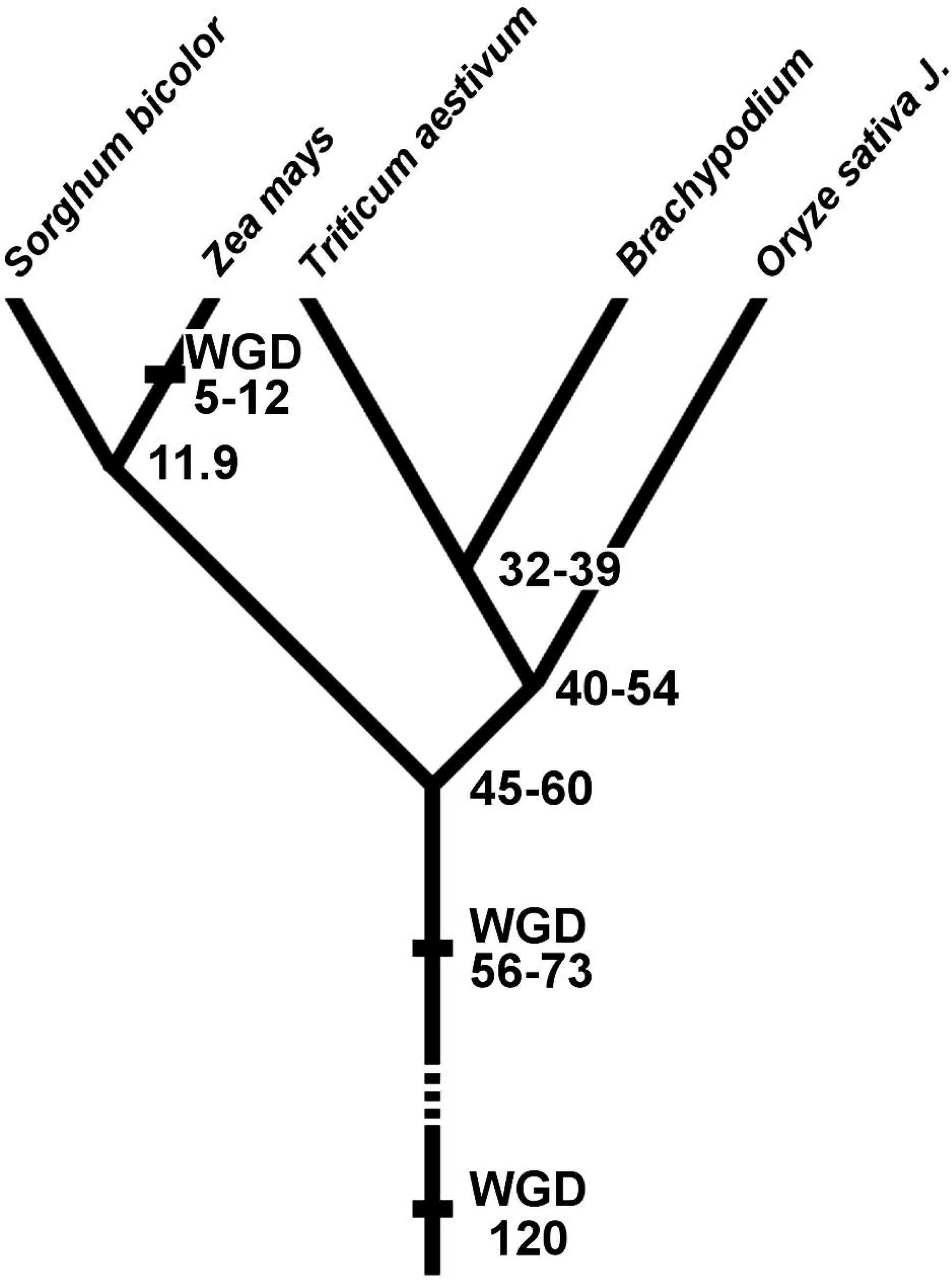
Divergence and WGD of grasses The time of WGD as earlier estimated (Schnable et al. 2009; Tang et al. 2010; Wang et al. 2011; Vogel et al. 2010). Time unit, mya (million years ago).

**Figure 7.**
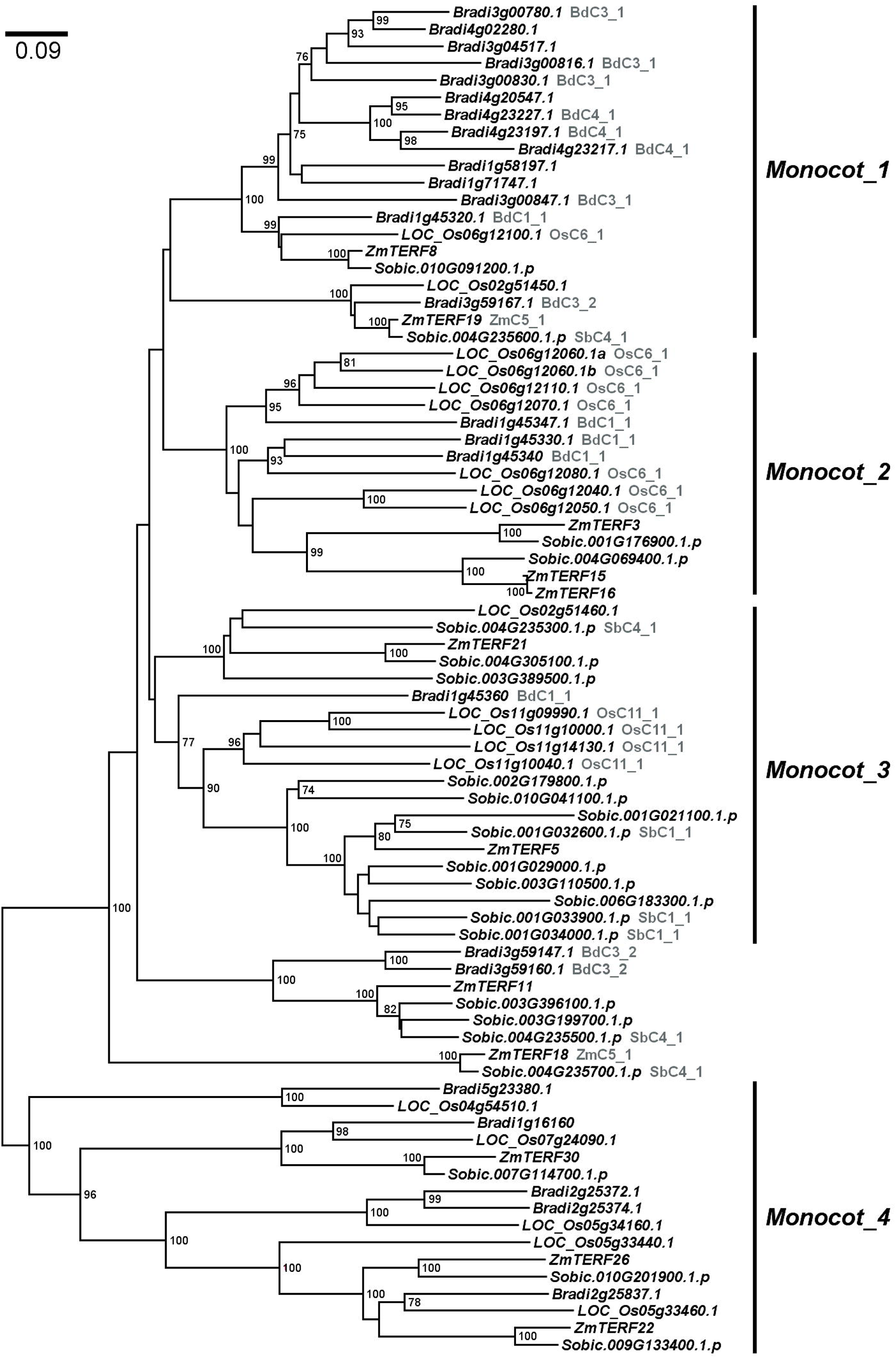
ML method-based phylogeny of the monocot M-class *mTERF* genes. The mTERF proteins were aligned using MUSCLE v3.8.31 (Edgar 2004) and the alignment was modified by CINEMA 5 (http://aig.cs.man.ac.uk/research/utopia/cinema/cinema.php). The ML tree was built by MEGA v6.0 (Tamura et al. 2013) with a JTT amino acid substitution model and 500 resampling for bootstrap test.

### Diversifying selection on M-class *mTERF* genes

M-class *mTERF* genes encode an average of 366 amino acids and have 3–8 mTERF motifs confirmed by using SMART (Letunic et al. 2012). The M-class proteins in dicots (5.8 motifs per M-class protein on average) have a higher number of mTERF motifs than monocots (4.6 on average) (supplementary table S1).Unlike non-M-class mTERF proteins, M-class proteins possess several exclusive conserved motifs that were identified by using MEME (Babiychuk et al. 2011; Zhao et al. 2014). The rapid evolution of the M-class genes was apparently driven by the diversifying selection of their protein sequences. Eight paralogous gene sets from four well-annotated plant genomes (*A. thaliana*, *A. lyrata*, rice, and *P. trichocarpa*) as representatives of higher plants were used to calculate the probabilities of diversifying selection on the M-class *mTERF* genes by using the PAML v4.7 package (Yang 2007). The data, including paralogous gene alignments, trees and parameters used for PAML analysis are presented in supplementary data S2 and supplementary table S2. Likelihood ratio (LR) tests were applied to determine the best codon substitution models that fit the observed values, including the M1, M2, M7 and M8 models, of which M1 and M7 are the null hypotheses that assume purifying or neutral selection and M2 and M8 allow diversifying selection (Yang et al. 2000; Yang 2007). Bayes empirical Bayes (BEB) was used to define the sites of interest that underwent diversifying selection when the M8 model was accepted (Yang et al. 2005) (see *Materials and Methods* for details).

Each of the eight gene sets has an estimate of ω > 1 under the M2 and M8 models and only one exception are the Osa_3 genes with ω = 1 under the M2 model (*P* = 0.99). A high degree of diversifying selection on M-class *mTERF* genes in plants was observed under the M8 model (at worst *P* = 0.0059 for the Osa_3 gene set) (table 1). The number of significantly positively selected sites obtained in different gene sets under the M8 model varied even in the same species, which ranged from 6 in Ptr_2 to 72 in Aly_1 with a posterior probability of 95% as a cutoff (table 1). Meanwhile, the positively selected sites were not conserved in these plants (supplementary table S2), suggesting that diversifying selection facilitated the emergence of different novel functions for the M-class *mTERF* genes.

**Table 1.**
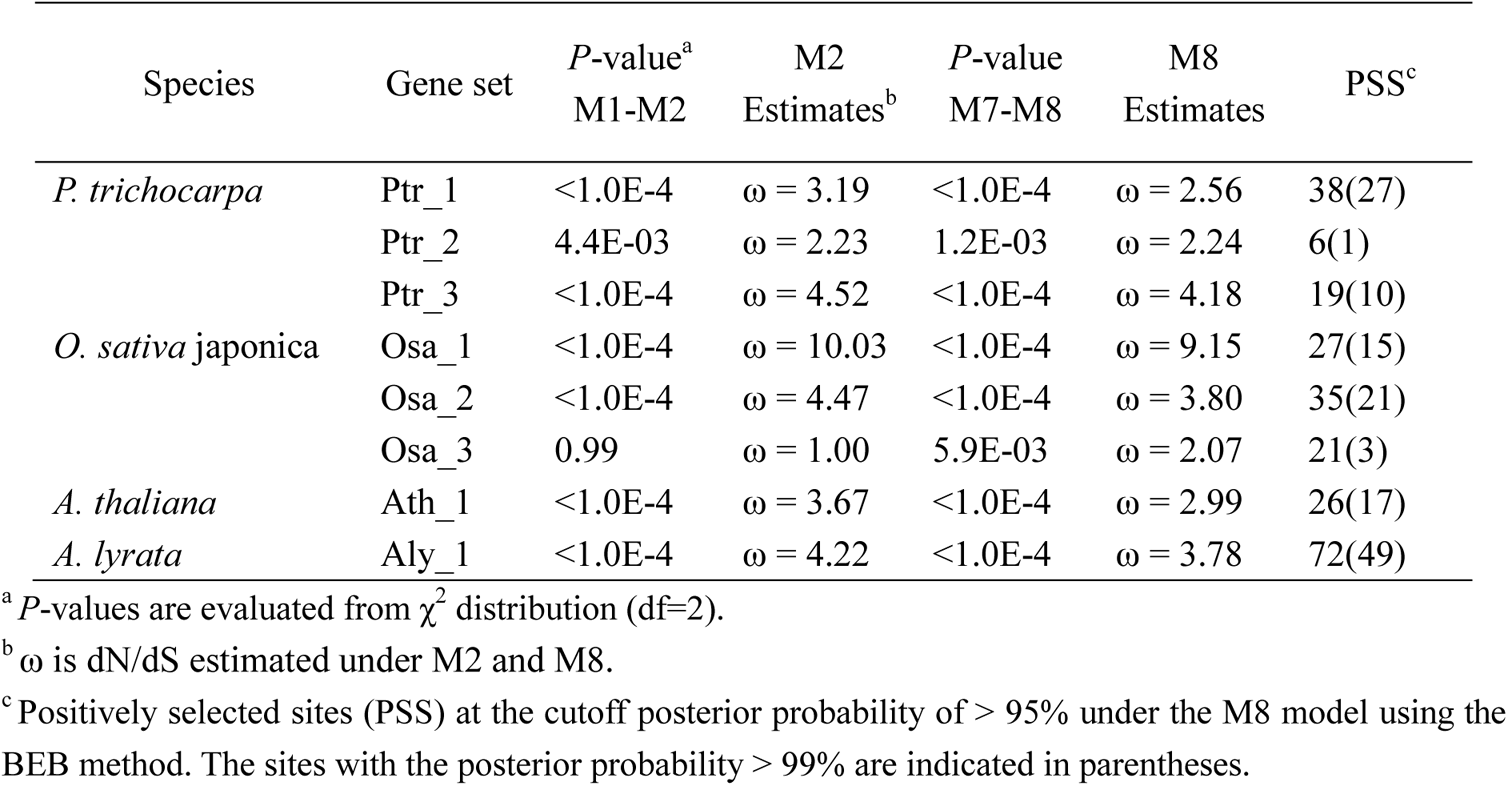
Comparison of codon substitution models in plant M-class *mTERF* genes

### Residues under positive selection in M-class mTERF proteins are involved in nucleotide-binding activation

Although the structures of the human mTERF proteins have been determined by X-ray crystallography (Jiménez-Menéndez et al. 2010; Spåhr et al. 2010; Yakubovskaya et al. 2010, 2012), no plant mTERF protein structures have been experimentally determined. Based on homology modeling, the structure of the LOC_Os0612100 protein, a representative of the M-class gene in rice, was developed using I-TASSER (Zhang 2008) and showed highest homology with human mTERF1 proteins (3N6S, TM-score > 0.77) in the PDB database. Similar to the structure of the human mTERF proteins, LOC_Os0612100 is a modular protein containing four mTERF modules, less than in human, each comprising two or three tandem α-helices. A left-handed superhelical module was observed, with a positively charged groove at its surface (fig 8, supplementary figure S7). Its structure was similar to that of other RNA/DNA-binding proteins such as HEAT (Sibanda et al. 2010), PUF (Edwards et al. 2001), TAL (Deng et al. 2012) and PPR (Yin et al. 2013). Combined with the evidence that *Arabidopsis* mTERF6 specifically interacts with a chloroplast RNA (cpRNA) sequence as indicated by *in vitro* bacterial one-hybrid screening, electrophoretic mobility shift assays and co-immunoprecipitation experiments (Romani et al. 2015), these findings suggest that M-class mTERF proteins might also act as RNA/DNA-binding proteins.

**Figure 8.**
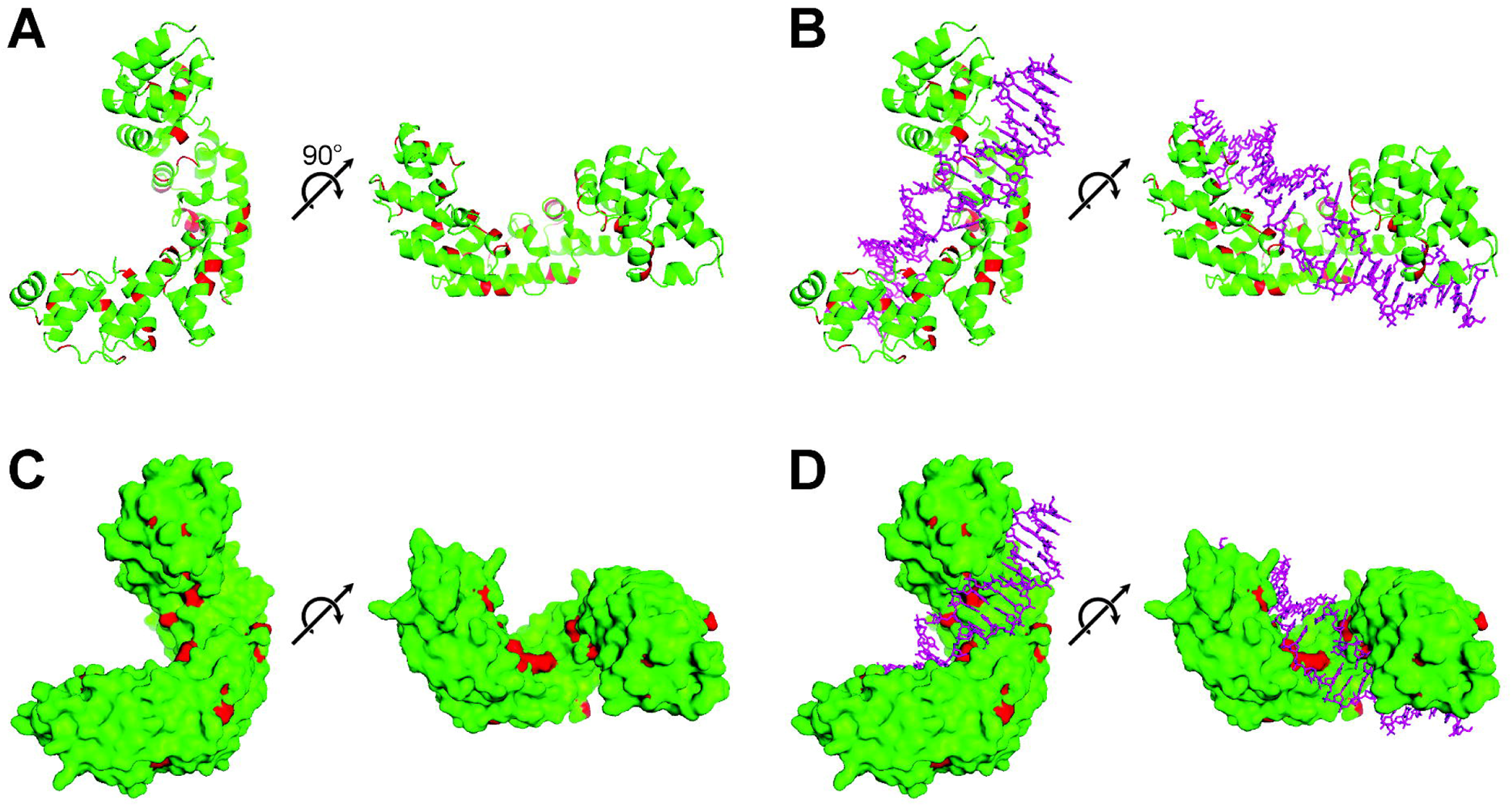
Positively selected sites of M-class mTERF proteins involved in nucleic acid-binding. The tertiary structure of LOC_Os0612100, a rice mTERF protein, was developed using I-TASSER (Zhang 2008) based on homology modeling and displayed in the different forms by PyMOL. The positively selected sites are in red and the DNA strands are marked in magenta (*B*, *D*).

*LOC_Os0612100* belongs to the Osa_2 gene set, which consists of 35 significantly positively selected sites with a posterior probability of > 95% and 21 sites with a posterior probability of > 99%. We mapped the positively selected sites onto the deduced protein structure of LOC_Os0612100 and found that 12 of the 35 sites were located on loops, whereas the others were all tucked into helices (fig. 8). Approximate 43 amino acid residues of the LOC_Os0612100 were predicted by I-TASSER (Zhang 2008) to be putative nucleic acid-binding sites, a result generated by sequence comparison with human mTERF1. Furthermore, eight of these sites were identified as positively selected sites (fig 8, supplementary figure S8). It is speculated that rapid evolution of the M-class genes occurred in higher plants to adapt to changes in substrate.

### M-class *mTERF* genes are associated with mitochondrial genome variation

In plants, non-M-class *mTERF* genes encode virtually all chloroplast-targeted mTERF proteins and a few mitochondria-targeted mTERF proteins. However, the present study has shown that mitochondria-targeting non-M-class *mTERF* genes form orthologous clades (fig. 2). To date, only eight plant *mTERFs* have been characterized, which all belong to the non-M-class subfamily and play important roles in the regulation of expression of chloroplast or mitochondrial genes (Quesada 2016). The gain in the number of M-class genes in higher plants thus provides gene that act on the mitochondria, implying that the birth and mass expansion of M-class genes might have occurred after mitochondrial genomic variation in plants. Therefore, we analyzed the correlation between the number of *mTERF* genes and the sizes of plant nuclear, chloroplast and mitochondrial genomes. As shown in fig. 9, the total number of *mTERF* genes (r = 0.608, *P* < 0.05) and M-class genes (r = 0.681, *P* < 0.05) were directly correlated with variation in the corresponding mitochondrial genome and not with that of the nuclear or chloroplast genomes. The mitochondrial genomes have significantly changed in size from unicellular green algae to flowering plants, ranging from ~16 kb (*C. reinhardtii*) to ~ 4 megabasepairs (Mbs) (*A. trichopoda*) (Rice et al. 2013). The high number of M-class *mTERF* genes in *P. abies*, *A. trichopoda*, and *C. sativus* are associated with the increase in size of the mitochondrial genomes, since no recent WGD events have occurred in these plants (Alverson et al. 2011; Nystedt et al. 2013; Rice et al. 2013). The DNA replication and gene expression machineries of plant mitochondria are relatively complex, particularly those related to gene expression, regulation of RNA transcription, RNA editing, RNA decay and translation (Liere and Borner 2011). Therefore, larger mitochondrial genomes will have a relatively high level of complexity. Based on these findings, we assume that the rapid evolution and positive selection of M-class genes were driven by the changes in plant mitochondrial genomes, in combination with the complexity of their gene expression machinery that influences plant evolution to precisely coordinate the interactions between the nucleus and mitochondria.

**Figure 9.**
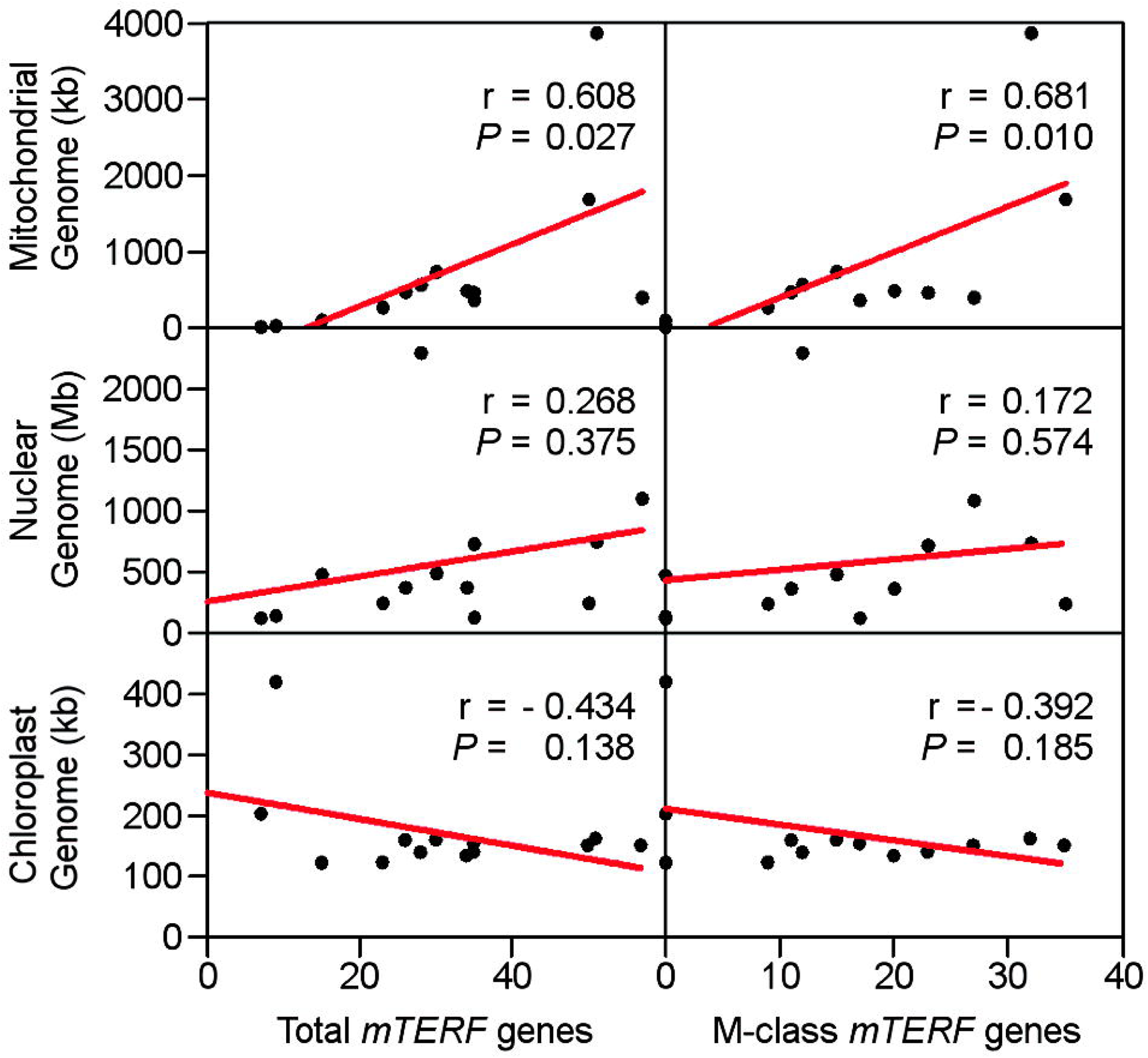
Correlation between *mTERF* gene expansion and plant genome variation. Analysis of the relationship between *mTERF* genes and nuclear or mitochondrial genomes was performed by using IBM SPSS Statistics 20 (IBM Corp. Released 2011. IBM SPSS Statistics for Windows, Version 20.0. Armonk, NY: IBM Corp) and only species with available genome information were included in this study.

## Discussion

Previous investigations on the systematic identification and expression analysis of the *mTERF* gene family have shown that there are more *mTERF* genes in plants than animals, and higher plants have a higher number of *mTERF* genes than lower plants (Kleine 2012; Zhao et al. 2014). Nevertheless, a comprehensive evolutionary estimation of the number of *mTERF* genes in different plants has not been conducted to date. The rapid expansion of sequence databases has facilitated in the identification of putative *mTERF* genes in different organisms. Here, on the basis of their protein properties and phylogenetic relationships, we have identified a novel subfamily of *mTERF* genes (M-class genes) in 18 plant species, ranging from algae to flowering plants that could be distinguished from previously identified non M-class genes.

Non-M-class genes from different species share independent evolution clades and in each of their subclades, only few *mTERF* genes belong to one certain plant (fig. 2), indicating that non M-class genes within the same clades have conserved function and possibly underwent strong purifying selection. All *mTERF* genes previously identified in lower plants belong to the non-M-class subfamily, some of which might represent the ancestral *mTERF* genes in plants. M-class genes are specific to higher plants and consist of group VII *mTERF* genes that have been earlier classified in *Arabidopsis* (Babiychuk et al. 2011) and group VIII-IX *mTERF* genes in maize (Zhao et al. 2014), which encode proteins that have several distinct motifs, as mined in the MEME webserver (Bailey et al. 2009). M-class *mTERF* genes account for more than half of the *mTERF* genes in higher plants and most of these are clustered in relatively narrow chromosomal regions (supplementary figure S2). Since plant gene families distributed in a cluster are mostly explained by tandem duplication, such as type I MADS box genes (Nam et al. 2004), *SKP1* genes (Kong et al. 2007), LRR genes (Meyers et al. 2003), PPR genes (Schmitz-Linneweber and Small 2008; Wang et al. 2006) and glutathione S-transferase (GST) genes (Lan et al. 2009), the increase in M-class genes was likely also due to tandem duplication (fig. 4), an important means of gene expansion resulting from unequal crossing-over and gene conversions. The rapid evolution and positive selection of duplicated M-class genes was observed in several well-annotated genomes, since massive numbers of M-class genes in these plants tend to form species-specific subclades or miniclades(fig. 5). After gene duplication, new copies then face the “to be or not to be” evolutionary fate. Three alternative outcomes for duplicated genes include gene reservation (subfunctionalization or neofunctionalization), silence or loss (nonfunctionalization) (Lynch and Conery 2000). In the ZmC5_1 cluster comprising *ZmTERF18*, *ZmTERF19* and *ZmTERF20* in maize, *ZmTERF20* is silenced by degenerative mutations (nonfunctionalization) and *ZmTERF18* shows a fairly low expression level among a variety of maize tissues, a different expression pattern from that of *ZmTERF19.* A similar scenario has been observed in the *Arabidopsis* AtC1_1 cluster, where M-class genes showed apparent expression divergence in different plant organs (Kleine 2012). These findings indicate that tandemly duplicated M-class genes undergo “birth-to-death” selection as proposed by Nei and Rooney (2005), although we have not experimentally investigated the function of duplicated M-class genes yet. The higher birth rate of M-class genes provides more opportunities to select for the proper divergent function during plant evolution. The “Birth-to-death” evolution model has been surveyed in type I MADS box genes (Nam et al. 2004), *SKP1* genes (Kong et al. 2004), LRR genes (Bergelson et al. 2001; Michelmore and Meyers 1998) and PPR genes (Wang et al. 2006).

The first *mTERF* gene, *hsmTERF1*, was identified in human as a mtDNA-binding factor that promotes the termination of transcription (Kruse et al. 1989). The first characterized plant *mTERF* genes included the *MOC1* gene in *C. reinhardtii* (Schönfeld et al. 2004). *hsmTERF1* and *MOC1* play similar roles in mtDNA transcription termination. The hsmTERF1 protein can stop mtDNA transcription from the H1 site at the 3’-termini of 16S rRNA by directly binding to a DNA sequence within the tRNA^Leu(UUR)^ (Fernandez-Silva et al. 1997; Kruse et al. 1989), whereas MOC1 acts as a mtDNA transcription termination factor by specifically binding to an octanucleotide sequence within the mitochondrial rRNA-coding module S3 (Wobbe and Nixon 2013). The evolutionarily conserved transcription termination activity of *hsmTERF1* and *MOC1* is probably explained by the observed similarities in their mitochondrial genome transcription machinery. In the small mitochondrial genomes of human (~17kb) and *C. reinhardtii* (~16kb), mtDNA is transcribed bidirectionally to produce two long primary transcripts, followed by RNA processing to yield mature mRNAs (Schönfeld et al. 2004; Taanman et al. 1999; Wobbe and Nixon 2013). These *mTERF* genes are required to regulate mtRNA transcription and maturation in human and *C. reinhardtii*.

One explanation for the higher number of *mTERF* genes in plants compared to that in animals is that there is an extra organelle, the plastid, in plants in addition to the mitochondria (Kleine and Leister 2015; Quesada 2016). Besides chloroplasts, the large and complicated mitochondrial genomes may account for the enlargement of the *mTERF* gene family in higher plants. Most mitochondrial genes are transcribed as independent transcriptional units except for the 18S rRNA and 5S rRNA, which are co-transcribed and further spliced into mature rRNAs in higher plants (Gagliardi and Binder 2007). Moreover, organellar RNA metabolism involved in gene expression regulation is unexpectedly complicated in higher plants (Barkan 2011; Hammani and Giegé 2014). The differences in mtDNA transcription mechanism and RNA process at the posttranscriptional level between higher and lower plants would certainly influence the roles of *mTERF* genes in higher plants. To date, a total of seven *mTERF* genes have been explored experimentally in higher plants, of which *Arabidopsis BSM/RUG2/mTERF4*, *mTERF6*, *MDA1/mTERF9*, *SOLDAT10*and maize *Zm-mTERF4* encode chloroplast proteins, whereas the other two *mTERF* genes, *Arabidopsis SHOT1* and *mTERF15*, encode mitochondria-targeted proteins (Babiychuk et al. 2011; Hammani and Barkan 2014; Hsu et al. 2014; Kim et al. 2012; Meskauskiene et al. 2009; Robles et al. 2012b; Robles et al. 2015; Sun et al. 2016b). Nevertheless, these *mTERF* genes all belong to the non-M-class gene subfamily. *Arabidopsis mTERF6* is the only gene that has been determined to be capable of terminating transcription of plastid DNA by binding to its plastid DNA target site *in vitro* (Romani et al. 2015). Mutation of *mTERF6* disrupts the translation and maturation of chloroplast rRNAs, which in turn impedes plastid development, ultimately resulting in seedling lethality (Romani et al. 2015). The putative transcription termination function inherited within *mTERF6* may be due to the existence of an rRNA operon and homolog, *mTERF6*, to *hsmTERF1* since they are included in a subclade in a previously reported ML tree (Zhao et al. 2014). For other well-known *mTERF* genes, no evident DNA- or RNA-binding and transcription termination activities were observed. *Arabidopsis BSM/RUG2/mTERF4* and maize *Zm-mTERF4* are orthologs and are required for RNA splicing of group II introns in the respective chloroplasts (Babiychuk et al. 2011; Hammani and Barkan 2014). *Arabidopsis mTERF15* plays an important role in mitochondrial *nad2* intron 3 splicing and is essential for normal plant growth and development (Hsu et al. 2014). *MDA1/mTERF9*, *SOLDAT10*and *SHOT1* have been shown to be involved in stress response and resistance by regulating expression of organellar genes (Kim et al. 2012; Meskauskiene et al. 2009; Robles et al. 2012b; Robles et al. 2015). Alterations in organellar gene expression induced by these *mTERF* mutants influence the activity of nuclear genes via a retrograde signaling pathway (Kim et al. 2012; Kleine and Leister 2016; Meskauskiene et al. 2009; Sun et al. 2016b). Therefore, these *mTERF* genes are required for organellar gene expression regulation and are essential for normal plant growth and development. The importance of these genes explains the purifying selection on non-M-class genes to a certain degree.

Given that duplicated M-class *mTERF* genes form species-specific clades that differ from non-M-class clades and have undergone rapid divergent selection, we can infer that M-class genes have probably evolved novel roles compared to those of the non M-class genes. The distinct roles of M-class *mTERF* genes may be related to mitochondrial biogenesis and development since M-class proteins have transit peptides for the mitochondria and their expansion in higher plants is associated with variations in mitochondrial genomes (fig. 9). mtDNA recombination occurs frequently in plants and has greatly changed the content and size of mitochondrial genomes (Logan 2006), causing that relatively differences were observed even within the same species (Allen et al. 2007). In plants, each mitochondrial gene always has more than one transcript because of their various promoters and complicated regulation of mtDNA transcription (Gagliardi and Binder 2007). The enlargement of the mitochondrial genome due to mtDNA recombination also affects the flanking sequences of mitochondrial genes, which further alters their expression. The extensive duplication of M-class genes under diversifying selection probably compensates for the alteration of mitochondrial genomes since positively selected sites identified in different plants or gene sets are different (supplementary table S2). Meanwhile, several positively selected sites within the M-class mTERF proteins are predicted to be involved in nucleic acid-binding activity. Similar to husmTERF1, M-class mTERF proteins have a superhelical structure that is theoretically capable to bind nucleic acids (fig. 8). It is speculated that M-class genes might be involved in mtDNA transcription or RNA processing by binding to nucleic acids or being recruited into a functional complex similar to that observed in *BSM/RUG2* (Babiychuk et al. 2011). Screening of mutant libraries did not detect any visible mutant phenotype caused by M-class *mTERF* genes and this may be due to the following reasons: (i) One copy of the duplicated M-class gene can functionally complement the loss of the other copy because of their similarity in protein sequence; and (ii) visible phenotypic changes in M-class mutants are dependent on specific conditions such as the particular type of cytoplasm and environmental stimulus.

Mitochondria play a vital role in the generation of energy and diverse metabolic intermediates for many cellular events in most eukaryotic cells (Logan 2006). The maintenance of structural integrity and fundamental functions of mitochondria are regulated by the nuclear genome. The conflict between nuclear and mitochondrial genomes causes diseases in animals and CMS in flowering plants. CMS has been used to investigate host-parasite relationships between nuclear and mitochondrial genomes (Sato Fujii et al. 2011; Touzet and Budar 2004) in which *RFL* genes are positively selected. M-class *mTERF* and *RFL* genes in higher plants are highly similar in terms of classification, distribution in a cluster, compartmentalization in organelle, higher order protein structure, nucleic acid-binding activity and rapid evolution and diversifying selection (Dahan and Mireau 2013; Fujii et al. 2011; Lurin et al. 2004; Schmitz-Linneweber and Small 2008; Wang et al. 2006; Yin et al. 2013). In the CMS system, positively selected residues within the PPR motif of RFL proteins are probably associated with rapid changes in chimeric mitochondrial genes that are related to male sterility (Fujii et al. 2011). Accordingly, rapid positive selection on M-class *mTERF* genes is probably associated with the variation in mitochondrial genomes, providing additional molecular evidence for the host-parasite relationship between nuclear and mitochondrial genomes. Therefore, in addition to the important roles in controlling gene expression in organelles, *mTERF* genes may be used as a molecular tool for investigating the retrograde signaling pathway and in better understanding the “arms-race” relationship between nuclear and mitochondrial genomes.

## Materials and Methods

### Sequences and Alignment

Identification of *mTERF* sequences in *A. thaliana*, *O. sativa ssp.* Japonica, and *Z. mays* was performed as previously described (Kleine 2012; Zhao et al. 2014). The genome sequence data and gene annotation of the following plant species for *mTERF* gene identification were retrieved from Phytozome v9.0 (http://www.phytozome.net/), *C. reinhardtii* (Merchant et al. 2007), *V. carteri* (Prochnik et al. 2010), *P. patens* (v1.6) (Rensing et al. 2008), *S. moellendorffii* (Banks et al. 2011), *B. distachyon* (Vogel et al. 2010), *S. bicolor* (v2.1) (Paterson et al. 2009), *V. vinifera* (Jaillon et al. 2007), *C. papaya* (Ming et al. 2008), *A. lyrata* (Hu et al. 2011), *C. sativus* (Huang et al. 2009), *G. max* (Schmutz et al. 2010), *M. truncatula* (Young et al. 2011), and *P. trichocarpa* (Tuskan et al. 2006). Gene models for *A. trichopoda* (v1.0) (Chamalaet al. 2013) were downloaded from the Amborella Genome Database (http://amborella.org/), and high-confidence gene annotation data for *P. abies* (Picea1.0) (Nystedtet al. 2013) were obtained from ConGenIE (http://congenie.org/).

To identify the putative *mTERF* genes in the plant genomes, the hmmsearch program in HMMER 3.0 package (Eddy 1998) was used to detect mTERF motifs in deduced protein sequences using the hidden Markov model PF02536 in the Pfam database (Punta et al. 2012) as described elsewhere (Zhao et al. 2014). Hits with an E-value under default inclusion threshold were collected, and the conserved mTERF motifs were investigated in the SMART database (Letunic et al. 2012). Mis-annotation gene models were corrected or removed, including those with no start codon, incorrect start codon, unsequenced genomic gap spanning, truncation, or gene fusion. Only mTERF proteins of > 100 amino acids were included in the analysis. The *mTERF* gene models were renamed in *A. trichopoda* (with “Atr” instead of “evm_27.model.AmTr_v1.0”), *C. papaya* (with “Cp_” instead of “evm.model.”), *S. moellendorffii* (with “Sm” as a prefix to the primary names given in Phytozome v9.0) and *A. lyrata* (with “Al_” as a prefix).

The complete amino acid sequences of mTERF proteins were aligned using MUSCLE v3.8.31 (Edgar 2004). The resulting alignment was manually refined by visual inspection in CINEMA 5 (http://aig.cs.man.ac.uk/research/utopia/cinema/cinema.php).

### Phylogenetic Analysis

ClustalW v2.0.8 (Larkin et al. 2007) was used to calculate distance trees using the NJ method with default parameters. RAxML v7.2.8 (Stamatakis 2006) was used to construct ML inference-based trees with the BLOSUM62 amino acid substitution model (Henikoff and Henikoff 1992). Bootstrap testing was performed with 500 samples to search for the best ML scoring tree. Phylogenetic trees were displayed and analyzed using FigTree v1.4.0 (http://tree.bio.ed.ac.uk/software/figtree/).

### Establishing Gene Sets for PAML Analysis

*mTERF* genes encoded in well-annotated genomes of *A. thaliana*, *A. lyrata*, *O. sativa ssp.* Japonica, and *P. trichocarpa* were collected and used for diversifying selection analysis. For the M-class *mTERF* genes, paralogous gene sets were constructed based on their phylogeny (fig. 5). The codons were aligned using PAL2NAL v14 (Suyama et al. 2006) based on the corresponding protein alignments.

### Testing for Positive Selection

The dN/dS ratio (ω) was calculated using the codeml program in PAML v4.7 (Yang 2007). Positive selection was detected using a likelihood ratio test (LRT) with two codon substitution model pairs M1 v.s. M2 and M7 v.s. M8 as described elsewhere (Yang et al. 2000). For the M1-M2 model pair, M1 is a neutral model that assumes two discrete codon substitutions, ω = 0 (purifying selection) or ω = 1 (neutral selection), and M2 is the positive-selection model with an extra ω > 1. For the M7-M8 model pair, M7 is a neutral model that assumes a continuous β distribution that restricts ω to the interval between 0 and 1, and M8 is the alternative model that adds an extra category with a ω > 1. M1 and M7 were compared as null models with M2 and M8, respectively, to test whether M2 or M8 fits the data. Positively selected sites were identified under M8 using the BEB approach as implemented in the codeml program (Yang et al. 2005). To verify which of the models best fit the data, likelihood ratio tests were performed by comparing twice the difference in log likelihood values between the pairs of the models using a χ^2^ distribution (Yang et al. 2000).

### Modeling of the Tertiary Structures of M-class mTERF proteins

Plant *mTERF* genes have homologs in humans for which the crystal structures of the proteins have been solved. The tertiary structures of one rice M-class mTERF protein without a transit peptide were predicted based on their homologs using I-TASSER (Zhang 2008). PyMOL v1.6.0.0 (The PyMOL Molecular Graphics System, Version 1.6.0.0 Schrödinger, LLC.) was used to map positive-selection sites to the 3D structure of the M-class mTERF protein and to display the three-dimensional (3D) structure.

## Acknowledgments

We are grateful to the members of the Zheng lab for helpful discussions on the results of this manuscript. We thank Dr. Hanhui Kuang (Huazhong Agricultural University, China) and Dr. David Jackson (Cold Spring Harbor Laboratory, USA) for critically reviewing the manuscript. This work was supported by the national key research and development program of China (2016YFD0100303), the National Basic Research Program of China (973 Program) (2014CB138203) and the grants from the National Natural Science Foundation of China (31171565 to Hailin Xiao, 31500253 to Yanxin Zhao).

## Notes

### Competing Interest Statement

The authors have declared no competing interest.

