## Supplementary Tables for "Expansion and Adaptive Evolution of the *mTERF* Gene Family in Plants"

**Supplementary Table S1. Detailed information of plant *mTERF* genes identified in this study.**

| Gene models[a] | Genbank accession[b] | Introns[c] | ORF | Protein[d] Length (aa) | MW (Da) | pI | Subcellular location[e] | mTERF motif[f] | M-class[g] | Note[h] |
|---|---|---|---|---|---|---|---|---|---|---|
| ***Chlamydomonas reinhardtii* (7)** | | | | | | | | | | |
| *Cre14.g611200.t1.2* | - | 5 | 621 | 206 | 22026.40 | 9.36 | C | 2 | | *MOC5* |
| *Cre12.g560750.t1.3* | XP_001703146.1 | 6 | 1689 | 562 | 58954.87 | 5.14 | M | 6 | | *MOC3* |
| *Cre12.g542500.t1.2* | XP_001693924 | 4 | 756 | 251 | 27462.49 | 10.00 | M | 3 | | *MOC1* |
| *Cre10.g445100.t1.3* | XP_001690319 | 6 | 642 | 213 | 23305.43 | 8.53 | C | 1 | | |
| *Cre10.g427000.t1.2* | XP_001702553 | 12 | 1500 | 499 | 53835.71 | 5.30 | M | 5 | | *MOC2* |
| *Cre09.g408050.t1.3* | - | 8 | 1824 | 607 | 61808.16 | 5.94 | M | 4 | | *MOC4* |
| *Cre07.g330650.t1.2* | XP_001690624.1 | 4 | 690 | 229 | 25206.97 | 8.37 | C | 1 | | *MOC6* |
| ***Volvox carteri* (9)** | | | | | | | | | | |
| *Vocar20012395m* | - | 5 | 708 | 235 | 25901.10 | 9.71 | C/- | 1 | | |
| *Vocar20011849m* | XP_002956517 | 4 | 642 | 213 | 22810.42 | 9.35 | C/M | 2 | | |
| *Vocar20011696m* | XP_002954087 | 7 | 1467 | 488 | 53656.03 | 5.92 | C/- | 6 | | |
| *Vocar20011615m* | XP_002954086 | 7 | 1587 | 528 | 58211.65 | 8.88 | C/M | 5 | | |
| *Vocar20011606m* | XP_002954117 | 1 | 1974 | 657 | 68859.13 | 11.17 | S/S | 6 | | |
| *Vocar20009503m* | XP_002958787 | 2 | 1068 | 355 | 38217.20 | 7.08 | _/_ | 3 | | |
| ***Vocar20006656m*** | - | 3 | 837 | 278 | 29673.09 | 10.12 | M/- | 4 | | |
| *Vocar20004652m* | XP_002954428 | 5 | 1224 | 407 | 44014.33 | 5.67 | M/- | 5 | | |
| *Vocar20002501m* | XP_002946108 | 10 | 2985 | 995 | 105925.64 | 9.38 | _/_ | 9 | | |
| ***Physcomitrella patens* (15)** | | | | | | | | | | |
| *Pp1s89_222V6.1* | XP_001767054 | 3 | 1566 | 521 | 57770.26 | 7.06 | C/C | 7 | | |
| *Pp1s88_175V6.1* | XP_001766931 | 3 | 960 | 319 | 35755.12 | 8.54 | _/_ | 2 | | |
| *Pp1s81_201V6.1* | XP_001766050 | 0 | 1368 | 455 | 51497.93 | 8.93 | _/C | 6 | | |
| *Pp1s64_149V6.1* | XP_001763927 | 2 | 1878 | 625 | 70534.46 | 9.38 | _/_ | 10 | | |
| *Pp1s36_216V6.1* | XP_001759211 | 0 | 1764 | 587 | 65246.51 | 4.7 | C/C | 9 | | |
| *Pp1s33_375V6.1* | XP_001758712 | 6 | 1887 | 628 | 71332.23 | 8.85 | _/S | 10 | | |

| | | | | | | | | | | |
|---|---|---|---|---|---|---|---|---|---|---|
| *Pp1s31_149V6.1* | XP_001758215 | 3 | 1437 | 478 | 54009.57 | 9.63 | _/_ | 7 | | |
| *Pp1s258_35V6.1* | XP_001779463 | 2 | 1422 | 473 | 53539.13 | 9.06 | _/_ | 7 | | |
| *Pp1s251_76V6.1* | - | 2 | 1593 | 530 | 59358.69 | 8.78 | ER/M | 9 | | |
| *Pp1s25_200V6.1* | XP_001757032 | 6 | 2889 | 962 | 109315.02 | 6.13 | C/M | 8 | | |
| *Pp1s219_66V6.1* | - | 0 | 2010 | 669 | 75152.11 | 9.27 | C/C | 10 | | |
| *Pp1s21_14V6.1* | XP_001756049 | 0 | 1677 | 558 | 62569.84 | 5.37 | C/_ | 9 | | |
| *Pp1s18_200V6.1* | XP_001755526 | 4 | 1641 | 546 | 61080.73 | 9.4 | _/M | 8 | | |
| *Pp1s16_107V6.1* | - | 3 | 732 | 243 | 27071.38 | 9.4 | _/_ | 2 | | |
| *Pp1s136_28V6.1* | XP_001771323 | 5 | 1800 | 599 | 67623.14 | 9.24 | C/_ | 7 | | |
| ***Selaginella moellendorffii* (13)** | | | | | | | | | | |
| *Sm92242* | XP_002970034 | 0 | 1545 | 514 | 57828.53 | 5.31 | C/C | 9 | | |
| *Sm103816* | XP_002975492 | 0 | 1446 | 481 | 53933.53 | 5.84 | C/_ | 8 | | |
| *Sm77318* | XP_002961886 | 1 | 975 | 324 | 37589.45 | 9.39 | _/_ | 7 | | |
| *Sm83040* | XP_002964972 | 0 | 1041 | 346 | 38367.35 | 9.36 | C/C | 7 | | |
| *Sm165772* | XP_002963159 | 5 | 1374 | 457 | 51866.64 | 9.22 | _/_ | 8 | | |
| ***Sm74633*** | XP_002960914 | 2 | 1263 | 420 | 46899.09 | 9.56 | M/C | 6 | | |
| *Sm444400* | XP_002979970 | 0 | 1728 | 575 | 65459.51 | 9.36 | C/C | 7 | M | |
| *Sm403419* | XP_002961964 | 0 | 1512 | 503 | 56271.35 | 9.6 | M/M | 6 | | |
| *Sm408106* | XP_002967240 | 1 | 1992 | 663 | 74541.35 | 9.98 | _/_ | 5 | | |
| *Sm419725* | XP_002980015 | 4 | 1311 | 436 | 49485.15 | 9.86 | M/M | 6 | | |
| *Sm410501* | XP_002969433 | 1 | 1998 | 665 | 72296.1 | 5.78 | M/M | 3 | | |
| *Sm431071* | XP_002992936 | 5 | 954 | 317 | 35954.56 | 9.24 | _/_ | 2 | | |
| ***Sm58739*** | XP_002980051 | 0 | 666 | 221 | 24225.86 | 8.26 | _/_ | 2 | | |
| ***Picea abies* (60)** | | | | | | | | | | |
| *MA_39589g0010* | - | 0 | 1413 | 470 | 53985.97 | 6.94 | _/_ | 9 | | high_confidence |
| *MA_15876g0010* | - | 0 | 1629 | 542 | 61109.03 | 5.45 | _/_ | 10 | | high_confidence |
| *MA_78196g0010* | - | 0 | 1629 | 542 | 61109.03 | 5.45 | S/_ | 10 | | high_confidence |
| *MA_10434956g0010* | - | 2 | 2148 | 715 | 79872.97 | 9.46 | _/_ | 11 | | high_confidence |

| | | | | | | | | | | |
|---|---|---|---|---|---|---|---|---|---|---|
| *MA_104598g0010* | - | 0 | 1137 | 378 | 42307.29 | 9.36 | S/M | 6 | M | high_confidence |
| *MA_9454422g0010* | - | 0 | 1092 | 363 | 41069.67 | 9.42 | C/C | 6 | | high_confidence |
| *MA_371670g0010* | - | 1 | 1314 | 437 | 49851.22 | 9.31 | _/_ | 7 | | high_confidence |
| *MA_453913g0010* | - | 1 | 1035 | 344 | 39222.42 | 9.69 | _/_ | 8 | | high_confidence |
| *MA_280158g0010* | - | 0 | 1008 | 335 | 37410 | 9.63 | S/_ | 6 | M | high_confidence |
| *MA_10429715g0010* | - | 0 | 1041 | 346 | 39598.02 | 9.44 | _/_ | 7 | M | high_confidence |
| *MA_2589g0010* | - | 0 | 1404 | 467 | 52668.74 | 9.67 | S/M | 7 | M | high_confidence |
| *MA_112645g0010* | - | 0 | 633 | 210 | 24472.84 | 9.44 | _/M | 5 | | high_confidence |
| *MA_10426325g0010* | - | 0 | 1038 | 345 | 39259.81 | 9.41 | _/_ | 6 | M | high_confidence |
| *MA_126865g0020* | - | 0 | 1782 | 593 | 67613.2 | 9.48 | M/M | 6 | | high_confidence |
| *MA_926365g0010* | - | 3 | 1233 | 410 | 46683.17 | 8.93 | C/C | 5 | | high_confidence |
| *MA_178046g0010* | - | 0 | 1062 | 353 | 40051.9 | 9.69 | S/M | 7 | M | high_confidence |
| *MA_122549g0010* | - | 0 | 1068 | 355 | 41134.3 | 10.02 | _/_ | 7 | M | high_confidence |
| *MA_125123g0020* | - | 2 | 1575 | 524 | 58656.79 | 8.78 | _/_ | 8 | | high_confidence |
| *MA_209070g0010* | - | 0 | 1128 | 375 | 42114.51 | 10.01 | C/_ | 7 | M | high_confidence |
| *MA_593288g0010* | - | 1 | 1254 | 365 | 41462.65 | 9.48 | _/M | 5 | M | high_confidence |
| *MA_493420g0010* | - | 0 | 690 | 229 | 26482.1 | 9.7 | _/_ | 5 | M | high_confidence |
| *MA_34619g0010* | - | 0 | 918 | 305 | 35155.67 | 10.07 | M/M | 7 | M | high_confidence |
| *MA_45859g0010* | - | 0 | 996 | 331 | 37334.76 | 9.64 | M/ER | 5 | M | high_confidence |
| *MA_6307g0010* | - | 0 | 660 | 219 | 25475.66 | 6.36 | _/_ | 5 | M | high_confidence |
| *MA_14049g0010* | - | 0 | 657 | 218 | 25141.6 | 9.03 | _/_ | 4 | M | high_confidence |
| *MA_10429074g0010* | - | 0 | 1683 | 360 | 40924.65 | 9.66 | S/_ | 5 | M | high_confidence |
| *MA_92135g0010* | - | 0 | 1038 | 345 | 39768.43 | 9.65 | _/C | 6 | M | high_confidence |
| *MA_590299g0010* | - | 0 | 924 | 307 | 34892.61 | 9.01 | _/_ | 3 | M | high_confidence |
| *MA_10041174g0010* | - | 0 | 1008 | 335 | 38179.79 | 9.11 | _/_ | 6 | M | high_confidence |
| *MA_139814g0010* | - | 0 | 990 | 329 | 37602.14 | 9.58 | M/M | 6 | M | high_confidence |
| *MA_10427666g0020* | - | 0 | 828 | 275 | 31488.76 | 9.5 | _/_ | 6 | M | high_confidence |
| *MA_33244g0010* | - | 0 | 1107 | 368 | 42042.07 | 9.46 | _/C | 6 | M | high_confidence |
| *MA_86824g0010* | - | 0 | 1185 | 394 | 44892.51 | 9.57 | M/M | 6 | M | high_confidence |
| *MA_10430524g0010* | - | 0 | 783 | 260 | 29240.17 | 8.91 | _/_ | 6 | M | high_confidence |

| | | | | | | | | | | |
|---|---|---|---|---|---|---|---|---|---|---|
| *MA_205355g0010* | - | 0 | 777 | 258 | 29742.78 | 9.74 | M/M | 6 | M | high_confidence |
| *MA_161434g0010* | - | 0 | 777 | 258 | 29873.06 | 9.77 | M/_ | 6 | M | high_confidence |
| *MA_10083329g0010* | - | 0 | 672 | 223 | 25352.75 | 9.34 | _/_ | 5 | M | high_confidence |
| *MA_161180g0010* | - | 4 | 1311 | 436 | 49898.83 | 9.81 | _/_ | 5 | | high_confidence |
| *MA_124772g0010* | - | 1 | 1041 | 346 | 38921.63 | 9.77 | C/C | 6 | M | high_confidence |
| *MA_10432566g0010* | - | 0 | 1554 | 517 | 57998.7 | 9.25 | _/_ | 6 | | high_confidence |
| *MA_760114g0010* | - | 0 | 768 | 255 | 29264.61 | 9.51 | _/_ | 6 | M | high_confidence |
| ***MA_187605g0020*** | - | 0 | 513 | 170 | 19558.98 | 9.76 | _/_ | 3 | M | high_confidence |
| ***MA_10209606g0010*** | - | 0 | 513 | 170 | 19608.89 | 9.66 | _/_ | 3 | M | high_confidence |
| *MA_804585g0010* | - | 1 | 1029 | 342 | 38241.25 | 9.57 | M/M | 5 | M | high_confidence |
| *MA_118798g0010* | - | 1 | 675 | 224 | 25468.6 | 9.69 | _/_ | 5 | M | high_confidence |
| ***MA_26585g0010*** | - | 0 | 438 | 145 | 16658.73 | 8.93 | _/M | 3 | M | high_confidence |
| *MA_385472g0010* | - | 2 | 1248 | 415 | 46385.46 | 8.7 | _/_ | 4 | | high_confidence |
| *MA_10435408g0010* | - | 0 | 708 | 235 | 26611.79 | 9.03 | _/_ | 5 | M | high_confidence |
| ***MA_28540g0010*** | - | 0 | 534 | 177 | 20444.96 | 9.62 | _/_ | 3 | M | high_confidence |
| *MA_628537g0010* | - | 1 | 669 | 222 | 25444.75 | 9.42 | _/_ | 4 | M | high_confidence |
| *MA_21437g0010* | - | 0 | 987 | 328 | 37046.49 | 10.1 | C/C | 6 | M | high_confidence |
| ***MA_17909g0010*** | - | 0 | 438 | 145 | 16725.9 | 9.81 | _/M | 3 | M | high_confidence |
| *MA_10430524g0020* | - | 2 | 681 | 226 | 25362.72 | 8.9 | _/_ | 3 | M | high_confidence |
| *MA_335595g0010* | - | 0 | 621 | 206 | 22805.29 | 9.14 | M/M | 3 | | high_confidence |
| *MA_10436083g0040* | - | 0 | 687 | 228 | 25383.32 | 9.72 | C/C | 4 | M | high_confidence |
| *MA_10425770g0010* | - | 0 | 765 | 254 | 28622.53 | 8.37 | _/_ | 1 | | high_confidence |
| *MA_88386g0010* | - | 0 | 627 | 208 | 23925.02 | 9.14 | _/_ | 3 | M | high_confidence |
| *MA_805240g0010* | - | 0 | 723 | 240 | 27406.06 | 9.71 | _/_ | 3 | | high_confidence |
| *MA_191994g0010* | - | 1 | 693 | 230 | 26488.15 | 8.33 | C/M | 1 | | high_confidence |
| ***MA_162625g0010*** | - | 0 | 357 | 118 | 13375.54 | 9.3 | S/_ | 1 | | high_confidence |
| ***Amborella trichopoda* (51)** | | | | | | | | | | |
| *ATr_scaffold00010.279* | ERM94303.1 | 0 | 1962 | 653 | 74100.85 | 9.2 | M/M | 9 | | *AmTrmTERF1* |
| *ATr_scaffold00025.37* | ERN12323.1 | 0 | 1554 | 517 | 58761.94 | 6.1 | _/_ | 10 | | *AmTrmTERF2* |

| | | | | | | | | | | |
|---|---|---|---|---|---|---|---|---|---|---|
| *ATr_scaffold00025.247* | ERN12533.1 | 6 | 1776 | 591 | 67619.2 | 8.58 | C/C | 9 | | *AmTrmTERF3* |
| *ATr_scaffold00010.300* | ERM94324.1 | 0 | 2232 | 743 | 83958.85 | 9.15 | C/C | 12 | M | *AmTrmTERF4* |
| *ATr_scaffold00010.287* | ERM94311.1 | 0 | 2232 | 743 | 83953.88 | 9.18 | C/C | 13 | M | *AmTrmTERF5* |
| *ATr_scaffold00088.129* | ERM99614.1 | 4 | 1536 | 511 | 58852.83 | 9.37 | C/C | 7 | | *AmTrmTERF6* |
| *ATr_scaffold00092.56* | ERM99200.1 | 0 | 1089 | 362 | 41487.07 | 6.19 | C/_ | 6 | | *AmTrmTERF7* |
| *ATr_scaffold00005.115* | ERN06841.1 | 3 | 1437 | 478 | 53867.4 | 7.11 | _/C | 7 | | *AmTrmTERF8* |
| *ATr_scaffold00010.280* | ERM94304.1 | 0 | 1218 | 405 | 45439.25 | 9.23 | S/_ | 7 | M | *AmTrmTERF9* |
| *ATr_scaffold00030.14* | ERN16000.1 | 5 | 2001 | 666 | 75288 | 9.07 | _/_ | 8 | | *AmTrmTERF10* |
| *ATr_scaffold00022.29* | ERN11430.1 | 0 | 1026 | 341 | 38785.81 | 9.49 | _/_ | 8 | | *AmTrmTERF11* |
| *ATr_scaffold00010.282* | ERM94306.1 | 0 | 1218 | 405 | 45795.65 | 9.36 | S/M | 6 | M | *AmTrmTERF12* |
| *ATr_scaffold00030.182* | ERN16168.1 | 0 | 1101 | 366 | 42034.04 | 8.92 | M/M | 6 | | *AmTrmTERF13* |
| *ATr_scaffold00045.13* | ERN01934.1 | 0 | 963 | 320 | 36516.9 | 9.64 | _/_ | 6 | M | *AmTrmTERF14* |
| *ATr_scaffold00010.281* | ERM94305.1 | 0 | 1011 | 336 | 38313 | 9.54 | _/C | 7 | M | *AmTrmTERF15* |
| *ATr_scaffold00024.308* | ERN11309.1 | 2 | 1773 | 590 | 66993.48 | 9.56 | M/M | 6 | | *AmTrmTERF16* |
| *ATr_scaffold00060.18* | ERM95820.1 | 0 | 1191 | 396 | 45148.48 | 9.22 | S/M | 8 | M | *AmTrmTERF17* |
| *ATr_scaffold00010.284* | ERM94308.1 | 0 | 1203 | 400 | 45329.23 | 9.69 | M/M | 8 | M | *AmTrmTERF18* |
| *ATr_scaffold00010.290* | ERM94314.1 | 0 | 1191 | 396 | 45001.34 | 9.27 | C/ER | 7 | M | *AmTrmTERF19* |
| *ATr_scaffold00010.247* | ERM94271.1 | 0 | 993 | 330 | 38335.96 | 9.41 | _/_ | 6 | M | *AmTrmTERF20* |
| *ATr_scaffold00058.26* | ERN06499.1 | 0 | 1005 | 334 | 38255.66 | 9.22 | C/C | 5 | | *AmTrmTERF21* |
| *ATr_scaffold01360.1* | ERN04397.1 | 0 | 993 | 330 | 38372.1 | 9.5 | _/_ | 6 | M | *AmTrmTERF22* |
| *ATr_scaffold00066.162* | ERN20246.1 | 0 | 1233 | 410 | 46825.75 | 9.58 | C/C | 8 | M | *AmTrmTERF23* |
| *ATr_scaffold00010.153* | ERM94177.1 | 0 | 1137 | 378 | 43555.26 | 9.5 | S/M | 7 | M | *AmTrmTERF24* |
| *ATr_scaffold00010.248* | ERM94272.1 | 0 | 993 | 330 | 38464.11 | 9.38 | _/_ | 6 | M | *AmTrmTERF25* |
| *ATr_scaffold00871.1* | ERM99306.1 | 0 | 1137 | 378 | 43745.43 | 9.47 | S/M | 7 | M | *AmTrmTERF26* |
| *ATr_scaffold00084.23* | ERN02296.1 | 1 | 1533 | 510 | 58470.05 | 8.4 | C/_ | 7 | | *AmTrmTERF27* |
| *ATr_scaffold00010.249* | ERM94273.1 | 0 | 987 | 328 | 38132.84 | 9.53 | _/_ | 6 | M | *AmTrmTERF28* |
| *ATr_scaffold00010.250* | ERM94273.1 | 0 | 987 | 328 | 8132.84 | 9.53 | _/_ | 6 | M | *AmTrmTERF29* |
| *ATr_scaffold00010.289* | ERM94313.1 | 0 | 1206 | 401 | 45926.86 | 9.52 | M/M | 7 | M | *AmTrmTERF30* |
| *ATr_scaffold00010.68* | ERM94092.1 | 0 | 1020 | 339 | 38916.5 | 9.39 | _/C | 7 | M | *AmTrmTERF31* |
| *ATr_scaffold00038.116* | ERN14616.1 | 0 | 1005 | 334 | 38635.4 | 9.8 | S/C | 7 | M | *AmTrmTERF32* |

| | | | | | | | | | | |
|---|---|---|---|---|---|---|---|---|---|---|
| *ATr_scaffold00010.262* | ERM94286.1 | 0 | 1182 | 393 | 45098.98 | 9.76 | S/ER | 7 | M | *AmTrmTERF33* |
| *ATr_scaffold00005.32* | ERN06758.1 | 2 | 933 | 310 | 35096.82 | 5.69 | _/ER | 5 | | *AmTrmTERF34* |
| *ATr_scaffold00066.163* | ERN20247.1 | 0 | 1167 | 388 | 44391.28 | 9.79 | S/C | 7 | M | *AmTrmTERF35* |
| *ATr_scaffold00010.69* | ERM94093.1 | 0 | 918 | 305 | 34132.05 | 9.52 | M/M | 6 | M | *AmTrmTERF36* |
| *ATr_scaffold02857.1* | ERN08273.1 | 0 | 801 | 266 | 30784.19 | 9.51 | _/_ | 6 | M | *AmTrmTERF37* |
| *ATr_scaffold00009.432* | ERM95245.1 | 0 | 1506 | 501 | 58023.62 | 9.03 | M/M | 5 | | *AmTrmTERF38* |
| *ATr_scaffold00020.26* | ERN11838.1 | 1 | 705 | 234 | 26971.23 | 6.67 | _/ER | 5 | | *AmTrmTERF39* |
| *ATr_scaffold00126.6* | ERM97101.1 | 0 | 516 | 171 | 19947.52 | 9.42 | _/_ | 4 | M | *AmTrmTERF40* |
| *ATr_scaffold00029.212* | ERN09581.1 | 0 | 1713 | 570 | 65233.21 | 9.35 | C/M | 4 | | *AmTrmTERF41* |
| *ATr_scaffold00010.388* | ERM94412.1 | 0 | 582 | 193 | 22944.63 | 9.33 | _/_ | 4 | M | *AmTrmTERF42* |
| *ATr_scaffold00126.8* | ERM97103.1 | 0 | 831 | 276 | 30766.91 | 9.57 | C/C | 5 | M | *AmTrmTERF43* |
| *ATr_scaffold00033.165* | ERN14326.1 | 0 | 1758 | 585 | 67281.95 | 9.42 | C/ER | 5 | | *AmTrmTERF44* |
| *ATr_scaffold00027.143* | ERN10839.1 | 0 | 1707 | 568 | 65346.82 | 8.75 | M/M | 5 | | *AmTrmTERF45* |
| *ATr_scaffold00029.208* | ERN09577.1 | 0 | 1953 | 650 | 74476.83 | 9 | _/_ | 5 | | *AmTrmTERF46* |
| *ATr_scaffold00126.7* | ERM97102.1 | 0 | 645 | 214 | 23768.25 | 9.45 | _/_ | 4 | M | *AmTrmTERF47* |
| *ATr_scaffold00612.2* | ERM98877.1\| | 0 | 867 | 288 | 32295.96 | 10 | S/ER | 5 | M | *AmTrmTERF48* |
| *ATr_scaffold00010.72* | ERM94096.1 | 0 | 681 | 226 | 25040.16 | 9.59 | C/C | 4 | M | *AmTrmTERF49* |
| *ATr_scaffold00126.5* | ERM97100.1 | 0 | 420 | 139 | 15518.92 | 9.27 | C/C | 2 | M | *AmTrmTERF50* |
| *ATr_scaffold00010.283* | ERM94307.1 | 0 | 318 | 105 | 12131.44 | 10.42 | _/_ | 2 | M | *AmTrmTERF51* |
| ***Brachypodium distachyon* (41)** | | | | | | | | | | |
| *Bradi1g02400.1* | XP_003562785 | 1 | 660 | 219 | 24166.71 | 8.58 | C/C | 1 | | |
| *Bradi1g06490.1* | XP_003558151 | 0 | 942 | 313 | 34178.83 | 8.57 | C/C | 5 | | |
| ***Bradi1g16160.1*** | XP_003559733 | 0 | 1173 | 390 | 43346.89 | 9.96 | M/M | 6 | M | |
| *Bradi1g23400.1* | XP_003560036 | 4 | 1503 | 500 | 57120.29 | 9.2 | M/C | 7 | | |
| *Bradi1g27300.1* | XP_003563104 | 0 | 1716 | 571 | 64540.96 | 6.53 | M/M | 5 | | |
| *Bradi1g45320.1* | XP_003560893 | 0 | 1242 | 413 | 45816.28 | 9.5 | M/M | 5 | M | |
| *Bradi1g45330.1* | - | 0 | 1143 | 380 | 41909.49 | 10.09 | M/M | 4 | M | |
| ***Bradi1g45340.1*** | XP_003564059 | 0 | 1176 | 391 | 43503.3 | 9.67 | M/M | 4 | M | |
| *Bradi1g45347.1* | XP_003564060 | 0 | 1164 | 387 | 42913.55 | 9.76 | C/M | 4 | M | |

| | | | | | | | | | |
|---|---|---|---|---|---|---|---|---|---|
| ***Bradi1g45360.1*** | XP_003564061 | 0 | 1155 | 384 | 41569.33 | 9.34 | M/M | 5 | M |
| *Bradi1g57920.1* | XP_003557618 | 5 | 1827 | 608 | 69119.27 | 9.17 | C/C | 8 | |
| *Bradi1g58197.1* | XP_003557632 | 1 | 1182 | 393 | 42755.91 | 9.35 | M/M | 5 | M |
| *Bradi1g61810.1* | XP_003557871 | 0 | 939 | 312 | 33954.43 | 9.79 | M/C | 5 | |
| *Bradi1g71747.1* | - | 1 | 1047 | 348 | 38350.73 | 9.46 | M/M | 3 | M |
| *Bradi2g25372.1* | XP_003566245 | 0 | 1152 | 383 | 41816.26 | 9.75 | M/M | 4 | M |
| *Bradi2g25374.1* | XP_003566246 | 0 | 1149 | 382 | 41790.09 | 9.79 | M/M | 3 | M |
| *Bradi2g25830.1* | XP_003568480 | 0 | 1515 | 504 | 56764.62 | 5.42 | C/C | 9 | |
| *Bradi2g25837.1* | XP_003568493 | 0 | 1185 | 394 | 43726.31 | 9.37 | M/M | 5 | M |
| *Bradi3g00780.1* | XP_003570782 | 0 | 1152 | 383 | 41875.86 | 9.4 | M/M | 5 | M |
| *Bradi3g00816.1* | - | 1 | 987 | 328 | 36379.92 | 9.83 | _/_ | 5 | M |
| *Bradi3g00830.1* | XP_003570704 | 0 | 1167 | 388 | 42571.57 | 9.58 | M/M | 4 | M |
| **Bradi3g00840.1** | - | 0 | 507 | 168 | 19063.01 | 9.74 | _/_ | 2 | M |
| *Bradi3g00847.1* | XP_003570800 | 0 | 1158 | 385 | 42312.42 | 9.72 | M/M | 4 | M |
| *Bradi3g04517.1* | XP_003574975 | 0 | 1164 | 387 | 42260.3 | 9.55 | M/M | 5 | M |
| *Bradi3g40280.2* | XP_003574768 | 0 | 1005 | 334 | 37660.32 | 9.35 | _/_ | 7 | |
| *Bradi3g40580.1* | XP_003572404 | 0 | 1947 | 648 | 73310.08 | 9.35 | M/M | 6 | |
| *Bradi3g46830.1* | XP_003575221 | 0 | 846 | 281 | 30490.78 | 9.91 | C/_ | 3 | |
| *Bradi3g48010.1* | XP_003572693 | 3 | 1500 | 499 | 56289.56 | 9.3 | M/M | 7 | |
| *Bradi3g59140.1* | - | 0 | 783 | 260 | 29211.65 | 9.19 | _/_ | 3 | M |
| *Bradi3g59147.1* | - | 0 | 1158 | 385 | 43043.65 | 9.56 | M/M | 4 | M |
| *Bradi3g59160.1* | - | 1 | 1188 | 395 | 44163.92 | 9.62 | M/M | 4 | M |
| *Bradi3g59167.1* | XP_003573096 | 0 | 1191 | 396 | 43310.62 | 9.58 | S/ER | 4 | M |
| *Bradi3g60150.1* | XP_003570710 | 1 | 1490 | 496 | 55238.6 | 9.03 | M/M | 9 | |
| *Bradi4g02280.1* | XP_003577836 | 0 | 1161 | 386 | 42459.68 | 9.47 | M/M | 6 | M |
| *Bradi4g07150.1* | XP_003575506 | 1 | 912 | 303 | 34628.71 | 7.27 | M/M | 1 | |
| *Bradi4g20547.1* | XP_003576092 | 0 | 1143 | 380 | 42029.36 | 9.86 | M/M | 4 | M |
| *Bradi4g23197.1* | XP_003577739 | 0 | 1146 | 381 | 41948.83 | 9.64 | M/M | 5 | M |
| *Bradi4g23217.1* | - | 1 | 1077 | 358 | 38821.96 | 9.72 | _/_ | 6 | M |
| *Bradi4g23227.1* | XP_003576092 | 0 | 1149 | 382 | 42009.93 | 9.58 | M/M | 4 | M |

| | | | | | | | | | |
|---|---|---|---|---|---|---|---|---|---|
| *Bradi4g37917.1* | XP_003576836 | 6 | 1797 | 598 | 66459.24 | 9.51 | C/C | 9 | |
| *Bradi5g23380.1* | XP_003579363 | 1 | 1209 | 402 | 45121.34 | 10.32 | M/M | 4 | M |
| ***Oryze sativa* ssp. japonica (34)** | | | | | | | | | |
| *LOC_Os01g27690.1* | BAD61377 | 0 | 420 | 139 | 15136.36 | 11.45 | C/C | None | M |
| *LOC_Os02g36780.1* | BAD34257 | 0 | 849 | 282 | 30633.12 | 11.44 | C/M | 4 | |
| *LOC_Os02g39040.1* | EEE57321 | 3 | 1473 | 490 | 55220.09 | 9.44 | C/C | 7 | |
| *LOC_Os02g51450.2* | BAD15633 | 0 | 1200 | 399 | 43868.09 | 9.07 | M/_ | 4 | M |
| *LOC_Os02g51460.1* | BAD15634 | 0 | 1149 | 382 | 42682.18 | 9.67 | M/M | 5 | M |
| *LOC_Os02g54200.1* | BAD19540 | 3 | 1725 | 574 | 64072.3 | 9.15 | M/M | 9 | |
| *LOC_Os03g24590.1* | NP_001050155 | 0 | 906 | 301 | 32895.3 | 10.12 | M/C | 5 | |
| *LOC_Os03g57149.1* | NP_001051481 | 0 | 933 | 310 | 33928.58 | 8.93 | C/C | 6 | |
| *LOC_Os03g62240.1* | BAG90218 | 1 | 645 | 214 | 24457.1 | 9.13 | _/M | 1 | |
| *LOC_Os04g54510.1* | BAG98323 | 0 | 1182 | 393 | 43640.33 | 10.45 | M/M | 5 | M |
| *LOC_Os05g33440.1* | AAT85213 | 0 | 1182 | 393 | 43598.83 | 10.09 | M/M | 5 | M |
| *LOC_Os05g33460.2* | BAG93261 | 1 | 1200 | 399 | 43892.18 | 9.87 | M/M | 6 | M |
| *LOC_Os05g33500.1* | BAG95733 | 0 | 1527 | 508 | 56686.4 | 5.26 | C/C | 9 | |
| *LOC_Os05g34160.1* | BAG86643 | 0 | 1188 | 395 | 43006.39 | 10.56 | M/ER | 4 | M |
| *LOC_Os06g12040.1* | BAD38189 | 0 | 1179 | 392 | 43272.86 | 9.68 | M/M | 5 | M |
| *LOC_Os06g12050.1* | NP_001174667 | 1 | 975 | 324 | 35590.1 | 9.95 | M/M | 3 | M |
| ***LOC_Os06g12060.1a*** | - | 0 | 1221 | 406 | 44563 | 9.72 | M/M | 4 | M |
| ***LOC_Os06g12060.1b*** | - | 0 | 1335 | 444 | 47978.11 | 10.24 | M/M | 4 | M |
| *LOC_Os06g12070.1* | BAD38194 | 0 | 1182 | 393 | 43116.33 | 9.67 | M/M | 4 | M |
| *LOC_Os06g12080.1* | BAD38196 | 0 | 1137 | 378 | 41477.35 | 9.72 | M/M | 4 | M |
| *LOC_Os06g12100.1* | BAD37286 | 0 | 1215 | 404 | 44309.68 | 9.39 | M/M | 4 | M |
| *LOC_Os06g12110.1* | BAD37287 | 0 | 1182 | 393 | 43599.98 | 9.75 | M/M | 4 | M |
| *LOC_Os07g04230.1* | BAC83514 | 5 | 1827 | 608 | 68425.36 | 9.36 | M/C | 8 | |
| *LOC_Os07g22670.1* | BAC20725 | 0 | 1728 | 575 | 64576.29 | 8.96 | M/M | 5 | |
| *LOC_Os07g24090.1* | BAC84114 | 0 | 1227 | 408 | 45899.69 | 9.92 | M/M | 5 | M |
| *LOC_Os07g39430.1* | BAC55604 | 4 | 1512 | 503 | 57654.05 | 8.87 | C/ER | 7 | |

| | | | | | | | | | | |
|---|---|---|---|---|---|---|---|---|---|---|
| *LOC_Os08g40430.1* | BAC56785 | 0 | 1002 | 333 | 37416.14 | 9.39 | _/_ | 7 | | |
| *LOC_Os08g40630.1* | BAC57325 | 0 | 1911 | 636 | 72283.08 | 9.57 | M/M | 4 | | |
| *LOC_Os09g38720.1* | BAD45958 | 9 | 1956 | 651 | 73192.03 | 9.41 | C/ER | 10 | | |
| *LOC_Os11g09990.1* | BAG91551 | 0 | 1221 | 406 | 43935.5 | 9.69 | M/M | 4 | M | |
| *LOC_Os11g10000.1* | AAX95040 | 0 | 1377 | 458 | 50376.38 | 9.53 | M/M | 6 | M | |
| *LOC_Os11g10040.1* | BAG98866 | 0 | 1179 | 392 | 42837.48 | 9.49 | M/M | 4 | M | |
| *LOC_Os11g14130.1* | AAX92923 | 1 | 1254 | 417 | 45602.82 | 9.69 | C/M | 5 | M | |
| *LOC_Os12g30610.1* | ABA98287 | 0 | 999 | 332 | 37774.87 | 10.75 | M/M | 2 | | |
| ***Zea mays* (28)** | | | | | | | | | | |
| *GRMZM2G168665* | DAA45475.1 | 0 | 912 | 303 | 32.57 | 10.44 | C/C | 5 | | *ZmTERF1* |
| *GRMZM2G061542* | DAA48102.1 | 0 | 1950 | 649 | 73.88 | 8.92 | M/M | 6 | | *ZmTERF2* |
| *GRMZM2G034217* | ACG24064.1 | 0 | 1185 | 394 | 44.2 | 9.99 | M/M | 5 | M | *ZmTERF3* |
| *GRMZM2G159766* | ACG24523.1 | 0 | 1203 | 400 | 43.67 | 10 | M/M | 6 | M | *ZmTERF5* |
| *GRMZM2G170137* | NP_001140442.1 | 1 | 645 | 214 | 23.65 | 8.93 | _/C | 1 | | *ZmTERF6* |
| *GRMZM2G060114* | ACG38629.1 | 0 | 1170 | 389 | 43.06 | 9.37 | M/M | 4 | M | *ZmTERF8* |
| *GRMZM2G130773* | DAA41241.1 | 4 | 1527 | 508 | 57.95 | 8.98 | M/M | 7 | | *ZmTERF9* |
| *GRMZM2G177019* | NP_001144077.1 | 0 | 990 | 329 | 36.92 | 10.33 | M/M | 2 | | *ZmTERF10* |
| *GRMZM2G023257* | NP_001151049.1 | 0 | 1167 | 388 | 42.98 | 9.93 | M/M | 5 | M | *ZmTERF11* |
| *GRMZM2G113181* | NP_001152615.1 | 0 | 1005 | 334 | 37.34 | 9.63 | _/_ | 7 | | *ZmTERF12* |
| *GRMZM2G312806* | NP_001141758.1 | 0 | 840 | 279 | 31.16 | 10.49 | M/M | 4 | | *ZmTERF13* |
| *GRMZM2G000610* | NP_001145894.1 | 0 | 1470 | 489 | 54.68 | 8.09 | M/M | 9 | | *ZmTERF14* |
| *GRMZM2G119921* | ACL_52777.1 | 0 | 1188 | 395 | 44.62 | 9.3 | M/M | 4 | M | *ZmTERF15* |
| *GRMZM2G395850* | NP_001143033.1 | 0 | 1068 | 355 | 40.12 | 9.47 | M/M | 4 | M | *ZmTERF16* |
| *GRMZM2G024550* | ACF87053 | 3 | 1461 | 486 | 54.33 | 9.24 | C/C | 7 | | *ZmTERF17* |
| *GRMZM2G017355* | AFW73453.1 | 0 | 1176 | 391 | 42.83 | 9.31 | M/C | 4 | M | *ZmTERF18* |
| *GRMZM2G017429* | NP_001169079.1 | 0 | 1173 | 390 | 42.43 | 9.11 | ER/S | 4 | M | *ZmTERF19* |
| *GRMZM2G161146* | DAA48102.1 | 0 | 1116 | 371 | 40.83 | 9.76 | _/M | 5 | M | *ZmTERF21* |
| *GRMZM2G012999* | AFW77957.1 | 0 | 1212 | 403 | 45.16 | 9.96 | M/M | 6 | M | *ZmTERF22* |
| *GRMZM2G426154* | NP_001152167.1 | 5 | 1836 | 611 | 68.67 | 9.52 | C/C | 9 | | *ZmTERF23* |

| | | | | | | | | | | |
|---|---|---|---|---|---|---|---|---|---|---|
| *GRMZM2G142150* | NP_001169565.1 | 6 | 1839 | 612 | 68.43 | 9.34 | C/C | 10 | | *ZmTERF24* |
| *GRMZM2G062910* | NP_001147866.1 | 0 | 1725 | 574 | 64.75 | 6.48 | M/M | 4 | | *ZmTERF25* |
| *GRMZM2G325350* | NP_001130068.1 | 0 | 1155 | 384 | 42.9 | 9.47 | M/M | 6 | M | *ZmTERF26* |
| *GRMZM2G029933* | NP_001149660.1 | 0 | 1485 | 494 | 55.23 | 5.7 | C/C | 9 | | *ZmTERF27* |
| ***GRMZM2G068462*** | - | 0 | 510 | 169 | 18.58 | 9.94 | _/_ | 3 | | *ZmTERF28* |
| *GRMZM2G157716* | AFW88319.1 | 0 | 906 | 301 | 32.47 | 9.81 | C/M | 5 | | *ZmTERF29* |
| **-** | NP_001152154.1 | - | 873 | 290 | 33.25 | 9.81 | ER/C | 6 | M | *ZmTERF30* |
| **-** | ACG24302.1 | - | 999 | 332 | 37.71 | 9.12 | _/_ | 7 | | *ZmTERF31* |
| ***Sorghum bicolor* (35)** | | | | | | | | | | |
| *Sobic.001G021100.1.p* | XP_002466136 | 1 | 909 | 302 | 33627.21 | 10.28 | M/ER | 3 | M | |
| ***Sobic.001G029000.1.p*** | XP_002463602 | 0 | 1197 | 398 | 44101.75 | 10.05 | M/M | 5 | M | |
| *Sobic.001G032600.1.p* | XP_002463622 | 0 | 1188 | 395 | 43454.25 | 10.02 | M/M | 6 | M | |
| ***Sobic.001G033900.1.p*** | XP_002466189 | 0 | 1194 | 397 | 43498.89 | 9.71 | M/M | 6 | M | |
| *Sobic.001G034000.1.p* | XP_002466190 | 1 | 1224 | 407 | 45169.56 | 9.54 | M/M | 6 | M | |
| *Sobic.001G066600.1.p* | XP_002463781 | 0 | 975 | 324 | 35651.36 | 8.19 | C/C | 6 | | |
| *Sobic.001G176900.1.p* | XP_002466862 | 0 | 1170 | 389 | 42805.99 | 9.68 | M/M | 4 | M | |
| *Sobic.001G364100.1.p* | XP_002467825 | 0 | 942 | 313 | 33521.3 | 10.55 | C/C | 5 | | |
| ***Sobic.002G028400.1.p*** | XP_002459325 | 5 | 1809 | 602 | 67808.12 | 9.53 | C/C | 8 | | |
| *Sobic.002G179800.1.p* | XP_002460105 | 0 | 1185 | 394 | 42762.28 | 9.72 | M/M | 5 | M | |
| *Sobic.002G296100.1.p* | XP_002460661 | 6 | 1830 | 609 | 67994.96 | 9.55 | C/C | 10 | | |
| *Sobic.002G305100.1.p* | XP_002460710 | 0 | 1731 | 576 | 65454.05 | 7.66 | M/M | 4 | | |
| *Sobic.002G354600.1.p* | XP_002460925 | 4 | 1521 | 506 | 57730.89 | 8.76 | M/M | 7 | | |
| ***Sobic.003G110500.1.p*** | XP_002455361 | 0 | 999 | 332 | 36667.99 | 10.14 | M/ER | 3 | M | |
| *Sobic.003G199700.1.p* | XP_002458051 | 0 | 1149 | 382 | 42321.49 | 9.73 | M/M | 4 | M | |
| *Sobic.003G389500.1.p* | - | 0 | 1149 | 382 | 41899.27 | 9.82 | M/M | 6 | M | |
| *Sobic.003G396100.1.p* | XP_002458954 | 0 | 1167 | 388 | 43401.74 | 9.89 | M/M | 4 | M | |
| *Sobic.004G069400.1.p* | XP_002453425 | 0 | 1188 | 395 | 44739.27 | 9.84 | M/M | 4 | M | |
| ***Sobic.004G190900.1.p*** | XP_002454060 | 0 | 840 | 279 | 30955.52 | 10.97 | M/M | 4 | | |
| *Sobic.004G206200.1.p* | XP_002452407 | 3 | 1479 | 492 | 55524.62 | 9.19 | M/C | 7 | | |

| | | | | | | | | | |
|---|---|---|---|---|---|---|---|---|---|
| *Sobic.004G235300.1.p* | XP_002454274 | 0 | 1143 | 380 | 41932.41 | 9.4 | M/M | 5 | M |
| *Sobic.004G235500.1.p* | XP_002454275 | 0 | 1167 | 388 | 43300.59 | 9.7 | M/M | 5 | M |
| *Sobic.004G235600.1.p* | XP_002454276 | 0 | 1173 | 390 | 42483.5 | 9.38 | C/ER | 4 | M |
| *Sobic.004G235700.1.p* | XP_002454277 | 0 | 1185 | 394 | 43146.93 | 8.87 | C/M | 4 | M |
| *Sobic.004G305100.1.p* | XP_002454582 | 0 | 1167 | 388 | 42861.5 | 9.9 | M/M | 5 | M |
| *Sobic.004G321700.1.p* | XP_002454669 | 0 | 1470 | 489 | 54663.88 | 7.52 | M/M | 9 | |
| *Sobic.006G183300.1.p* | XP_002446953 | 3 | 966 | 321 | 35532.6 | 9.29 | C/C | 5 | M |
| *Sobic.007G114700.1.p* | XP_002444251 | 1 | 1182 | 393 | 43780.82 | 9.63 | M/M | 6 | M |
| *Sobic.007G206100.1.p* | XP_002444756 | 0 | 1005 | 334 | 37411.97 | 9.42 | _/_ | 7 | |
| *Sobic.008G137800.1.p* | XP_002442362 | 0 | 984 | 327 | 36625.6 | 10.44 | M/M | 2 | |
| ***Sobic.009G131600.1.p*** | XP_002439769 | 0 | 1497 | 498 | 55924.65 | 5.61 | C/C | 9 | |
| *Sobic.009G133400.1.p* | XP_002439781 | 0 | 1197 | 398 | 44381.15 | 9.91 | M/M | 6 | M |
| *Sobic.010G041100.1.p* | XP_002436491 | 0 | 1155 | 384 | 41688.91 | 9.74 | M/M | 6 | M |
| ***Sobic.010G091200.1.p*** | XP_002436739 | 0 | 1209 | 402 | 44691.05 | 9.27 | M/M | 5 | M |
| *Sobic.010G201900.1.p* | XP_002437330 | 0 | 1188 | 395 | 44100.92 | 10.01 | M/M | 5 | M |
| ***Vitis vinifera* (30)** | | | | | | | | | |
| *GSVIVT01001819001* | CBI40062 | 4 | 1395 | 464 | 52254.83 | 9.08 | M/_ | 7 | |
| *GSVIVT01008120001* | XP_002265430 | 0 | 1356 | 451 | 52130.3 | 9.26 | M/M | 6 | |
| *GSVIVT01009012001* | CBI19144 | 3 | 1278 | 425 | 48794.25 | 9.82 | C/M | 7 | M |
| *GSVIVT01010970001* | CBI32246 | 1 | 1227 | 408 | 46296.29 | 9.62 | M/M | 7 | |
| *GSVIVT01011061001* | CBI32316 | 1 | 1596 | 531 | 60031.8 | 9.14 | _/_ | 10 | |
| *GSVIVT01012810001* | CBI25532 | 6 | 2160 | 719 | 80381.1 | 8.86 | _/ER | 4 | |
| *GSVIVT01015207001* | CBI27915 | 5 | 1662 | 553 | 63246.22 | 9.19 | M/M | 8 | |
| *GSVIVT01017772001* | CBI26090 | 4 | 1284 | 427 | 49953.82 | 9.44 | _/_ | 6 | M |
| *GSVIVT01021544001* | CBI30943 | 4 | 783 | 260 | 29372.02 | 9.18 | _/C | 3 | |
| *GSVIVT01022213001* | CBI21495 | 0 | 1110 | 369 | 42492.67 | 9.27 | C/_ | 7 | M |
| *GSVIVT01023845001* | CBI37711 | 2 | 1065 | 354 | 40312.58 | 9.45 | C/C | 7 | |
| *GSVIVT01026275001* | CBI29073 | 6 | 1692 | 563 | 65101.63 | 8.86 | S/ER | 4 | |
| *GSVIVT01028380001* | XP_002271898 | 0 | 1239 | 412 | 46689.26 | 9.86 | M/M | 6 | M |

| | | | | | | | | | |
|---|---|---|---|---|---|---|---|---|---|
| ***GSVIVT01028382001a*** | - | 0 | 1233 | 410 | 46831.17 | 9.71 | M/M | 7 | M |
| ***GSVIVT01028382001b*** | - | 0 | 1164 | 387 | 43897.24 | 9.64 | M/_ | 6 | M |
| ***GSVIVT01028382001c*** | - | 0 | 1158 | 385 | 43814.38 | 9.64 | M/M | 6 | M |
| ***GSVIVT01028382001d*** | - | 0 | 1155 | 384 | 43559.08 | 9.56 | M/_ | 5 | M |
| ***GSVIVT01028382001e*** | - | 0 | 1164 | 387 | 43565.04 | 9.6 | M/M | 6 | M |
| ***GSVIVT01028383001a*** | - | 0 | 1173 | 390 | 43837.47 | 9.49 | M/M | 7 | M |
| ***GSVIVT01028383001b*** | - | 0 | 1170 | 389 | 43564.57 | 9.51 | M/M | 7 | M |
| ***GSVIVT01028383001c*** | - | 0 | 1206 | 401 | 44789.82 | 9.45 | M/M | 6 | M |
| *GSVIVT01028384001* | CBI37156 | 0 | 1101 | 366 | 41230.9 | 9.41 | M/_ | 6 | M |
| *GSVIVT01029533001* | CBI18428 | 9 | 2427 | 808 | 93392.3 | 9.26 | C/_ | 10 | |
| *GSVIVT01031956001* | XP_003631646 | 0 | 831 | 276 | 31949.09 | 9.47 | C/C | 5 | |
| ***GSVIVT01031970001*** | XP_003631610 | 5 | 2037 | 678 | 78101.24 | 8.86 | C/C | 8 | M |
| *GSVIVT01033517001* | CAN83222 | 0 | 1770 | 589 | 67766.52 | 8.96 | M/M | 5 | |
| *GSVIVT01034475001* | CBI18165 | 1 | 639 | 212 | 23724 | 5.73 | C/ER | 1 | |
| *GSVIVT01036787001* | CBI24192 | 0 | 1233 | 410 | 46683.06 | 10.15 | S/ER | 7 | M |
| *GSVIVT01037780001* | CBI26741 | 5 | 1443 | 480 | 55233.22 | 8.77 | C/C | 6 | |
| ***GSVIVT01038641001*** | CAN80174 | 1 | 1437 | 478 | 53464.26 | 6.81 | C/C | 6 | |
| ***Carica papaya*** **(26)** | | | | | | | | | |
| *Cp_supercontig_1.44* | - | 0 | 861 | 286 | 32234.8 | 10.17 | C/M | 5 | |
| *Cp_supercontig_101.32* | - | 0 | 1671 | 556 | 64304.17 | 8.61 | M/M | 5 | |
| *Cp_supercontig_104.58* | - | 1 | 1305 | 434 | 48900.51 | 9.31 | C/_ | 6 | |
| *Cp_supercontig_13.147* | - | 0 | 1365 | 454 | 51399.87 | 9.4 | M/M | 6 | M |
| *Cp_supercontig_130.3* | - | 0 | 1161 | 386 | 43769.42 | 9.28 | M/M | 6 | M |
| *Cp_supercontig_135.43* | - | 0 | 1737 | 578 | 65732.77 | 9.22 | M/_ | 5 | |
| *Cp_supercontig_139.30* | - | 0 | 924 | 307 | 34461.36 | 9.47 | C/C | 7 | |
| *Cp_supercontig_151.12* | - | 0 | 1524 | 507 | 58020.79 | 8.74 | M/M | 9 | |
| *Cp_supercontig_16.125* | - | 0 | 813 | 270 | 30993.45 | 9.75 | _/M | 4 | M |
| *Cp_supercontig_17.174* | - | 0 | 1188 | 395 | 45126.59 | 9.52 | _/M | 7 | M |
| *Cp_supercontig_171.37* | - | 3 | 1446 | 481 | 54631.84 | 9.1 | M/M | 8 | |

| | | | | | | | | | |
|---|---|---|---|---|---|---|---|---|---|
| *Cp_supercontig_176.24* | - | 6 | 1794 | 597 | 67362.35 | 8.57 | C/_ | 9 | |
| *Cp_supercontig_2154.2* | - | 0 | 1230 | 409 | 46348.74 | 9.82 | M/M | 5 | M |
| ***Cp_supercontig_245.9*** | - | 3 | 1089 | 362 | 41779.59 | 9.54 | _/_ | 4 | M |
| *Cp_supercontig_287.2* | - | 0 | 1122 | 373 | 42849.02 | 9.82 | M/M | 7 | M |
| *Cp_supercontig_3.154* | - | 1 | 1506 | 501 | 56789.11 | 8.58 | C/C | 8 | |
| *Cp_supercontig_36.118* | - | 0 | 1149 | 382 | 43476.15 | 9.85 | M/M | 7 | M |
| *Cp_supercontig_36.174* | - | 0 | 1221 | 406 | 46334.35 | 9.31 | M/M | 6 | M |
| *Cp_supercontig_441.3* | - | 1 | 1113 | 370 | 42256.81 | 9.59 | C/M | 6 | M |
| *Cp_supercontig_60.7* | - | 0 | 1062 | 353 | 40655.84 | 9.34 | _/_ | 7 | |
| *Cp_supercontig_62.107* | - | 0 | 852 | 283 | 32767.18 | 9.78 | C/C | 4 | |
| *Cp_supercontig_62.154* | - | 6 | 1974 | 657 | 75327.69 | 7.26 | C/C | 6 | |
| ***Cp_supercontig_66.48*** | - | 2 | 570 | 189 | 21524.69 | 4.93 | _/_ | 1 | M |
| *Cp_supercontig_9.127* | - | 0 | 1260 | 419 | 46691.39 | 8.7 | _/_ | 8 | |
| *Cp_supercontig_94.76* | - | 1 | 1338 | 445 | 51770.65 | 9.07 | M/M | 8 | |
| *Cp_contig_28389.2* | - | 2 | 1392 | 463 | 53421.76 | 6.26 | M/M | 3 | |
| ***Arabidopsis thaliana* (35)** | | | | | | | | | |
| *AT1G21150.1* | NM_101969 | 0 | 1173 | 390 | 44289.4 | 9.794 | C/C | 5 | M" |
| *AT1G56380.1* | NM_104517 | 2 | 1167 | 388 | 44562.3 | 4.5772 | _/_ | 7 | M |
| *AT1G61960.1* | NM_104876 | 0 | 1374 | 457 | 51477.4 | 10.0324 | M/M | 7 | M |
| *AT1G61970.1* | NM_104877 | 0 | 1257 | 418 | 46951.4 | 10.05 | M/M | 6 | M |
| *AT1G61980.1* | NM_104878 | 0 | 1257 | 418 | 47236.5 | 10.32 | M/M | 6 | M |
| *AT1G61990.1* | NM_104879 | 0 | 1245 | 414 | 46247.7 | 10.23 | M/M | 5 | M |
| *AT1G62010.1* | NM_104881 | 0 | 1248 | 415 | 51564.7 | 9.8057 | M/M | 6 | M |
| *AT1G62085.1* | - | 0 | 1386 | 461 | 51564.7 | 9.8057 | M/M | 7 | M |
| *AT1G62110.1* | NM_104892 | 0 | 1389 | 462 | 52242.3 | 9.769 | M/M | 7 | M |
| *AT1G62120.1* | NM_104893 | 0 | 1314 | 437 | 48939.5 | 10.078 | M/M | 6 | M |
| *AT1G62150.1* | NM_104896 | 0 | 1392 | 463 | 52065.2 | 8.6788 | M/C | 7 | M |
| *AT1G62490.1* | NM_104928 | 1 | 1005 | 334 | 37854.4 | 9.7905 | _/_ | 3 | M |
| *AT1G74120.1* | NM_106072 | 0 | 1338 | 445 | 51030.6 | 10.1353 | M/M | 5 | |

| | | | | | | | | | |
|---|---|---|---|---|---|---|---|---|---|
| *AT1G78930.1* | NM_106542 | 6 | 1776 | 591 | 67563.0 | 8.3097 | C/C | 8 | |
| *AT1G79220.1* | NM_106573 | 0 | 1200 | 399 | 46464.3 | 10.18 | M/ER | 6 | M' |
| *AT2G03050.1* | NM_126357 | 0 | 852 | 283 | 32259.2 | 9.195 | M/M | 5 | |
| *AT2G21710.1* | NM_127741 | 5 | 1926 | 641 | 74744.4 | 8.6933 | C/C | 8 | |
| *AT2G34620.1* | NM_129016 | 1 | 912 | 303 | 34252.7 | 8.2744 | C/C | 6 | |
| *AT2G36000.1* | NM_129159 | 0 | 1002 | 333 | 37952.3 | 8.9732 | C/_ | 5 | |
| *AT2G44020.1* | NM_129964 | 0 | 1524 | 507 | 58073.8 | 8.314 | M/M | 9 | |
| *AT3G18870.1* | NM_112773 | 0 | 825 | 274 | 31431.2 | 10.21 | C/C | 5 | |
| *AT3G46950.1* | NM_114562 | 0 | 1353 | 450 | 51336.0 | 10.17 | M/M | 7 | M |
| *AT3G60400.1* | NM_115904 | 0 | 1677 | 558 | 64561.8 | 8.315 | M/M | 5 | |
| *AT4G02990.1* | NM_116533 | 0 | 1626 | 541 | 61548.4 | 6.7197 | M/_ | 10 | |
| *AT4G09620.1* | NM_117030 | 2 | 639 | 212 | 24009.3 | 6.651 | C/C | 1 | |
| *AT4G14605.1* | - | 3 | 1482 | 493 | 55960.4 | 9.608 | C/_ | 8 | |
| *AT4G19650.1* | - | 3 | 1728 | 575 | 66744.6 | 9.6911 | _/_ | 6 | |
| *AT4G38160.1* | NM_119977 | 0 | 1002 | 333 | 37888.1 | 9.336 | _/_ | 7 | |
| *AT5G06810.1* | NM_120764 | 1 | 3426 | 1141 | 131204.6 | 8.671 | M/_ | 8 | |
| *AT5G07900.1* | NM_120872 | 0 | 1218 | 405 | 45845.3 | 9.487 | C/C | 5 | M" |
| *AT5G23930.1* | NM_122298 | 0 | 1374 | 457 | 51885.3 | 10.3168 | M/M | 7 | M |
| *AT5G45113.1* | NM_148090 | 0 | 1245 | 414 | 48240.6 | 9.633 | _/_ | None | |
| *AT5G54180.1* | NM_124798 | 1 | 1503 | 500 | 56348.9 | 7.5444 | C/ER | 8 | |
| *AT5G55580.1* | NM_124940 | 4 | 1491 | 496 | 57201.2 | 9.1545 | C/C | 6 | |
| *AT5G64950.1* | NM_125894 | 0 | 1176 | 391 | 44696.1 | 10.24 | M/M | 7 | M' |
| ***Arabidopsis lyrata* (46)** | | | | | | | | | |
| ***Al_313019*** | EFH6941 | 0 | 1206 | 401 | 45594.36 | 9.41 | M/M | 7 | M |
| ***Al_315177a*** | - | 0 | 1251 | 416 | 46291.59 | 9.43 | M/M | 7 | M |
| ***Al_315177b*** | - | 0 | 1389 | 462 | 51890.5 | 9.13 | M/M | 7 | M |
| ***Al_315190*** | EFH64314 | 1 | 1338 | 445 | 50400.8 | 9.43 | M/M | 6 | M |
| ***Al_338172*** | EFH64304 | 0 | 1260 | 419 | 46879.44 | 9.7 | M/M | 6 | M |
| *Al_475136* | EFH64311 | 0 | 1245 | 414 | 46778.86 | 9.77 | M/M | 5 | M |

| | | | | | | | | | |
|---|---|---|---|---|---|---|---|---|---|
| *Al_475137* | EFH64312 | 0 | 1260 | 419 | 47459.14 | 9.8 | M/M | 6 | M |
| *Al_476555* | EFH65225 | 0 | 1350 | 449 | 51464.54 | 9.42 | C/M | 6 | |
| *Al_477061* | EFH65479 | 7 | 1779 | 592 | 67716.77 | 6.29 | C/_ | 9 | |
| *Al_477089* | EFH64032 | 0 | 1218 | 405 | 47006.74 | 9.75 | M/ER | 6 | M |
| *Al_481036* | XP_002878567 | 5 | 1842 | 613 | 71696.06 | 9.26 | M/M | 9 | |
| *Al_482425* | XP_002862643 | 1 | 912 | 303 | 34047.92 | 8.73 | C/C | 6 | |
| *Al_482571* | XP_002862184 | 1 | 993 | 330 | 37826.89 | 9 | C/_ | 5 | |
| *Al_484214* | XP_002876855 | 0 | 855 | 284 | 32385.8 | 8.88 | M/C | 5 | |
| ***Al_485072*** | XP_002875821 | 1 | 1302 | 433 | 49394.06 | 9.59 | M/M | 6 | M |
| *Al_487623* | XP_002871294 | 0 | 1215 | 404 | 45614.55 | 9.18 | C/C | 5 | M |
| *Al_489254* | XP_002874153 | 0 | 1365 | 454 | 51775.53 | 9.48 | M/_ | 7 | M |
| *Al_489815* | XP_002874579 | 2 | 645 | 214 | 24154.79 | 6.53 | _/_ | 1 | |
| *Al_490263* | XP_002874884 | 0 | 1605 | 534 | 60789.18 | 6.07 | M/_ | 10 | |
| *Al_490575* | XP_002866785 | 0 | 1020 | 339 | 38645.21 | 9.05 | _/_ | 7 | |
| *Al_495499* | XP_002864311 | 2 | 1503 | 500 | 56501.75 | 6.59 | C/ER | 8 | |
| *Al_495645* | XP_002864399 | 4 | 1482 | 493 | 57224.92 | 9.08 | C/C | 6 | |
| *Al_497370* | XP_002862643 | 1 | 912 | 303 | 34047.92 | 8.73 | C/C | 6 | |
| *Al_497554* | XP_002862184 | 1 | 993 | 330 | 37826.89 | 9 | C/_ | 5 | |
| *Al_861095* | XP_002871231 | 1 | 3435 | 1144 | 131842.34 | 8.93 | C/M | 8 | |
| *Al_893233* | EFH62725 | 0 | 1422 | 473 | 53675.67 | 9.58 | M/M | 8 | M |
| *Al_893270* | EFH62747 | 0 | 1392 | 463 | 52310.99 | 9.02 | C/C | 8 | M |
| *Al_893275* | EFH62750 | 1 | 1440 | 479 | 54296.09 | 9.81 | C/M | 7 | M |
| *Al_893280* | EFH64299 | 0 | 1161 | 386 | 43427.69 | 9.56 | M/M | 7 | M |
| *Al_893283* | EFH64302 | 0 | 1272 | 423 | 47938.02 | 9.86 | M/M | 6 | M |
| ***Al_893289*** | EFH64307 | 0 | 1101 | 366 | 41336.74 | 9.36 | _/ER | 7 | M |
| ***Al_893294*** | EFH64309 | 1 | 663 | 220 | 23223.87 | 9.99 | S/ER | 2 | M |
| ***Al_893298a*** | - | 0 | 1236 | 411 | 45770.63 | 9.69 | M/M | 6 | M |
| ***Al_893298b*** | - | 0 | 1260 | 419 | 47522.57 | 9.43 | M/M | 6 | M |
| *Al_893300* | EFH64315 | 0 | 1284 | 427 | 47997.02 | 9.5 | M/M | 6 | M |
| ***Al_894067*** | EFH64654 | 1 | 1164 | 387 | 43137.06 | 9.64 | M/M | 5 | M |

| | | | | | | | | | |
|---|---|---|---|---|---|---|---|---|---|
| *Al_898263* | XP_002885283 | 0 | 831 | 276 | 31643.84 | 9.68 | C/C | 5 | |
| *Al_903808* | XP_002881945 | 0 | 1527 | 508 | 58259.54 | 8.38 | M/M | 9 | |
| *Al_907562* | XP_002876552 | 0 | 1683 | 560 | 64479.56 | 8.15 | M/M | 5 | |
| ***Al_907730*** | XP_002876637 | 0 | 1308 | 435 | 49531.49 | 9.83 | M/M | 4 | M |
| ***Al_907868*** | XP_002876694 | 0 | 1164 | 387 | 43934.65 | 9.48 | C/M | 5 | M |
| *Al_914700* | XP_002869971 | 3 | 1653 | 550 | 63662.45 | 9.18 | M/_ | 5 | |
| *Al_915404* | XP_002870303 | 2 | 1497 | 498 | 56686.84 | 9.25 | C/_ | 8 | |
| *Al_917019* | XP_002863524 | 1 | 1389 | 462 | 53025.3 | 9.31 | M/M | 5 | |
| ***Al_917622*** | XP_002865578 | 0 | 441 | 146 | 16698.96 | 8.84 | _/M | 3 | M |
| *Al_919822* | XP_002864933 | 1 | 1149 | 382 | 44082.41 | 9.48 | M/M | 6 | M |
| ***Cucumis sativus* (50)** | | | | | | | | | |
| *Cucsa.033690.1* | XP_004143818.1 | 0 | 1575 | 524 | 59679.42 | 6.82 | _/_ | 10 | |
| ***Cucsa.035460.1*** | XP_004169604.1 | 1 | 978 | 325 | 36696.09 | 9.81 | C/M | 6 | M |
| *Cucsa.075910.1* | - | 0 | 1122 | 373 | 42425.95 | 9.61 | C/M | 7 | M |
| *Cucsa.076310.1* | XP_004135362.1 | 0 | 1125 | 374 | 42676.01 | 9.74 | C/M | 7 | M |
| *Cucsa.076320.1* | XP_004147124.1 | 0 | 1125 | 374 | 42782.15 | 9.77 | C/M | 7 | M |
| ***Cucsa.076330.1*** | XP_004135533.1 | 0 | 924 | 307 | 34794.67 | 9.18 | _/_ | 7 | M |
| *Cucsa.076340.1* | XP_004135534.1 | 0 | 1122 | 373 | 42436.64 | 9.59 | C/M | 8 | M |
| ***Cucsa.076460.1*** | XP_004170657.1 | 0 | 1128 | 375 | 43068.56 | 9.53 | C/M | 7 | M |
| *Cucsa.076490.1* | XP_004147125.1 | 0 | 1128 | 375 | 42682.27 | 9.71 | C/M | 7 | M |
| *Cucsa.076510.1* | XP_004168906.1 | 0 | 1131 | 376 | 42271.8 | 9.66 | S/M | 7 | M |
| *Cucsa.076520.1* | XP_004168905.1 | 0 | 1122 | 373 | 42187.68 | 9.86 | C/M | 7 | M |
| ***Cucsa.076540.1*** | XP_004172786.1 | 0 | 1119 | 372 | 42383.68 | 9.76 | C/M | 7 | M |
| ***Cucsa.083730.1*** | XP_004159265.1 | 0 | 957 | 318 | 36632.61 | 8.88 | C/_ | 6 | |
| *Cucsa.088330.1* | XP_004158513.1 | 1 | 651 | 216 | 24487.18 | 8.56 | C/C | 1 | |
| *Cucsa.088410.1* | XP_004138819.1 | 0 | 1158 | 385 | 44591.99 | 9.56 | M/M | 5 | M |
| *Cucsa.100000.1* | XP_004144567.1 | 0 | 987 | 328 | 37605.6 | 9.37 | _/_ | 8 | |
| *Cucsa.101810.1* | XP_004142030.1 | 4 | 1710 | 569 | 64689.41 | 9.22 | S/ER | 7 | |
| ***Cucsa.120010.1*** | - | 0 | 609 | 202 | 22839.18 | 8.29 | _/_ | 5 | |

| | | | | | | | | | |
|---|---|---|---|---|---|---|---|---|---|
| *Cucsa.151720.1* | XP_004145310.1 | 6 | 1785 | 594 | 66843.05 | 8.34 | C/C | 9 | |
| *Cucsa.152130.1* | XP_004145298.1 | 0 | 915 | 304 | 34099.54 | 9.01 | C/C | 6 | |
| *Cucsa.165080.1* | - | 0 | 1122 | 373 | 42531.99 | 9.7 | C/M | 7 | M |
| *Cucsa.165090.1* | XP_004165692.1 | 0 | 1122 | 373 | 42458.85 | 9.92 | C/M | 7 | M |
| ***Cucsa.165480.1*** | - | 1 | 789 | 262 | 30255.99 | 10.03 | C/M | 6 | M |
| *Cucsa.166200.1* | - | 0 | 1122 | 373 | 43023.36 | 9.64 | C/M | 6 | M |
| ***Cucsa.167430.1*** | XP_004135745.1 | 0 | 1362 | 453 | 52270.1 | 9.35 | C/M | 7 | |
| *Cucsa.172140.1* | - | 5 | 882 | 293 | 33541.17 | 9.75 | _/ER | 3 | M |
| *Cucsa.201490.1* | XP_004149163.1 | 0 | 1161 | 386 | 43696.14 | 9.81 | C/C | 7 | M |
| *Cucsa.220350.1* | XP_004162746.1 | 0 | 849 | 282 | 32766.86 | 9.72 | C/ER | 6 | |
| ***Cucsa.240560.1*** | XP_004169767.1 | 5 | 1980 | 659 | 75722.65 | 8.78 | C/C | 9 | |
| *Cucsa.253200.1* | XP_004151613.1 | 0 | 1116 | 371 | 42034.01 | 9.67 | C/M | 6 | M |
| *Cucsa.253240.1* | XP_004173915.1 | 0 | 1080 | 359 | 41048.8 | 9.6 | C/M | 6 | M |
| *Cucsa.253310.1* | XP_004134480.1 | 0 | 1098 | 365 | 41596.43 | 9.73 | C/ER | 6 | M |
| *Cucsa.253320.1* | XP_004159847.1 | 0 | 1098 | 365 | 41989.19 | 10.17 | M/M | 6 | M |
| ***Cucsa.253330.1*** | XP_004159848.1 | 0 | 1095 | 364 | 41640.75 | 9.79 | M/M | 6 | M |
| ***Cucsa.253340.1*** | XP_004134477.1 | 0 | 1107 | 368 | 42025.12 | 9.64 | C/M | 6 | M |
| ***Cucsa.253350.1*** | - | 0 | 1113 | 370 | 42377.87 | 9.9 | M/M | 7 | M |
| ***Cucsa.253370.1*** | XP_004159850.1 | 0 | 1098 | 365 | 42163.93 | 9.66 | M/M | 6 | M |
| ***Cucsa.253380.1a*** | XP_004134476.1 | 0 | 1113 | 370 | 42627.85 | 9.78 | C/M | 6 | M |
| ***Cucsa.253380.1b*** | XP_004134476.1 | 0 | 1095 | 364 | 41142.96 | 9.75 | M/M | 6 | M |
| *Cucsa.263970.1* | XP_004172643.1 | 0 | 1785 | 594 | 68607.94 | 9.06 | S/_ | 3 | |
| *Cucsa.281080.1* | XP_004172367.1 | 0 | 1122 | 373 | 42773.4 | 9.64 | C/ER | 7 | M |
| *Cucsa.281090.1* | XP_004147133.1 | 0 | 1122 | 373 | 42234.74 | 9.6 | C/M | 7 | M |
| *Cucsa.281210.1* | XP_004162047.1 | 0 | 1125 | 374 | 42538.95 | 9.7 | C/M | 7 | M |
| ***Cucsa.284970.1*** | XP_004155384.1 | 0 | 837 | 278 | 31732.31 | 8.23 | S/_ | 6 | |
| *Cucsa.304690.1* | XP_004151387.1 | 4 | 1566 | 521 | 60526.03 | 9.47 | C/C | 6 | |
| ***Cucsa.324410.1*** | XP_004161949.1 | 0 | 1197 | 398 | 45609.5 | 9.27 | M/M | 7 | M |
| *Cucsa.340350.1* | XP_004158334.1 | 0 | 1551 | 516 | 58688.25 | 8.56 | S/M | 9 | |
| ***Cucsa.347310.1*** | XP_004135531.1 | 0 | 882 | 293 | 33581.36 | 9.53 | _/M | 7 | M |

| | | | | | | | | | |
|---|---|---|---|---|---|---|---|---|---|
| ***Cucsa.371610.1*** | XP_004172427.1 | 0 | 738 | 245 | 27393.8 | 9.79 | C/M | 4 | M |
| ***Cucsa.393940.1*** | XP_004161098.1 | 0 | 1209 | 402 | 45951.49 | 9.28 | _/ER | 5 | M |
| ***Glycine max* (57)** | | | | | | | | | |
| ***Glyma01g06010.1*** | XP_003518006.1 | 0 | 849 | 282 | 32046.81 | 9.97 | C/M | 6 | |
| *Glyma02g12120.1* | XP_003518738.1 | 0 | 888 | 295 | 33692.86 | 10.07 | C/ER | 6 | |
| *Glyma02g38800.1* | - | 0 | 1557 | 518 | 58712.29 | 6.32 | _/_ | 10 | |
| *Glyma02g46750.1* | XP_003518542.1 | 4 | 1467 | 488 | 56750.35 | 9.46 | C/C | 6 | |
| *Glyma03g26731.1* | XP_003520435.1 | 0 | 1701 | 566 | 64283.19 | 7.87 | M/M | 4 | |
| *Glyma03g29920.2* | - | 3 | 1008 | 335 | 38233.74 | 9.64 | M/M | 2 | |
| ***Glyma04g32192.1*** | XP_006579258.1 | 0 | 1131 | 376 | 42614.12 | 9.62 | M/M | 5 | M |
| ***Glyma04g40661.1*** | XP_003522526.2 | 0 | 1572 | 523 | 59579.63 | 9.14 | S/ER | 7 | |
| *Glyma05g15170.1* | XP_003524621.1 | 3 | 1443 | 480 | 54536.38 | 9.51 | M/M | 8 | |
| *Glyma05g34550.2* | - | 1 | 1113 | 370 | 42307.53 | 7 | _/_ | 7 | |
| *Glyma06g22340.2* | XP_006582565.1 | 1 | 762 | 253 | 28974.11 | 8.77 | _/_ | 5 | |
| *Glyma07g14330.2* | XP_006583538.1 | 0 | 1791 | 596 | 67678.5 | 9.14 | _/_ | 3 | |
| *Glyma07g37870.1* | XP_006584032.1 | 0 | 1146 | 381 | 43867.71 | 10.04 | S/M | 5 | M |
| *Glyma07g37970.1* | XP_003529583.1 | 0 | 1158 | 385 | 43897.92 | 9.82 | M/M | 6 | M |
| *Glyma08g05110.1* | XP_003530919.1 | 0 | 1500 | 499 | 56767.54 | 8.7 | M/M | 9 | |
| *Glyma08g11270.1* | XP_006585139.1 | 0 | 1149 | 382 | 44155.06 | 9.65 | S/C | 5 | M |
| *Glyma08g17840.1* | XP_006598215.1 | 1 | 1017 | 338 | 38913.2 | 9.07 | C/C | 6 | |
| *Glyma08g37480.1* | XP_003531975.1 | 0 | 1203 | 400 | 46005.55 | 9.9 | M/M | 6 | M |
| *Glyma08g41780.1* | XP_003530716.2 | 0 | 1137 | 378 | 43139.99 | 10.06 | M/M | 5 | M |
| *Glyma08g41790.1* | XP_003530716.2 | 0 | 1140 | 379 | 43613.63 | 9.93 | M/M | 4 | M |
| *Glyma08g41850.1* | ACU19460.1 | 1 | 1074 | 357 | 40040.05 | 9.81 | _/_ | 6 | M |
| *Glyma08g41870.1* | XP_003532083.1 | 0 | 1212 | 403 | 45669.02 | 9.53 | M/M | 4 | M |
| *Glyma08g41880.1* | XP_003530719.1 | 0 | 1200 | 399 | 45272.77 | 9.69 | M/C | 4 | M |
| *Glyma09g05130.2* | XP_006586932.1 | 0 | 1188 | 395 | 45189.34 | 9.42 | M/M | 6 | M |
| ***Glyma09g05211.1*** | - | 0 | 489 | 162 | 18325.94 | 9.57 | _/_ | 2 | M |
| *Glyma09g11740.1* | XP_003533849.2 | 0 | 969 | 322 | 35922.01 | 8.93 | C/C | 6 | |

| | | | | | | | | | |
|---|---|---|---|---|---|---|---|---|---|
| *Glyma09g30200.2* | - | 4 | 1386 | 461 | 52385.02 | 9.03 | _/_ | 7 | |
| *Glyma09g37940.2* | - | 0 | 333 | 110 | 13420.88 | 9.71 | S/_ | 2 | |
| *Glyma10g06160.2* | ACU19409.1 | 2 | 1239 | 412 | 47178.22 | 9.21 | _/_ | 8 | |
| *Glyma10g36001.1* | - | 1 | 423 | 140 | 15991.62 | 9.36 | C/_ | 1 | |
| *Glyma11g12520.2* | - | 6 | 1539 | 512 | 59239.57 | 8.96 | _/_ | 6 | |
| *Glyma12g04720.1* | XP_006591910.1 | 5 | 1875 | 624 | 72061.2 | 9.2 | C/_ | 8 | |
| *Glyma13g20470.2* | XP_003542522.1 | 0 | 1008 | 335 | 38165.88 | 9.37 | _/_ | 8 | |
| *Glyma13g28790.1* | - | 1 | 948 | 315 | 36669.8 | 10.06 | M/M | 5 | M |
| *Glyma14g01940.1* | XP_006595700.1 | 4 | 1488 | 495 | 57734.25 | 8.87 | C/C | 7 | |
| *Glyma14g05540.1* | XP_006595840.1 | 0 | 1695 | 564 | 64859.57 | 9.01 | C/M | 5 | |
| *Glyma15g00290.1* | XP_003546903.1 | 4 | 1779 | 592 | 67858.52 | 9.36 | _/_ | 10 | |
| *Glyma15g10270.1* | XP_006597531.1 | 0 | 1146 | 381 | 44138.62 | 9.66 | M/M | 4 | M |
| ***Glyma15g16400.2*** | XP_006597741.1 | 0 | 1188 | 395 | 45137.86 | 9.82 | M/M | 5 | M |
| *Glyma15g16410.3* | XP_006597742.1 | 0 | 888 | 295 | 33293.35 | 9.2 | M/M | 3 | M |
| *Glyma15g16420.2* | - | 0 | 1170 | 389 | 44901.71 | 9.81 | M/M | 6 | M |
| *Glyma15g16430.2* | XP_003547396.1 | 1 | 1011 | 336 | 38574.23 | 10.04 | M/M | 4 | M |
| ***Glyma15g16530.1*** | - | 0 | 462 | 153 | 17432.66 | 9.2 | _/_ | 1 | M |
| *Glyma15g23480.1* | XP_003546577.1 | 0 | 909 | 302 | 33971.88 | 9.15 | C/C | 5 | |
| *Glyma15g41300.1* | XP_006598215.1 | 1 | 1023 | 340 | 39173.57 | 9.26 | C/C | 6 | |
| *Glyma16g09990.1* | XP_003547773.1 | 0 | 1119 | 372 | 42402.73 | 9.18 | S/_ | 6 | M |
| ***Glyma18g12810.2*** | XP_006602249.1 | 0 | 1182 | 393 | 44934.08 | 9.83 | M/M | 5 | M |
| *Glyma18g13720.1* | XP_003553150.2 | 0 | 1209 | 402 | 45866.34 | 9.69 | M/C | 5 | M |
| *Glyma18g13740.1* | XP_003551935.1 | 0 | 1206 | 401 | 45770.32 | 9.62 | M/C | 4 | M |
| *Glyma18g13750.1* | XP_006602266.1 | 0 | 1215 | 404 | 45567.37 | 9.8 | M/M | 5 | M |
| ***Glyma18g13780.2*** | - | 1 | 1158 | 385 | 43890.18 | 9.28 | M/M | 4 | M |
| *Glyma18g13790.2* | XP_003553152.1 | 0 | 1215 | 404 | 45862.46 | 9.64 | M/C | 5 | M |
| *Glyma18g13800.1* | XP_003551936.1 | 0 | 1209 | 402 | 45787.34 | 9.84 | M/M | 5 | M |
| *Glyma18g48450.1* | XP_006602863.1 | 1 | 813 | 270 | 32024.18 | 9.64 | C/C | 4 | |
| *Glyma19g22410.1* | XP_006604090.1 | 3 | 1422 | 473 | 53936.73 | 9.22 | M/M | 8 | |
| *Glyma19g32820.2* | XP_006604383.1 | 0 | 1731 | 576 | 65824.79 | 9.35 | M/M | 6 | |

| | | | | | | | | | |
|---|---|---|---|---|---|---|---|---|---|
| *Glyma20g31580.1* | XP_003556199.1 | 2 | 708 | 235 | 26407.4 | 8.29 | C/_ | 1 | |
| ***Medicago truncatula* (23)** | | | | | | | | | |
| *AC233682_35.1* | AES84904.1 | 1 | 987 | 328 | 38022.19 | 9.51 | C/C | 6 | |
| *Medtr2g019810.1* | AES64213.1 | 0 | 1101 | 366 | 42147.07 | 9.48 | M/M | 4 | M |
| ***Medtr2g019840.1*** | AES64216.1 | 0 | 1158 | 385 | 44803.67 | 9.75 | M/M | 4 | M |
| *Medtr2g034600.1* | AES65036.1 | 4 | 1704 | 567 | 63817.17 | 9.21 | _/_ | 8 | |
| *Medtr2g049780.1* | AES65844.1 | 0 | 942 | 313 | 35374.45 | 8.96 | C/C | 6 | |
| *Medtr2g060620.1* | AES66019.1 | 1 | 585 | 194 | 22492.82 | 6.17 | S/_ | 1 | |
| ***Medtr2g076320.1*** | AES66608.1 | 0 | 1272 | 423 | 48478.46 | 9.78 | C/_ | 4 | M |
| *Medtr3g085240.1* | AES72024.1 | 3 | 768 | 255 | 29289.01 | 9.71 | C/_ | 1 | M |
| *Medtr3g092710.1* | AES72631.1 | 1 | 1521 | 506 | 57893.45 | 9.29 | C/C | 7 | |
| *Medtr4g049400.1* | AES88145.1 | 0 | 702 | 233 | 26886.86 | 9.99 | S/M | 1 | M |
| *Medtr4g070060.1* | AES89105.1 | 5 | 1854 | 617 | 71478.59 | 9.38 | _/_ | 8 | |
| *Medtr4g119550.1* | AES91847.1 | 0 | 1179 | 392 | 44154.04 | 9.83 | C/M | 5 | M |
| *Medtr4g119570.1* | AES91849.1 | 2 | 921 | 306 | 34678.75 | 9.35 | S/ER | 4 | M |
| ***Medtr4g119580.1*** | AES91850.1 | 0 | 1134 | 377 | 42817.01 | 9.59 | M/M | 6 | M |
| *Medtr5g041630.1* | AES96801.1 | 1 | 927 | 308 | 35369.6 | 9.19 | _/_ | 6 | |
| *Medtr5g068860.1* | AES98461.1 | 0 | 1593 | 530 | 59944.28 | 5.09 | _/_ | 9 | |
| *Medtr5g084810.1* | AES99796.1 | 0 | 1779 | 592 | 68350.25 | 8.59 | C/C | 5 | |
| *Medtr5g094610.1* | AET00669.1 | 4 | 1587 | 528 | 61033.86 | 7.12 | C/C | 8 | |
| *Medtr7g080620.1* | AES80446.1 | 4 | 2214 | 737 | 85261.56 | 8.31 | M/M | 5 | |
| *Medtr7g081270.1* | ABN05921.1 | 0 | 855 | 284 | 33301.44 | 9.64 | M/_ | 5 | |
| *Medtr7g093000.1* | AES81460.1 | 0 | 1716 | 571 | 65078.95 | 9.06 | M/M | 5 | |
| *Medtr8g012210.1* | AET01407.1 | 0 | 1113 | 370 | 42988.12 | 9.45 | M/M | 6 | M |
| *Medtr8g105560.1* | AET05439.1 | 3 | 1683 | 560 | 64730.9 | 8.23 | _/_ | 8 | |
| ***Populus trichocarpa* (52)** | | | | | | | | | |
| *Potri.001G029400.1* | - | 0 | 1176 | 391 | 44429.55 | 9.87 | M/ER | 7 | M |
| *Potri.001G030300.1* | XP_002297734.1 | 0 | 1167 | 388 | 43791.63 | 9.83 | M/M | 7 | M |

| | | | | | | | | | |
|---|---|---|---|---|---|---|---|---|---|
| *Potri.001G034500.1* | XP_002297718.1 | 0 | 1116 | 371 | 42680.45 | 9.87 | M/ER | 6 | M |
| *Potri.001G034600.1* | XP_002297715.1 | 0 | 1161 | 386 | 44212.88 | 9.82 | C/M | 8 | M |
| *Potri.001G034700.1* | XP_002297714.1 | 0 | 1137 | 378 | 43391.15 | 9.76 | C/M | 8 | M |
| *Potri.001G034900.1* | XP_002297713.1 | 0 | 1116 | 371 | 42365.8 | 9.91 | C/M | 7 | M |
| *Potri.001G035000.1* | ABK95347.1 | 0 | 1161 | 386 | 43768.38 | 9.87 | C/M | 7 | M |
| ***Potri.001G035200.1*** | XP_002297707 | 0 | 1098 | 365 | 41755.19 | 9.39 | C/C | 6 | M |
| ***Potri.001G035300.1*** | | 0 | 1290 | 429 | 49425.87 | 9.07 | C/M | 6 | M |
| *Potri.001G035600.1* | XP_002297704 | 1 | 810 | 269 | 31034.46 | 9.75 | S/_ | 6 | M |
| ***Potri.001G361800.1*** | XP_002298854.2 | 5 | 1503 | 500 | 57295.75 | 9.6 | C/C | 7 | |
| ***Potri.003G189300.1*** | | 0 | 1185 | 394 | 44307.13 | 9.66 | C/C | 8 | M |
| ***Potri.003G190300.1*** | XP_002304820.1 | 0 | 1149 | 382 | 43606.04 | 9.72 | C/C | 7 | M |
| *Potri.003G190400.1* | - | 0 | 1131 | 376 | 43128.6 | 9.87 | C/_ | 7 | M |
| *Potri.003G190700.1* | XP_002304819.1 | 0 | 1101 | 366 | 41845.4 | 9.92 | M/M | 7 | M |
| *Potri.004G012400.1* | XP_002305563 | 0 | 1176 | 391 | 44661.97 | 9.29 | M/M | 7 | M |
| *Potri.004G012900.1* | XP_002304896.2 | 0 | 1185 | 394 | 45034.67 | 9.51 | S/M | 8 | M |
| *Potri.004G013000.1* | - | 0 | 1176 | 391 | 44631.1 | 9.69 | M/M | 7 | M |
| *Potri.004G013100.1* | - | 0 | 1176 | 391 | 44902.62 | 9.68 | M/M | 7 | M |
| *Potri.004G150600.1* | XP_002305414.1 | 0 | 846 | 281 | 32756.14 | 9.72 | C/C | 5 | |
| *Potri.004G209400.1* | XP_002328286.1 | 0 | 1002 | 333 | 37973.84 | 9.59 | _/_ | 7 | |
| *Potri.004G222000.1* | XP_002328768.1 | 0 | 1095 | 364 | 41861 | 9.7 | M/M | 6 | M |
| *Potri.006G205000.1* | XP_002308443.2 | 0 | 1056 | 351 | 40266.51 | 9.23 | C/_ | 6 | |
| *Potri.007G001800.1* | XP_002310354.2 | 6 | 1755 | 584 | 66912.27 | 8.95 | C/_ | 10 | |
| *Potri.007G069100.1* | XP_002310671.2 | 0 | 909 | 302 | 34745.19 | 9.85 | M/M | 4 | M |
| *Potri.008G216200.1* | XP_002330358.1 | 0 | 1104 | 367 | 42157.26 | 9.1 | C/_ | 6 | M |
| *Potri.009G116200.1* | XP_002313046.2 | 5 | 2022 | 673 | 78345.42 | 8.66 | C/_ | 8 | |
| ***Potri.009G170300.1*** | XP_002312791.1 | 1 | 1005 | 334 | 38068.22 | 9.78 | M/_ | 7 | |
| *Potri.010G022700.1* | - | 0 | 1134 | 377 | 43209.32 | 9.78 | C/_ | 5 | M |
| *Potri.010G167400.1* | XP_002315044.1 | 0 | 882 | 293 | 33652.76 | 9.73 | C/_ | 5 | |
| *Potri.011G005100.1* | XP_002317119.1 | 0 | 1188 | 395 | 45122.77 | 9.65 | M/M | 6 | M |
| *Potri.011G081400.1* | XP_002317291.2 | 0 | 984 | 327 | 36808.19 | 9.32 | M/ER | 6 | |

| | | | | | | | | | |
|---|---|---|---|---|---|---|---|---|---|
| ***Potri.011G085600.1*** | XP_002331073.1 | 2 | 474 | 157 | 18465.66 | 9.43 | S/ER | 3 | |
| *Potri.012G046700.1* | XP_002317871.1 | 0 | 1167 | 388 | 44672.14 | 9.37 | M/_ | 6 | M |
| *Potri.012G046800.1* | - | 0 | 1167 | 388 | 44802.48 | 9.43 | M/M | 6 | M |
| *Potri.012G118200.1* | XP_002332130.1 | 0 | 1779 | 592 | 67228.19 | 9.11 | C/M | 5 | |
| *Potri.012G118400.1* | - | 0 | 693 | 230 | 25865.05 | 8.37 | _/_ | 2 | |
| *Potri.013G113500.1* | XP_002327239.1 | 1 | 648 | 215 | 24041.56 | 8.49 | C/M | 1 | |
| *Potri.013G116700.1* | XP_002327325.1 | 0 | 1545 | 514 | 58710.99 | 8.18 | M/M | 10 | |
| *Potri.014G044900.1* | XP_002326874.1 | 1 | 1728 | 575 | 66193.71 | 8.06 | C/M | 5 | |
| *Potri.014G133200.1* | XP_002321027.1 | 0 | 1203 | 400 | 46151.95 | 9.19 | C/_ | 6 | M |
| *Potri.014G137400.1* | XP_002321054.1 | 0 | 1566 | 521 | 59079.66 | 6.37 | _/_ | 10 | |
| *Potri.015G005200.1* | XP_002321365.1 | 1 | 1554 | 517 | 57654.53 | 8.9 | C/C | 7 | |
| *Potri.015G038400.1* | XP_002322107.1 | 0 | 1164 | 387 | 44675.25 | 9.39 | M/M | 6 | M |
| *Potri.015G038500.1* | XP_002322106.1 | 0 | 1167 | 388 | 44676.56 | 9.57 | M/M | 6 | M |
| *Potri.016G048200.1* | XP_002322675.1 | 2 | 1887 | 628 | 72484.17 | 8.98 | M/M | 6 | |
| *Potri.016G048300.1* | XP_002322676.1 | 0 | 1704 | 567 | 65553.33 | 8.16 | S/M | 4 | |
| *Potri.016G048400.1* | XP_002322677.1 | 0 | 1704 | 567 | 65461.45 | 8.82 | C/M | 3 | |
| *Potri.016G072200.1* | XP_002322796.1 | 0 | 1065 | 354 | 40590.95 | 8.87 | _/_ | 6 | |
| *Potri.017G067600.1* | - | 3 | 1509 | 502 | 56644.7 | 9.3 | C/_ | 7 | |
| *Potri.018G051600.1* | XP_002324772.2 | 1 | 1314 | 437 | 50484.45 | 9.45 | M/M | 8 | |
| *Potri.T104200.1* | - | 0 | 1176 | 391 | 44661.97 | 9.29 | M/M | 7 | M |

[a] Gene models annotated used here were from Phytozome v9.0 (http://www.phytozome.net/) except that for *Picea abies* (Norway spruce) which genome annotation data was from ConGenIE.(http://congenie. org/) (Nystedt et al. 2013) and *Amborella trichopoda* which was from Amborella Genome Database (http:// amborella.org/) (Chamala et al. 2013). The *mTERF* gene models have been renamed for *Amborella trichopoda* (with “ATr” instead of “evm_27.model.AmTr_ v1.0”), *Arabidopsis lyrata* (with “Al_” as a prefix), *Carica papaya* (with “Cp_” instead of “evm.model.”) and *Selaginella moellendorffii* (with “Sm” as a prefix). The *mTERF* genes in bold were corrected based on mRNA sequences in NCBI or their homologs in relatives species, and a lower letter following the *mTERF* genes indicated there were more than one gene model re-annotated.

[b] Genebank accessions entries of plant mTERF proteins were determined by BLAST tools (http://blast.ncbi.nlm.nih.gov/Blast.cgi).

[c] Only introns dispersed in DNA coding sequence were considered here.

[d] Length in amino acids (aa), molecular weight (MW) and isoelectric point (pI) of deduced proteins were calculated by Expasy online tools (http://expasy.org/tools/).

[e] Subcellular location of mTERF proteins were predicted *in silico*, by using Predotar (https://urgi.versailles.inra.fr/predotar/predotar.html) (Small et al. 2004) and (/) TargetP (http://www.cbs.dtu.dk/services/TargetP/) (Emanuelsson et al. 2007) for higher plants; Predalgo (https://giavap-genomes.ibpc.fr/cgi-bin/predalgodb.perl?page=main) (Tardif et al. 2012) for algae (*Chlamydomonas reinhardtii* and *Volvox carteri*) (Wobbe and Nixon 2013), and (/) HECTAR (http://www.sb-roscoff.fr/hectar/) (Gschloessl et al. 2008) for *V.carteri*. C, chloroplast or plastid; M, mitochondria; ER or S, Secretory; _, none or other place.

[f] The mTERF motif model used here was from SMART database (Letunic et al. 2012) and about 32 aa in length (Roberti et al. 2006a; Roberti et al. 2006b).

[g] Higher plant mTERF proteins were divided into two main subfamilies on the basis of their conserved evolutional relationships in phylogenetic tree and organelle targeting. M, M-class *mTERF* gene.

[h] Notes for plant *mTERF* genes. *C. reinhardtii mTERF* genes were identified as *MOCs* (Wobbe and Nixon 2013); Gene models of *P. abies mTERF* genes were of high quality (Nystedt et al. 2013); The names for maize *mTERF* genes were given previously (Zhao et al. 2014).

**Supplementary Table S2. The paralogous gene sets for mining positive selection on plant M-class *mTERF* genes.**

| Gene sets | Name | Gene model | Positively Selected Sites[b] |
|---|---|---|---|
| ***Oryze sativa* ssp. japonica (332 aa)[a]** | | | |
| Osa_1 | Osa_001 | LOC_Os02g51460.1 | **103**, 125, **135**, **167**, 179, **180**, 182, 194, |
| | Osa_002 | LOC_Os11g09990.1 | **195**, **201**, 205, **234**, 236, **238**, **241**, 245, |
| | Osa_003 | LOC_Os11g10000.1 | **267**, **270**, **291**, 304, **305**, **307**, 308, 311, |
| | Osa_004 | LOC_Os11g10040.1 | **312**, 315, 323 |
| | Osa_005 | LOC_Os11g14130.1 | |
| Osa_2 | Osa_006 | LOC_Os02g51450.2 | 16, **20**, 69, **98**, 100, 122, 129, **131**, |
| | Osa_007 | LOC_Os06g12040.1 | **132**, **155**, **157**, **161**, **162**, **163**, 175, |
| | Osa_008 | LOC_Os06g12050.1 | 178, **190**, 201, 213, 215, 216, 230, |
| | Osa_009 | LOC_Os06g12060.1a | **236**, **238**, 241, **242**, **250**, **262**, **267**, |
| | Osa_010 | LOC_Os06g12060.1b | **273**, **292**, **298**, **304**, 315, **319** |
| | Osa_011 | LOC_Os06g12070.1 | |
| | Osa_012 | LOC_Os06g12080.1 | |
| | Osa_013 | LOC_Os06g12100.1 | |
| | Osa_014 | LOC_Os06g12110.1 | |
| Osa_3 | Osa_015 | LOC_Os05g33440.1 | 21, 26, 58, 61, 69, 85, 88, 132, 157, |
| | Osa_016 | LOC_Os05g33460.2 | 190, 193, 194, 213, 238, **239**, 251, |
| | Osa_017 | LOC_Os05g34160.1 | **252**, **304**, 305, 310, 314 |
| | Osa_018 | LOC_Os07g24090.1 | |
| ***Populus trichocarpa* (339 aa)** | | | |
| Ptr_1 | Ptr001 | Potri.001G035300.1 | **48**, 59, **61**, **65**, **69**, **97**, **100**, 102, |
| | Ptr002 | Potri.001G029400.1 | **106**, **133**, 138, **139**, **142**, 165, **166**, |
| | Ptr003 | Potri.001G030300.1 | **167**, 170, **173**, 202, **206**, **231**, 236, |
| | Ptr004 | Potri.001G034500.1 | **238**, **241**, **242**, **249**, 250, **270**, **273**, |
| | Ptr005 | Potri.001G034600.1 | 274, 298, **300**, **303**, **304**, **319**, **333**, |
| | Ptr006 | Potri.001G034700.1 | **336**, 338 |
| | Ptr007 | Potri.001G034900.1 | |
| | Ptr008 | Potri.001G035000.1 | |
| | Ptr009 | Potri.001G035200.1 | |
| | Ptr010 | Potri.001G035600.1 | |
| | Ptr011 | Potri.003G189300.1 | |
| | Ptr012 | Potri.003G190300.1 | |
| | Ptr013 | Potri.003G190400.1 | |
| | Ptr014 | Potri.003G190700.1 | |
| | Ptr015 | Potri.004G222000.1 | |
| Ptr_2 | Ptr016 | Potri.012G046700.1 | 94, 95, 100, **106**, 139, 336 |
| | Ptr017 | Potri.012G046800.1 | |

| | | | |
|---|---|---|---|
| | Ptr018 | Potri.015G038400.1 | |
| | Ptr019 | Potri.015G038500.1 | |
| Ptr_3 | Ptr020 | Potri.004G012400.1 | **61**, **101**, 106, 123, **131**, 136, **166**, |
| | Ptr021 | Potri.004G012900.1 | **167**, **169**, **183**, **199**, **202**, 205, 238, |
| | Ptr022 | Potri.004G013000.1 | 241, 242, 272, 320, **333** |
| | Ptr023 | Potri.004G013100.1 | |
| | Ptr024 | Potri.008G216200.1 | |
| | Ptr025 | Potri.010G022700.1 | |
| | Ptr026 | Potri.011G005100.1 | |
| | Ptr027 | Potri.T104200.1 | |
| ***Arabidopsis thaliana* (414 aa)** | | | |
| Ath | | AT1G56380.1 | **13**, **76**, 120, 172, **177**, **178**, 202, |
| | | AT1G61960.1 | 208, **211**, **215**, **236**, 251, 269, 274, |
| | | AT1G61970.1 | **280**, **282**, **285**, **286**, **292**, **309**, 320, |
| | | AT1G61980.1 | **349**, 381, **387**, **392**, **399** |
| | | AT1G61990.1 | |
| | | AT1G62010.1 | |
| | | AT1G62085.1 | |
| | | AT1G62110.1 | |
| | | AT1G62120.1 | |
| | | AT1G62150.1 | |
| | | AT1G62490.1 | |
| | | AT3G46950.1 | |
| | | AT5G23930.1 | |
| ***Arabidopsis lyrata* (446 aa)** | | | |
| Aly | | Al_315177a | 6, 11, 12, **13**, 18, 25, 27, **32**, 44, **48**, |
| | | Al_315177b | 52, **55**, **70**, 81, 84, **86**, **107**, **111**, |
| | | Al_315190 | **114**, **118**, 147, **157**, **162**, **169**, 172, |
| | | Al_338172 | **179**, **188**, **213**, 218, **219**, **220**, 227, |
| | | Al_475136 | **245**, **249**, **252**, **255**, **256**, 268, **275**, |
| | | Al_475137 | **277**, 288, 291, 292, **310**, **316**, 317, |
| | | Al_485072 | **321**, **323**, **325**, **326**, **327**, **333**, 341, |
| | | Al_489254 | **350**, **358**, **361**, **362**, 385, **388**, **390**, |
| | | Al_893233 | **396**, 422, **423**, **424**, **428**, 431, **432**, |
| | | Al_893270 | **433**, **437**, **440**, **446**, **449** |
| | | Al_893275 | |
| | | Al_893280 | |
| | | Al_893283 | |
| | | Al_893289 | |
| | | Al_893298a | |

| |
| --- |
| Al_893298b |
| Al_893300 |
| Al_894067 |
| Al_907730 |

[a] The length of multiple protein sequence alignments of *mTERF* gene sets is shown in parentheses.

[b] Positively selected sites under the M8 model are identified using BEB method. The sites with the posterior probability (P) of > 99% are marked in bold.
