## Supplementary Figures for "Expansion and Adaptive Evolution of the *mTERF* Gene Family in Plants"

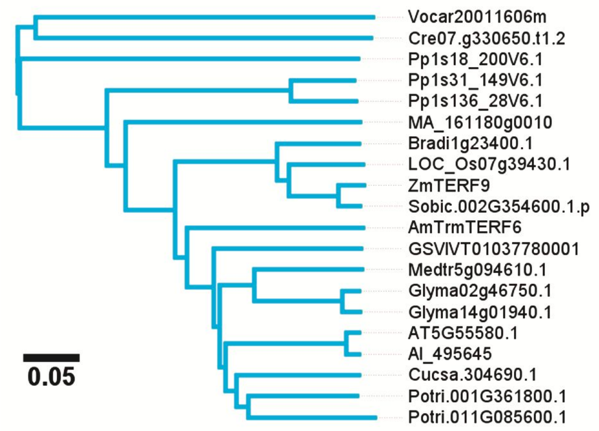


**Supplementary Figure S1. The conserved clade IV of fig. 1.**

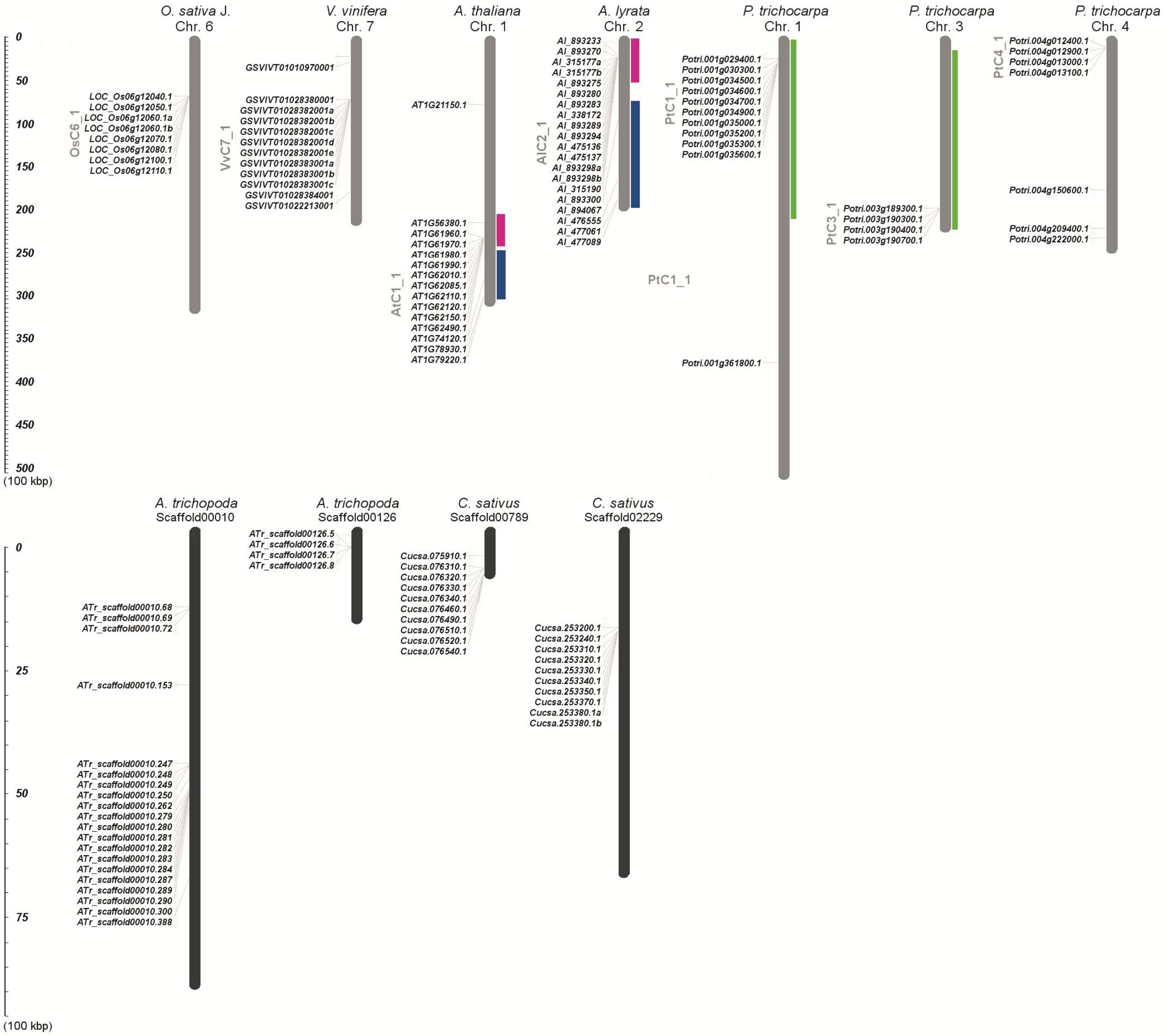


**Supplementary Figure S2. The chromosomal location of *mTERF* genes involving tandem duplication event.** The location of *mTERF* genes on respective chromosomes was obtained from Phytozome database (https://www.phytozome.net). The blocks in color denote the homologous regions.

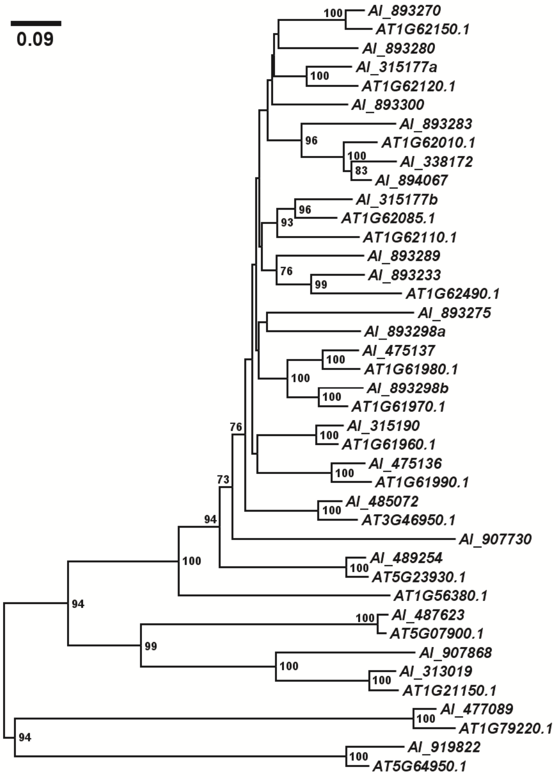


**Supplementary Figure S3. The ML tree of M-class *mTERF* genes of *Arabidopsis thaliana* and *Arabidopsis lyrata*.** Multiple sequence alignment of *A. thaliana* and *A. lyrata* M-class mTERF proteins was performed using MUSCLE v3.8.31 (Edgar 2004) and then was used to built the ML tree with MEGA 6.0 as described in fig. 7. The bootstrap values more than 70 percent were shown.

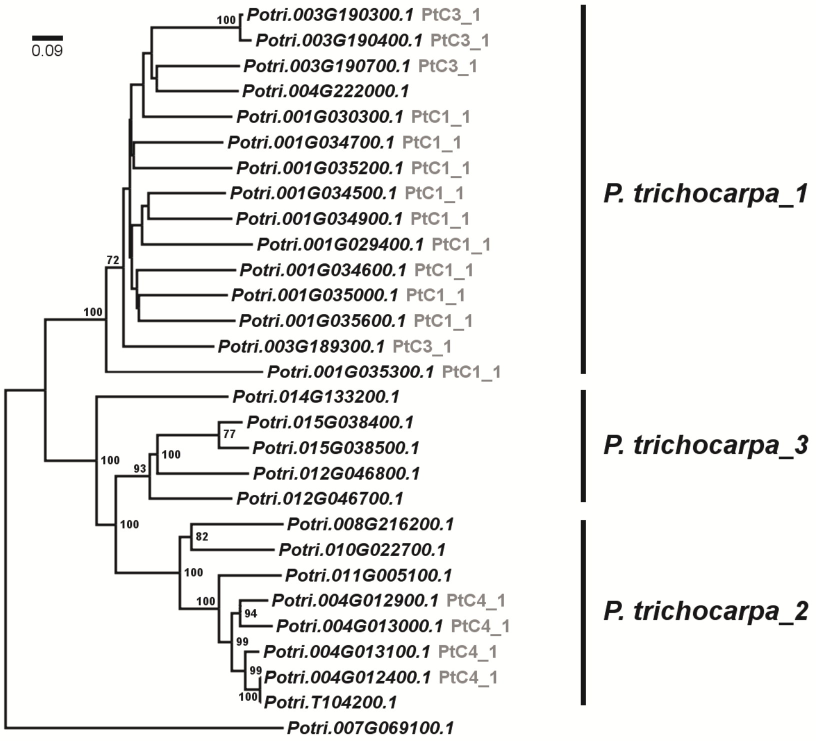


**Supplementary Figure S4**. The ML tree of *Populus trichocarpa* M-class *mTERF* genes. Multiple sequence alignment of *P. trichocarpa* M-class mTERF proteins was conducted by MUSCLE v3.8.31 (Edgar 2004).The ML tree was constructed using MEGA 6.0 as described in fig. 7. Only the bootstrap values more than 70 percent were shown.

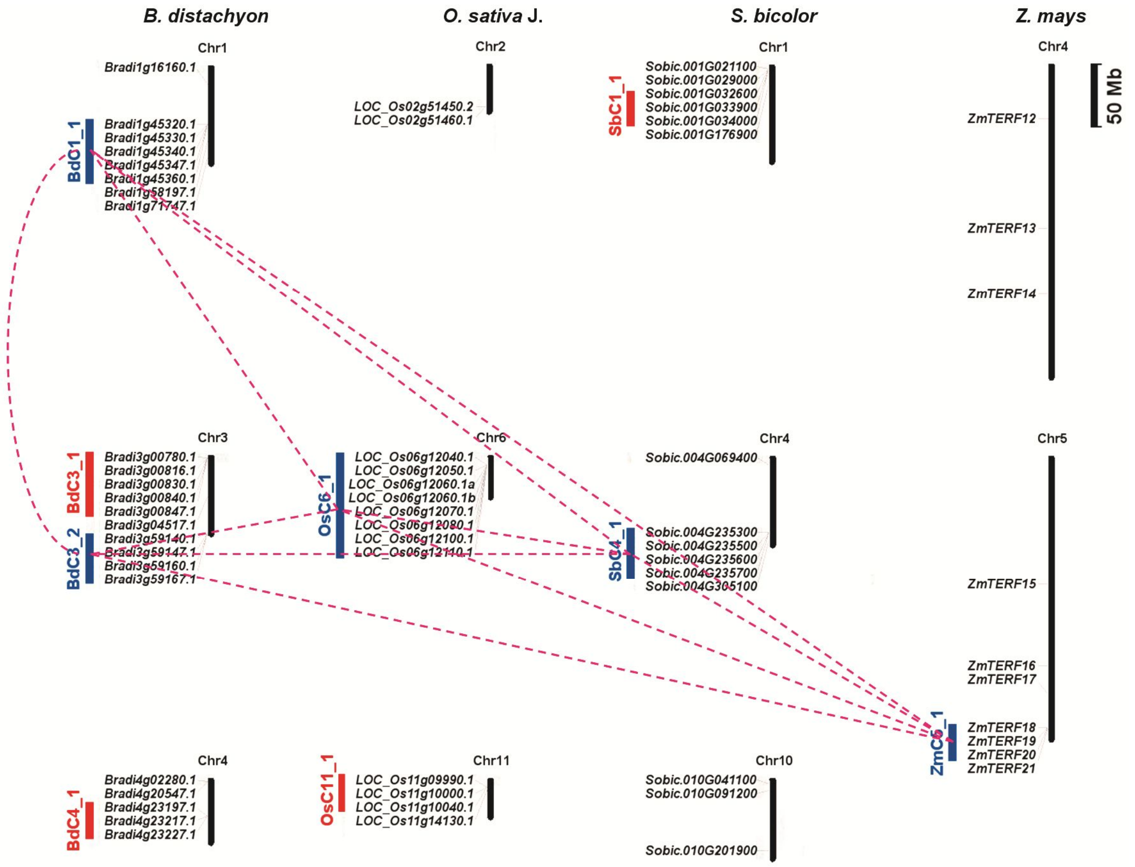


**Supplementary Figure S5. Distribution of monocot M-class *mTERF* gene clusters on chromosomes.** The *mTERF* genes marked by blue bars are homologs and located in the syntenic regions of different species. There was no *mTERF* gene homologous to the *mTERF* gene clusters marked by red bars found in their syntenic regions. The syntenic regions were predicted using SyMAP v4.3 Synteny (Soderlund et al. 2011) (http://symapdb.org/projects/poaceae/).

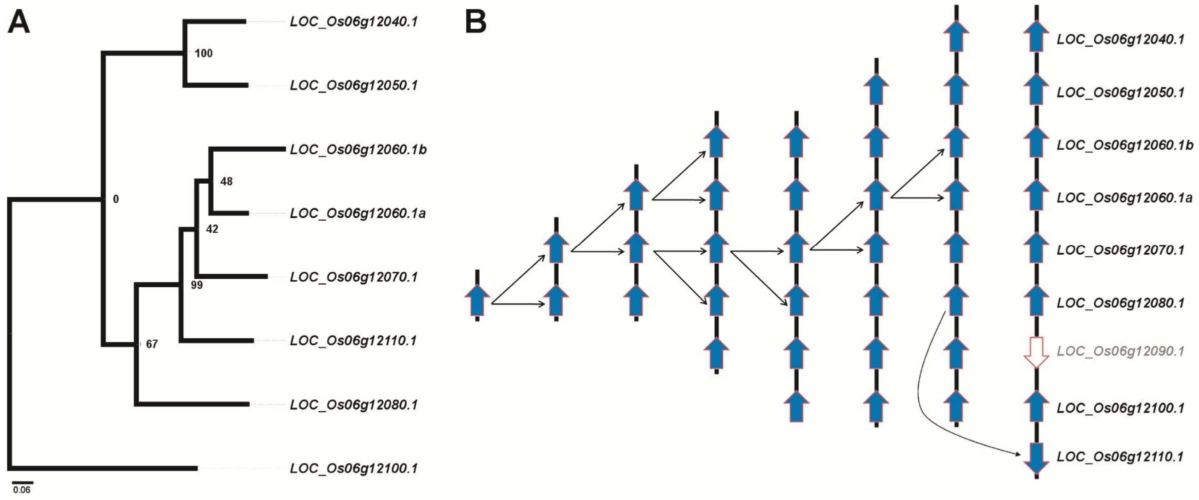


**Supplementary Figure S6. Hypothetical origins of OsC6_1 *mTERF* genes by tandem duplication.** (*A*) The ML tree of OsC6_1 *mTERF* genes was built using MEGA 6.0 (Tamura et al. 2013) with their protein sequence alignment as described in fig. 7. (*B*) The hypothetical origins of OsC6_1 *mTERF* genes by tandem duplication were analyzed based on the left ML tree and their locations on chromosome 6 of rice.

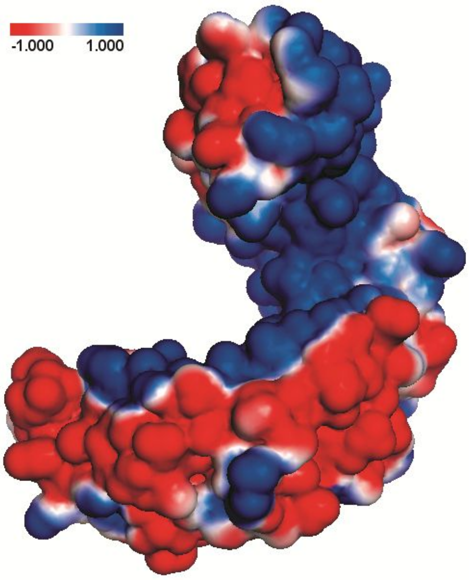


**Supplementary Figure S7. Electrostatic analysis of M-class mTERF protein.**

The electrostate of LOC_Os06g12100 protein was calculated using PDB2PQR (Baker et al. 2001) and APBS (Dolinsky et al. 2007) and was displayed with PyMOL v1.6.0.0 (The PyMOL Molecular Graphics System, Version 1.6.0.0 Schrödinger, LLC.).

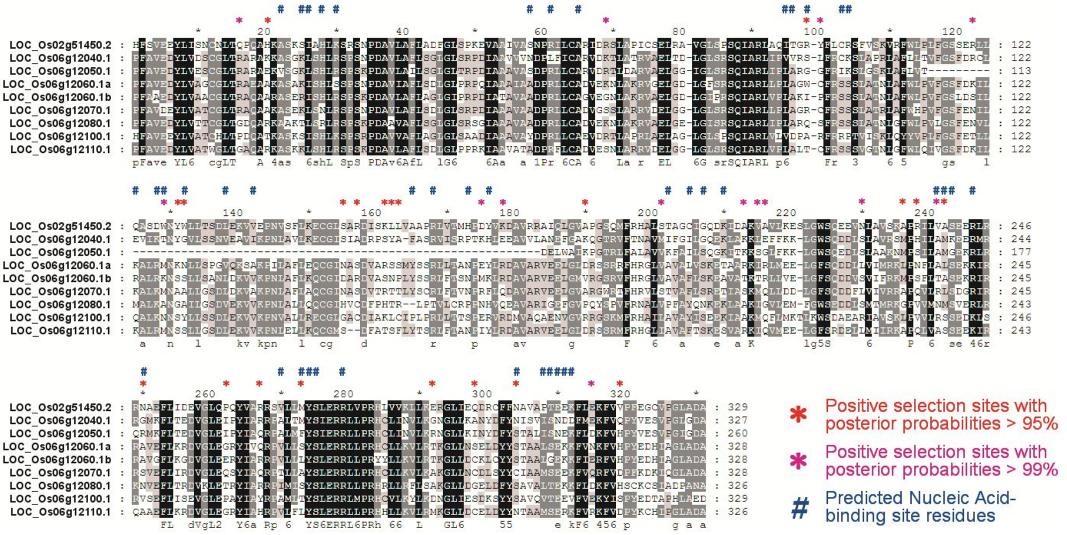


**Supplementary Figure S8. The positively selected sites and putative nucleic acid-binding sites in OsC6_1 mTERF proteins.** The positively selected sites of OsC6_1 M-class mTERF proteins were identified using PAML v4.7 (Yang 2007) and the putative nucleic acid-binding sites were predicted by I-TASSER (Zhang 2008). The alignment of OsC6_1 mTERF proteins was displayed by GeneDoc (http://www.nrbsc.org/gfx/genedoc/ebinet.htm). The symbols, * and # above the alignment, are used to denote the positively selected sites and nucleic acid-binding sites, respectively.
