## Supplementary Data for "Expansion and Adaptive Evolution of the *mTERF* Gene Family in Plants"

### Supplementary Data S1. The phylogenetic tree of plant M-class *mTERF* genes (in newick).

('AT1G74120.1':0.554651068029568,('Glyma04g32192.1':0.8158716738002092,(('Bradi5g23380.1':0.32463763531145967,'LOC_Os04g54510.1':0.12471560
524005403):0.8045168730329297,((((((Sm444400:1.1376039500707213,MA_10436083g0040:1.154998295577526):0.31743786388825446,((MA_209070g0
010:0.1884193321202953,MA_21437g0010:0.3069823783851236):0.10048183024754942,(MA_124772g0010:0.4034840306576703,(MA_34619g0010:0.10
912501704248144,MA_760114g0010:0.14724370584610916):0.10461155590891395):0.2272338461637227):0.6898288935002137):0.05818527918704303,
((MA_2589g0010:0.5295453789912195,((MA_280158g0010:0.2746490717740158,MA_45859g0010:0.2634982363800894):0.041200916775054684,(MA_
104598g0010:0.24663142853863654,(MA_590299g0010:0.3107765096294784,MA_139814g0010:0.6058424903096158):0.3113316626180256):0.0881197
090504446):0.25157229111111856):0.16553849645504606,((Al_477089:0.08382282137494261,'AT1G79220.1':0.11490315859040111):0.278519419346686
84,(AmTrmTERF14:0.49541746122347,(('Glyma16g09990.1':0.26746483409163224,'Medtr8g012210.1':0.2967740909553298):0.32387777004484336,(GS
VIVT01036787001:0.3142865354322393,('evm.model.supercontig_36.174':0.1936748969144501,'Potri.007G069100.1':0.196454630680011):0.05202666451
8668476):0.05114545234108186):0.0939688877193834):0.15703376490297594):0.6182204794562659):0.18985579136590566):0.04089260714256303,(M
A_118798g0010:0.6820394135466472,((MA_10426325g0010:0.2627768364670314,MA_86824g0010:0.36689185483746534):0.09966251417075365,((MA
_493420g0010:0.29005044107198563,(MA_10083329g0010:0.1393374550967235,MA_804585g0010:0.20292080153698044):0.239068688787966):0.0562
16264437009084,(MA_122549g0010:0.5318812560772183,(((MA_628537g0010:0.2720751609799517,(MA_10430524g0010:0.24604189723649048,MA_1
0430524g0020:0.06750646558495778):0.18019417963564696):0.030777740434517434,(MA_593288g0010:0.2973708873542269,(MA_10435408g0010:0.2
5197667157082165,(MA_10429074g0010:0.12140529612200181,MA_10427666g0020:0.057941162188865455):0.2114304398414991):0.055218275743389
7):0.040845314925899996):0.07509999245086058,((MA_161434g0010:0.15355230112417545,(MA_10429715g0010:0.19189349025414085,MA_205355g
0010:0.02412666089918411):0.17678300680429038):0.13771398262421622,((MA_92135g0010:0.11608240926595413,MA_33244g0010:0.2093054215287
4101):0.18432630963258415,(MA_178046g0010:0.25818087032838666,(MA_10041174g0010:0.1063697183602003,(MA_6307g0010:0.158216156291021
95,MA_14049g0010:0.07515721757766483):0.08370349662429642):0.01783953241032003):0.10095871881368991):0.03972234823069465):0.0517723446
47585504):0.06122743126019626):0.023247481213385005):0.056047575542296287):0.2090333460687876):0.3543310528013187):0.2223850195830929,((
(ZmTERF18:0.056314477379809116,'Sobic.004G235700.1.p':0.050991429424291704):0.738983679083427,(('Bradi3g59147.1':0.1238354258766685,'Bradi
3g59160.1':0.1128367912480471):0.2188059451468894,(ZmTERF11:0.1120890699055761,('Sobic.003G396100.1.p':0.05060123458388567,('Sobic.003G19
9700.1.p':0.15220761914863715,'Sobic.004G235500.1.p':0.08074873253809019):0.03817782890363416):0.04168487379519294):0.18384132973960202):0.
34402059659611406):0.07115354495546986,((('Sobic.003G389500.1.p':0.3660464259305331,(('LOC_Os02g51460.1':0.40356900877126606,'Sobic.004G23
5300.1.p':0.3205047789903716):0.05877446991275352,(ZmTERF21:0.11207154289089637,'Sobic.004G305100.1.p':0.0876041600013843):0.315332138075
43875):0.038746639155041146):0.20064312378998306,((Bradi1g45360:0.4983288997355025,('LOC_Os11g10040.1':0.3377795973804219,('LOC_Os11g14
130.1':0.4011891496897061,('LOC_Os11g09990.1':0.23993499022879008,'LOC_Os11g10000.1':0.3016817812005875):0.08989585061014735):0.05669446
088451133):0.10074832348857604):0.07041889745668624,('Sobic.006G183300.1.p':0.2785700879117116,(('Sobic.002G179800.1.p':0.26723204954454965,
'Sobic.010G041100.1.p':0.31056511278683224):0.13467444820494515,(ZmTERF5:0.3037332309885555,(('Sobic.001G029000.1.p':0.17087075637981325,'
Sobic.003G110500.1.p':0.1663322929261207):0.04907683698707129,(('Sobic.001G021100.1.p':0.3860579130720773,'Sobic.001G032600.1.p':0.1197756556
8572614):0.08139609018335787,('Sobic.001G033900.1.p':0.17243152796363212,'Sobic.001G034000.1.p':0.15750718020803586):0.06123143335702751):0.
02661970365248007):1.34750946315897E-6):0.057066094892164757):0.006229314560957365):0.30148485549787307):0.12928791176028967):0.0534926
1319731831,((('Bradi1g45347.1':0.3550883899942749,('LOC_Os06g12110.1':0.21679755505825465,('LOC_Os06g12060.1b':0.21177247652541464,('LOC_
Os06g12060.1a':0.15452502538519924,'LOC_Os06g12070.1':0.23606991109882644):0.03253230978031014):0.06373874550609686):0.0819247749572018
4):0.09300579864136886,((Bradi1g45340:0.34817749632498685,('Bradi1g45330.1':0.2708142616989549,'LOC_Os06g12080.1':0.3611207853616464):0.05
617815183758764):0.05855844315929498,(('LOC_Os06g12040.1':0.18482122384389468,'LOC_Os06g12050.1':0.2951467906061513):0.209913513894673
44,((ZmTERF3:0.12149186193287596,'Sobic.001G176900.1.p':0.09390410035916243):0.3894505628645737,('Sobic.004G069400.1.p':0.1469223684840947
6,(ZmTERF15:0.025972374148323145,ZmTERF16:0.004315223154467117):0.10034694190619656):0.3077320924837837):0.09466185844431388):0.0630
017544998358):0.06205494219208053):0.24392508140350613,(('Bradi3g59167.1':0.1175774938668821,('LOC_Os02g51450.1':0.15115883732915253,(Zm
TERF19:1.34750946315897E-6,'Sobic.004G235600.1.p':0.037024441577832926):0.05920787968972873):0.015263598007392668):0.3725168321165426,('B
radi3g00847.1':0.36278558070791317,(('LOC_Os06g12100.1':0.1988233678035673,('Bradi1g45320.1':0.182890760218677,(ZmTERF8:0.021794393745081
68,'Sobic.010G091200.1.p':0.02948995451420427):0.1250803176595301):0.007191044514536644):0.12745671058248406,(('Bradi3g00816.1':0.2725282096
4605223,('Bradi3g00830.1':0.22269619786110767,('Bradi3g04517.1':0.17538511796828485,('Bradi3g00780.1':0.14280797650785595,'Bradi4g02280.1':0.08
436917223648757):0.020887740646894644):0.0625402730793738):0.028573179547704268):0.043343970756626764,(('Bradi1g58197.1':0.26503809468141
565,'Bradi1g71747.1':0.22852215383783087):0.0408172524617261,(('Bradi4g20547.1':0.08618795830155346,'Bradi4g23227.1':0.06521907324876315):0.03

0654406221087865,('Bradi4g23197.1':0.07696499862404677,'Bradi4g23217.1':0.21296368490064033):0.08465626220235398):0.1725396584288144):0.008
209259784952202):0.07940103109956885):0.019771384888226218):0.20515556954307557):0.022342931517319888):0.0656833828736064):0.0340269610
5062295):0.4833581329303192):1.34750946315897E-6,((((Al_919822:0.08938807419395656,'AT5G64950.1':0.06577930946052006):0.472288988039938,('
Cucsa.201490.1':0.5067206163414223,('Glyma08g11270.1':0.4749390582495951,(GSVIVT01022213001:0.22950347440884722,'evm.model.supercontig_17
.174':0.36441719171210984):0.07904298323595789):0.0841331761660767):0.062461192796048295):0.308549838711425,(((AmTrmTERF24:0.0273167256
96277536,AmTrmTERF26:0.02360964675852251):0.4907857957329296,(GSVIVT01009012001:0.39637184355794,'evm.model.supercontig_36.118':0.390
24256677655267):0.31160458582514794):0.15860575440100175,((AmTrmTERF4:0.011849794903784539,AmTrmTERF5:0.039979497437371406):0.4504
664263727846,((AmTrmTERF30:0.21062468507846355,(AmTrmTERF17:0.05534359752277623,AmTrmTERF19:0.031568027816174536):0.14949263561
214884):0.30868794725659005,(((AmTrmTERF12:0.03595248745740284,(AmTrmTERF15:0.03329528286554931,(AmTrmTERF9:0.021438277921354793
,AmTrmTERF37:0.09049668945408691):1.34750946315897E-6):0.016171854494019627):0.4531491203205529,(AmTrmTERF22:0.030803037744279824,(
AmTrmTERF20:0.0443536480646102,(AmTrmTERF25:0.02113177108100509,(AmTrmTERF28:1.34750946315897E-6,AmTrmTERF29:1.3475094631589
7E-6):0.025989860145082557):0.013839564891982509):1.34750946315897E-6):0.28567995978231775):0.03926420219639864,((AmTrmTERF18:0.021421
76364536387,AmTrmTERF36:0.04491789653075402):0.2521459500712385,((AmTrmTERF31:0.07562043191065886,(AmTrmTERF43:0.07520672620806
595,(AmTrmTERF23:0.027926979490862588,AmTrmTERF49:0.03849543367382765):0.0171607366711703):0.00461592443950224):0.1031583414194527
5,(AmTrmTERF33:0.04276016669466406,(AmTrmTERF35:0.07606095151042865,(AmTrmTERF32:1.34750946315897E-6,AmTrmTERF48:0.0073212752
97775419):0.031083207790640447):1.34750946315897E-6):0.2173806816895473):0.15358800558426675):0.17809668192848574):0.024810967067215053
):0.03500544087237179):0.18435286966328296):0.035607857846112506):0.13058615244140648,((('Potri.001G035300.1':0.5514482931402752,(('Potri.003
G190300.1':0.004903201099786824,'Potri.003G190400.1':0.07847552555717201):0.34894161029343806,('Potri.001G034600.1':0.34937162986678155,(('Po
tri.001G029400.1':0.42888951929404706,(('Potri.001G034500.1':0.3024701901281304,'Potri.001G034900.1':0.28597310004155346):0.04852760430184593,
('Potri.003G190700.1':0.2968176692989514,'Potri.004G222000.1':0.30244524041107346):0.07624179924819757):1.34750946315897E-6):0.0664095149615
5129,('Potri.003G189300.1':0.35524900481678806,(('Potri.001G034700.1':0.2792391823104714,'Potri.001G035200.1':0.327788038593229):0.066780660118
5578,('Potri.001G030300.1':0.34153890097568956,('Potri.001G035000.1':0.2802079462540702,'Potri.001G035600.1':0.316959328916188):0.058855408940
538506):0.03352216549104063):0.015331179473503553):0.024655048509148188):0.014601913666994851):0.004592119292904091):0.0479319008557311
4):0.1476878516745148,((('Cucsa.165480.1':0.1391465943962998,'Cucsa.166200.1':0.13397714490564927):0.0656362149113675,(('Cucsa.076540.1':0.1611
8388804947098,'Cucsa.281080.1':0.13706020248034445):0.08865491910835026,('Cucsa.281090.1':0.1279260967869313,'Cucsa.281210.1':0.093471441185
15699):0.047652017030903736):0.022685283666499502):0.011092010161981098,('Cucsa.076490.1':0.1394045855245088,(('Cucsa.165090.1':0.1441938279
28454,('Cucsa.076510.1':0.1536986619581525,('Cucsa.076310.1':0.12999303663286066,('Cucsa.035460.1':0.062021009028140964,'Cucsa.076320.1':0.1196
583979295255):0.003893500661401723):0.03938704620574404):0.05417300377883055):0.004397895065964538,('Cucsa.076340.1':0.1495292805576236,((
'Cucsa.165080.1':0.12230943011920793,('Cucsa.075910.1':0.10606826854508308,'Cucsa.076460.1':0.12131296866881996):0.039960584758116764):0.0141
08162700552629,('Cucsa.076520.1':0.137039980062862,('Cucsa.076330.1':0.21899435478278598,('Cucsa.324410.1':0.21647706463396527,('Cucsa.347310.
1':0.007265218637231227,'Cucsa.371610.1':0.004293493820516312):0.09090607601496031):0.04120788441349355):0.02329673535087526):0.0377303112
7555503):0.01864991906690235):0.02161933426106325):0.019197378056071895):0.03720367943497667):0.37369336613797866):0.2879023822042064,((
((Bradi1g16160:0.21957382221886626,'LOC_Os07g24090.1':0.278479418674744):0.13501712177869132,(ZmTERF30:0.06928018709065302,'Sobic.007G
114700.1.p':0.06791836540614611):0.245428203191933):0.48296041433070147,(('LOC_Os05g34160.1':0.2854103507651636,('Bradi2g25372.1':0.2101542
7084313118,'Bradi2g25374.1':0.17709401664701716):0.09156289409413905):0.5055757734145805,('LOC_Os05g33440.1':0.5135488830462651,((ZmTER
F26:0.2334688388691157,'Sobic.010G201900.1.p':0.21574308053904162):0.14063184501509443,(('Bradi2g25837.1':0.27323519265485546,'LOC_Os05g33
460.1':0.26723082741000703):0.08403099652799173,(ZmTERF22:0.07686352331859031,'Sobic.009G133400.1.p':0.10504374081568836):0.235468148028
91933):0.029162344057673865):0.14973110756438804):0.2757248822569142):0.19148663087658815):0.18163148630835058,(('AT1G56380.1':0.76406366
55425973,(Al_907730:0.6620258658128013,((Al_489254:0.040529493003451204,'AT5G23930.1':0.03510457165593854):0.27928613972609156,((Al_4850
72:0.04963466011090688,'AT3G46950.1':0.07285907444026038):0.1936640566620233,((Al_894067:0.1763031530779293,(Al_893283:0.294284707227814
47,(Al_338172:0.24726823932440756,'AT1G62010.1':0.11831535322745662):1.34750946315897E-6):0.019524381402479807):0.1626371151464985,((Al_
475136:0.059738081984719256,'AT1G61990.1':0.0692474936759927):0.18668670347873212,(Al_893275:0.4143679175864,((Al_893298a:0.273430712490
8571,((Al_475137:0.05956683727434585,'AT1G61980.1':0.09967772910498608):0.0707979263845415,(Al_893298b:0.10534026099057456,'AT1G61970.1'
:0.10105293410472083):0.06679248567271716):0.07208217869807963):0.045677881631766115,((Al_893280:0.16677270998154542,(Al_315177a:0.09524
598587959865,'AT1G62120.1':0.10256930166327914):0.08170463397542206):0.04704355481405479,(((Al_315190:0.041958143148589526,'AT1G61960.1'
:0.042554086629124):0.1320827279164004,(Al_893289:0.16055961835565496,(Al_893233:0.11439498864168914,'AT1G62490.1':0.6974247085698404):0.
07491231799010828):0.037216356822708366):0.009033771075178363,(Al_893300:0.20680837693256587,((Al_893270:0.03199726145530442,'AT1G6215

0.1':0.05777364753231149):0.19799438783065462,('AT1G62110.1':0.1641996179958547,(Al_315177b:0.11078264586824332,'AT1G62085.1':0.108283708
875905):0.04111064907273525):0.016414729654095326):0.018587814411161297):0.007385719173691359):0.01862851995888152):0.0168349733335484):
0.00940450088337248):0.022858092897496227):0.02060794354037797):0.06480819212661794):0.03380865046111368):0.02237063860645645):0.232567
71864791):0.5322869501520222,((('Cucsa.088410.1':0.14389491803742968,'Cucsa.393940.1':0.1817626177453531):0.6038998372831407,('evm.model.supe
rcontig_245.9':0.9138216101459639,('evm.model.supercontig_13.147':0.48366522227793546,'evm.model.supercontig_130.3':0.5902594097281936):0.14216
711312268376):0.013944378585145852):0.05254841457731616,(((('evm.model.supercontig_2154.2':0.5583110019827198,(Al_907868:0.267378809095267
84,(Al_313019:0.04938288823780155,'AT1G21150.1':0.10592701189831778):0.2281074469150426):0.431608004875042):0.16918667753256497,((GSVIV
T01028380001:0.12571215037644606,GSVIVT01028382001a:0.1124011051455523):0.20244421963732553,(GSVIVT01010970001:0.32906817306694114
,((GSVIVT01028383001b:0.0819070897471551,(GSVIVT01028383001c:0.12709439476660508,GSVIVT01028384001:0.0667478055463003):0.029854405
744970492):0.15730315228845387,(GSVIVT01028382001b:0.050463723293734206,((GSVIVT01028382001c:0.07935866514982551,GSVIVT0102838300
1a:0.08319194608542797):0.02206573012181716,(GSVIVT01028382001d:0.08274931088420998,GSVIVT01028382001e:0.07232054465414661):0.01184
7128780915114):0.0399081175431935):0.17597384560851495):0.09747457150802093):0.12918178868790672):0.05089485609600821):0.06602769211049
14,(('evm.model.supercontig_441.3':0.2350813542472581,('Potri.014G133200.1':0.22332346192115363,(Al_487623:0.035380337837681804,'AT5G07900.1':
0.03842277452458452):0.338466119279275):0.03028975273958557):0.1972612641965977,(('Glyma18g13800.1':0.4525357333787078,('Glyma08g41780.1':
0.15815665111868896,'Glyma08g41790.1':0.14040212132139063):0.24361804808333684):0.05538632996387556,('Glyma18g12810.2':0.266418160570816
24,(('Glyma18g13780.2':0.2550076144534777,'Glyma18g13790.2':0.18078061144746554):0.0741558297719796,(('Glyma08g41850.1':0.1562250788387395
7,'Glyma18g13750.1':0.05346667482630756):0.09517567728908861,(('Glyma08g37480.1':0.0499286278284272,'Glyma08g41870.1':0.07528118796709009)
:0.17732690038188512,('Glyma18g13740.1':0.14261190394416443,('Glyma08g41880.1':0.08654768550582072,'Glyma18g13720.1':0.13268330316496693):
0.09488153347864749):0.02814496673492003):0.04201382618078189):0.05283069255889447):0.0939980570042343):0.16113504098815445):0.28506668
75586994):1.34750946315897E-6):0.1337766426038928,(((('evm.model.supercontig_16.125':0.47610904630701584,'evm.model.supercontig_287.2':0.47863
81636407586):0.06421961048075969,('Potri.012G046700.1':0.24579903700811465,('Potri.012G046800.1':0.3148270929123643,('Potri.015G038400.1':0.119
430412328266,'Potri.015G038500.1':0.09265084894143182):0.2415089678782198):0.0517547105590293):0.2598388399396999):0.01445592106711793,(G
SVIVT01017772001:0.7142516548972945,(('Cucsa.253350.1':0.10668588407110785,'Cucsa.253380.1a':0.10245190822363195):0.09519313362849782,('Cu
csa.253320.1':0.1348572762898766,(('Cucsa.253240.1':0.1392213658354197,'Cucsa.253370.1':0.03173968164923063):0.13769812513634447,('Cucsa.25334
0.1':0.1355667481646941,(('Cucsa.253200.1':0.11748286384421212,'Cucsa.253380.1b':0.10720461631582098):0.06832619243643077,('Cucsa.253310.1':0.0
9255869767488704,'Cucsa.253330.1':0.11032682594596552):0.009974314134626343):0.0037573642805839645):0.015498966964636984):0.012270524826
848418):0.037424544123259414):0.4330626221697801):0.10154734547867664):0.037940310903332074,(('Cucsa.172140.1':0.6851263025318257,(('Potri.0
08G216200.1':0.3737125702298404,'Potri.010G022700.1':0.23526889920525507):0.02374941199223421,('Potri.011G005100.1':0.21601938240488042,(('Po
tri.004G012900.1':0.12340068387153413,'Potri.004G013000.1':0.0775525310957458):0.03394518369429175,('Potri.004G013100.1':0.04651735940079841,(
'Potri.004G012400.1':1.34750946315897E-6,'Potri.T104200.1':1.34750946315897E-6):0.024603077493160663):0.029079254078286522):0.08903989414784
465):0.1969367696772892):0.18196584954539838):0.030815863233195948,((('Glyma13g28790.1':0.1756553313968312,'Glyma15g10270.1':0.0963291778
9190075):0.25412517683296015,('Medtr2g019810.1':0.15531579047888677,'Medtr2g019840.1':0.13748004286742552):0.16997836152568155):0.18197862
561534817,((('Glyma15g16400.2':0.32565528704934527,'Glyma15g16430.2':0.25158285467147634):0.11816103771218814,('Medtr4g119570.1':0.35714502
084436806,'Medtr4g119580.1':0.22083019015678204):0.3179343421182528):0.06708742568704329,(('Glyma07g37870.1':0.29543613526226226,('Glyma07
g37970.1':0.5377173886673074,'Medtr4g119550.1':0.21187630353103074):0.10433164315675227):0.11883894315348775,('Medtr2g076320.1':0.527614592
1447551,('Glyma15g16420.2':0.2421719459543505,('Glyma09g05130.2':0.15899934792494308,'Glyma15g16410.3':0.18563599202533107):0.07456163448
105159):0.2337802296115617):0.054640043655971554):0.036639545803892115):0.2068688229682056):0.22572715963857645):0.0502016080595246):0.1
9123796791946793):0.08603150874194632):0.045415915948862864):0.006190517231999321):0.10282887343840265):0.07746454769500277):0.10580124
616711833):0.10745859660066363):0.90396849014633):0.554651068029568);

### Supplementary Data S2. Alignments, trees and control file for PAML analysis of M-class *mTERF* genes.

#### 1) Multiple sequence alignments of paralogous *mTERF* gene sets.

**Osa_1_CDS.phylipi**

5 996

Osa_001

CGCTTCGTCGACGAGGACGCCCTCGTCGCCGCCTGCGGCCTCACCGGAGCCGAAGCCCTCAAGGCCTCGAAGCGTCTACAGAAGGTA------CCC
TCAAATCTAGACGCCGCCCTCACCTTCCTCGCCGACTTCCGCCTCTCCAAAGACGACATCGCCGCCGCAAGCTCACGGTACCCGCGGTTCCTCC
ATCTCAAGGTGGACGAAACCCTAACCTCCCAAGTAGCCCGGCTCCGGGAT---ATCGGCCTATCCACTCCCGAGATTGGCCGCCTCATTACCATCGC
CCCCTGC---ATCCTCTCCAATCCTCGCACGATCTCCCGCCTTGAATTCTACCTCTCCTTTTTGGGCTCCTACCCCAGGGTGCACTCCGCCCTTCGG
AACAATTCCTCCCTTCTACGAAACAACATCGAGAGTGAGGTCAAGCCCAACATTGCATTCCTGGAGCAGTGCGGCCTAACTACTTGTGATATTGC
CAAGATCCTCATGTCTGGCTCTAGGATACTCATTATGCAGCCAGAGCATGTCAAGGAAATTGTGGCGTGTGCAGACAAGTTCGGTATGCCCCGTG
AGTCAGCTGGTTTCAGGTATGCCCTGATGGCTGTTACCGGCATCAGCCCAGTGAGGGTCAGCGCAAAACTTGATTTCTTGAGGATGGTTATTGGA
TGCTCAGATGCCCAGCTACACATTGCGGTGTCTAGGTTTCCACTAATCCTGACATACTCAGAAGTTAAGCTGAGCCGCTCACTGGAATTCCTGAA
AGCGGAGGTTGGGTTGGAGCCTCAGTACATTGTGCTCCGGCCAGCATTGCTCGGCTATAGCATTCAGAAACGGCTGATGCCGCGTTATCATGTCA
TGAAGGTTCTGAATGAAAAGGGCCTGCTGAAGAAAGACACTGATTTTTACTCTATGGTTAAAATAGTTGAAGAGAGCTTTTTTAAGAAGTTCCTT
CTCCCATACCACAGGTCTGTTCCAGGACTTGAAAAAGCTTAT

Osa_002

CCCTTCTCTGTCGAGGAGTACCTCGTCGACACCTGCGGCCTCACCGGCGCCCAGGCGCTCAAAGCATCCAAGAAGCTCTCCCACCTCAGGTCGC
CCGCCAAGCCCGACGCCGTGCTCGCCGTCCTCTCCGGCGTCGGCCTCTCCCGCGCCGACCTCGCCGCCGTCGTCGCCTCCGACCCGCAGCTGCT
CTGCGCCAGGGCCGAC---AACATCGCCCGCCGCATCGCCTCCCTGCGCGACCGCGTGGGCTTGTCGGACCCCCAGATCGGCAGCTTCCTCCTCG
CGGGCGGCGCCAAGGGCATCCACGCGTGTGACGTCGCCTCGAGGCTGGAGTTCTGGATCCCCTTCTTGGGATCCTTCGAGACGCTCCTCAGGAT
CTTGAAGGGGAACAACGTGCTGGTGCTCTCTGACCTTGAGAAGGTGATCAAGCCCAACATCGCGCTGCTCCAGGAGTGCGGGCTAACTGTTTG
TGATATTGCCAAGATGGCCAGGTTTGCTCCACGGATGTTCACGTCCAATCCAAAGCAGGTGGAAGGGTTTGTGCGGCGTGCCGATGAGCTTGGC
GTGCCACGCACCTCAGGCCAATTCAAGTACATGGTGGGTATTTTTGCTAACATCAGCGAAGGCAGTGCCACCGCGAGGATGGAGTACTTGAGTC
GCAGTCTTGGTTGCTCCATGGACAAGCTCCGCTCTGCGGTCCAAAAGTTACCACAAATTCTAGGATTATCTGAGACAAATCTTGGCAGCAAGATA
GAGTTCTTGGTCGGCAAGGTCAGATTGGAGCCTGAATACCTTTTGAAGACGCCAAAGCTGTTCACGTACAGCTTGGAGAAGCGGTTGGTGGCG
AGGCATTATATTGTGCAAGTCCTTGCTGCGAAGGGATTG---AAGAAAGATGTTCCCTTTTGCTCATATGTACAGCTAGGCGAAAGTTGTTTTGTAA
AAAACTTCATTGATCAGCACGAGAACGTTGTCCCTGGTCTTTCTGATGCCTAT

Osa_003

CCCTTCTCCGCCGAGGACTACCTCGTCGCCACCTGCGGCCTCACCGGCGACCAGGCGCTCAAGGCGTCCAAGAAGATCTCCCACCTCAGGTCC
GCCGCCAACCCGGACGCCGTGCTCGCCGTCCTCTCCGGCGTCGGCCTCTCCCGCGCCGACCTCGCCGCCGTCGTCGCCTCCGACCCGCACCTCC
TCTGCGCCAGGCCCGAC---AACGTCTCCCGCCGCGTCACGTCTCTGCGCGACCGCGTGGGCTTGTCGGACCCTCAAATCGGCAGGTTCCTCCTC
GCCGGCGGCGCGATGGCCGTCCGCAAATGCGATGTCGCCGAGCGGCTGGAGTTCTGGATCCCCTTCTTGGGATCCTTCGAGACGCTTCTCAAGA
TGTTGAGGAGGAACAACGCAATTGTCAGAGCTGATGTTGAGAAGGTGATCAAGCCCAACATCGCGCTGTTCCAGGAGAGCGGGCTAACTGTTC
GAGATATTGTCAAGATGCCC---------GGATGGCTGTTCACGTTCAATCCAAAGCGGGTGGAAGCGGCTGTGGAGCGTACTGGCAAGCTTGGTGTC
GAGCTCGCCTCCAGTAGATTAAAGTACATGCTGTCTATTGCTGGCAACATCACCGAAGGCAATGCATCTGCGAGGATGAAGTACTTGAGTAGCAC
TCTTAATTGCTCCATGGACAAGGTCGAGTATATGGTCGGCAAAATGCCAACTATTATAACACTTTCTGAGGAAAAGCTTCGCAGAAAGATAGAGT
TTTTGGTCACCAAAGTCGGGTTGGAGCCTGATTACATTTTGAGTAAGCCTGTTTTGTTCGCGTGCAGCCTGGAGAAGCGGTTGATGCCGAGGCAT
TATATTGTAGAAGTCCTTTTGGCTAAGGGATTGATAAAA---AATGCTGGCTTTTTAACATATGCCATACTGCGGGAGAAGGATTTCGTAGCAAGGT
ACATTGATCAACACAAGAATGCTGTTCCTGGGCTCGCTGATGCCTAT

Osa_004

CCCTTCTCCGTCGAGGAGTACCTCGTCGCCACCTGCGGCCTCACGGGCGCCCAGGCGCTCAAAGCGTCCAAGAAGCTCTCCCACCTCAGGTCG
CCCGCCAAGCCCGACGCCGTGCTCGCCGTCCTCTCCGGCGTCGGCCTCTCCCGCGCCGACCTCGCCGCCGTGGTCGCCGCCGACCCGATGCTGC
TCTGCGCCAGGGCGCGC---AACGTCGCCCGCCGGCTGCATTCGCTCCGCGACCGCGTCGGTCTGTCCGATGCCGACGTCGCCCGTTTCCTCCTCG
CCGGCGGCGCAATGGGCCTCCGCAAATGCGACATCGCCCCGAGGCTCGAGTTCTGGATCGGCTTCGTCGGATCGTTCGACAAGCTCCTACCGGC

ACTGAAGGGGAACAACGGAATCCTCATGAGTGACCTGGACAAGGTCGTGAAGCCGAACATCGCGCTGCTCCAGGAGTGCGGGCTAAGTGTTTG
TGAAATTGCCAAGCTGTCTACGCTCAAGTGGACGGTGCTGTCGTTGAGTCCAGAGCGAGTCAAGGCGTCGGTGCTGTGCGTCGAGAAGTTGGT
CGTGCCCCGCTCCTCGGATCGGTTCAAGCACGTGCTCAAGTCCGCCTGTTGGATCTCCGAAGATATGCTCGCCATGAAAATGGAGTTCTTGAGG
AGCACTCTTGGTTGCTCCGAGGACAAGCTCCGTGCCGCGGTCTGCATTTCGCCACATATTTTCTACCTGTCCGACAAGAACCTTTGCCGAAAGAT
AGACTTCCTGATCTCCGAGGTTGGCTTGGAGCGTGAGTTCATTGTGGAAAGGCCTTGGGTGCTCGGCTACAGCCTCGAGAAGCGTATGGTGCCA
CGGCATTCTGTCATGAAGATCCTGAGGACCATGGGATTGATGAAGGATGCTGTTGACTTCAGCAGTTCACTTGTTTACTCTGAGAAAAAATTCGT
TGCAAGATACATTGATCCTTACAAGCAAGCAGCTCCCACGCTCGCAGATTCCTAC

Osa_005

CCCTTCTCCGTCGAGCACTACCTCATCGCCACCTGCGGCCTCACCGCCGCTCAGGCGCGCAGGGCGTCCCCGAAGCTCTCCCGTCTCAACTCCT
CCTCCAACCCCGACGCCGTGCTCGCCCTCCTCTCCTCCCTCTCCCTCTCCCGCGCGGACCTCGCCGCCGTCGTCGCCGCCGAGCCGCGGCTGCT
CCGCGCGAGGCCCGGC---ACCATCGCCCGCCGCATCGCCTCCCTCCGCGGCCGCGCCAACCTGTCTGCGCCCCAGATCCGCAGCTTCCTCATGTC
CGGCGGCGCGGCGCACCTCGCCTCGTCCGACGTCTCCCCGAAGCTCGCGTTCTGGGTCCCCTTCCTCGGCTCCTTCGACATGCTCCTCAAGATC
CTCAGGAGGTGCAACGCGATCCTCGCCACCGACGTCGACAGGGTGGTCAGGCCCAACGTCGCGCTGCTCGGGGAGTGCGGGCTAGGTGTTTGT
GATATTGTCCAGATGACCCAGACCGCCGCGTGGCTGCTCACGTTCAACCCAGAGCGACTGAAGATTGTCGTGCGGCGCGCCGAGGAGCTCGGC
GTGCCCACTTCATCGTGGGCGTTCAAGGACGCCGTGTGCACCGTCGCCCGTAACAACGAGGGCACCATCGCCGCGAGGATGGAGTTCTTGCGC
GGCACTCTTGGTTGCTCCATGGACAAGCTACGCTCTGCGATCAGCAGGAAGCCGAGCATTCTAGGATTCTCGGAGAAGACGCTTCGCGGCAAAA
TAGAGTTCTTGCTCACTAAGGTCCAATTGGAGACTGAATACATCTTGCAGAGGCCTGTGATGCTCACGCTCAGCCTGGACAAGCGGTTGGCGCC
GCGGCATTATGTTCTACAAGCCCTTGTGGAGAAGGGGTTGATAAAAAATGATGTTGATTACTACAGTTGTGTTTGTTTTGGCAATGAACATTTCGT
AGCGAGGTACATTGATCGCCACGAGGATGCTCTTCCTGGTCTCACGGATGCCTAT

**Osa_2_CDS.phylipi**

9    996

Osa_006

CACTTCTCTGTCGAGGAATACCTCATCTCCAACTGCAATCTCACCCAGCCACAGGCCCACAAGGCGTCCAAGAGCATCGCCCACCTCAAGTCCC
GCTCCAACCCCGACGCCGTCCTCGCCTTCCTCGCCGACTTCGGCCTCTCCCCCAAGGAGGTCGCGGCCATTGTTGCCTCCAACCCGCGCATCCT
GTGCGCCCGCATCGACCGCTCCCTTGCACCCATCTGCTCGGAGCTCCGCGCC---GTCGGCCTCTCCCCCTCCCAGATCGCCCGCCTCGCCCAGAT
CACCGGCCGC---TACTTCCTCTGCCGCAGCTTCGTCTCCAAGGTGCGGTTCTGGCTCCCCCTCTTCGGCTCATCGGAGAGGCTTCTCCAGGCCAG
CGACTGGAACTACTGGCTCCTCACCTCCGACCTCGAGAAGGTGGTCGAGCCCAACGTCTCCTTCCTCAAGGAATGCGGGCTAAGTGCTCGTGAT
ATTTCCAAGCTGCTCGTGGCCGCGCCGCGGCTTGTCACCATGCACCCGGATTACGTCAAGGACGCCGTGCGGCGTGCCATCCAGCTCGGCGTGG
CCCCGGGCTCCCAGATGTTCCGGCACGCGCTCTCCACCGCCGGCTGCATCGGCCAGGACAAGATCGACGCCAAGGTCGCCGTCCTGAAGGAGA
GCCTCGGGTGGTCGCAGGAGGAGGTGAACCTGGCCGTGAGCAAGGCGCCGCGGATCCTGGTCGCGTCCGAGGAGAGGCTGCGCCGCAACGCG
GAGTTCTTGATCGACGAGGTCGGGCTGCAGCCGCAGTACGTCGCGCGGCGGTCGGTGCTGCTCATGTACAGCCTCGAGAGGAGGCTGGTGCCC
CGCCATCTCGTGGTGAAACTTCTCAAGGAGAGGGGTCTGATCGAGCAAGACCGGTGCTTCTTCAACGCGGTGGCGCCAACTGAAGAGAAGTTC
TTGGAGAAGTTTGTTGTCCCCTTTGAGGGGTGCGTCCCGGGTCTCGCCGACGCCTAC

Osa_007

CCCTTCGCCGTCGAGGACTACCTCGTCGACTCCTGCGGCCTGACGAGAGCCCGAGCCAAGAAGGCCTCCGGGAAGCTCTCCCACCTCAGGTCC
CCCTCCAACCCCGATGCCGTCCTCGCCTTCCTCTCCGGCCTCGGCCTCTCCCGCCCGGACATCGCCGCCGTGGTCGTCAACGACCCGCTCTTCAT
CTGCGCCAGGGTGGACAAGACGCTGGCCACCCGCGTCGCCGAGCTCACCGAC---CTCGGCCTGTCCCGCTCCCAGATCGCGCGCCTCATCCCGG
TGGTCCGCAGC---CTCTTCCGCTGCAAGTCGCTCGCCCCAAGGCTCGCCTTCTTGCTCACCGTGTTTGGCTCCTTCGACAGGTGCCTCGAGGTTA
TCAAGACCAACTATGGCGTCCTTAGCTCCAACGTTGAGGCGGTCATCAAGCCCAACCTCGCCGTACTCAAGGAATGCGGGATATCTATAGCGGAT
AGGCCCTCATACGCG---TTCGCATCGCGGGTGATCAGCAGGCCTACTAAGCACCTGGAAGAAGCTGTGGTGCTTGCTAACGAGTTTGGAGCTAA
GCAGGGAACTCGGGTGTTTACTAATGCGGTCATGATTTTTGGAATCCTCGGACAGGAGAAGCTCGCCAAGAAGTTGGAGTTCTTCAAGAAG---C
TTGGTTGGTCACAGGACGATTTGTCATTGGCGGTGAGGAGCATGCCACACATCCTAGCCATGAAGGAAGAAAGGATGCGTCGAGGTATGAAGTT
CTTGACCGAGGATGTTGGTCTAGAGATACCATACATTGCACGAAGGCCTGCACTCACGATGTATAGCATTGAGCGCCGGTTGTTGCCACGGCATT
GCTTGATCAATGTTCTCAAGGGGAACGGATTGCTCAAGGCCAATTATGACTTCTATAATATCTCTGTGATTAGCAACGATGATTTCATGGAAAAAT
TCGTGCAACCATATGTGGAGAGTGTTCCAGGCCTTGGTGATGCATAT

Osa_008

CCCTTCGCCGTCGAGGACTACCTCGTGGAGTCCTGCGGCCTGACGAGAGCCCGCGCGGAGAAGGTCTCCGGGAAGCTCTCCCACCTCAGGTCC
CCCTCCAAGCCCGACGCCGTCCTCGCCATCCTCTCCGGCCTCGGCCTCACGCGCCCCGACATCGCCGCCGCCGTCGCCTCCGACCCAAGGCTGC
TGTGCGCCCGTGTGGACAGGACGCTGGATGCCCGCGTCGCCGAGCTCGGCGGC---ATCGGCCTCTCCCGCTCCCAGATCGCGCGCCTCATCCCG
CTCGCTCGCGGC---GGATTCCGCATCAAGTCTCTTGGTTCAAAGCTCGCCTTCTTGGTCACA-------------------------------------------------------------------
---------------------------------------------------------------------------------------------------------------------------------------GATGAACTGTGGGCAATTAAGCCGGGA
ACACGTCTATTTGCTCTCGCAGTTGTGAAGTTTGCAATCCTCAGCCAGGGGAAGATCACCAAGAAATCAGGGCTGTTTAAGAAG---CTTGGATGG
TCACAGGAAGATTTGTCATTAGCAGCGAAGAACATGCCAAGTATCCTTGCCATGGGTGAGAAAAGGTTGAGGCAAAGAATGAAGTTCTTGACG
GAGGATGTTGGTCTAGAGATACCATACATTGCACAAAGGCCTGCACTCATGTTTTATAGCATTGAGCGCCGCTTATTGCCAAGGCATTGCTTGATC
AATGTTCTCAAGAGGAACGGATTGCTTAAGATCAATTATGACTTCTATTCTACTGCCTTGATTAGCAATGAGAAATTTCTGGATAAATTCGTGCATC
CTTATGTGGAGAGTGTTCCAGGCATTGGTGATGCCTAT

Osa_009

CCCTTCGCCGTCGAGGACTACCTCGTCGCCGGCTGCGGCCTCACCCGAGCCGAAGCAGCCAAGGCTTCGGCGAAGATCTCCCACCTCAGCTCC
CCCTCCAATCCCGACGCCGTCATCGCCTTCCTCTCCGACCTCGGCCTCCCCCGCCCCCAAATCGCCGCCGCCATCGCCGCCGACCCGAGGTTGCT
GTGCGCGGACGTGGAGAAAAATCTCGCCAAACGCGTCGGCGAGCTCGGCGAC---CTGGGCTTCTCCCGCTCCCAGATCGCGCGCCTCCTCCCGC
TCGCCGGCTGG---TGCTTCCGCAGCAGCTCCCTCGCCACCAACCTCGCCTTCTGGCTCCCGGTCTTCGGCTCGTTCGATAAGATCCTCAAGGCTC
TCAGGATGAACAAAAACCTCCTCAGCCCCGGCGTCCAGAAGTCGGCGAAGCCGATCCTGGCCTTCCTCGAGCAATGCGGGATCAATGCCTCTGA
TGTTGCCCGTAGTTCAATGTACTCCAGCCGGCTGCTCACCGCGAATCCCGAGTACCTCCGAGACGCCGTGGCACGGGTGGAGGAGTTGGGCCTG
GACCGCAGTTCGAGGAGGTTCCACCGTGGGCTCGTTGCGGTGGCTTTGGTAAGCAAGGAGACTGCTGCGAGGAAGATACGATTGATGGAGGAG
---CTTGGGTTTTCGCAGGATGATCTGTTGGTGATCATGAGGAAGTTGCCAAACTTTTTGGCTTTATCCGAGAAAAAGATACGGCGAGCTGTGGAG
TTCTTGAAGAGGGATGTTGGTTTGGAGGGACGGTACATTGTGCAAAGGCCAGTGTTGTTGTCGTATAGCCTTGAACGGCGGCTGTTGCCTCGGC
ATTGCTTGCTCAAGGTCCTCAGGACAAAGGGATTGCTGAATAGTGAGTTGGATTACTACTCCACAGCTGCATTGAGCGAGAAGAAGTTTGTCAA
CAAGTTTGTGCATCCTTATGAGGACCACATTGCAGGCCTTGCTGATGCTTAT

Osa_010

CCCTTCGCCGCCGAGGACTACCTCGTCGCCGCCTGCGGCCTCACCCGAGCCCAAGCAGCCAGGGCTTCGGAGAGGATCTCCCACCTCAGGTCC
CCCTCCAAGCCCGACGCCGTCCTCGCCTTCCTCGCCGGCCTCGGCATCCCGCGCCCCGACATCGCCACCGCCGTCGCCGCCGACCCGAGGCTGC
TGTGCGCCGGCGTGGAGGGAAACCTCGCCAAGCGCGTCGCCGAGCTCGGCGAC---CTGGGCATCCCGCGCTCCCAGATCGCGCGCCTCGTCCCC
CTCGCCAAGATC---CCCTTCCGCAGCAGCTCCCTCGCCACCAACCTCGCCTTCTGGCTCCCGGTGTTCGGCTCGTTGGACAGCATCCTCAGGGCC
CTCAGGAAGAACTCCTCCCTCCTCAGCGCCAACCTCGACAAGGTGGTGAAGCCGAACCTGGCATTCCTCAAGCAATGCGGGATAGATGCCCGT
GATGTTGCCAGCAATCCACTGTACTCCAGCCGGCTGTTCACCTCGAATCCCATGAAACTCCGGGACGCCGTGGCGCGAGTGGAGGAGCTAGGCA
TGGTCCGCGGCTCGAGGGTGTTCCACCGTGGGCTCGTTGCGGTGGCTTTCCTAAGCAAGGAGGCTGTCGCGACGAAAACACGGCTGCTGGTGG
AG---CTTGGGTTTTCGCAGGATGATGTTTCGGTGATCTTCAGGAAGATGCCATCCTTCCTGACCGCATCCGAGAAAAGGATACGGCGAGCCGTGG
GGTTCTTGAAGGGGGATGTTGGTCTGGAGGAACGGTACATTGCGCGAAGGCCGGTATTGCTCTTGTATAGCCTTGAACGGCGGCTATTGCCTCGG
TATTACTTGCTCAAGGTCCTCAGGACAAAGGGATTGCTGGATTGCAAGCTGTGTTACTACTCAACAGCCGCACTGGGCGAGAAGAAGTTTATAG
AAAGGTTTGTGCATCCTTATGAGGACCACATTGCAGGCCTTGCTGATGCTTAT

Osa_011

CCCTTCGCCGTCGACGAGTACCTCGTCGCCACCTGCGGCCTCACCCGAGCCCAGGCAGCCAAGGCTTCGGAGAAGCTCTCCAACCTCAGGTCC
CCCTCCAATCCCGACGCCGTCCTCGCCTTCCTCTCCGACCTCGGCCTCTCGCGCCCCGACATCGCCGCCGCCGTCGCCGCCGACCCAAGGCTGC
TGTGCGCCGACGTGGGGAGCAGCCTCGCCAGGCGCGTCGACGAGCTCGGCGGC---CTGGGCCTCTCCCGCTCCCAGATCGCGCGCCTCCTCCCG
CTCGCCGGCAGG---TGCTTCCGGAGCAGCTCCCTCGCCACAAGGCTCGCCTTCTGGCACCCGGTGTTCGGCTCGTTCGAGAATATCCTCAAGGC
GCTCAAGATGAACGCCGCCCTCCTCGGCTCCGACCTCGACAAGGTGGCGAAGCCGAACCTGGCCTTCCTCGCGCAATGCGGGATAAACGCCTC
CGATGTCACTCGCACAACCCTGTACTCCTGCCGGCTCTTCACCGTGAACCCTAGGTTCCTCCAGGACGCTGTGGCAAGAGTGGAGGAGTTAGGC
GTGGCCCGTGGTTGGAGGACGTTCCACCGCGTGCTCAGTACAGTGGCTTTCTTAAGCAGGGAGACTATTGCCAGTAAAATGCAGCTGCTGGATG
AT---CTTGGGTTTTCGCAGGATGATTTCCTGGTAATCGTAAGGAGGGCGCCACAAGTGCTGCGTTTATCCGATGGAAGGATACGGCGATCTGTTGA
GTTCTTGATAAGGGATGTCGGTCTGGAGCAAAGCTACATCGCACAAAGGCCAACGTTGTTGGCGTATAGCCTTGAGCGGCGGTTGTTGCCTCGG
CATTGCTTGCTCAAGGTCCTCAAGGCAAAGGGATTGCTGAACTGTGACTTGTCTTACTACTGCATAGCCGCGATGAGCGAGGAGAAGTTTGTGC
AAAGGTTTGTGGATCCTTTTAAGGACAAAATTCAAGGCCTTGCTGATGCTTAT

Osa_012

CCCTTCGCCGTCGAGGACTACCTGGTCACCACCTGCGGCCTCACCGGGGACCAAGCGCGGAAGGCGGCCAAGACTCTCTCCCGCCTCAGGTCC

CCCTCCAAGCCCGACGCCGCCGTCGCCTTCCTCTCCGGCCTCGGCCTCTCGCGCTCCGGCATCGCCGCCGCCGTGGCCGCCGACCCGAGGTTGC

TGTGCGCCGACGTGGAGAAGAACCTGGCCAAGCGCGTCGCCGAGCTCGGTGAG---CTGGGCATCTCGCGCTCCCAGATCGCGCGCCTCATCCCG

CTCGCCCGTCAG---TCCTTCCGCAGCAGCTCGCTCGCCACGAACCTTGGCTTCTGGCTCCCGGTCTTGGGCTCTTTCGAGAATGTGCTCATGGCG

CTCAAGGCGAACGGAGCGATTCTTGGCTCCGACGTCGAGAAGGTGGTCAAGCCCAACCTGGCCCTCCTCCAGCAATGTGGGATACATGTCTGC

GATTTCCCCCATACACGC------CTGCCCACTGTGCTCTGCAGGCCACCCAACCATGTTCAAGAAGCTGTGGCACGCATCGGTGAGTTTGGCGTGC

CACAGTACTCACCGGTGTTCCGCAATGCGCTTGTGCCATTTGCTTACCAAAACAAGGAGAAGCTCGCTGCCAAAATTGGAGTCCTTGAAATG---T

TTGGGTGGTCAGAGGACGATCTCTCGATGACTATGAGGAAGGGTCCTGTCGTCATGAACATGTCTGTGGAAAGGCTGAGGAAAAATGTGGAGTT

CTTGACAAGGGATGTTAAGCTCGAGACGCGTTACATTGCTCGAAGGCCAATCATGATCAGTTATAGCCTTGAGCGTCGGTTGTTACCCCGGCATC

GCTTGCTCAGGTTTCTCAGTGCAAAGGGATTGCTTGATGGTGAATTGGACTTCTATTCAGCGGTTGCATTGACTGAGAAGAAGTTTCTTGACAAG

TTTGTGCATTCTTGCAAGTGTAGCATTGCAGATCCTGCTAATGCTTAT

Osa_013

CACTTCGCCGTCGAGGAGTACCTCGTGGCCACCTGCCACCTCACCCCCGACCAGGCTACCAAGGCCTCCAAGTCCATCTCCCACCTCAAGTCCC

CGTCCAGGCCCGACGCCGTCGTCGCCTTCCTCGCCGGCCTCGGCCTCTCCGCCGCCGACATCGCCGCCGCCGTCGCCTACGACCCGCGCCTCCT

CTGCGCGGAGGTCGACAGGACGCTCGCCCCGCGCCTCGCGGAGCTCGCGGGC---CTCGGCCTCTCCCCCTCGCAGATCGCGCGCCTCGTCCTCG

TCGACCCCGCC---CGCTTCCGCCGCCCCACCGTCATCTCCAAGCTCCAGTACTACGTCCCGCTGTTCGGATCCTTCGAGACCCTCCTCCAGGCGC

TCAAGAACAACTCCTACCTCCTCAGCTCCGACCTCGAGAAGGTTGTCAAGCCCAACGTCGCGCTCCTGCGGGAGTGCGGGCTAGGTGCTTGTG

ATATTGCTAAGCTGTGCATCCCGTTGCCGAGGCTGCTCACCACCAGCCCAGAGCGTGTCCGAGATATGGTGGCTCAAGCAGAAAACGTCGGCGT

GCGCCGTGGCTCCAAGATGTTCAGGCACGCTATTCTGGCCGTCGCGTACATCAGCGAGGAAAAGATCGCCGCCAAAATGCAGTTCTTGATGAAG

ACATTAAAGTGGTCAGACGCAGAGGCGAGGATTGCGGTGTCCAAGCTTCCTGTGGTGCTGAGGAGCTCGGAGGACAAGCTGAGCCGTGTGTCG

GAATTCTTGATCTCTGAGGTCGGGTTGGAGCCGGCATACATTGCTTACAGGCCGGCAATGCTCACTTATAGCCTGGAACGCCGGCTCATGCCCCG

GCACTGCGTTCTGAAGTATCTTAAGGACAATGGATTGATAGAATCCGACAAGAGCTACTATAGCGCGGTCCAGGTGACCGAGGAGGTCTTTGTG

GAGAAATACATAAGTCCATACGAGGACACTGCGCCGCACCTTGCTGAAGACTAT

Osa_014

CCCTTCGCCGTCGAGGACTACCTCGTCGCCACCTGGGGCCTCACCGGAGCCCAAGCGCGCAAGGCCTCGAAGAAGCTCTCCCACCTCAGGTCC

CCCTCCAAGCCCGACGCCGTCCTCGCCTTCCTCTCCGACCTCGGCCTCCCGCCCCGCAAGATCGCCGCCGTCGCCACCGCAGACCCGCGCTTCC

TCTGCGCCGACGTGGAGAGCAACCTCGCCAGGCGCGTCGACGAGCTCGGCGGC---CTGGGACTCTCCCGCTCCCAGATCGCGCGCCTCGTCCCG

CTCGCCCTCACC---TGCTTCCGGTCCAGCTCCGTCGGCACGAATCTAGGGTTCTGGCTCCAGATCGTGGGCTCGTTCGACAAGATCCTCAAGGCC

CTCAGGATGAACTCCTCGCTACTCGGCTCCGACCTCGAGAAGGTGGTGAAGCCGAACCTGGAATTGCTAAAGCAATGTGGGATGAGT------GATT

TTGCCACTAGTTTCTTGTACACCAGCAGGCTCTTCACCGCGAACCCTATCTATCTCCGGGACGCCGTGGCACGGGTGGAGGAGTTAGGCCTGGA

CCGCAGCTCGAGGATGTTCCGGCATGGGCTCATCGCGGTGGCTTTCACGAGCAAGGAGTCCGTTGCGAGGAAGATACAGGTGATGGAGGAG---C

TTGGGTTTTCGCGGGATGAATTGTTGATGATCATCAGGAAGGCACCGCAGTTGGTGGCCTCATCGGAGGAAAAGATACGGCAAGCCGCTGAATT

CTTGAAGAGGGATGTTGGTTTGGAGGGACGGTACATTGCGCATAGGCCAGTGTTGTTTTTGTACAGCCTTGAGCGGCGATTGTTGCCTCGGCATC

ACTTGCTCAAGGTTCTCAGGATGAAGGGATTGCTGGATTGTGAGTTGGATTACTACAACACAGCCGCGATGAGTGAGAGGAAGTTTGTGCGGAA

GTTTGTGGATCCTTACAAGTGCCACATTCCAGGCCTCGCTGATGCTTAT

### Osa_3_CDS.phylipi

4 996

Osa_015

TCCGCCGCCGCGGCCTCCTACCTCGTCGCCTCCGTCGGCCTCTCCCCGGCCGCCGCCGCCAGGATCTCC---CGCAAGGCACGCTTCCGTTCCAAC

GCCGACGCCGAAGCCGTCGTCTCCCTCCTCCGCGGCCACGGCTTCTCCGACGCCAACATCGCGCAGGTGCTCCCAAAAATCCCGGGCCTCCTCA

TCCTCAACCCGGACAAGATCCTCCGGCCAAAGCTCGAGTACTTCGCCTCC---CTCGGGGTGGTCCCCTCGGCACTGTCCCGCGCACCCCTCCTCG

CGCGCAGCCTG---------GAGAAGCACCTCGTCCCGTGCGTCGAGTTCATCCGCGGCGTCGTTGGCACCGACGCCAACCTCTGCGCGGCCATCTCC

CGGAACCCCTGGGCACTCTGGTGCGACATTAACAGCAGCATGCGCCCCGCCGTGGAATCCCTTCGCCGCCACGGCCTCGCTGAGGCGAACATCT

CCAGGCTCGTCGTCATCAATCTCAGCGCTCTCACGATGTCACCCGACCGCATTGACGGGATCTTTGGGGACCTGGAGGCGCTCGAGCTGCCCAT

CTCGCATTCGCGCTTCGTGTACGGCTTCTGGGCACTGTCCAGACTCAAGAGGGGGGCATGGGAGGAACGGATGTCCGTGTTCATGAGA---TTTG

GGGTTTCCAGGAGCGAGTTGCTCAAGGCATTCAGGGAGCAGCCCGGTATACTAGTGTTCACAGCTAAAACTATCCAGCGGAAGCTCAGTTTCTA
CCAGGAAAAGCTGAAGGTTGCACCGGCAGATGTGATTGCACATCCTCTTCTCTTGACGTTCAGCTTGGAGAAGAACATCATACCTAAATGTGCC
GTGCTGAATGTGCTATTGAGGGAGGGGAAGATTAAGAGAGAGATGGACTTGTTGAGACCCTTGCAGCGTAGCAATATAAGTTTCTTTGAGAGAT
TTGTGAGGAAGTATGAGGAAGATGTGCCAGATGTTGTAAAAGCGTAC

Osa_016

CCATGCCCCGACACCTCTTACCTGGTCTCC---TGCGGCCTCCCCCCGGCCGTTGCCCGTCACACGGCCGCCCGCGGCCTCCGGATCCGATCCACC
GAGAAGGCGGACGCCGTCCGCACCCTCCTCCGCAGCTACGGCTTCTCCGACGCGGACGTCGCCCGGATTGCCCGCAGCGCCCCGCTGCTCCTC
ACCGTCGATCCCGACCGCATCATCCGCCCCAAGCTCGAGTTCTTCGCCACG---ATGGGGTTCCAGCCCAGCAAGCTGTCCACCGCGCCGCTCCTC
CTCGCGCGCAGCCTC------GAGAAGCACCTCGTCCCCACCATCCAGTTCCTCCGCAGCATCATCGGCAGCGACGACGGGATCCGCCGCGGGTTCT
CCCGCATCCCTCGCGCACTCCTGGTCAGCCTGGACAACTGCATGCGCCCCGCGGTGGAAGCCCTCCACAGGCACGGCCTCACCGGCCGCGACG
TGTCCAAAGTCCTCGTCCTCCAGATGGGCGTGCTCATGCTGTCGCCGGTCCGCATCGGCGAAATCTTCGAGGATCTCAAGGCTATGGGTATGTCC
ATCACGGACGGCCGCTTTGCCAACTCGTTCCGCGCGATGTGCAGCATGAGGAGGGCGACCTGGCTGCGGAAGGTGGCGCTGTACCGGAGC---TT
TGGCTTGTCCGAGAGTGAGGTGTTCGAGGCCTTCAAGAAGCAGCCCACGGCACTGCTTGGCGCTGACGAGACCATAAAAAAGAATGCGTCGTT
CTTCCGCGACGCGCTAAAGCTTGAAATGCGCGAGGTAATGGTGCATCCTGTTGTTATGGCATATAGCTTTGAGAAGACCATCCTGCCAAGGTGTG
CCGTGTTGAGCGTGTTGATGAGAGAGGGGAAGATTAATCCAGATATCCAGTTGCTTCATGCATTGCTTGGCAGTGCCAAGACATTCTCAGGGAG
GTATGTAGATAGGTTTGCTGCTGATGTGCCAGACGTTGTTGAAGCATAT

Osa_017

CCGAAGGTCGTA---TCCTACCTGATATCCTCCTGTGGCCTCACCCCTGCCGCCGCCGCCCGCGCCGCCGCCCCCTGGCTCCCCCTCGCCTCTCCG
AGCAACGCCGACGCCGTCGTCGCGCTCCTCCGCCGCTACGGGTTCACCGACGCCGACATCTCCGCCACCGTCCGGGCGTTCTCCAGGATCCTCG
CGTCCGACCCCGCCCGGACCCTCCAGCCCAAGCTCGACTACCTCCGCTCC---GTCGGCATCACGGCGCCGCTCCTCCCCCGGGTCGTCTCCCTCA
GCCCCGTC---ATCCTCCACGAGTCCCACCTTGCCCCGCTCATCGCCTCCCTCCGCGAGGTCCTCGGCTCCGACTCGCGCATCGTCACCGCGCTCC
GCCAGATGCCGTTCGCCATGCGCTGCAGCCCCAAGGCCACCTTCCTCCGGACGCTCCCCGTGCTCCGCGACCACGGCCTTACGCCCAGCGAGCT
GTCCAAGCTCGTCGCCAGCCAGCCCGGGGTCATCCTGCTGGGCCCCGGCCGCGCCGGGGAGATCGTCCAGGCCGTCAAGGACGCCGGCGTCGA
GCCAGGAAGCCCCATGTTCGTCTACATCTTCGCCGCCTTCTCCAAGCTGAAGGCCCCCACGCTGGAGAACAAGTTCGCGATCTACCGCAGC---CT
CGGCTTCGGCAAGGACGACATCGCCGTGATGCTGCGGCGGCTCCCGAACGCGGCGGGGATCTCGGAGGAGAGGCTGAAGAGGACCGTCGGCT
TCTTGACCGGCAAGGCGGGGCTGCGCCGCGAGGACATCGTCGCGTACCCGAACCTGCTGTCGCGGAGCCTGGACTCCCAC---GCCCGGAGGTG
CGCGGTGCTCGCCGTGCTGAGGAGGGAAGGGAAGCCGGAGGGGCAGCACCGCGTGCCCCATGTGCTCGTGGCCAGCCTCGCACGGTTCATGA
AGGCGTACGTGAGGCGGTACGAGGGGGAGGTCCCCGATGTCTTGCGGGCTATC

Osa_018

CCCTGCTCCCTCACGCACTTCCTCCGCAACACCTGCGGCCTCTCGGAGGATGAGGCGGCCGCGGCCGCCGCGCGCGTC---CGCCTGCGCTCCAC
CAAGAAGGCGCACGCCATCGTCGCCCTCTTCCGCGGCATCGGCTTCTCGGCCGCCGACATCGCCCGCCTCGTCACCTCGAACCCGTCCTTGCTC
TCTTACCGGGCCGACGCCACCCTCATGCCCAAGATCGAGTTCTTCCGCCGCGAGCTCGGCCTCACCGACGCCGAGATCCGCCGCCTCGTCCTCG
CCAACCCCTACCGCGTCCTCCGTAAGCGCTGCATCCGCCCCAACTACCTCATCCTCAGAGACCTCCTCGGCAGCGACAAGAACGTGACCGCTGC
CGTCTTGCAGTCCACGGATCTCATCCACGGGGACGTTCGTGGCATCCTCCTCCCCAAGATCAAGATCCTCCAGGACTACGGCGCCACGAACGAT
GTCATCGTCAAGCTGGTCACCACGCACCCCAGGGCACTCATGCATAGGGCCTCCCGCTTCGAGGAGAGCTTGGCTGCCATGAAGGAGCTCGGG
GTCAGACCTTCCTCTGGAATGTTCCCCTATTCCTTCGGGCTCTTTGCCAGATTGCATCCAAGGAAATGGAAGGGCAGGATGGACAATTTCTTGAG
C---TTGGGATGGACAAAGGAGCAAGTGATTGAGGCGTTTGTGAGGCACCCGTACTGCATGTCCGTGTCGAATGACAAGGTGAAGCTCATCTGGC
AGTTTCTCGCGAAGAAGCTCAGGTGGACCACCGATTATGTCGCAAGGAGCCCGATGGTTCTCTCCTTCAGCTACGATAAGCGAATCTTGCCAAG
GTGCACGGTGCTGAACTTGTTGGCATCGAGGGGCATATTCAACAGGGACATCAAG---ACATCCCATCTGGTACTGGGGGAGAAGAAGTTTAAGG
AGAAGTATGTCACCCCTTACCAGGATGAAATTCCTGAGGTACTGGAAGCTTAT

**Ptr_1_CDS.phylipi**

15 1020

Ptr001

TCATTTACAGTTGAGTGCCTCGTAAACTCATGTGGGCTTCCCTCAAAATCAGCTCTTGAATTCTCCCGAGACTTCCACCTGCATGAAAACAACCT
CCAAAGCTTCCAATCTGTATTCCGCTGCTTCCAGTCCCACAACATTCCCAGCATCCGCATAACCAAATTGATTAAAAGGCGGCCTCAAATCCTCA
ATTACAATGTAGAAGACAACCTCAAGCCTAAACTCCAGCTCCTCGTGCAAAATGGCATCGTGGGTCATCACATGTGCAAGGTTTTCGTATCAAAT

CCGGTAATTTTGAACGCTGACTTAGATTCTCAAATTAAACCATGTTTTCAGTTTTTAAAGTCTGTTCTTGGTAGTAATAGGAATGTCGTAGAAGCTA
TCAATCGTTCTTCCAATTTGCTGACTTGTGATTTGAAGGGTTGTTTGAAACCAAATATTGATTTTTTGATTAGAGAGGGAGTGCCTTTTGATGGGG
TTGCTGAATTTCTTATTAGGGACGCAATAACTGTACAGCACAAGCATAATAGTATGGTCAATGCAGTGAATGATCTTAAGAATCTGGGATTTGATC
CAAAGGCTCCTGTTTTTCTTGAGGCTGTTAGAGTGAGGATTCACATGAGTGAGTCAATTTGGAGGGAGAAAATTGAAGTGATGAAGAGTTTGGG
GTGGAGTGAAGAGGAGATTTTTTCGGCTTTTAAGCGAGATCCGATTTTTTTAAAATCCCCAGTAGAGAAAATCAGGGTGGCGACGGATTTCTTTG
TGAATACATTGAAATTGGGACGGCAAATTTTAAGTGAAGATCCTGAATTTTTTACCCTTAAAATTGATAAAAGTTGTCGAAGGAGGTATGATGTTT
TCAAACTTTTGGAGTCGGAGAAGCTGCTCGAAGGTGGTGTGAAGATTGAGGAAGTGCTCAAAATGAGGGATAAGGAGTTCTTGGTGAAATACG
TTAAGAAGTATGTGGACAAAGTCCCAGGTTTATGGGAGACTTTCAATGGTAGGAAGCAACAACAATCT

Ptr002

TCGTTTACACTTCAATTTCTTGTTAACTCATGTGGGCTTCCTTTAGAAACTGCTCTTACTGCATCCAAGAAGTTCCATCTCAACGAGAAGAACATC
CACAAGACTCAATCTTTGCTCCGTTTCTTGAAATCTCACCACTTTGTGGACACGCATATCGCCGATCTGATAAAAAAGATGCCCGCTTTCCTTGGT
TGTAAAGTAGAAAACAATCTAAAGCCCAAGTTCGAGTTCCTCGCCGCAAATGGCGTTGCTGGTAACCTCCTTCCCCAACTTATCCTATCAAATCC
TGATATTTTGTGGACGTCCTTGGATTCTACTATCAAACCATCTTTTAAGTTTTTGAAGTCTTTTCTTGGTACCAATGTGAAAATTACAGCGGCTCTT
AAGCGTTGTTCATGGTTGTTGAGAGTTAATCGGAACAGAACTATGCAAACAAATATTGATTTGTTGATTAAAGAAGGATTGCCTCTTGACCGGCT
TGCAAAACTCATTATCTCGAGTCCAAGATCTCTGCTATCTAAGCATGATAAGATAGTTTATGCAGTGAATTCTGTTAAGAATCTGGGCCTTGAAAC
AAACGATACTATGTTCATATATGCTCTTGGGGTGAAGATGAAAATGACTGACACGACTTGGAAGAAGAAAATTGAGGTGATGAAGAGTTTGGGT
TGGAGTGAAGAGGAGATTTTCGGCACTTTCAAGCGATGCCCTCAAATATTGCAATACTCGGAGAAGAAAATCAGGATTACCGTGGATTTTTTTAT
CAATTCCGTTGATTTGGGACCAGAAATTCTACTGGTCTATCCTTCTTTGTTTTGCCTCTCAGTTGATAAGAGGGTTCGTCCATGGTATAATGTTATT
AATGTTTTGAAGTCAAAGAACTTGATTAAAAGAGAAAAGAAGTTTCCTTCTTTGCTACTGATGAGTGAGAAGAAATTCTTGGAGAATTATGTTGA
TAAGTATGCGGATGATGTTCCTGGTTTATGGGAGGTGTATACGGGCACTGCCAACACAAAGAAG

Ptr003

TCATTTGATGTTCATTTCCTCATGAATTCATGTGGGCTTTCCTCAAAATCTGCCCTTTCAGTTTCCCACAAGCTCCACCTCCAACAAAATAAGCTC
CAAAAACCACAATCTGTGCTCCTCTTCTTGAAATCTCACGGATTTGACGATTCCCACATCGCCCAATTAATTGAAAAGCGACCCAAAATCCTCCA
TAGCGGAGTAGACGACACTTTGAAGCCCAAATTCGACTTCTTCGTTAAAAATGGTTTCACGGGTAAGCTTCTGCCTCAGCTTATCGCCTCCGATC
CGAATATTTTGTCAGCGGCTGTAGATTCTCATCTTAAACCATGTTTTGAGCTTCTCAAGTTATTTCTGGGCAGTCCTGACAGAATTGTAGTTGCCTT
AAAGCGTGCGCCTTTTTTAATGTCTTTTAGTTTCAAGGGTGCTGTGCAGCCAAATATTGAGTTGTTGATAAAAGAGGGAATGCATGTTGATAGGG
TTGCAAAACTACTAAGTTTGCATGCAAGAGTGATACTGGTTAAGCATGATAGGATGGTTTATGCAGTGAATGCTTTGAAGAATTTGGGTGTTGAA
CCCAAGACTCCTGTGTTTCTACATGCTGCTAAGGTAATGTTATCCATAAGTAAGTCGAATTGGAGGAAGAAAATTGAGGTTATGAAGAGCCTTGG
ATGGAGTGAAGAGGAGATTATTGTGGCTTTTAAACGATACCCTTATCTATTAGCATGCTCAGAGGAGAAAATCAGAAAATCGTTGGATTTCTTTGT
GAATACTTTAAAGTTAGAACCACAAGCTATAATTACATGTCCCGAGTATTTGTCATATTCGGTTGATAGAAGGCTTCGTCCCAGGCATAATGTTTTG
AAGGTTTTGGTGTCAAAGAAGCTGGTTAAAGAAGATGAGAAGATTGTGCGGGCAGTAACCATTAGTGATAGAGATTTCCTTGAAAAGTATGTTA
CTAAATATGCAGACAAGGTGACAGGTTTATTGGAAATATATGGTGGTATCAGCAAGGCAAAAAAG

Ptr004

TCATTTAAGGTTCAATACCTCATCGATTCATGCGGGCTTCCTTCACAGTTGGCCCTTTCAACTTACCAGAAGCTCCAACACGACAAGAAGAACCT
CCCAAATGCTTATTCTGTGCTTCAGTACTTGAAAGATCATGACTTTAGCAACACCCACATTTCCAAACTAATCGACAAGTATCCCCGAGTCCTTCA
AGTCAGAGTGGGAAGCAATCTAAAACCCAAATTCGACTTCTTCACTGAAAATGGGTTTGTGGGTCAGCTTCTGCCTCAGCTTATTCTATCAAATC
CGTCTGTTTTGAGAAGAGCTTTAGATTCTCAAATAAAACCATGTTTTGAGTTATTGAACTCACTTCTTGGTTGTAAGGAGAACCTTGTAGTGGCTC
TTAAGCGTGCTTCTTGGCTGTTGACGGTGAATCTTAAGGTTGTTATTCAACCAAATGTTGATTTGTTGATTAAAGAGGGATTGCCTCTTGATAGGG
TTGCAAAGCTAATTCTCTGGCAACCAAGAGCTGTACTACAAAAGATGGACAGGATGGTTTATGCATTACATGCTCTTAAGAGTATGGGCCTTGAT
GTAGAGGATAATATCTTTATACATGCTCTTCGCGTGAGGATACAGTTGCCTGAAACGACTTGGAAGAAGAAAATTGAAGGGATGAAGAGTTCGC
AGTGGAGTGAAGAGGAGATTTTGGGGGCTTTTAAGCGCTACCCCCCAATACTAGCATTGTCAGAGAAGAAAATCAGGAGTTCAATGGATTTCTT
TATCAATACAATGGAGCTGGAGAGACAAAACATAATTGCCTGTCCCCTTTTTCTTGGCTATTCAATTGATAAAAGGGTTCGTCCAAGGTATAATGT
TATAAAGGTTTTGAAGTCGAAGAAGCTGACAAGTAGAGACAAGAAGATGACTACTTTGCTAACAATAAACGAGAAGAATTTCTTGACAAATTAT
GTCCACAGGTATGTAGATGTAGTCCCTGGTTTATTGGAGCTGTACATGGGTAATGGTAAGACAAAAAAG

Ptr005

TCATTTACAGTTAAGTTCCTCGTCAATTCATGCGGGCTTCCTTTAAAATCCGCTCTTTCAGTTTCTAAAAAGTTCCAAATCCATGAAAAGGAACTC
CATAAGTCCCTTTCTGTACTTGAGTTCTTGAAAGCTCACGACTTTAACGAGACCCAGATAGGCAGATTGATCGAAAAGTGGCCCCGTGTCCTCCT

CTGCAGAGTAGAAAGCACCCTCAAGCTCAAGTTCGATTTCCTCACTCAGAACGGTTTTTCGGGTCAGATTCTGCCTCAGCTTATCGTCTTAGTTC
CGGCGATTTTGAATAGGAAGGTAGATTCTTGTATTAAGCCATGCTTTGAGTTTTTGAAGTCCTTTCTTGACAACAATGAGAAACTTTTAGCAGCTA
TTAAGCGTTATCCGTGGTATTTTACCTTTAATTTCAATTCTGCTCTGAAACCAAATACTGTTTTCTTGATAAAAGAGGGAGTGCCTCATGATAGGGT
TGCGAAACTGATTTTGATGTATCCGAGAACTTTACAGATGAAACCTGATAGGATGGTCCGTGTAGTGAATTCTGTTAAGAATCTGGGCCTTGAAC
CAAAGGCTCCTGTGTTTGTACATGCTCTTAGAGTGATGATAGGAATGAGTGAATCCACTTGGAAGAGGAAAATTGAGTATATGAAGAGTTTGGGG
TGGACTGAAGATGAGGTTTTGCTGACCTTTAAACGAAATCCAGATATACTGGCATGCTCGGAGGACAAAATCGGGAGGGCCATGGATTTCTTTGT
GAATACTGTGAGATTAGGATCACAAACTGTAGTTGCAAATCCAGTTTTACTCCAGTACTCAATTGATAAAAGGGTTCGTCCGAGGTATAATGTTTT
GAAGGTTTTGGAGTCAAAGAATCTGATCGAAGTGAACCAGAGGGTTTTTTGGTTGCTAACAAGAAGTGAGATGAAATTCCGGGAAAATTATGTT
GCAAGGTATGCAGATAAAGTACCAGGCTTATTGGAGATATATCGCGGTACTGTGGAAGCAAAAAAG

Ptr006

TCATTTACAGTCGACTTCCTCATCAACTCATGTGGGCTTCCCTCAAAATCTGCTCTTTCTGTCTCCCAGAAGTTACAACTCGACGAGAAGAGCAT
CCAGAAGCCCCAATCAGTACTCGAGTTTCTGAAAGCTCACGGCTTTAAAGAAACCCACGTCGTCAAATTAATTGAGAAGCGGCCTGATGTCCTC
AGACGTGGAGTAGACACCAATCTCAAGCCAAAATTTGAGTTCCTCATCGCAAATGGCTTTGTGGGTAAGCTTCTTCCTGAGCTTATTACATCAAA
TCCGAATGTTTTGGAAAGGGCTTTAGAATCTAATATGAAGCCATGTTTTGAGTATTTTAAGTCCATTCTTGGTAGTAATGACATGATTGTAGCGGCT
TCTAAGCGATGTGCAGTGTTCTTGACCTATGATTGGAAGAGTATTATACAGCCAAATGTTGAGTTGTTGATAAAAGAGGGAGTGCCTGAAGAGAG
AGTAGTAAAAATGATTGTTGCACAACCAAGAATTATATATCAAAGGCGCGATAGGATGGTTTATGCAGTCAATGCTGTCAAGAATTTGGGTCTTGA
GCCAAAGGCTCCCATGTTTATATATGCTCTTAGATCGATCTTATCCATGAATGAGTTCACTTGGAAGAAGAAGATTGAGGTGATGAAAAGTTTTGG
GTGGACTGAAGAAGAGATTTTGCGGGCATTTAAGCAATACCCGTTTCAATTATCATCCTCGGAGGAAAAAATGAGGAAATCGATGGATTTCTTGT
TGAATACTATTAAGATGGAAAGGCAGGCCATCATTGCCTGTCCTAAGTTTCTTATGTATTCAACCGAGAAAAGGCTCCGTCCTAGGTATGATGTTT
TGAAGATTTTGAAGTCTAAGAAGCTAATTGAAATCGGCAAGAAGACGAACTATCTGCTAACAGTTAGTGAGAAGAATTTCTTGGAGAATTATGTT
ACTAAGTATGCTGATAAAGTGCCAGGTTTATTGGAGGTATACAGGGGCACAACAAAGACCGAAAGA

Ptr007

TCATTTACGGTGGATTTTCTCGTCAACTCATGTGGGCTTCCTTTAAAATCTGCACTTTTAGCTTCCCGGAAGCTGAAGCTGGACAAAAAAAATCT
TGGAAATCCTCCTTTTGTGCTACAATTCTTGAAATCTCACAACTTCGAAGAAACCCACATCTCCAAATTGATTGAGAGGCGGCCTCAAGTCCTCC
AGTCCAGAGTAGAAGGCAATCTCGCTCCCAGATTCAAGTTTCTGATCGCAAATGGCTTTGTGGGTAAGCTCCTTCACGACCTTATCATACACCAT
ACTGAAATACTTACAAGTGCCTTAGATTCTCGTATTAAACCAGCTTTTTACCTTTTGAAGTCGTTTCTATACTGTAATGAGAATATTGTTGCGGCTC
TTAAGCGTTCTTCCAGGTTGTTGACGGCTGATTTGAATGTTAATGCGCAACCTAACATTGATTTTTTGAGAAAAGAGGGAGTTCCTGTTAACATG
GTAGCAAAATTGATTATCTTAAATCCAGGAACTATACTGAGTAAGCGTGGCAGGATGGTTTATGCGATGAATGCTATTAAGAATTTGGGTCTGGAG
CCAGATAAGACGATGTTTGTACGTGCTCTTAGCGTGAGGTTACAAATGACTGAAACAACCTGGAATAAGAAAATTGAAGTGATGAAGAGTTTGC
AGTGGAGTGAAGAGGAGATTCTGAGGGCTTTTAAGAGGTACCCACAAATATTAGCATTCTCGGAGGAGAAAATCAGGAGTGCAATAGATTTCTA
TATCAATACTATGGAGTTGGAAAGACAAATTATAATTGCTAATCCTAATTTTATTGGCTTCTCAATTGATAAAAGGATTCGCCCAAGGTATAATGTTA
TAAATGTCTTAGAGTCGAAGGAACTGATTAAAGGAGACATGAAAATTTCTACTCTGCTAGCTACGAGTGAGAAGAAGTTCTTTATAAATTATGTC
AGCAGGTTTGCGGATGAAGTCCCTGGTCTATTAGAGCTGTACAAGGGTACAGCCATGAGAACAGAG

Ptr008

TCCTTTACAGTTCATTTCCTTGTGAATTCATGCGGGCTCACTTCAAAATCTGCTCTTTCAGTTTCCAAGAAGTTCCAAATCCGTGAAAACAACCTC
CAAAACCCTCAATCCGTACTCCAGTTCTTGAAAGCTCATGACTTTAGCGAAACCCATATTTCCAAATTGATTGAAAAGCGACCCAAGATTCTCCT
GCGCAGAATTGAAGACAATCTGAAAGCCAAGTTTGACTTCTTTATTGAGAATGGTTTTGCGGGTCAGCTTCTGCCCCAACTTATTTTATCAAATCC
GGTGATTTTGGAAAGGGCCTTAGATTCCCATATTAAACCATCTCTTCTGTACTTCAAGTCTATTCTCGGTACCAGCGAGAAAGTTATTGCAGCTTC
CAAACGTTCTGTGTTTTTGTTGACTTGTGATTGGAACAGTATCGTGCTACCAAATGTTGATTTCTTGATTAAAGAGGGAGTGCCTGTTGATAGGGT
TGCAAAGTTGTTTCTGTTTCACCCACAGGTTGTGCAACGGAAGCATGATAGGATGGTTTATGCAGTGAATACTGTCAAGGATTTGGGACTTGAAC
CAGAGGTTTCTATTTTTATATATGCTCTTACTACCATGATGCAATCGAGCGAGTCCACTTTGAAGAAGAAAGTTGAAGTGTTGAAGAGTTTGGGG
TGGACGGAGGAGGAGATTTTCCGGGCTTTTAAGCAAGACCCTGCTATTTTGCGATTTTCAGAGGAAAAGATCAGGGGTGTGATGGATTTCTTGG
TGAATACTGTGGGATTGAGGCCACAAACTATTATTGCAAACCCTTTGTTTCTTCATTACTCAATTAACAAAAGGCTTCGACCAAGGTATAATGTTT
TGAAGGCTTTGGAGTCGAAGAAGCTCTTTGATGAGGGCATGAGCATTGGGTCGGCGCTAAAAATGAGCGAGAAGAAATTCATGAAGAATTATGT
TTCCAAGTATGTACACAGTGTCCCCGGTATATTGGACACATATAAGGGTATCATCAAGCCTATAAAG

Ptr009

TCATTTACTGTACAACACCTCCTTAGCTCATCTGGGCTTCACTTAGAATCTGTTCATTCAGTTTCCCAGAAGCTCCAAATCGATGAAAGTGATCTT

CAAAACCCCCATTATGTCATCGGGTTCTTGAAAGCTCACGATTTTAAGGATGCCCACATAGCCAAATTGATCCACAAGTGGCCTGCTGTCCTCCAT
TGCAAAGTAGAACACAATTTGAAGCCTAAATTTGAGTTCTTCATAGAAAATGGTTTTGTTGGTGAGATTCTGCCCGAGCTTATTGTATCAAATCCG
GATGTTTTGAGAAGGGCTTTAGATTCTCGAATTATTCCATGCTTTGAGCTTTTGAAGTCGGTTCTTGGTTGCAGTGAGAAAGCTGCATCGGCTTTT
AAGCGTTGT------------TCGGTATCAATGATGTCTGCCATGGAACCAAACATTGATTTATTGATAAAAGAGGGAGTGCCTGTCGATAGGATTGCAAA
ACTGATTATGTTGCAACCAAGAACTATACAGCAAAAGCATCAGAGGATGGTTTATGCTGTGAAAGCTCTTAAAGATTTGGAAATCGATTCAAAAA
CTACCGTTTTTATACATGCTCTTAGAGTGATGTTACAAATGAGCGAGTCCACTTGGAACAAGAAAGTTGAAGTGTTGAAGAGTTTGGGGTGGAC
CGAGGAGGAGATTTTACAGGCTTTTAAGCGATGCCCCTTTTGTTTTACATGTTCGGAGGAGAAAATCAGGAGTGTCGTGGATTTCCTGGTGAACA
CATTGAAGATGGAACTGCGAACTGTAATTGGACGTCCCGAGTTTCTTATGCTTTCTGTTGATAAAAGGATTCGTCCAAGGTACAATGTGTTAAAG
ATTTTGGAGTCGAAGAAGCTGGTTATTGGGAAGAAGAATATGAAGCAGTTGCTAACAATGAGGGAGAACAATTTCTTCCAGAACTATGTCATTA
AGTATGCAGACAAGGTCCCAGGTTTATTGGAGGCATATGAGGTTCCAAAGCAA---------

Ptr010

----------------------------------------------------------------------------------------------------AACTCCAAATTTGTACTTGAGTTCTTGAAAGCTCACAACTTCAGC
GACACCCTGATCACCCAACTGATCCAGAATCATTCCCGAATCCTCCAAAGCAGAGTAGAAAGCAATATCAAGCCGAAGTTCGACTTCTTCGTTA
AACATGGGTTAGCGGGCCAACTTCTGCCTGAGCTTATT------------------------------------------------------------------------------------------------------------------CG
TTCTCCATGGCTGTTGACGTATAATGTGAAAGGTATTATGCAACCAAATATTGATTTGTTGATAAAAGAGGGGGTAACTTTTGATAGGGTAGCAAA
ACTGATTATCTCACAACCAGGAGCCATACAGCAAAAGCATTCTAGGATGGTCTATACTGTGAATGCTCTTAAAAACTTGGGCATTGAACCAAATA
CTCCCATGTTTATGCATGCTCTTAGAGTGATGTTACAAACGAGCGACCCCACTCGGAAGAAGAAAGTTGGAGTGTTGAAGAGTTTGGGTTGGAC
TGAGGAGGAAATTTTAAAGGATTTTAAGCATGATCCCCTTATTTTGGGATGTTCAGAGGAGAAGATCAGGGATGTCATGGATTTTTTTGCGGGTAC
CTTGAGATTGAAACCTCAAACTGTTATTACGAACTCCTGGTTCCTTCATTACTCAATTGATAAAAGGCTCCGTCCAAGGTACAATGTTTTGAAGAC
TTTGAAGTCAAAGAACCCGATTGATGGGGACATCAGGATTGCGTGGCTGTTAACATTGAGTGAGAAGAAATTCTTGGAAAACTTTGTTACTAAG
TATGCAGACAATGTCCCAGGTTTATTAGATTTTCTGCAATGTGGT---------------

Ptr011

TCTTCTACGGCCCAATTCCTCGTCAATTCATGCGGTCTTTCTCTGCAATCTGCTCTTTCAGTTTCTAAAAAGTTCCAAATCCATGAGAATGATCTCC
ATAAGCTCCGATCTGTTGTCCAATTCATGAAAGCTCATGGTTTTAGCGAGACCCACCTGGCCAAATTGATCCAAAGTCGGCCAGGAGTTCTCCAT
TGCAGATTAGAAGGTAATCTCAAACCCAAATTTGAATTTCTCACAGAGAATGGCGTTGTGGGTCAGCTTCTGCCAGAGCTCATTTTATCAAATAC
TCATATTTTAAGAAGGGGTTTAGATTCTCAAATTAAACCATGTTTTGAGTTTTTAAAGTCTGTTCTTGGTTGCAATGAGAACGTTTTGGTGGCTCTT
AAACATTCTTCTTGGCTGTTGACGTTGCGTTTGAAGGGCACTATGCAACCAAATTATGATTTGTTGATAGAAGAGGGAGTGGCTGTTGATAGGAT
TGCTAAACTGATTATGATGGACCCAAGAGCCATACAGAATAAGCGTGATAAAATGATTTCTACAGTGAGTACTCTCAAGGATTTAGGCCTTGAAC
CAAATGCTCCTGTTTTTATACACGCTCTTAGAGCGATGTTATCTATCAGTGAATCGACTCGGAAGAAGAAAATTGAGGTGCTGAAGAGTTTGGGT
TGGAGTGAGAAAGATATTTGGCATGCCTTTAAGCGACATCCTCTTCTTTTGGGATATTCAGAGGAGAAAATCAGGGCTGGGATGGATTTTTTTGT
GAATACATTGAAGTTGGGATTGCAACATGTGATTGCCTGCCCCCAGCTTCTTACTTATTCTATTGATAAAAGGTTGCTTCCCAGGTATAATGTTTTA
AAGATTTTGGAGTCAAAGAAGCTGATTAAAAAGGATTTGAAGATTAAGAATTTGCTTAAAATAAATGAGAAGGAATTCTTGAAGAATTATGTTTA
CAAGTATGTGGACGATATTCCAGGGTTGTTGGATTTATATGGAGGAGCCGACAAAGCAAAAAAG

Ptr012

TCTTTTACTGTTCAATATCTCATCAACTCATGCGGCCTTCCTCTACAATCTGCTCTTTCCGTCTCCAAAAAGTTCCAAATCGATGAAAATAACCTTC
AGAAGCCTCAATCCGTCATCCAATTCTTGAAATCTTACGATTTTCAAGATTCCCACATTGCCAAATTGATCGAAAAGTGGCCCGCTGTCCTCCGTT
CCAGAACCGAAGATAATCTCAAGCCTAAGTTTGACTTCTTCATAAAAAATGGCTTTGTGGGTCAGCTTCTGCCTCAGCTTGCCGTATTAGATCCG
GTAATTTTTAGAACGTCCTTGGATGCTTCTATCAAGCCATGTTTTGAGCTTTTGAAACGATTTCTTGAAAGTAATGAGAATATTTTAGCTGCTCTTA
GTCGTGCTCCTTTTTTAATGTCCTTTAGTTTCAATGCTACCGTGCGACCAAACCTTGACTTGTTGAAAAAGGAGGGAGTGACCGCTGATAGGGTT
GCAAAACTGCTATTGTCGCAGCCAAGATCTTTACAACATAGTAATGATAGGATGGTTTATGCTGTCACTTATCTTAAGCAACTGGGCATCGAGCCA
GACAAGACAATGTATATACATGCTCTTACAGTGATTGCACGAATGAGTGAATCTGCTTGGAGGAAGAAAATTGATATGTTTAAGAGTGTTGGGTG
GACCGAAGAGGAGGTTTTATGGGCTTTTAAGCGATTCCCTTATATATTATTAACTTCAGAGGAGAAAATCCGGAGCATGATGGATTTCTTTCTGAA
TAAGATGAAGTTGGAACGGCAAACTATAGTTGCCAATCCTGCGCTTCTTAAATACTCTTTTGGCAATAGGATTCTTCCGAGGTGCAATGTTTTGGA
GGTTTTGAAGTCAAAGAAGCTGATTAAAGGAGACCCAAACATTGCTACTTTATTAAAATTAAGCGAGAAGGATTTCATGGAGAGATGTGTTACC
AAGTATGAGGATAAAGTTCCTGGTTTACTGGAGATGTATGGAGGAATTGACAAAGGAAAAAGA

Ptr013

TCTTTTACTGTTCAATACCTCGTCAACTCATGCGGCCTTCCTCTACAATCTGCTCTTTCCGTCTCCAAAAAGTTCCAAATCGATGAGAATAACCTT

CAGAAGCCTCAATCCGTCATCCAGTTCTTGAAATCTTACGATTTTCAGGATGCCCACGTTGCCAAATTGATCGAAAAGTGGCCTGCTGTCCTCCG

TTCCAGAACAGAAGGTAATCTCAAGCCTAAGTTTGACTTCTTCATAAAAAATGGCTTTGTGGGTCAGCTTCTGCCTCAGCTTGCCGTATTGGATC

CGAGAATTTTTAGAGCCTCCTTACATGCTCATATCAAGCCATGTTTTGAGCTTTTGAAACGTTTTCTTGAAAGTAATGAGAATATTTTAGCTGCTCT

TAGTCGTGCTCCTTTTTTAATGTCCTTTAGTTTCAATGCTACCGTGCGACCAAACCTTGACTTGTTGAAAAAGGAGGGAGTGACCGCTGATAGGG

TTGCAAAACTGCTGTTGTCGCAGCCAAGATCTTTACAACATAGTAATAATAGGATGGTTTATGCTGTCACTTATCTTAAGCAACTGGGCATCGAAC

CAGACAAGACAATGTATATACATGCTCTTACAGTGATTGCACGAATGAATGAATCTGCTTGGAGGAAGAAAATTGATATGTTTAAGAGTGTTGGG

TGGACCGAAGAGGAGGTTTTATGGGCTTTTAAGCGATTCCCTTGTCTATTATTAAGTTCAGAGGAGAAAATCCGGAGCATGATGGATTTCTTTCTG

AATAAGATGAAGTTGGAACGGCAAACTATAGTTGCCAATCCTGCGCTTCTTAAATACTCTTTTGGCAATAGGATTCTTCCGAGGTGCAATGTTTTG

GAGGTTTTGAAGTCAAAGAAGCTGATTAAAGGAGACACAAACATTGCTACTTTTCTAAAATTAAGCGAGAAGGATTTCATGGAGAGATGTGTTG

CCAAGTATGAGAATAAAGTTCCTGGTTTACTGGAGATGTATGGAGGAATTGACAAAGGAAAAAGA

Ptr014

TCATTTACTGTTCAGTACCTCATTACCTCATGTGGGCTTTCCTTGCAGTCAGCTTGTTCTGTTTCCAAAAAGTTCCAAATCGATGAACAGAATCTC

CAAAAGCCCCTATCCGTTATCCAACTTTTGAAATCTCACGATTTTAAGGATGCCCACATTGCCAAAATGATCGAAAAGCGACCCCGATTGCTTCA

CTGCAGTACACAAGACAATTTGAAGCCCAAATTCGACTTCTTCATCAAGAATGGATTCGTGGGTCGACTTCTCCCTGAGTTATTAGTTTCAGATC

CGGTAATTTTGACAAGGAACTTGGGTTCTCGTATTAAGCCATGTTTTAAGCTCTTGAAGTCATATGTACAGAGTCGTGAAGGCGTTGTAGCTCTTC

TTAAGCGTGCTCCGTTTTTTTTATCGTATGGTTCCATGGACTCAATGCGACTAAATATTGATTTGTTGGTAAAAGAGGGTGTGGCTGCTGATAGGAT

TGCAAAACTGTTGATCTGGCAACCAAGAAGTATACTCTATAAGCCTGATAGGATAGTTTATGCGCTAAATGCTCTTAAGAACCTGGGACTTCAGC

CAGGGGATAAACCATTTATACAGGCTCTTAGCGTGAGGATACAATCGAATGACACGGCTTGGAAGAAGAAAATTGAAGTTATCAAGAGTTTGGG

CTGGAGTGAAGAGGAGGTTTTGAGGAGTTTTAAACGACACCCGCCATTATTTGGATACTCAGAGAAGAAAATCAGGACTGCAATGGATTTCTTT

ATCAACACTATGGAGTTGGAACGACAATTTATAATAAACAGTCCCAATTTTCTTGGGATGTCAATTGATAAAAGGATTCGTCCAAGGTATAACGTT

ATCAAGGTTTTGGAGTCAAAGGAACTGATTAAAAGAGATAAGAAGATTAGTACTTTGTTGTCATTAAGTGAGAAAAATTTCTGGGCGAATTATGT

CATCAAGTATGCGGACGAGGTCCCAGGTTTACTAGAGATATATGGTGGTGCTGGCAAGGCAAAA---

Ptr015

---TTTACTGTTCAATTCCTTGTCAACTCTTGTGGGCTTCCTCTACAATCTGCCCTCTCAGTTTCCAAAAAGTTCCAAATCGACGAAAATAACCTCC

ATAAGCCCCAATCTGTCATCCAATTCTTGAAATCTAACGACTTTAAGGACACCCACATCGCTAAAACGATCGAGAAGTGGCCTGCTGTCCTACAT

TCCAGAACAGAAGATACTCTCAAGCCTAAGTTTGACTTCTTCATAAAAAATGGTTTTGCGGGTCAGCTTCTGCCCCAGCTTATTGTATCAAACCC

GGACGTTTTGAGAAGACATCTGGGTTCTCATATTAAGCCATTTTTTGAGTTTTTGAAGCCTTTTTATGCCAGTAACGAGGAAGTTGTAGAAGCTAT

TATGCGTGCTCCATGGTTGTTGTCGATTCCTTTAAATGGTGATATGCAGTTAAATACCGATTTCCTGATAAAAGAGGGAGTGTCCATCGATAGAATA

GCAAAACTGATGCAGTGGCAACCAAGGGTTATGGGGCAGAAGCATGATAAGATGGTTTATGCAGTGGCTGCTACTAAAAAATTGGGCGTTCAAC

CAGGGGATAGTATGTTTGTACGTGTTCTTGCTGTGCTGGTTATAGTGTCTGAATCAACCTGGAGGAAGAGAATTGAGGTGATGAAAAGTATGGGT

TGGAGTGAAGGTGAGGTTTTGTGTGCCTTTAAGCGATTCCCCCCGCTATTAACTTGTTCAGAGGAGAAAATCAGGGGCGCAATGGATTTTTTCTT

TAATACCATGGAATTGGGACGGCAATCTTTAATTACCTATCCCTATTTTATTGGTTTCTCAATTGATAAAAGAGTTCGTCCAAGGTACAATGTTATG

AAGGTTTTGGAGTCGAGAAAGCTAATTGAAGGAGATTGGAATATTGCTACTCCGCTAACAATCAGCGAGAAGAAGTTCTTGCTGAATTATGTTAC

CAAATATGCGGACAAAGCTCCAGACTTATTGCAGATATATGGCGGTACTGACAAGTCAAAAAGA

**Ptr_2_CDS.phylipi**

4 1020

Ptr016

TCATTTGCAGCCTCTTATCTCATAAAAAAAATGTGGGTTCTCTCCAGAATCTGCTCTTTCAGCTTCCAAGCATCTCAAATTCGAAACCCCAGAT------

AAACCTGACTCAGTCATCGACACATTCAGGCGCTATGGTTTCCCCGAAGACAAAATCTTTAAACTCGTTAAGAAATTCCCGAAGGTGCTTTCCTG

TAATCCTGACAAAACCCTTTTGCCTAGACTGGATTTTTTTCTCTCCAGAGGCATGTCAAGCACTGAACTTGCCACCCTTTTCTGCATAATTCCTCC

TCTTTTGCATAGAAGCTTGGAAAACATTATAACTCCTACTTTTAACTTCCTTAGTGATTTGCTTCAATCCAATGACAAGGCAATTACTGTTGCCAA

AACCTATCCGTTTATTATCTATCATCGTCCTGAGAGCTATTTGCAACCTTACGTCAGCATTTTGAGAGAAAATGGAATTCCCAAATCACATATTGCT

TCCTTAATTTATAAGTGGCCTAGGACTGTGAGAGCGTGTCCAATTCGTTTTAGAAATACTGTTGAAACGGTAAAAGAAATGGGTTTTGACCCTTC

CAAGTTAGTGTTTACTCTAGCAGTGCTGGCACGGTCTGCGCAGAGTAAATCTGGATGGGAAAAGAAGGTTGGTGTTTATAAGAGATGGGGTTGG

TCAGACGAAGAGGTTCTGGCTGCTTTCAAAAGGAATCCGTGGTGTATGATGAGTTCCGAGGATAAGATTATGGCAGTGATGGACTTTTTGGTTAA

CAATATGGGCTGTGAGTCTTCTTATGTTGCCGAACATCCAATTCTTTTGTTGCTGAGCTTGGAGAAGAGACTTATACCAAGGGCTTCTGTCCTTCA

ATTCCTACAATCAAATAAGTTGATTGATGAAAAGCCTAACTTGGCCACTTTGTTTAAGTATTCGGAGAAGTCATTTCTTCATAAGTTCGTGGATGG

TTTT---GATGAAGCTCCTCAGCTATTAAAACTATACAGGGAAAAACTGAATCTTTCAAAA

Ptr017

TCATTCCCAGTCTCTTACCTCATAAATAAATGTGGGTTCTCTCTAGGATCTGCTCTATCAGCTTCTAAACGTCTCACTTTTGAAATCCCAGAT------C

AACCTGACGCAGTGATTCACATGTTCAAGCGCTATGGTTTTTGTGATGCCGATATCTTTAGACTAATTAAAAGATATCCACGAGTCCTTTCGTGCA

ATCCTGACAAAACCCTCTTGCCAAGACTCGAGTTTTTTATTCTAAAGGCATGTCAAACACTGATATTGCCCACATTTTATGCGGATGCCCTCTTC

TTTTAGGCAGAAGCTTGCAAAATCATGTAACTCCTAATTTCAATTTACTTAGTGATCTGCTTCTATCCGACAACAAAGTCATTTCTGCTGCCAGAA

ACGACCCGTTTATTCTATTTCGTCACGGTGACAGATATTTGAAACCTTTTATCAACCTTTTGCAAGATCATGGAGTTCCCAGAACACAGATTGCGT

CCTTAATTTGCAACTGGCCTAGGTCTATCGGTGTATGCCTAAATCATTTCAGAAAGTGCGTCGAGGAAGTTAGAGAAAAGGGTTATGATCCTTCTA

GTGCAGAGTTTATTAGAGCAGTGGTGGTATTGAGTCAATTGGGAAAATCTGGGTGGGAAAGAAAGGCTGTTGTTTATAAGAGTTGGGGTTGGTC

TGAGAAAGACATTCTTGTTGCTTTCAGAAAGAACCCGTGGAGCTTGATGACATCGGAGAGTAAGATTATGGCAGTGATGAAATTTTTCGTTGAG

AAAATGGATTGTGAGTCTTTATATCTTGCAAAACACCCAAATCTTTTGATGTATAGCTTGGAGAAGAGACTCATACCAAGGGCTTTGGTTCTTCAT

TTTCTGTTAAGGAATAAGTTGATTGACAAGAAGCCTAACTTGAATACTTTGTTTGTGTATTCTGAGAAGTTGTTCCTTGAGAAGTTCGTGAATTGT

TTT---GATGAAGCCCCCCAGCTATTAAAGCTATATAGAGATCAATCAGATCTTGCAAAA

Ptr018

TCATTTGCAGCCTCTTATCTCATAAGCAAATTTGGGTTCTCTCCTGAATCTGCTCTTTCAGCTTCTAAACATCTCAATTTTACAACCACAGAG------A

AACCTGACTCTGTGATTCACATTTTCAAGCACTATGGTTTCTCTCAGGTTCAAACCTTAAAACTCGTTAAAAAATACCCACGAGTGCTTTCGTGC

AACCCTGAGAAAACCCTTTTGCCTAAACTCGAGTTTTTCCACTCTAAAGGCATGTCAAACAATGACATTGCACGCATTTTATGTACATATCCTCAT

ATTTTAGTTAGAAGCTTGGAAAACTGTATAACTCTTAATTTTAACTTCCTTGGCAATTTGCTTCAATCTAATGACAAGACAATTGCTGCTGCTAAA

AGATACTCGCCTATTCTATATCATAAACCCGATAGATTCTTGAAACCTTGTATCGACATTTTGGAAGAGTATGGAGTCCCCAAAAAGCATATTGCTT

CATTGGTTCACCGTTGGCCTAGGTCTGTCATGATGTCTCCGAATTATCTGAGAAGAATTGTAGAGAAGGTGAGAGAAATGGGATGTGATCCTCTT

AAGCCACAGTTTACTACGGCAGTGATGGTGATGAGTCTATTGAGCGAATCTGGGTGGGAAAGGAGGCTGGGTGTTTATAAGAGTTGGGGTTGGT

CTGAGGAAGACGTTCATGCTGCTTTCATAAAGGAGCCTTGGTGCATGATGACATCAGATGATAAGATTATGGCAGTGATGGATTTTTTGGTCAAC

AACATGGATTGTGAGCCTTCATTTATTGTGAAGAACCCCTATCTTTTGAAGCCAGGCTTGAAGACAACATTTATACCAAGAGCTTCAGTTGTTCA

CTTTTTGTTATCGAAACAGTTGATTGAAACGAAGCCAAACTTGGTTACTCTGTTTTTGTGTTCTGAGAAGATGTTCCTTGAGAAATTCGTGTATCG

TTTT---GAAGAAGCCCCTCAGCTATTAAAGCTGTATGGAGATCAATCAAATCTTTCAAAA

Ptr019

TCATTTGCAGCCTCTTATCTCATAAACAAATTTGTGTTCTCTCCTGAATCTGCTCTTTCAGCTTCTAAACATCTCAGTTTTAAAACCCCAGAT------A

ACCCTGACTCAGTGATTCGCATGTTCCAGCACTATGGTCTCTCTCAAGACCAAATCTTTAAACTCGTTAAAAAATACCCACGAGTGCTTTCGTGC

AAACCTGAGAAAACCCTTTTGCCGAAACTCAAGTTTTTTCACTCTAAAGGCATGTCAGGCAATGACATTGCCCACATTTTATGTGCACATCCTTG

TATTTTGAATCGAAGCTTGGAAAACCAAATAATTCTTAATTTCAACTTCCTTGGCAATTTGCTTCAATCTAATGAAAAGACAATTGCTGCTGTTAA

AAGATACTCGCCTATTCTCTATCATAAAATCGATACATATTTGAAACCTTGTATCGACATTTTGGAAGAGTATGGAGTCCCCAAAAGGCATATTGCT

ACATTGGTTCACCGTTCTCCTAGGTCTGTCATGATGTCTCCGAATCATTTGAGAAGCATCGCTGAGACGGTGAGAGAAATGGGATGTGATCCTCT

TAAGCCGCATTTTGCTACGGCAGTGATGGTGATGGGTCTATTGAGCAAATCTGGGTGGGAAAGGAGGCTGGGTGTTTATAAGAGTTGGGGTTGG

TCCGAGGAAGACGTTCTTGCTGCTTTCATAAAGGAGCCTTGGTGCATGATGACATCGGATGATAAGATTATGGCAGTGATGGATTTTTTGGTTAAT

AACATGGATTGTGAGCCTTCATTTATTGTGAAAAACCCCTATCTTTTGAAGCCAGGCTTGAAGACAACATTTATACCAAGAGCTTCAGTTGCTCA

GTTTTTGTTATCGAAACAGTTGATTAAAAGGAAGCCTAACTTGGTTACTCTGTTTTTGTGTTCTGAGAAGTTGTTCCTTGAGAAGTTCGTGAATTG

TTTT---GATGAAGCCCCACAGCTATTGAAGCTGTATAACAAAAGAAATCAAATCTTTCAA

**Ptr_3_CDS.phylipi**

8 1020

Ptr020

TCCTTTACAGTCTCATACCTTATGAACAAATGTGGGTTCTCTCTAAAATCTGCTTTAGAAGTTTCTAAGCAGGTCCATTTTGAAACCCCAGAT------

AAGCCTGATTCTGTTCTTGCCGTTTTCAAGAACTGTGGCTTCTCAAAGTCCCATATCTTGAACCTTGTGAGGAGACGGCCAGCGGTGCTTTTGTC

TAAACCTAACACAACACTTTTGCCCAAGCTTGAGTTTTTCCAATCTAAAGGTTTTTCAAGCCCTGATGGTATCAAAATCATATCATCCTACCCGTG

GGTTTTTAAGTACAGCTTAGAAAACCAGTTAGTTCCTGCTTTTGATTTCCTTGAAAACTCGCTCCAATCTGATGCCGTGGCCATCAAAGCAATCA

AGCGTTTCCCTCGTATTCTAAATGTTACTGTTGAA---AACATGGCACGTGTTGTCGATGTTTTACTAGACAATGGAGTCCCTGAAAAGAATATTGC

TCTGCTAATTCGTTCCCGGCCTTCTATTATGGTTTCAAATCTGGAGAATTTAAAAAAGCTTATAGAGGAAGTGACTCTAATGGGATTTCATCCTTCC
AAGAGTCAGTTTGTTGTGGCAATCAGGGTGCTGACGTCCGTGACCAGAACCACGTGGGAGAAGAAGCTTGATGTGCATAGAAAGTGGGGGTTG
TCTGAGGAAGAAATTCTTGAAGCATTTGTAAAGTTTCCATGGTTTATGTCCCTATCTGAAGAGAAGATCATGGCAGTTATGGATCTTTTTGTCAAC
AACTTGGGCTGGGAGTCTTCCTATATTGCCAAAAATCCAACTTTTTCATCATATAGCCTTGAGAAAAGGCTAATTCCAAGGGCTTTGGTATTGCAA
TTTTTAGTTTCTAAAGGCTTGGTTGAGAAGAGTTTCAGAAGCCTTGCATTCTTCAATACACCTGAAGATAAGTTCCGGCAGATGTTTATAGATCAC
CATGCTGACTCTACC---CAGATACTGAAATTTTACGAGGAAAAACTGAATCTTTCATCA

Ptr021

TCCTTTACAGTCTCATACCTTATGAACAAATGTGGGTTTTCTCTAAAATCTGCTTTAGAAGTTTCTAAACAAGTTCATTTTGAAACCCCAGAT------
AAGCCTGATACTGTTCTTGCCGTTTTCAAGAACTATGGCTTCTCAAAATCCCATATTCTGAACCTTGTGACGAGACGGCCAACGGTGCTTTTGTCT
AAACCTAACACAACACTTTTGCCCAAGCTTGAGTTTTTCCAATCTAAAGGTTTTTCAAGCCCTGACCATGTCAAAATCATATCATCCTACCCAAG
GATTTTGATGTGTAGCTTAGAAAACCAGTTAGTCCCTGCTTTTGATTTCCTTGAAAACTTGCTCCAATCTGATGCCTCGGTCATCAAAGCAATCAA
GCGTTACCCTGGTATTCTATATATTAATGTTGAA---AGCATGGCACGTGTTGTCGATGTTTTACGAGACAATGGAGTCCCTAAAAAGAATATTGCTC
TGCTAATCCGTTCCAAGCCTTCTATTATGATTTCAAATCTGGAGAATTTTAAAAACCTTATACAGAAAGTGGCTCTAATGGGATTTCGCCCTTCAA
AGAGTCAATTTGTTTGTGCAATCATGGTGCTGATGTCCTTGAGCAGATCCACTTGGGAAAAGAAGTTTGCTGTGTATAGGAGGTGGGGTTTGTCT
GAAGAAGAAATTCTTACAGCATTTGTAAAATTTCCAATGTTTATGCGCATATCTGCAGAGAAGATCGCTGGATCAATGGATCTTTTTGTCAACAAA
TTGGGTTGGGAGTCTTCCTATATTGCCAAAAATCCAACTTTTTCATCATATAGCCTTGAGCAAAGGCTAATTCCAAGGGCTTTGGTGTTGCAATTT
TTAGTTTCTAAAGGCTTGGTTGAGAAGAGTTTCAGAAGCCTTGCATTCTTCAATACACCTGAAGATAAGTTCCGGCAGATGTTTATTGATCACCAT
GCTGAATCTACC---CAGATATTGAGATTTTATGAGGAAAAACTTAATCTTTCATCA

Ptr022

TCCTTTACAGTCTCATACCTTATGAACAAATGTGGGTTCTCTCTAAAATCTGCTTTAGAAGTTTCTAAACAGGTTCATTTTGAAACCCCAGAT------
AAGCCTGATTCTGTTCTTGCCGTTTTCAAGAACTATGGTTTCTCAAAATCCCAAATTTTGAACCTTGTGAGGAGACGGCCACCGGTGCTTTTGTC
TAGACCTAACACAACACTTTTGCCCAAGCTTGAGTTTTTCCAATCTAAAGGTTTTTCCAGCCATGATGTTATCAAAATCATATCATCCTACCCATGG
GTTTTGATGTACAGCTTAGAAAACAAATTAGTCCCTGCTTTTGATTTCCTTGAAAACTTGCTCCAATCTGATGCCGTGGTCATCATAGTAATCATGC
GCTCCCCTCGTATTCTAAATAGTAATGTTGAA---AACATGGCACGTATTGTTGATGTTTTACAAGGCAATGGAGTCCCTAAAAAGAATATTGCTCTG
CCAATTCGTTGCCAGCCTTCTATTATGATTTCAAATCTGGAGAATTTTAAAAAGCTTATAGAGGAAGTGACTCTAATGGGATTTCATCCTTCCAAG
AGTCAATTTGTTTCTGCAATCACAGTGCTGAGGTCCATGAGCGGATCCACTTGGGAAAAGAAGCTTACTGTGTATAGGAGATGGGGTTTGTCTGA
AGAAGAAATTCTTACAGCATTTGTAAAATTTCCAATGTTTATGCGCAAATCTGCAGAGAAGATCGCAGCATCAATGGATCTTTTTGTCAACAAATT
AGGGTGGGAGTCGTCCTATCTTGCCAAAAATCCAACTTGCTCATCATATAGCCTTGAGAAAAGGCTAATTCCAAGGGCTTTGGTGTTGCAATTTTT
AGTTTCTAAACGCTTGGTTGAGAAGAGTTTCAGAAGCCTTGCATTCTTCAATACACCTGAAGATAAGTTCCGGCGGATATTTATAGACCAGCATG
CTGAGTCTACC---CAGATATTGAAATTTTACAGGGAAAAACTGAATCTTTCATCA

Ptr023

TCCTTTACAGTCTCATACCTTATGAACAAATGTGGGTTCTCTCTAAAATCTGCTTTAGAAGTTTCTAAACAGGTCCATTTTGAAACCCCATAT------A
AGCCTGATTCTGTTCTTGCCGTTTTCAAGAACTATGGCTTCTCAAAGTCCCATATCTTGAACCTTGTGAGGAGACGGCCAGCGGTGCTTTTGTCT
AAACCTAAAACAACACTTTTGCCCAAGCTTGAGTTTTTCAAATCTAAAGGTTTTTCGAGCCCAGATGTTATCAAAATCATATCATCCTACCCATGG
GTTTTTAAGTACAGCTTAGAAAACCAGTTAGTTCCTGCTTTTGATTTCCTTGAAAACTCGCTTCAATCTGATGCCGTGGCCATCAAAGCAATCAA
GCGTTTCCCTCGTATTCTAAATGTTACTGTTGAA---AACATGGCACGTGTTGTTGATGTTTTACGAGACAGTGGAGTCCCTGAAAAGAATATTGCT
CTGCTAATTCGTTCCCGGCCTTCTATTATGGTTACAAATCTGGAGAATTTTAAAAAGCTTATAAAGGAAGTGACCCTAATGGGATTTCATCCTTGC
AAGAGTCAGTTTATTGAAGCAATCAGGGTGCTGACGTCCATGAGCAGATCCACTTGGGAGAAGAAGCTTGATGTGCATAGAAAGTGGGGGTTGT
CCGAGGAAGAAATTCTTGAAGCATTTGTAAAGTGTCCATGGTTTATGTCCCTATCTGAAGAGAAGATCATGGCAGTTATGGATCTTTTTGTCAACA
AACTGGGCTGGGAGTCTGCCTATATTGCCAAAAATCCAACCTTTTCATCATATAGCCTTGAGAAAAGGCTAATTCCAAGGGCTTTGGTATTGCAAT
TTTTAGTTTCTAAAGGCTTGGTTGAGAAGAGTTTCAGAAGCCTTGCGTTCTTCAATACACCCGAAGACAAGTTCCGGCATATGTTTATAGATCGC
CATGGTGAATCTACC---CGGATATTGAAATTTTACGAGGAAAAACTGAATCTTTCGGCA

Ptr024

GCATTTACTGTGTCTTATCTCATAAATTCTTGTGGGTTGTCTCCAAAATCTGCATTAGCAGCTTCCAAAGATGTCCATTTTGATGACCCACAT------A
AACCTGATGTTGTCCTTAGCTTTTTCAAGAACCATGGCTTTTCAAAAGCCCAAATCTTTAACATTATCAAAGGATACCCTGGAGTGCTCTTGACCA
ATCCTGACAAAACCCTTTTGCCCAAACTTGAGTTTTTGCAATCTAAAGGTGTTTCAAGCCCTGACATTGCCAAGATCATATCATCACATCCTTGGA
CTTTGCAGCGGAGGTAC------TGCTTTGTCCCTATTTTTTATTTCTTTAAACACTTGGTTCAATCTGATGATACAACCATCAAAGTATTCAAGCGCTA

TCCCGGTCTATTTGGACTTGATCTTGCA---ATTGTGACATCTATGCTCAATATTTTGCGAGATAATGGAGTGCCTGAAAGTAATATTCCTATGTTAGC
TCGTTGCTATCCCTTAACTATGATGTTGACCCTGGAGAAATTTCAGAAGCTTGTGGAGGAATTGAGGGCAATGGGCTTTGATACTTCCACGAGTC
GGTTTATTTTGGCGATGAATGTTCTGTGCTTGATGAGTAGAGTCAAGTGGGAAAGGAAACTTGATGCGTACAGAGACTGGGGTTTGTCTCATGA
AGAGATTCTTGCAGCATTCAGAAAGTATCCATATTTTATGACAGCATCTGAATATAAGATTATGGAAGTGATGTGTCTTTTTGTCAACAAATTGGGC
TGGGAGCCTTCTTTTATTGCTAAACACCCATCTCTTATGTTATACAGTGTGGAGAAGACGCTCATTCCAAGGGCTTCAGTTTTGGAGTTTCTAGTT
TCCAGAGGCTTGATTGAGAAGAGTTTTAGAAGCTATGAATTCTTCCAGAGTCCAGAAAATAAGTTCCTGCAAAATGTCATAAGCAGCTATGCTGA
ATCCACT---GAGCTGTTGCAGTTGTACAGGGAGAAACAGAATCTTTCAAGA

Ptr025

TCATTTACTGTTTCTTACCTCGTCAACAAATGTGGGTTTTCACTAAAATCTGCATTAGAAGCTTCTAAACGTGTCCATTATGAAACCCCACAT------A
AACCTGATTCTTTACTTAGCTTCTTCAAGGACCATGGCTTCTCAGAGAACCAAACTTTAAAACTTACCAGAAAATGCCCTGAACTGCTTTTGTAC
AATCCTGACAAAACCCTTTTGCCCAAACTTGAGTTTTTGTATTCTAAAGGAGTTTCAACCGCTGATGTTGCTAAAATCATATCCTTCTACCCTTGG
ATTTTGAGATGCAGCTTAGAAAACCAGTTAGTCCCTACTTTTGATTTTTTGAAAAATTGGTTC---CCAAATGATACCATTGTTCAGGTATTCAAAAG
TACTCCTCTTGTTCTTCAACTTAATCCTGTA---ACTGTGAAATATATTTCCCAGATTTTGCGAGATAATGGAGTGCCTGACAAGAATATTGTTATGCT
AGTTCGTAGCCACCCTAAAACATTGTTGTTGAGTCCAAAGAAATTTAACAAGGTGCTGTGTAAAGTGAGGAAAATGGGATTAGATCCTTGCAAG
ACTCAATTTGTCGTTGCGATCCTAGCATTGACGTCCATGAGCAGATCCACATGGGAAAAGAAACTTGATGTATATAGGAGATGGGGTTTGTCCCA
TGAAGAGATTCTTGCAGCATTTGCAAAGTCTCCATGGTTTATGACCCTATCTGAAGAGAAGGTTGTGGCAGTGATGGATCTTTTTGTCAACAAAT
TGGGCTGGGAATCTTCTTTTATTGCCAAAAACCCAACACTTGTCTCGTATAGCTTGGAAAAGAGGCTCACTCCAAGGGCTTCGGTTTTGCAATTT
CTAGTGTCTCAAGGTTTGATTGAGAAGAGTTTCAGAAGCACTACATTCTTCATTGCATCTGAAAATAAGTTCCTGCAGCAGTTCATAAATCAGCG
AGCTGAATCTACC---CAGATATTGAAACTGTACCAGGAGAAATTGAATCTTTCAAGA

Ptr026

TCCTTTACAGTATCATACCTTATGAACATTTGTGGGTTCTCTCTAAAACCTGCTTTAGAAGTTTCTAAACAGGTTCATTTTGAAACCCCAGGT------
AATGCTGATTCTGTTCTTGAGATTTTCAAGAACCATGGCTTCTCAAAGGCCCATATTTTGAACCTTGTGAGGAGATGGCCTAGAGTGCTTTTGTGT
AAACCTCACAGAACACTCTTGCCCAAGCTTGGATTTTTCCATTCTAAAGGGTTTTCAAGCCCTGATGTTGTCAAAATTATATCAACCTATCCGTGG
ATTTTGAGAATTAGCTTTGAAAACAAGTTAGTCCCTGCTTTTGATTTCTTTGAAAATTTGCTCCAATCTGATGCCATGGCTATCAAAGCAGTCAAA
CTTGACCCGCGCCTTCTAGATGCTGGTCTTGAA---AAGGCCGCACGTATTGTTGATATTTTGTTAGAAAATGGAGTCCCTATGAAGAATATTGCTCT
ATCAGTTCGGATAAAGCCTGGTATTATGCTTTCGAATCTGGAGAATTTCAAAAGGCTTGTTCAGAAAGCAAGTCTAATGGGATTTCATCCTTCCAA
GAGTCAATTTGTTGTAGCAATTGTGTTGCTGAGGTCCATGACCACATCCACATGGGAAAAAAAGCTTGATGTGTATAGGAGATGGGGTTTGTCTC
AAGAAGAAATTCTTGCAGCATTTGTAAAGAATCCTTGGTTTATGAGCTTATCTGAAGAGAAGATCACGGCAGTGATGGATCTTTTTGTCAACCAA
TTGGGTTGGGAGTCTTCCTATCTTGCCAAAAACCCAACTATACCATCATATAGCCTGGACAAAAGGCTTGTTCCAAGGGCTTTGTTATTGCAATTT
CTAGTTTCTAAAGGCTTGGTTGAGAAGAGTTTCAGAAGCACTGCGTTCTTCTACACACCTGAAAATAAGTTCCGGCAGATGTTCATAAATCACCG
TTCTGAATCTACC---CAGATATTGAAATTTTACAATGAAAAACTGAATCTTTCATCA

Ptr027

TCCTTTACAGTCTCATACCTTATGAACAAATGTGGGTTCTCTCTAAAATCTGCTTTAGAAGTTTCTAAGCAGGTCCATTTTGAAACCCCAGAT------
AAGCCTGATTCTGTTCTTGCCGTTTTCAAGAACTGTGGCTTCTCAAAGTCCCATATCTTGAACCTTGTGAGGAGACGGCCAGCGGTGCTTTTGTC
TAAACCTAACACAACACTTTTGCCCAAGCTTGAGTTTTTCCAATCTAAAGGTTTTTCAAGCCCTGATGGTATCAAAATCATATCATCCTACCCGTG
GGTTTTTAAGTACAGCTTAGAAAACCAGTTAGTTCCTGCTTTTGATTTCCTTGAAAACTCGCTCCAATCTGATGCCGTGGCCATCAAAGCAATCA
AGCGTTTCCCTCGTATTCTAAATGTTACTGTTGAA---AACATGGCACGTGTTGTCGATGTTTTACTAGACAATGGAGTCCCTGAAAAGAATATTGC
TCTGCTAATTCGTTCCCGGCCTTCTATTATGGTTTCAAATCTGGAGAATTTAAAAAAGCTTATAGAGGAAGTGACTCTAATGGGATTTCATCCTTCC
AAGAGTCAGTTTGTTGTGGCAATCAGGGTGCTGACGTCCGTGACCAGAACCACGTGGGAGAAGAAGCTTGATGTGCATAGAAAGTGGGGGTTG
TCTGAGGAAGAAATTCTTGAAGCATTTGTAAAGTTTCCATGGTTTATGTCCCTATCTGAAGAGAAGATCATGGCAGTTATGGATCTTTTTGTCAAC
AACTTGGGCTGGGAGTCTTCCTATATTGCCAAAAATCCAACTTTTTCATCATATAGCCTTGAGAAAAGGCTAATTCCAAGGGCTTTGGTATTGCAA
TTTTTAGTTTCTAAAGGCTTGGTTGAGAAGAGTTTCAGAAGCCTTGCATTCTTCAATACACCTGAAGATAAGTTCCGGCAGATGTTTATAGATCAC
CATGCTGACTCTACC---CAGATACTGAAATTTTACGAGGAAAAACTGAATCTTTCATCA

**Ath_CDS.phylipi**

13 1242

AT1G56380

GCTGATGATGTGATCCCTAGAGATGCTCAAAAAGAGCCGAAATTAACGGTCTCTTACCTTGTTGATTCAGTGGGTATACCGATAAAATTCGCAGA
ATCAATCTTGAAGGAAGTCAGCTCAAAGGACAAGTGCAATCCCAATTCTGTTCTGAATCTTCTAAGAAGTTATGATTTCACAGATTCTCAGATATC
AAGCATCATTACGACTGATCCAGAATTACTTATGGAAGATGCTGAGAATTCCCTTTGTCCCAAGCTTAAGTTTTTGGAGTCCAGAGAAATTTTAAG
CTCTAGGCTCAATGACATTGTTACTAGAGTTCCGAAAATCTTGAGGATGGAAGAGGAAAAATCTATGATCACATACTATGACTTTGTCAAAACCA
TTACACTAACGAGTTCGCGTTCGGACTTCTATAAGGTTTGCGAACTTTATCCATAT--------------------------------------------------------------------------------
----------------------------------------ATTGAAAGTTCTATTAGGAAGGTTATTGAGATGGGTTTTGATCCTTTCGCCCCAAAAATTTTCGATGCTACGGTCGT
TGTTTGCACATTAAGCAACGAAACATTAGAAGAAAGAGTCAATATCTATAAAACGTTAGGCTTTGATGTCAGAGATGTATGGGAAATGTTCAAGA
AGTGTCCTACTTTTCTGAATATCTCGGAGAAGAAGATAACTCAGAGTTTCGAAACATTGAAGAAGTGTGGACTTGTCGAAGAAGAAGTCATCTC
TATGTTCCAGAAGTCTCCACAATGTATTGATTTTTCAGAGCTGGATATAACTCAGAATTTTGAATTCTTGAAGGGATGTGGATTAGTCGAAGAAGA
GGTCCTCTCTATGTTCAAGAGGTATCCGCAATGTATTGGTTTTTCAGAGAAAAAAATATTGAACGCGGTTGAAACATTTTTAGGCCAAATGATGGT
CAATCGAGAAGGCGTGGTTTCGATCCCTGTGGTACTTGAATTCAGCATGGAGAAAATGATTGTACCGAGGTGCAATGTTATCAAAGCTCTTACGT
CCAAAAGATTGCTCAAAACTGAAGTCTCTTCAATGTTTTCTGTCTTGATATGTCCCGATGAGGTGTTTCTTGAAAGGTATGTGAGTAAGCATGATG
ATCAAGAACTTGTCGATGAGTTGATGTCTATCTTC

AT1G61960

GTTGCTGATGCGACCCTTATAGATAGCCTTAAAGGTAACAACTTTACAGTCTCTTACCTAGTTGATTCATTGGGTTTAACCACAAAGCTCGCAGAA
TCGATCTCCAAGAAAGTAAGCTTCGAGGAGAGGAGAAATCCGGATTCTGTTCTGAGTCTTCTTACAAGTTATGGGTTCACAAAGTCTCAGATTTC
AAGTATCATTACTATCTACCCACGATTGCTTGCATTAGATGCTGAGAAATCCATTGCTCCCAAGCTTCAGTCTCTGCAGTCCAGAGGAGCTTCAAG
CTCTGAGCTCACTCAGATTGTTTCTACTGTTCCTAAAATCTTGGGGAAGAGAGGGCACAAATCTATAACTGTATACTATGATTTCGTCAAAGACAT
TATAGAAGCTGATAAGAGTTCTAGTTACGAAAAGTTATGTCATTCTTTTCCACAGGGTAAC---AAGAAGAATAAGATCAGAAATATCTCTGTTTTG
AGAGAATTGGGTGTGGCTCAGCGTTTGTTATTCCCTTTGCTCATCTCAGATGGCCAACCTGTCTGTGGTAAAGAAAGATTTGAAGAATCTCTCAA
GAAGGTTGTTGAGATGGGTTTTGATCCGGAGACTACAAAGTTCGTTGAAGCTCTGCGAGTTATTTACCGTATGAGTGACAAAACAATAGAAGAA
AAAGTCAATGTCTACAAAAGGTTAGGCTTTGGTGTGGCTGATGTATGGGCAATCTTCAAGAAGTGGCCTTCTTTTTTGTCATACTCAGAGAAGAA
GATAACTCATACTTTTGAAACCCTTAAGAGTTGTGGTCTGCTTAAACACGAGGTCCTGTTATTGTTGAAGAAGCATCCAAAATGCATTTGTTCTTC
AGAGCAGAAGATTGTGAACTCCATAGAAACATTCTTAGGCCTAGGATTCAGCAGAGATGAGTTTGCGATGATGGTCAAGCGATATCCTCAGTGCA
TTGACTATACTGCTGAGACAGTGAAGAAGAAAACTGAGTTTATTGTGAAGAATATGAATTGGCCACTAGAGGCTTTGGTTTCGATCCCTCAGGTA
TTTGGATACAGCCTGGAGAAGAGAACTGTCCCAAGGTGTAATGTAATCAAAACTCTCATCTCAAAAGGACTGATGGGAAGTGAAGCCCCTCCAA
TGTCATCTGTTTTAACAAGTACCGATCAGGCGTTTTTAAGGAGGTATGTGATGAAGCACGAC------AAGTTGGCACCTGAGTTGATGGCTATCTTC

AT1G61970

GCTGCGTGTTTGAGTCCTAGAGTTGGTCGAAAAGGAAACAACTTTACAGTCTCTTACCTTGTTGATTCATTGGGCTTAACCACAAAACTCGCTGA
ATCCATTTCACGGAAGGTTAGCTTCGAGGACAAGAACAATCCTGATTCTGTTCTGAATCTTTTGACAAGTCATGGGTTCACAGGTTCTCAGATCT
CCACCATCATTAGGGATTACCCACAGTTGCTTATAGCAGATGCTGAGAAATCACTTGGTCCCAAGCTTCAGTTTTTGCAGTCTAGAGGAGCTTCA
AGCTCTGAAATCACTGAGATTGTTTCTTCAGTTCCGGAAATATTGGGAAAGAAAGGGCACAAAACTATCAGCGTATACTATGATTTCATCAAAGA
CACTTTG---CTTGAGAAGAGTTCCAAGAACGAAAAGTTATGTCATTCTTTGCCACAGGGTAAT---CTGGAGAACAAGATCAGAAATGTATCGGTTT
TAAGAGAATTGGGAATGCCTCACAAGTTGTTATTCTCCTTGCTCATCTCCGATAGCCAACCTGTATGTGGAAAAGAAAAATTTGAAGAAACCCTC
AAGAAAGTTGTTGAGATGGGTTTTGATCCTACCACCTCAAAGTTTGTCGAAGCTCTACAAGTTATTTACAAAATGAACGAGAAAACAATCGAAG
AAAAAGTCCATCTCTATAAAAGCTTAGGCTTTGATGTGGGAGATGTATGGTCAAGTTTT------------------------------------------------------------------------
------------------------------AAAAAGTGGCCAATCTCGCTGAGAGTATCGGAGAAGAAGATGTTGGACTCCATTGAAACATTTCTTGGCCTAGGATTCA
GCAGAGATGAGTTTGCCAAGATGGTCAAGCACTTTCCTCCGTGCATTGGATTATCTACAGAGACGGTCAAAAAGAAAACTGAGTTTCTGGTGAA
GAAGATGAATTGGCCACTAAAGGCTGTTGTTTCAAACCCTGCGGTATTTGGATATAGCCTGGAGAAGAGGATTGTTCCAAGGGGTAACGTAATCA
AAGCTCTCATGTCCAAAGGATTGATGAGAAATGAACTCCCTTCGATATCGTGTGTCCTCATGTGTACCAAGCAGGTGTTTTTAAACAGGTATGTG
GCGAATCACGTGGACAAGCAGCTGGTGACGGAGTTGATGGCTATCTAT

AT1G61980

GCTGCGCGTTTGAGTCCTAGAGTTGGTCGAAAAGGTAAGAGCTTTACAGTCTCTTACCTCGTTGATTCATTGGGTTTACCCAAAAAGCTCGCTGA
ATCGGTCTCAAGGAAGGTTAGCTTCGAGGACAAGGACAACCCTGACTCTGTTCTGAATCTTTTGAGAAGTCATGGGTTCACTGATTCTCAGATCT

CCACCATCGTTACGGATTATCCACAATTGCTTGTAGCTGATGCTGAGAAATCACTTGCTCCCAAGCTTCAGTTTTTGCAGTCTAGAGGAGCTTCA
AGCTCTGAGCTCACTGAGATTGTTTCTACAGTTCCTAAAATCTTGGGAAAGAGAGGCCACAAAACTATCAGCGTATTCTATGATTTCATCAAAGA
AACTTTA---CTTGATAAGAGTTCCAAGTCCGAAAAGTCATGTCAACCTTTTCCACAGGGTAAT---CTGGAGAACAAGATCAGAAATTTATCTGTTT
TGAGAGAATTGGGGATGCCTCACAAGTTGTTATTCCCCTTGCTCATCTCTTGTGACGTACCTGTCTTCGGTAAAGAAAAATTTGAAGAATCCCTC
AAGAAAGTTGTTGAGATGGGTTTTGATCCATCCACCTCAAAGTTTGTGGAAGCTTTGTGCGTTGTTCAAAGATTGAGCGACAAAAACATAGAAG
ATAAAGTCAATGCGTATAAAAGGTTAGGCTTTGATGTGGAATATGTATGGACAGTGTTC--------------------------------------------------------------------------------
---------------------------------AAGAGGTGGCCAAACTTTCTGACACACTCGGAGAAGAAGATATTGAACACCATTGAAACATTTCTAGGCCTAGGGTTC
AGCAGAGATGAGTTTAGCGTGTTAATTAAGCGCTTTCCTCAGGGCATTGGATTATCCGCGGAGATGGTTAAAAAGAAGACTGAGTTTTTGGTAAA
GAAGATGAATTGGCCACTAAAGGCTCTTGTTTCAAACCCTGCGGTGCTTGGATACAGCCTAGAGAAGAGGACTGTCCCAAGGGGTAACGTAGTT
CAAGCTCTCATCTCCAAGGGATTGATCGGAAGTGAACTCCCTTCGATATCGCGTGTCTTCGTATGTACCGATCAGGTGTTCTTAAACAGGTATGTG
AAGAGGCATGAGGACAAGCAGCTGGAGACTGAGTTGATGGCTATCTAT

AT1G61990

GCTGCTGATGTGAGCATTAGAGATGGTCGAAAAGGGAAGAACTTTACAGTCTCTTACCTCGTTGATTCATTGGGTTTATCTAAAAAGCTCGCTGA
ATCCATCTCAAGGAAAGTTAGCTTTGAGGACAAGGTCAATCCTGATTCTGTTCTAAGTCTGTTTAGAAGTTACGGGTTCACAGATTCTCAGATCT
CTACCATCATTACGGATTATCCGTTATTGCTTGTAGCAGATGCTAAGAAAGCACTTGGTCGCAAGCTTCAGATTCTGCAGTCTAGAGGAGCTTCAA
GCTCTGAGATCACTGAGATTGTTTCTACAGTACCGAGAATCTTAGGAAAG---------AAATCTATTACTGTATACTATGATGCTGTCAAAGACATTATA
GTAGCCGATACGAGTTCGAGTTATGAA---------------CTCCCACAGGGTTCT---CAGGGGAATAAAATCCGAAATGTATCGGCTTTGAGAGAATTAGG
CATGCCTTCTCGGTTGTTATTACCCTTGCTCGTCTCCAAATCGCAGCCTGTGTGTGGAAAAGAAAATTTTGATGCATCCCTTAAGAAGGTTGTTGA
GATGGGGTTTGATCCCACCACCACTAAATTTGTTTTAGCTTTGCGGATGCTTTACCAAATGAGCGAGAAAACGATTGAAGAAAAAGTTGTAGTCT
TTAGAAGTTTAGGTTTTACTGTGGATGATGTATGGGAAATTTTC-----------------------------------------------------------------------------------------------------------------
AAGAAGACTCCTTCTGTTTTGAAAGTCTCCAAGAAGAAGATATTGAAGTCGGCTGAAACGTTTCTAGACCTAGGATATAGCAGAGCTGAGTTTC
TGATGATGGTTAAGCGTTATCCTCCTTGCATTGAATATTCTGTAGAGTCTGTTAAAAAGAAGAATGAGTTTCTGGTGAAGAAGATGAAATGGCCA
CGAAACGCCTTGGTTTTGCACCCTCAGGTGTTTGGATACAGCATGGAGAAGAGGATTATACCAAGGTGTAACATACTCGAAGCTCTCTTGTCGAA
AGGACTGCTCGGAAGTGAACTACCTGCAGTGTCCTCTGTTTTGTCGTGTACCGATGAAGGATTTTTAGACAGGTATGTGATGAAGCACAAC------
GAGCTGGTGCCTACGTTGATGGCTATATTC

AT1G62010

GCTGCTGATGTGAGCCCT------------AAAGGGGAGACCTTTAAATCCTCGTCCTTCCTTGATTCACTGCGTTTAGCTACAAAACTCACT-----------------
-------------GACAAGGGAAATTCTGATTCCGTTCTGGATCTTCTCAGAAGTTATGGGTTCACTGATTCTCAGATCTCCAGCATCATCAGGAGTGATTC
AAGAGTGCTTATAGACAATGATGCGACATCCCTTGGTTCAAAGCTTCAGTTTTTGCAGTCCAGAGGAGCTTCAAGCTCTGAGCTCACTGAGGTT
GTTTCCACAGTTCCTAAAATCTTGGGGAAGAGAGAGGGCAAATCTCTCAGCAGATATTATGATTTCATCAAAGTCATTATAGAAGCTGATAAGAG
TTCCAAGTATGAAAAGATATCTCATTCTTTGGCACAGGGT------------AACAAGATCAGAAATATCTTGGTTTTGAGAGAACTAGGCGTGCCTCAAA
AGCGGTTACTGCTTTTGCTCATCTCCAAATCGCAGCCTGTTTGTGGAAAAGAAAAGTTTGATGCATCACTCAAGAAGGTAGTTGAGATGGGTTTT
GATCCAACCACCTCAACGTTTGTGCACGCTTTGCATATGCTTTACCAAATGAGCGACAAAACGATTGAAGAAAAAATCCGAGTCTATAGAAGTGT
AGGTTTCTCTGTTGATGATGTATGGGCAATGTTC------------------------------------------------------------------------------------------------------------------AAGAAGTGG
CCACGCTCTTTGACACATTCGGAGAAGAAGGTAGCCAACTCTATAGAAACATTTCTAGGTCTAGGATTCAGCAGAGATGTGTTTATGATGATGTT
CAAGCGGTTTCCTCCGTGTATTGGATATTCTACAGAGGCGGTGAAGAAGAAGACTGAGTTTTTGGTCAAGGAGATGAATTGGCCAGTAAAGGCT
GTGGCTTCAATCCCTCAGGTACTTGGATACAGCCTAGAGAAGAGGACTGTCCCAAGGTGTAATGTTATCAAAGTTCTCATGTCAAAAGGATTGCT
TGAAAGTGAACTCCCTCCAATGTCTTCTGTTTTGACAAGTACTAGTGAGTCGTTTTTAAATTTGTATGTTAGCAAACATGATGACAAGCAGCTTGT
GGCTGAGCTGATGGCAATCTTC

AT1G62085

GCTACAGATGCGAGCCTTAGAGCTGGTCGAAAAGGGTTGAGTTTTTCTGTCTCTTACCTCGTTGATTCATTGGGTTTACCTAAAAAGGTCGCAGA
ATCGATCTCAAAGAAAGTCAGCTTCGAGGACAAAGGCAATCCTGATTCTGTTCTGAGTCTATTGAGAAGTCATGGGTTCACTGATTCTCAGATCT
CAAGCATCATTACGGATTATCCACAATTGCTAGTAGCTGATGCTGAGAAATCAATTGGTCCCAAGCTTCAGTTTCTGCAGTCCAGAGGAGCTTCA
AGGTCTGAGCTCACTCATATTGTATCTACAGTTCCTGAAATCTTGGGGAAGAGAGGGGACAAAACTATCAGCATATACTATGATTTTGTTAAAGAG
ATTATAGAAGCGGATAAGAGTTCCAAGTTCGAAAAGTTATGTCATTCTTTGCCAGAGGGTAGTAAGCAGGAGAACAAAATCAGAAATGTTTTGG
TTTTGAGAGAATTGGGTGTTCCTCAAAGGTTGTTGTTCCCATTGCTTATATCTGATCACCAACCTGTCTGTGGAAAAGAAAATTTTGAAGAATCA
CTCAAGAAAGTTGTTGAGATGGGTTTTGATCCAACCACCTCAAAGTTTGTCAAAGCTCTTAGGGTTGTTTACAGATTCCGCGACAAAACTATAGA

AGCAAAAGTTAATGTCTGTAAAAGTTTAGGCTTTTCTGTTGGAGATGTATGGGCAATGTTTAAGAAGTGTCCTTCCTTTCTGAATTTCTCTGAGAA
CAAGATAGTTCAAACTTGGGAAACCCTGAAGAAGTGTGGACTACTCGAAGACGATGTCCTCTCAGTGTTGAAGAAGTTTCCACAATGCATTAAT
GCTTCAGAGCAGAAGATAATGAACTCCATTGAAACATTTCTAGGCTTAGGATTTAGCAGAGATGAGGTTGCAATGATAGCCAAGCGCTTTCCTCA
ATGCCTTATTTTATCTGCAGAGACAGTGAAGAAAAAGACTGAGTTTCTGGTGAAGAAGATGAATTGGCCACTAAAGGCTGTGGTTTCAACCCCT
GCGGTACTTGGATACAGCTTAGAGAAGAGGACTATCCCAAGGTGTAATGTAATTAAAGCTCTCATGTCCAAAGGATCGCTTGGAAGTGAACTCCC
TGGAATGTCGTCTGTTTTGGTATGTACCAATGAAGAATTCTTATGCAGGTATGTGAAGAACCATGATGACAAGAAGATAGTACCTGAGTTGATGG
CTATTTTC

AT1G62110

GCTACTGATTTGAGCTCAAGAGATGGTCGAAAAGTGAAGAACTTTACTGTCTCTTACCTCGTTGATTCATTGGGTTTAGCTACAAAACTCGCTGA
ATCAATCTCAAAGAAAGTCAGTTTCGTGAACAAGGGCAATCCTGATTTAGTTCTGAGTCTTTTTAGAAGTTATGGGTTCACAAATTCTCAGATCT
CCAGCATCATTACGGATTATCCTAGATTGCTTCTAATAGATGCTGAGAAATCACTTGATATCAAGCTTCAGTTTTTGGAGTCTAGAGGAGCTTCAA
GCCCTGAGCTCACTCAGATTGTTTCTACAGTTCCTAAAATATTGGGAATGAAAGAGGGAAAGTCTCTAGGCAGATACTATGATTTCGTCAAAGAG
ATTATAGAAGCAGATAAGAGTTCCAAGTACGAAACTTTGTGTCAACCTTTGCCAGAGGCTAATAGGCAGGGTAATAAAATCAGAAATGTTTCGGT
TCTTAGAGACTTGGGTGTGCCTCAGAAGTTGTTATTCTCCTTGCTCATCTCTGATGCCCAACCTGTTTGTGGAAAAGAAAATTTTGAAGAATCAC
TCAAGAAGGTTGTTGAGATGGGTTTTGATCCAACCACCTCAAAGTTTGTCCAAGCTCTTAGGGCTGTTTACAGATTCACCGACAAAACAATAGA
AGAAAGAGTTAATGTGTATAAAGGGTTCGGCTTTGCTGTGGAAGATGTATGGGCAATGTTCAAGAAATGTCCTTACTTTTTGAATAGTTCGGAGA
AGAAGATTGGTCAGACAATTGAAACCCTGAAGAAGTGTGGACTACTCGAAGACGAGGTCATCTCAGTGTTGAAAAAGTATCCGCAGTGCATTG
GTACTTCAGAGCAGAAGATATTGAACTCCATTGAAATATTTCTAGGCCTAGGCTTCAGCAGAGATGAGTTCATAACAATGGTTAAGCGCTTTCCTC
AGTGTCTTATTTTATCTGCAGAGACGGTGAAGAAGAAGATTGAGTTTGTGGTGAAGAAGATGAATTGGCCACTGAAGGATGTGGTTTCAAATCC
TACGGTACTTGGATACAACCTGGAGAAGAGGACTGTCCCAAGGTGTAATGTTATTGAAGCTCTCATGTCAAAAGGAGACACAGGAAGTGAACTC
CCTCCAATGTCGTCTGTTTTGGTATGTACTGATGAGTTGTTCTTAAAAAGGTATGTGAGGAACCATGGTGACAAGGAGCTGGTGCTTGAGTTGAT
GACTATTTAC

AT1G62120

GCTGCTGATGTGAGCTCAAGAGATGGTCGAAAAGGTCACAACTTTACAGTTTCTTACCTCGTTGATTCATTAGGTTTAGCTACAAAAGTCGCTGA
ATCGATCTCCATGAAAGTCAGTTTCGATAACAAGGGCAACCCTGATTCTGTTCTGAGTCTTTTGAGAAGTCATGGGTTCACTGATTCTCAGATCTC
CAACATCATTAGGACTTTCCCAAGATTGCTTATCCTAGATGCTGAGAAATCCCTTGCTCCCAAGCTTCAGTTTCTGCAGTCCATAGGTGCTTCAAG
CTCTGAGCTCACCGAGACTGTTTCTGCAGTTCCGAAAATATTGGGGAAGAGAAAGGGCAAATCTCTCAGCAGATACTATGATTTCGTCAAAGTC
ATTATAGAAGCGGATAAGAGTTCCAAGTTAGAAAAGTTATGTCATTCTTTGCCAGAGGGTAGTAAGCAGGAGAACAAAATCAGAAATTTATTGGT
GTTGAGAGAAATGGGTGTGCCTCAGAGGTTGTTATTCTCATTGCTTATATCCGATGCCGGAGATGTTTGTGGTAAAGAAAAATTTAAAGAATCAC
TCAAGAAGGCTGTTGAGATTGGTTTTGATCCAACCACCGCAACTTTTGTCAAAGCTCTTAACGTTCTTTACGGATTAAGCGACAAAGGAATTGA
AAATAAATTCAATGCCTGTAAAAGGTTAGGCTTGGCTGTGGACGATGTATGGGCAATGTTC--------------------------------------------------------------------
------------------------------------AAGAAGTGGCCAAACATTCTGACAAAGTCTGAGAAGAAGATAGAGAACTCCGTTGAAACATTTCTAGGCCTAGGAT
TCAGCAGAGATGAGTTTTTGATGATGGTCAAGCGATTCCCTCAGTGCATTGGATATTCTACAGAGTTGATGAAGACGAAGACTGAGTTTCTGGTG
ACGGAGATGAATTGGCCACTAAAGGCTGTGGCTTCAATCCCTCAGGTACTCGGATACAGCCTAGAGAAGAGGACTGTCCCAAGGTGTAATGTTA
TCAAAGTTCTCATCTCAAAAGGATTGCTCGAAAGTGAACTCCCTCCAATATCCTCTGTTTTGACAAGTACCAGTGAGGTGTTTTTATATATGTATG
TTAGGAAACATGATGACAAGCAGCTTGTGGCTGAGTTGATGGCTATCTTC

AT1G62150

GCTGCTGATGTGAGCTTTAGAGATAGTCGAAAAGGTAACAACTTTACTGTCTCTTACCTCGTTGATTCATTGGGTTTAGCTTCAAAGCTCGCAGA
ATCGATCTCGATGAAAGTCAGTTTCGAGAACAAGGGCAATCCTGATACAGTTCTGAATCTTCTTAGAAGTCATGAGTTCACAGATTCTCAGATCT
CCAGCATCATTTCGGATTATCCAACATTGCTTGTAGCAGATGCTGAAAATTCACTTGGTCCCAAGCTTCTGTTGATGCAGTCTAGAGGAGCTTCAA
GCTCTGAGCTCACTGAGATTGTTTCTAAAGTTCCTAAAATCTTAGGAATGAAAGGGGACAAAAGTATCGGCAGATACTATGATATCGTCAAAGAG
ATTATAGAAGCGGATAAGAGCTCCAAGTTCGAAAAGCTGTGTCATTCTTTGCCTGAGGGTAGTAAGCAGGAGAATAAAATCAGGAATGTATTGGT
TTTGAGAGATTTGGGTGTGCCTCAAAGGTTGTTATTCTCATTGCTCTTCTCTAATCACCATGTCTGCTGTGGAAAAGAAAAGTTTGAAGAATCAC
TCAACAAGGTTGTTGGGATGGGTTTTGATCCCACCACTCCAAAGTTTGTGGAAGCTTTATGCATTGTTTACGGATTGAGCGACAAAAGACTAGA
AGAAAACTTCAATGTCTACAAGAGATTTGGCTTAACTGTAAACGATGTATGGGAACTTTTCAAGAAGTGCCCTGCCTTTCTTGGATACTCTGAGA
ATAGGATAATTCAGACATTTGAAGCCTTAAAGAGGTGTGGTCTGTGCGAAGATGAGGTATTGTCAGTGTTCAAGAAGAATCCACTATGCTTACGT
GCTTCAGAGCAGCAGATATTGAACTCCATGGAAACATTTATAGGCTTAGGATTCAGCAGAGATGAGTTTGTCATGATGGTCAAGTGCCTTCCTCA

GTGCATTGGATATTCTGCAGAGATGGTGAAGAAGAAGACTGAGTTTGTGGTGAAGAAGATGAATTGGCCACTTAAGGTCATTACTTTGTTCCCTC
AGGTACTTGGATATAGCATGGAGAAGAGGACTGTCCCAAGGTGTAATGTAATCAAAGCTCTCATGTCCAAAGGATTGCTTGGAAGCGAACTCCC
TCCAATGGCGTCTGTTTTGGCATGTACCGATCAGACCTTCTTAAAAAGGTATGTGGTTGAGCATGATGAA---AAGCTGGTGCTTGAGTTGATGTCT
ATCTTC

AT1G62490

GCTGCTGATGTGAGCCTTATAGATAGTCAAAAAGGTAAGAACTTTACAGTCTCTTACCTCGTTGATTCATTGGGTTTACCTAAAAAGCTCGCTGAA
TCGATTTCAAAGAAATTTAGATTCGAGGACAAGGCCAATCCTGATTCTGTTCTGAGTCTTTTGAGAAGTCATGGGTTCACAGTTTCTCAGATCTC
AATT------------------------------------------------------------------------------------------------------------------------CCTAAACTCTTGGGAAAAAGAGGGCACAA
AACTCTCAGCTTATACTATGATTTCGTTAAAGAGAGTTTAGAAGCAGATAAGAGTTCTAAGTATGAAACTTTATGTCAGTCTTTTCCACAGGGTAA
T---CTGGAGAACAAGAAGAGAAATGTATCGGTTTTAAGAGAATTGGGAATGCCTCACAAGTTGTTATTCCCCTTGCTTATATCCGTTGGTCAACCT
GTATGTGGAAAAGATAGATTTAATACGTCTCTCAAGAAGGTAGTTGAGATGGGTTTTGATCCGACCACCGCAAAGTTTGTCAAAGCTCTCCACGT
ATCTTACGAAATGAACGACAAAACAATAGAAGAAAAAGTCAATGTCTACAAAATGTTAGGCTTTGCTGTGGAAGATGTATGGGTAATCTTC--------
-----------------------------------------------------------------------------------------------AAGAAGTGGCCATACTCTTTGAAATACTCAGAGGAAAAGATTACTC
AGACGATTGAAACCTTGAAGATGTGCGGTCTACGA---------------------------------------------------------------------------------------------------------------------
----------------------------------------------------------------------------------GGT------CCTTCAAGTTTTGAAGAAGTATCCTCAATTCATACG-----------------------
---TACTTGAGAGCAGAGGAT---------------ATTGAGCTTAAT---------

AT3G46950

GCTAAAGATTCGAGTCCA------------AAGGGAAGTACTTTCACAGTCTCTTATCTCGTTGAATCATTGGGTTTGACTAAAAAACTTGCAGAAACAAT
CTCAAAGAAAGTCACCTTCGAAGACAAGGTAAATCCAGATTCTGTTCTGAATCTTCTTAGAAGTAATGGGTTTAAAGATTCTCAGATATCTAGGA
TCATAAGGGCTTACCCACGATTGCTTGTAACAGATGCTGAGAAATCTCTTCGTCCAAAGCTTCAGTTTTTAAAGTCTAGAGGAGCTTCAAGCTCT
GAGGTCATAGAGATTGTTTCGAATGTTCCAACAATCTTGGATAAGAAAGGTGAAGAATCTGTTAGTTTATACTATGATTTCGTTAAGGACATTATG
CAAGATGGTAAAAGT---------------TTGTGTATTTCTTGTCCAGAGGGAAAG---AAAGGTAACAGAATTCGAAATATATCGGTTTTGAGAGAATTGGG
AGTACCTCAGAAGTTGTTGTTTTCATTGCTTATATCTAGATATCAACCTGTTTGTGGTAAAGAGAAGTTTGAAGAAAGTTTAAAGAAAGTTGTTGA
TATGGGTTTTGATCCGGCTAAATCCAAGTTTGTGGAAGCTTTGCATGTAGTTTACGAAATGAGCGAGAAAACGATAGAAGAGAAAGTGAATGTG
TATAAGAGGTTAGGCTTTAGTGAAGCCGAGATATGGGCGATTTTCAAGAAGTGGCCTTACTTTTTAAAATTCTCTGAGAAGAAGATAATTCTGATG
TTTGAAACACTGAAGAAGTGTGGTTTGGTAGAGGAAGAGATTATTTCGGTGTTGAAGAGTCGTCCACAATGCATACGATCTTCTGAGCAGAAGA
TATTAGACTCTATTGAAATGTTCTTAGGCTTAGGATTCAGTAGAGATGATTTCAAGATGATGGTTAAGCGATATCCTTGTTGCACTGCATATTCTGG
CGAGACGCTGAGGAAGAAGTTTGAGGTTTTGGTGAAGATGATGAATTGGCCGCTAGAGGCTGTGGTCATGATCCCTACGGTACTTGGATACAGC
TTGGAGAAGAGGATAGTGCCAAGGTCTAACGTGATCAAAGCTCTCATGTCGAAAGGATTGATCGGAAGTGAAAACCCTCCGATTTCATCTGTCT
TGGTATGTACTGATCAGGAGTTCTTGAAAAGGTATGTGATGAAGCATGAC------AAGCTAGTGCCTAAGTTGATGGCTATCTTC

AT5G23930

ACTAAAGATCTAAGTCTTGAAGATGAACGAAAGAGGAAAACTTTCACAGTCTCTTACCTTATTGATTCATTAGGGTTAACTACAAAACTCGCTGA
ATCAATCTCAATGAAAGCTAACTTCGATGAAAAAGGTAATCCAGATTCAGTGCTTAAACTCTTAAGAAGCTATGGATTCAAAGATTCTCAAATCT
CAAGTATTATTTCCACTTACCCACGATTTCTCATTGAGAATCCTGAGAAAACTCTTCGTGCTAAACTTCATTTCTTGAAGCTTAACGGAGCTTCAA
GCTCTGAGCTTACAGAGATTGTTTCAAAAGTTCCTAAGATCTTGGGAAAGAGAGGAGGGAAATGGATTAGTCATTACTATGATTATGTCAAAGAG
ATTTTGCAAGATCAAGATTCT--------------------AGTTCTTCTTCTTCCAAGAGAAAGCAGACGAATCGAAATCGAAATGTATCGGTTTTGAGAAAAT
TAGGAGTACCACAGAGACTGTTATTGAACTTGTTGATCTCTAGAGCTAAACCAGTTTGTGGTAAAGAAAGATTTGAAGAATCGGTTAAGAAGATT
GTTGAGATGGGTTTTGATCCGAAAAGTCCAAAGTTTGTTAATGCTTTGTATGTTTTCTATGAGCTTAGTGATAAGACTATAGAGGAGAAAGTGAAT
GCTTATATAAGGTTAGGACTTAGTGTTAATGAAGTTTGGGCAGTTTTCAAGAAATGGCCTTTCTCTTTGAAATACTCTGAGAAGAATATAATTCAA
AAGTTTGAGACTTTGAAACGTGTTGGTCTAACGAAAGAAGAGGTCTGCTTAGTGGTTAAGAAGTACCCTGAATGTGTAGGGACTTCAGAGGAG
AAGATAGTGAAGTCAGTTAAAACTTTTCTTGAGCTAGGATTTACGAAAGACGAGGTTTTGATGATCATTAAGCGACATCCTCAGTGCATTGGATT
AGCTGCGGATTCGGTGAAGAAGAAGACCGAGTTTCTTGTGAAGACAATGGGTTGGCCTTTAAAGGTTGTAGCTTCGACTCCTATTGTACTTGGTT
TCAGCTTGGAGAAGTTCGTTTTACCGCGGTGTAATGTAATCAAAGCTCTTATGTCGAACGGGCTGATC---GGTGAAATGCCTGCGATATCGTCTGT
TTTGACGAGTCCTAAGCTGAAGTTTTTGAAGCTTTTTGTGGAGAAGCATCAG------GATGTGTTGCCTGAGTTGAACTCTATCTTC

**Aly_CDS.phylipi**

19 1368

Al_315177a

ATGTATTCTCTGATTCTCCATGGAAGAAAGTTGGTCCAGTTTCAGAAATGGCGTCACTTGAGGGTTTCGGTGAATCTTTATCAAGATGGATCTGCT
TTTTCCAATTCCTTCTCCTCCGCTGCTAGTGCTTCTGATGTGAGTCCTAGAGTTGGTCGAAAAGGAAACAACTTTACAGTCTCTTACCTCGTTGAT
TCATTGGGTTTACCTAACAAGCTCGCAGAATCTATTTCAAGGAAAGTCAGCTTCGAGGACAAGGGCAATCCTGATTCTGTTCTGAGTCTTTTGAG
AAGTCATGGGTTCACAGATTCTCAGATCTCAAGCATCATTACGGATTATCCAGTATTGCTTATAGCTGATGCTGACAAATCACTTGGTCCCAAGCT
TCAGTTTTTGCAGTCCAGAGGAGCTTCAAGCTCTGAGCTCACTGAGATTGTTTCTGCAGTTCCGAAAATCTTGGGAAAGAAAGAGGGAAAATCT
ATCAGCGCATACTATGATTTCGTCAAAGTCATTATAGAAGCAGATAAGAGTTCCAATATGGGAAGGATATGTCATTCTTTGCCAGAGGGTAAGCAG
GAGAATAAAATTAGAAATGTTTTGGTTTTGAGAGAATTGGGGGTACCTCAGAGGGTTTTATTCTCATTGCTTCTATCCGATGGCAGACATGTTTGT
GGAAAAGAAAAATTTAAAGAATCACTCAAGAAGGTAGTTAAGATTGGTTTTGATCCAACCACCTCAATGTTTGTCGAAGCTCTCAAGGTTCTTTA
CACATTGAGCGACAAAGGAATTGAAAGTAAATTCAATGCCTTTAAAAGGTTAGGCTTGGCTGTGGGCGAT-------------------------------------------------
----------------------------------------------------------------------------------------------------------TCTGAGAAGAAGATAGAGAACTCCATTGAAACATTTCTAGGTCTAG
GATTCAGCAGAGATGAGTTTTTGATGATGGTCAAGCGGTTTCCTCAGTGCATTGGATATTCG---------------------ACAGAGTATTTGGTGAAGGAGAT
GAATTGGCCACTTAAGGCTGTGGCTTCAATCCCTCAGGTACTTGGATACAGCCTAGAGAAGAGGACTGTCCCAAGGTGTAATGTAATCAAAGTT
CTCATCTCAAAAGGATTGTTCGGAAGTGAACTCCCTCCAATATCATCTGTTTTGACAAGTCCTAGGGAGAAGTTTTTAAATTGTTATGTTAGGAAA
CATGATGACAAGCAGCTTGTTGCTGAGTTGTTGGTTATCTTCACT

Al_315177b

ATGTATACTCTGATACTCCATGGAAAAAGGTTGGTCCAGTTGCAGAAATGGCGTCATTTGAGAGTTTCAGTGAACCTTCTCGAAAAGCCATCTCC
TTTTCCCAATTCCTTCTCCTAT---GCTACTGCTACAGATGCGAGCCTTAGAGCTGGTCGAAAAAGGTTGAATTTTTCTGTCTCTTACCTCGTTGCTT
CATTGGGTTTAACTAAAGAAGTCGCAGAATCGATCTCAAGGAAAGTCTGTTTGGTTGACAAGGGCAATCCTGATTCTGTTCTGAGTCTTTTGAGA
AGTTATGCGTTCACTGATTCTCAGATCTCCACCATCGTTACGGATTATCCACAATTGCTAATAGCTGATGCTGAGAAATCACTTGCTCCCAAGCTTC
AGTTTCTGCTGTCCAGAGGAGCTTCAAGCTCTGAGCTCGCTGTGATTGTATCTACAGTTCCTAAAATTTTGGGAAAGAAAGGGGACAAAACTAT
CAGCATATACTATGATATCGTCAAAGAGATTATAGAAGCGGATAAGAGTTCCAAGTTCGAAAAATTATGTCATTCTTTTCCACAGGGTAATCTGGA
GAACAAGATCAGAAATGTTTCGGTTCTGAGAGAGTTAGGTGTGCCTCAGAGGGTTTTATTCTCCTTGCTCATATCTGATCACCAACCTGTCTGTG
GAAAAGAAAATTTTGAAGAATCACTCAAGAAAGTTGTTGAGATGGGTTTTGATCCGACCACCTCAAAGTTTGTCGAAGCTCTAAACGTTGTTTA
TAGACTGAGCGACGAAACAATAGAAGAAAAAGTTAGTGTCTGTAAAGGGTTAGGTTTTTCTGTTGGAGATGTATGGGAAATGTTCAAGAAGTGG
CCTTGCTTTCTGAATAACTCGGAGAAGAAGATAAGTCAAACATTTGAAACCCTGAAGAAGTGTGGACTACCCGAAGACGAGGTCCTCTCACTGT
TGAAGAAGTTTCCACAATGCATTAACGCTTCAGAGCAGAAGATTTTGAACACCATTGAAACATTTCAAGACCTAGGATTTAGCAGAGATGAGTT
TGCAATGATAGCCATGCGCTTTCCTCCATGCCTTATTTTATCTGCAGCGACGGTGAAGAAGAAGACTGAGTTTGTGGTAAAGAAGATGAATTGGC
CACTAAAGGCTGTGGTTTCAACCCCTGCGGTACTTGGATACAGCCTCGAGAAGAGGACTGTACCCAGGTGTAATGTAATTAAAGCTCTCATGTCA
AAACGATTGCCCGGAAGTGAACTGCCTCCAATGTCGTCTGTTTTGGTATGTACCAATGAAGAATTCTTAAACAGGTATGTGAGGAACCATGAAG
ACAAGGAGCTGGTGCCTGAGTTGATGGCTATTTTCACC

Al_315190

ATGTATTCTCTGATACACCATGGAAGAAGGTTGGTCCAGTTGGAGAAATTGCCTAATTTCAGAGTTTCAGTG---------CAAACCGCATCTTCTTTGT
CAAATTCGTTCTCTTTT---ACTAGTGTTGCTGATGCGAGCCTTAGAGATGGTCTTAAAGGGAACAACTTTACCATCTCTTACCTCGTTGATTCATTG
GGTTTAACCACAAAACTCGCAGAATCGATCTCAAAGAAAGTCAGCCTCGAGGACAAGGAAAATCCGGATTCTGTTGTGAGTCTTCTTACAAGTT
ATGGCTTCACAAAGTCTCAGATTTCAAGTATCATTACGATTTACCCACGATTGCTTATATTACATGCTGACAAATCC---------------------------------AGAG
GAGCTTCAAGCTCTGAGCTTACTGAGATTGTTTCTACAGTTCCTAAGATCTTGGGAAAGAGAGGGCACAAATCTATATCTGTATACTATGATTTCG
TCAAAGACATCATAGAAGCTGATAAGAGTTCTAGTTACGAAAAACTATGCCATTCTTTTCCACAGGGTAACAAGGAAAACAAGATCAGAAATATC
TCTGTTTTGAGAGAATTGGGTGTGGCTCAGCGTTTGTTATTCCCCTTGCTCATCTCTGATAGCCAACCTGTTTGTGGGAAAGAAAGATTTGAAGA
ATCACTCAAGAAGGTTGTTGAGATGGGTTTTGATCCGGAGACCTCAAAGTTTGTTGAAGCTCTGCGCGTTATTTACAGAATGAGCGACAAAACA
ATAAAAGAAAAAGTCAATGTTTACAAAAGGTTAGGCTTTGGTGTGGCTGATGTATGGGCAATCTTCAAGAAGTGGCCTTCTTTTTTGTCATACTC
AGAGAAGAAGATAACTCATACGTTTGAAACCCTTATGAGGTGCGGTCTGCTCAAACACGAGGTCCTGTCATTGATAAAGAAGCATCCAAAGTGC
ATTTGTTCTTCAGAGCAGAAGATAGTAAACTCCATTGAAAATTTTTTAGGCCTAGGATTCAGCAGAGACGAGTTTGCGATGATGATCAAGCGTTA
TCCTCAATGCATTGACTATACTGCCGAGACAGTGAAGAAGAAAACTGATTTTATTGTGAAGAAGATGAATTGGCCACTAGAGGGTTTGGTTTTGA
TCCCTCAGATATTTGGATACAGCCTGGAGAAGAGGACTGTCCCAAGGTGTAATGTAATCAAAACTCTCATGTCAAAAGGATTGCTTGGAAGTGA

AATCCCTCCAATGTCATCTATTTTGACAAGTACCGATCAGGCATTTTTGAGGAGGTATGTGATGAAGTACGACAAG------CTGGTGCCTGAGTTGAT
GGCTATCTTCACT
Al_338172
ATGAATTCTCTTATACTCGGTGCAAGAAGGTTCGTGGGGTTGCAGAAATGGCGTAACTTGAGAGTTTCATTG---------CAAAATGGATCTGCTTTTT
CTAATTCCTTCTCCTCT---GCTACTGCTGCTGATGTAAACCCTAAAGATGGTGGAAAAGGGGAGACCTTTAAAGCTTCTTCATTCCTTGATTCACT
GCGTTTAGCTGCAAAACTC------------------------------ACAAGTAAAGTCAATGCTGATTCGGTTCTGGATCTTCTTAGAAGTTATGGGTTCACAGATT
CTCAGATCTCCAGCATCATTAGGAGTGATCCACAAGTGCTTATTGCCAATTCTGCGACATCCCTGGGTTCAAAGCTTGAGTTTTTGCAGTCCAGA
GGAGCTTCAAGCTCTGAGCTCACTGAGATTGTTTCTACAGTTCCTAAAATCTTGGGGAAGAGAGCGGGCAAATCTATCAGCAGATACTATGATTT
CATCAAAGTCATTATAGAGGCGGATAAGAGTTCCAAGTATGTAAAGTTATCTCATTCTTTGCCACAGGGT---------AACAAGATCAGAAATGTTTTG
GTTTTGAGAGATTTGGGTGTGCCTCGAAAGCGGTTATTATCTTTGCTCATCTCCAAATTTCAGCCCGTGTGTGGAAAAGAAAATTTTGATGCATCA
CTCAAGAAGGTCGTTGAGATGGGTTTTGATCCAACCACCTCAACGTTTGTGCACGCTTTGCACATGCTTTACCAAATGAGCGACAAAACGATTG
AAGAAAAAGTTGAAGTCTATAGAAGTATAGGTTTCACCGTGGACGATGTATGG--------------------------------------------------------------------------------
---------------------GCAATGTTCAAGAAGTGGCCACGCTCTCTGAGACACTCGGAGAAGAAGGTAGCCAACTCCGTAGAAACATTTCTAGGTCTAGG
ATTCAGCAGAGATGAGTTTTTGATGATGTTCAAGCGGTTTCCTCAGTGCATTGGATATTCGACAGAGTTAGTGAAGAAGAAGACTGAGTTTCTGG
TGAAGGAGATGAATTGGCCAGTAAAGGCTGTGGCTTCAGTCCCTCAGAGATTAGCTTTTGTGTACGAGTGTCTGAACCTT------------------------------
------------------------TTACCC--------------------------------------------------------------------------------------------------------
Al_475136
ATGTATTCTCTGATTCTCCATGGAAGAAGGTTGGTCGAATTGCATAAATGGCGCCATTTGAGATTTGCTCTG---------CAAAATGCATCTCCTTTGTC
CAATTCCTTCTCTTCT---GCTGCTGCTGCTGATGTGAGGTCAAGAGATGGTCGAAAAGGAAAGAACTTTACGGTCTCTTACCTCGTTGATTCATTG
GGTTTCACTACAAAACTCGCAGAATCGATCTCAAGGAAAGTTCATTTCACCGACAAGGCAAATCCGGATTCAGTTCTGAGTCTTTTGAGAAGTC
ATGGCTTCATAGATTCTCAGATCTCCTGCATCATTACCGATTATCCAGAATTGCTTATACTAGATGCTGAGAAATCACTTGGTCGCAAGCTTCAGAT
TCTACAGTCTAGAGGAGCTTCAAGCTCTGAACTCACTGAGATTGTTTCTACAGTTCCGAGAATCTTAGGAAGG---------AAATCTATCACTGTATAC
TATGATGCTGTCAAAGAAATTATAGTAGCCGATAAGAGTTCGAGTTATGAA---------------CTCCCACGGGGTTCTCAGGGGAATAAAATCCGAAATG
TATCTGTTTTGAGACAATTGGGCATGCCTCAGTGGTTGTTATTACCCTTGCTTGTCTCCAAATCGCAGCCTGTGTGTGGAAAAGAAAATTTTGAA
GAATCCCTTAAGAAGGTTGTTGAGATGGGTTTTGATCCCACCACCTCTAAGTTTGTTGTAGCTTTGCGCATGCTTTACCAAATGAGCGAGAAAAC
GATTGAAGAAAAAGTTGTAGTCTATACAAGTGTAGGTTTTACTCTGGATGATGTATGG----------------------------------------------------------------------------
-----------------------------GAAATTTTCAAGAAGACTCCTTCCGTTTTGAAAGTCTCCAAGAAGAAGATATTGAAGTCTGCTGAAACATTTCTAGCCCT
AGGATTTAGCAGAGCTGAGTTTTTGATGATGGTCAAGCGTTATCCTCCTTGCATTGAATATTCTCTAGAGTCGGTTAAAAGGAAGAATGAGTTTCT
GGTGAAGAAGATGAATTGGCCACTAAATGCCTTGGTTTTGCACCCTCAGGTGTTTGGATACAGCATGGAGAAGAGGATTATACCAAGGTGTAAC
GTACTCAAAGTTCTCTTGTCGAAAGGACTGCTCAAAAGCGAACTACCTGCAGTGTCCTCTGTTTTGTCGTGTACCGATGAAGGATTTTTAAATAG
GTATGTGATGAAGCACAACGAG------CTGGCGCCTACGTTGATGGCTATCTTCACC
Al_475137
ATGTATTCTCTGATTCGCCATGGAAGAAGGTTGGTCGAGTTGCAGAAATGGCGTCATTTGAGTTTTTTAGTG---------CAAAAGGCATCTCCTTTAT
CCAATTCATTCTCCTCT---GCTACTGTTGCACGTACGAGTTCTAGAGTTGGTCGAAAAGGTAACAACTTTACAGTCTCTTACCTCGTTGATTCTTTG
GGTTTAGCTAGAAAGCTCGCAGAATCGATTTCAAGGAAAGTTAGCTTCGAGGACAAGGCCAATCCTGATTCTGTTCTGAATCTTTTTAGAAGTCA
TGGGTTCACTGATTCTCAGATCTCAAGCATTGTTACGGATTATCCACAATTGCTTATAGCTGATGCTGAGAAATCACTTGGTCCCAAGCTTCAGTT
TTTGCAGTCTAGAGAAGCTTCAAGCTCTGAGCTCACTGAGATTGTTTCACAAGTTCCGAAAATCTTGGGAAAGAGAGGGCACAAAACTATCAGC
GTATACTATGATTTCATCAAAGACACTTTATTACATGATAAGAGTTCCAAGAAGGAAAAGTCATGTCATTCTTTTCCACAGGGTAATCTGGAGAAC
AAGATCAGAAATATATCGGTTTTGAGAGAATTGGGAATGCCTCACAAGTTGTTATTCCCCTTGCTCATCTCTTGTGACGTACCTGTCTTCGGTAAA
GAAAAATTTGAAGAATCCCTCAAGAAAGTTGTTGACATGGGTTTTGATCCGACCTCTGCAAAGTTTCTCGAAGCTCTACGAGTTGTTCAAAGAT
TGAGCGACAAAACAATAGAAGAAAAAGTCAATGCGTATGAAAGGTTAGGCTTTGATGTGGGAAATGTATGG----------------------------------------------
-----------------------------------------------------------GCAGTGTTCAAGAGGTGGCCAAACTTTCTGACACATTCAGAGAAGAAGATATTGAGCACCATTGA
AACATTTCTAGGCCTGGGATTCACCAGAGATGAGTTTTCCATGTTGGTTAAGCGCTTTCCTCAGGGCATTGGATTATCCCCAGAGACGGTTAAAA
AGAAGACCGAGTTTCTGGTCAAGAAGATGAACTGGCCAATAAAGGCTTTGGTTTCAAACCCTGCGATACTTGGATACAGCATGGAGAAGAGGA
CTGTCCCAAGGGGTAACGTAATCAAAGCTCTCATCTCCAAAGGATTGATCGGAAGTGAACTCCCTTCGATATCACATGTCTTTATATGTACCAATC
AGGTGTTCTTAAACAGGTATGTGAAGAAGCATGAGGACAAGCAGCTGGTGACTGAGTTGATGGCTATCTATCGT
Al_485072

ATGTTTTCTTTAATAATCCATGGAAGAAGGTCTGTGGAATTGCAGAAATGGCGTAATTTAAGAGTTTCATTGAGTATTTTGCAAAATGCAGCTTTT

ACCACCAAATCGTTCTCTTCT---GCAATAGCTAAAGATGTGAGTCCAAAG------------GGAACAACTTTCACTGTCACTTATCTAGTTGAATCATTGG

GTTTAACTAAAAAACTTGCAGAATCAATCTCAAAGAAAGTAAGTTTCGAAGACAAGGTAAATCCGGATTCTGTTCTGAATCTTTTTAGAAGTAAT

GGGTTTAAAGATTCTCAGATCTCTAGGATCATTAGGGCTTACCCAAGATTGCTTGTAATTGATGCTGAGAAGTCTCTTCGTCCAAAGCTTCAGTTT

TTAAAGTCTAGAGGAGCTTCAAGCTCTGAGGTCACAGAAATTGTTTCGAATGTTCCTACAATCTTGGGAAAGAAAGGGGAGAAATCTATTAGCT

TGTACTATGATTTCGTTAAGGACATTATGGAAGATGGT---------------AAAAGTTTAGGTCATTCTTGGCCAGAGGGTAAGAAAGGGAACAAAATCC

GAAATATCTCGGTTTTAAGAGAATTGGGAGTGCCTCAGAAGTTGTTGTTTCCATTGGTCATCTCTAATTATCAACCTGTTTGTGGTAAAGAAAAAT

TTGAAGAAACTCTCAAGAAAGTTGTTGATATGGGTTTTGATCCGACCAAATCAACGTTTGTGGAAGCTTTGCATGTAGTTTACAAAATGAGCGAG

AAAACGATAGAAGAGAAAGTGAATGTTTATAAAAGGTTAGGCTTTAGTGAAGTGGATATATGGGCAATTTTCAAGAAGTGGCCTTTCTTTTTGAA

ATTCTCGGAGAAGAAGATAATT------------------------------------------------CTGATGTATGAAACACTGAAGAAGTGTGGTTTGGTGGAGGAAGAGGT

CATTTCGGACTCTATTGAAACGTTTCTAGACTTAGGTTTCAGTAGAGATGAGTTCAAGATGATGGTTAAGCGATATCCTCAGTGCACTGCATATAC

TGCAGAGACGGTGAGGAAGAAGTTTGAGGTTTTGGTGAAGAAAATGAATTGGCCACTAGAAGATGTAGTTTTGATCCCTGCGGTACTTGGATAC

AGCTTGGAGAAGAGGATAGTGCCAAGGACTAACGTAATCAAAGCTCTCATGTCGAAAGGATTGATTGGTAGTGAAAACCCTCCGATTTCATCTG

TCTTGGTATGTACTGATCAGGAGTTCTTGAAAAGGTACGTGATGAAGCACGACAAG------CTAGTGCCTAAGTTGATGGCTATCTTCACC

Al_489254

ATGTATTCATTGATACTCCAAGGTAGAAGGTCACGCGAGTTACATCAATGGCAGAAT---CGAGTAATGAATCTTCTTCTCCAAAACGGCTCTACTT

TCACTGAATCGTTCTCTTCTGTTGCTACCGCTAAAGATCTAAGTTTTGAAGATGAAAGAAAGAGGAAGACTTTCACAGTCTCTTACCTTATTGATT

CATTAGGGTTAACTACAAAACTCGCTGAATCAATCTCAATGAAAGCTAATTTCGATGAAAAAGGTAATCCAGATTCAGTCTTGAAACTTCTAAGA

AGCTATGGATTCAAAGATTGTCAAATCTCAAGTATCATTGCTACTTACCCACGATTTCTTGTTGAGAGTCCTGAGAAGTCTCTTCGTGCTAAGCTT

CATTTCTTGAAGCTAAATGGAGCTTCAAGCTCTGAGCTTACTGAGATTGTTTCGAAAGTTCCTAAAATCTTGGGGAAGAGAGGAGGGAAATGGA

TTATTCATTACTATGATTATGTCAAAGAGATCTTGCAAGATCAAGATAGTTCTTCTTCCAAGAGA------------------------AAGCAGACGAATCGAAATC

GAAATGTATCGGTTTTGAGAGAATTAGGAGTGCCTCAGAGACTGTTGTTGAACTTGTTGATATCTAGAGCTAAACCAGTTTGTGGGAAAGAAAG

ATTTGAAGAATCGGTTAAGAAGATTGTTGAGATGGGTTTTGATCCAAAAAGTCCAAAGTTTGTCAGTGCTTTGTATGTTTTCTATGATCTTAGTGA

TAAGACTATAGAGGAGAAAGTGAATGCTTATAAGAGGTTAGGACTTTCTTTGGATGAAGTTTGGGTAGTTTTCAAAAAATGGCCTTTCTCTTTGA

AATACTCGGAGAAGAAGATAATTCAGACGTTTGAGACTTTGAAACGGGTTGGTCTAAGGGAAGAAGAGGTATGCTTAATGGTTAAGAGGTACCC

TGAATGTGTAGGGACTTCAGAGGAGAAGATAGTGAAGTCAGTTGAAACTTTTCTTGAGTTAGGATTCACAAAAGACGAGTTTGTGATGATCATC

AAGCGGCATCCTCAGTGTATTGGATTAGCTGCGGATTCGGTGAAGAAGAAGACCGAGTTTCTTGTGAAGACAATGGGTTGGCCTTTAAAGGTTG

TAGCTTCAACCCCCATTGTACTTGGATTCAGCTTGGAGAAGTTTGTTTTACCGCGGTGTAATGTAATCAAAGCTCTTTTGTCTAAAGGGCTGATC--

-GATGAAATCCCTGCGATCTCGTCTGTATTGACGAGTCCTAAGCTGAAGTTTTTGAAGCTCTTTGTGGAGAAGCACCAAGAT------GTGTTGCCTG

AGTTGAACTCTATCTTCACT

Al_893233

ATGTATTCTCTGATACTCCATGGAAGAAGATCTACTGAGTTACATAAATGGCGTAACTTTAGTGTTTCAGTGAAGCTTTTGCAAAATGTATCTGCTT

TTTCCAATTCCTTCTCCTCT---------GCTGCTGATGTGAGCCTTAGAGATGGTCGAAAAGGTAAGAACTTTACAATCTCTTACCTTGTTGATTCATTG

GGTTTACCTATAAAGCTCGCAGGATCGATTTCAAGGAAAGTCAGGTTTGAGAACAAGGCCAATCCCGATTCTGTTCTGAGTCTTCTGAGAAGTCA

TGGGTTCACAGATTCTCAGATCTCCACCATCATTACGGATTTTCCAACATTGCTTATATTAGATGCTGAGAAATCACTTGCTCCCAAGTTTCAGTTT

TTGCAGTCCAGAGGAGCTTCAAGCTCTGAGCTCACTCAGATTGTTTCTACAGTTCCGGAAATCTTGGGAAAGAGAGGGGACAAAACTCTCAGC

TTATGCTATGATTTCGTCAAAGAGAGTTTAGTAGCGGATAAGAGTTCCAAGTTGGAAAAGTTGTGTCATTCTTTGCCAGAGGGTAAGCAGGAGGA

TAAAATCAGAAATGTATCGGTTTTGAGAGAATTGGGCATGCCTCACAAGTTGTTATTCTCCTTGCTTACATCCGTTGGTCAACCTGTATGTGGAAA

AGATAGATTTGATGCATCCCTCAAGAAGATAGTTGAGATGGGTTTTGATCCGACCACCGCAAAGTTTGTCAAAGCTCTATACGTTGTTTACAATTT

GAGTGACAAAACAATAGAAGAAAAAGTCCATATTTACAAAAGGTTAGGCTTTGCTGTGGAAGATGTATGGGTAATCTTCAAGAAGTGGCCTTTC

TCTTTGAAATTCTCAGAGGAAAAGATAACTCAGACGATTGAAACCTTGAAGATGTGTGGTTTGAACGAAAACGAGGTCCTTCAAGTATTGAAGA

AGTATCCGCAGTTTATACGTATGTCACAGCAGAAGATATTGAACTTCATTGAAACATTTTTAAGTCTAGGATTCAGCAGAGATGAGTTTACGATGA

TAGTCAAGTGCTTTCCTATGTGCTTTGGATTGTCTGGAGAGACGGTGAAAAAGAAGACTGAGTTTGTGGTGAAGAAGACTAATTGGTCACTAAA

GGACACTACTTCGTTCCCTCAGGTATTTGGATACAGCCTGGAGAAGAGGATTGTACCAAGGTGTAACGTAATCAAAGCACTCATGTCCAGAGGA

TTGCTTGGAAGCGAACTCCCTTCAATGGCGTCTGTCTTGGCATGTAACGATCATGCTTTTGTTAAAAGGTATGTGAGGAAGCAGAATGACAAGG

AGCTTGTGGCTGAGTTGATGGCTATTTTCATT

Al_893270

TGTTCTTCTCTGATTCCTCATGGAAAAAGGTTGGTCCAGTTTCAGAAAGGGCATAACTTTAGTGTCTCAGTGAAGCTTTTCCAAAATGTATCTGCT
TTTTCCAATTCCTTCTCATCTGTTGCTAGTTCTCATGATGTGAGCATTAGAGATGGTCGAAAAGGTAACAACTTTACAGTCTCTTACCTCATTGATT
CACTTGGTTTAACTAAAAAGCTCGCAGAGTCGATTTCAATAAAAGTCCGTTTCGAGAACAAGGCCAATCCTGATTCTGTTCTGAGTCTTTTGAGA
AGTCATGGGTTTACAGATTCTCAGATCTCCAACATCATTACGGATTATCCACTATTGCTCATAGCAGATGCTGAGAATTCCCTTGGTCCCAAGCTTA
AGTTGCTGCAGTCAAGAGGAGCTTCAAGCTCTGAGCTCACTGAGATTGTTTCTAAAGTTCCTAAAATCTTAGCAATGAAAGGGGACAAAAGTAT
CAGCAGATACTATGATATCGTCAAAGAGATTGTAGAAGCGGATAAGAGTTCCAAGTTCGAAAAGTTGTGTCATTCTTTGCCAGAGGGTAAGCAG
GAGAATAAAATCAGAAATGTATTGGTTTTGAGAGAATTGGGCGTGCCTCAAAGGTTGTTATTCTCCTTGCTCATCTCTAATCACCATGTCTGCTGT
GGAAAAGAAAAATTTGAAGAATCACTCGAGAAGGTTGTTGGGATGGGTTTTGATCCCACCACCCCAAAGTTTGTGGAAGCCTTATGCATTGTTT
ACGGATTGAGCGACAAAAGGCTAGAAGAAAACTTCAATGTCTATAAAAGGTTCGGCTTAACTGTAAACGATATATGGGAACTTTTCAAGAAGTG
CCCTGCCTTTCTTGGATACTCAGAGAATAGGATAATTCAGACATTTGAAGCCTTAAAGAGGTGTGGTCTGTGCGAAGATGAGGTCATGTCAGTGT
TCAAGAAGAATCCACTATGCTTACGTGCTTCAGAGCAGCAGATATTGAATTCCATGGAAACATTTATTGGCTTAGGATTTAGCAGAGATGAGTTTG
TGATGATGGTCAAGCGCTTTCCTCAGTGCATTGGATATTCTGCAGAGATGGTGAAGAAGAAGACTGAGTTTGTGGTGAAGAAGATGAATTGGCC
ACTAAAGGTCATAACTTTGTTCCCTCAGGTACTTGGATACAGCATGGAGAAGAGGATTGTCCCAAGGTGTAACGTAATCAAAGCTCTTATGTCCA
AAGGATCGCTTGGAAGCGAACTCCCTCCAATGCCGTCTGTCTTGGCATGTACCGATCAGACCTTCTTAAACAGGTATGTGGTGGAGCATGATGAA
---AAGCTGGTGCTTGAGTTGATGGCTATCTTCAAC

Al_893275

ATGTCTTCTCTGATACTCCACCATAGGTCTCTTAAGTTACTACACAAGTGCCCTAATTTGAGAGTCTCACCG---------CAAAACCCATTTTCTTTTTC
CAGTTCCTTCTCTTCT-----------------ACTAGAGCTAAAGATGGTCAAAAAGGTCAGATTTTCACTATCTCTTACCTCATTGATTCATTGGGTTTAACC
GCAAACCTCGCTGAATCGATCTCAAGAAAAGTAAGCTTCGAGGAGAGGAGAAATCCTGACTCTGTTCTGAATCTTTTTAGAAGTTATGGTTTCA
CAGATCCTCAGATCGCAAGCATCATTACTGATTATCCACGATTGCTTATAGTAGATGCTAAGAAATCTCTTGGTCACAAGCTTCAAGTTTTGCAGT
CTAGAGGAGTTTCAAGCTCTGAGCTCACTGAGACTGTTTCTAAAGTTCCTAAAATCTTGGCAATGAAAGGGGACAAAACTATAAGCAGATACTAT
GATTTCGTCAGAGAGATTATCGAAGCGGGTAAGAGTTCTAAGTTCGAAAAGTTATGTCAATCTATGCCACAGGGTATGCAAGAGAATAAAATCCG
AAATTTATCTGTTTTGAGAGAACTTGGCGTGCCTCAGAGGTTATTATTCCCTTTGCTCGTCTCTGATCGTAAACTTGTATGTGGAAAAGAAAAATT
TGAAGAATCTCTTAAGAAGGTTGTTGAGATGGGTTTTGAACCGACAACCTCAAAGTTTGTCAACGCTCTACGTGTTGTTCAAAGAATCAGCGAG
AAAGAAATAGAAGAAAAGGTCAGTTTCTATAAAAGGTTAGGCTTTGATGTGGGAGATGTATCTGAAATGTTCAAGAAGTATCCAGTCTCTATGAG
ATTATCCGAAAAGAAGATAACTCAGAAGTTTGAAACCTTGAAGAAGTGTGGACTACTCGAAGAC---------GAGATCCTCTCAGTGTTTCCGCAAT
GCATTGGTGCTTCGGAGCAGAAGATAGCAAAATCCATTGAAACATTTAAAGACCTCGGATTCAGCAAAAATGAGTTTGCGTTTATGGTCAAGCA
CTTTCCTATGTGTCTTAACATATCCGCAGAGACGGTGAAGAAGAAGACGAAGTTTCTGGTGAAGAAAATGAAATGGCCACTAAATTCTGTGGCT
TTTTACCCGCAAGTACTTGGACTAAGCATGGAGAAGAGGATTGTACCAAGATGTAACGTTATGAAAGCACTCATGTCGAAAGGATTGCTTGAAA
GCAAATTACCTTTAAAGGAACGTCCAAGACTGTGTGTGGTCTGCAAAATCTTCAAGAAACTTATAGCAGCTCGTATGATTAATAATGCTTTCGAG
ACTCTTGCCTTTGTCTTAAGT

Al_893280

------------------------------------------------------------------------------------------------------------------------------------------------ATGGTCAAAAAGGTAAGAACT
TTTACTGTCTCTTACCTCGTTGATTCACTTGGTTTAGCTAAAAAGGTTGCAGAATCGATTTCAAGAAAAGTCAGTTTCGAGAACAAGGGCAATCC
TGATTCTGTTCTGAGTCTTTTGAGAAGTCATGGGTTCACAGATACTCAGATCTCCAGCATCATTACGGATTATCCACTATTGCTCATAGCAGATGGT
GAAAATTCCATTGGTCCCAAACTTAAGTTTTTGCAGTCTAGAGGAGCTTCAAGCTCTGAGCTCACAGAGATTGTTTCTAAAGTTCCGAGAATCTT
GGGAAAGAGAGGGCACAAAACTATCAGCAGATACTATGATACCGTCAAAGAGATTGTAGAAGCGGATAAGAGTTCCAAGTTCGAAAAGTTATGT
CATTCTTTGCCACAGGGTAAGCAGGAGAACAACATTAGAAATGTTTTGGTTTTGAGAGAATTGGGCGTGCCTCAGAGGTTGCTATTCTCCTTGCT
CATCTCTGATAACGGACATGTATGTGGAAAAAAAAAGATTTGAAGAATCACTCAACAAGGTTGTTGAGATGGGTTTTGATCCGACCACTGCAAGT
TTTGTCCGTGCTCTGCACGTTATTCAAGGATTTAGCGACAAAACAATAGAAGAAAAGGTCAATCTCTATAAAAGGTTAGGCTTTGATGTGGGAGA
TGTATGG-------------------------------------------------------------------------------------------------------GAAATGTTCAAGAAGTTTCCAACGTTTCTAGGACTC
TCGGAGAAGAAAATAGCCAACTCCATTGAAACATTTGTAAGCCTAAGATTCACCAGAGATGAGATTGTGGTCATGGTCAAGCGCTTTCCTCCGT
GCATTGGATGTTCTGCAGAGTCGGTGAAGAAGAAGACTGAATTTCTGGTAAAGAAGATGAATTGGCCACTAAAGGCTGTGGCTTCATTCCCTCA
GGTAATTGGATACAGCCTAGAGAAGAGGACTGTCCCAAGGTGTAATGTAATCAAAGTTCTCATCTCAAAAGGATTGCTCGGAAGTGAACTCCCT
CCATTGTCGTGTGTGTTGTCAATTACCGATCCAGCTTTCTTAAACAAGTATGTGGTGAAACATGATGACACGCAGCTCGTGCGTGAGTTAATCGCT
ATCTTCACC

Al_893283

ATGAAGTCTCTGATAATC------CGTAGGTTCGTGGGGTTGCAGAAATGGCGTAACTTGAGAGTTTCATTG---------CAAAATGGATCTTCTTTTTCCA
ATTCGTTCTCCTCT---GCTAGTGATGCTGATGTGAGCCTTAGAGATGGTCTAAAAGGTAACAAATTTAAAGCCTCGTGCCTCGTTGATTCACTGGG
TTTAGCTTCAAATCGCACTACATCGGTCTCAAGTGAAGTCAGTTTCACGGACAAGGTAAATCCTGATTCTATTCTAAATCTGTTTAGAAGTTATGG
GTTCACAGATTCTCAGATCTCCAACATCATTAGGACTTACCCACGGTTGCTTATAGCAGATTCTCAGAAATCACTGGGTTTCAAGCTTAAGTTTCT
GCAGTCTAGAGGAGCCTCAAGCTCTGAGCTCACTGAGATTGTTTCTTCACTTCCAAAAATCTTGAGAAAGAGAGGTCACAAAACACTCAGCTTA
TTTTATGATTTCGTCAAAGAGATTATACAAGTGGATAAGAAAAGAAAC---------TTATCTCAATCTTTTCTGCAGGAG---------AACAAGATCAGAAA
TATTTTTGTTTTGAGAGAATTGGGTGTGCCGCGAAAGCGGTTATTATCTTTGCTCATCTCCAAATCTCAGCCCGTGTGTGGAACAGAAAGATTTGA
TGCATCACTCAAGAAGGTCGTTGAGATGGGTTTTGATCCAACCACCTTAATGTTTCTCCAAGCTTTGCACATGCTTCACCAAATGAGCGACAAAA
CGATTGAAGAAAAGATTCAAGTCTATACAAGTGTAGGTTTCACTGTGGATGATGTATGG------------------------------------------------------------------------------
---------------------------------GCAATGTTCAAGAAGTGGCCACTCTCTCTGACACACTCGGAGAAGAAGGTAGCCAACTCCATTGAAACATTTTTTAGT
CTAGGATTCAGCAGAGATGACTTTGTAAGGATGGTCAAGCGGTTTCCTCAGTGCATTGGATTATCGGCAGAGTTGGTGAAGAAGAAGACTGAGT
TTCTGGTGAAGAAGATGAATTGGCCACTAAAGGCTGTGGTTTCAAACCCTACGGTACTTGGATACAGCCTAGAGAAGAGGACTGTCCCAAGGTG
TAACGTAATCAAAGCTCTTATGTTAAAAGGATTGCTCGGAAGTGAACTCCCGCCAATGATGTCTGTGTTGGCGATTACTGATAAAGCTTTCTTAAA
CAGGTATGTTATGAAGCATGATGATAAGCAGCTAGTGCCTGAGCTAATGGCTATCTTACTC

Al_893289

---------------------------------------------------------------------------------------------------------------------------------------------------------------------------------------------------------------
-----------------------------------------------------------------------------------------------------------------------------------ATGTCCAACATCATTAAGATGTATCCACTATTGCTT
ATAGCAGATGCTGACAAGTCACTTGGTCCCAAGCTTCAGTTTTTGCAGTCAAGAGGAGCTTCAAGCTCTGAGCTCACTCAGGTTGTTTCTAAAG
TTCCTAAAATCTTGGGGAAGAGAGAGGGCAAATCTCTTAGCAGATACTATGATTTCATCAAAGTCATTATAGAAGCAGATAAGTCCTCCAAGTAT
GAAAAGTTATGTCATGCTTTGCCAGAGGGTAGGCAGGATAATAAAATCCGAAATGTTTTGGTTCTGAGAGAATTGGGTGTGCCTCAGAGGTTGTT
ATTCTCCTTGCTCATCTCTGATAGCGGACCTGTATGTGGAAAAGAAAAATTTGAAGAATCCCTCAAGAAGGTTGTTGAGATGGGTTTTGATCCAA
CCACTTCGAAGTTTGTCAAAGCTCTCCATGGCTTTTACCAAATGAGTGACAAAACAATTGAAGAAAAACTCGATGTCTACAAAAGGTTAGGCTT
TTCTGTGGAAGATGTATGGGTAATCTTCAAGAAGTGGCCTTGCTCTTTGAAATTCTCAGAGGAAAAGATAACTCAGACGATTGAAACCTTGAAG
ATGTGTGGTCTGGACGAAAACGAAGTCCTTCAAGTGCTGAAGAAGTATCCGCAATTCATACGTATTTCAGAGCAGAAGATATTGAGTTTAATTGA
AACATTTTTAGGTGTAGGATTCAGTAGAGATGAATGCGTGATGATAATCAAGGGCTTTCCTATGTGCTTTGGATTGTCTGCAGAGACGGTGAAAA
AGAAGACTGAGTTTTTGGTGAAAAAGATGAATTGGCCACTAAAGTCTGTGGTTTCGAACCCTGCGGGACTTGGATACAGCTTGCAGAAGAGGA
TTGTTCCAAGGTGTAATGTAATCAAAGCTCTCATGTCCAAAGGATCGCTTGGAAGTGAACTCCCTTCAGTGGCTTCTGTCTTGGCATGTACTGAT
CAGGCGTTCTTAAACAGGTATGTGGTGAAGGATGATGATAAGCTCCTTGTGCTTAAATTAATGGCATGGCTATCT

Al_893298a

ATGTGTTCTCTGATACTCCATGGAAGAAGGTTGGTCCAGTTACAGAAATGGTGTCATTTGAGTTTTTCAGTG----------------------CAAAAGGCTTCTT
CTTTTTCCACT-------------------GTGAGCACTAAAGATTGTCGAAAAGGGGAGATTTTTACGATCAGTTACCTCGTTGATTCTCTTGGTTTAACTAGAA
AGCTCGCAGAATCAATCTCC----------------GAGGGCAAGGCAAATCCTGAGTCAGTTCTGAGTCTTTTGACTAGTCATGGGTTCACAGATTCTCAGA
TCTCCAGCATCATTACGATTTATCCACGATTGTTTTTATTAGATGCTAAGAAATCACTTGCTCCCAAGCTTAAGTTTTTGCAGTCCAGAGGAGCTTC
AAGCTCTGAGCTCACCGAGATTGTTTCAAAGGTTCCGGAAATATTGGCAAAGAAAGGAGACAAAACGCTAAGCAGATACTATGATTTCGTCAAA
GTCATTGTTGAAGCAGATAAGAGTTCCAATTACGACAAGTTATGTCATTCTTTGCCTGTGGGTAATCTAGAGAATAAAATCCGGAATATATCGGTT
TTAAGAGAACTCGGTGTGCCTCAGAGGTTGTTATTCCCCTTGCTTATCTCTAGTGGGGGACCTGTCAATGGGAAAGAAAGGTTTGGAGAATCAAT
TAAGAAGCTTGTTGAGATGGGTTTTGATCCGACCACTACAAAGTTTGTTAAAGCTCTGCGCATTGTTCAAGGATTAAGCGCAAAAACAATAGAA
GAAAAAGCCAATCTTTATAAAAGTTTAGGCTTT------GATGATGTATGG--------------------------------------------------------------------------------------------------------
---------GAAATTTTCAACAAATATCCAATCTTTCTGGCGCTCTCTGAGAAGAACATATTGAACTCTGTAGAAACATTCCTAGGTCTAGGATTCAGCA
GAGATGAATTTGCAAATATGGTCAAGAGCTTTCCTCAGGGCATTGGATTATCCGCAGAGACGGTTAAAAAGAAGACCGAATTTCTGGTGAAGAA
GATGAATTGGCCGCTAAAGGCCCTGGTTTTAAACCCTGCGGTACTTGGATACAACATGGAGAAGAGGATTGTACCAAGGTGTAACGTAATTAAA
GCTCTCATGTCCAAAGGACTGCTCGGAAGCAAACTCCCTCCAATCGGGTCTGTTTTGAAGAGTACCAATCAGGTCTTCTTTAAGAGGTATGTGAT
CAAGCACAATGACAAGAAGCTTGTGGCTGAGTTGATGACAATCCTCACT

Al_893298b

ATGTATTCTCTGATACTCCATGGAAGAAGGTCAGTCGAGTTGCAGAAATGGCGTAACTTGAGTGTTTCAGTG---------CCAAATGCATCTTCTTTTT
TCAATTTGTTTTCCTTTGCTACTACTGCTGCACATTTGAGTCCTAGAGTTGGTCGAAAAGGAAAGGACTTTACTTTCTCTTACCTCGTTGATTCATT
AGGTTTACCTAAAAAGCTCGCAGAATCGATTTCAAGGAAAGTCAGCTTCGAGGACAAGGGCAATCCTGATTCTGTTCTGAGTCTTTTTAGATGTC

AGGGGTTCACTGATTCTCAGATCTCTAGCATGATTGAGATTTACCCACGTTTGCTTATACTAGATGCTGAGAAATCCCTTGGTCCCAAGCTTCAGT
TTTTGCAGTCCAGAGAAGCGTCAAGCTTTGAGCTCACTCAGATTGTTTCAAAGGTTCCGGAAATATTGGGAAAGAAAGGGGACAAAACTATCA
GCGTATACTATGATTTCATCAAAGACACTTTACATGATAAG---AGTTTCAAGTACGAAAAGTTATGTCATTCTTTTCCACCGGGTAATCTAGAGAAC
AAGATCAGAAATGTATCGGTTTTAAGAGAATTGGGCATGCCTCACAAGTTGTTATTCTCCTTGCTCATCTCCGATAGCCAACCTGTATGTGGAAAA
GAAAAATTTGAAGGAACCCTCAAGAAGGTTGTTGAGATGGGTTTTGATCCAACCACCGGAAAGTTTGTCGAAGCTCTAAACGTTATTTACAAAA
TGAACGAGAAAACAATCGAAGAAAGATTCAATCTCTATAAAAGCTTAGGCTTTGATGCGGGAGATGTATGG-------------------------------------------------
----------------------------------------------------------TCAAGTTTCAAAAAGTGGCCAATCTCTCTGAGAGTAACGGAGAAGAAGATGTTGGACTCCATTGA
AACATTTCTTGGCCTAGGATTCAGCAGAGATGAGTTTGCGAAGATGGTCAAGCACTTTCCTCCGTGCATTGGATTATCTACAGAGATGGTCAAAA
AGAAAACTGAGTTCTTGGTGAAGAAGATGAATTGGCCACTAAAGGCTCTGGTTTCAAACCCTGCAGTACTTGGATACAGCCTGGAGAAGAGGA
TCGTCCCAAGGGGTAACGTAATCAAAGCTCTCATGTCCAAAGGATTGATCGGAAATGAACTCCCTTCGATATCGTGTGTCCTCATGTGTTCCAAA
CTGGTGTTTTTAAATAGGTATGTGGAGAATCACGAGGACAAGCAGCTGGTGACTGAGTTGATGGCTATCTATCGT

Al_893300

ATGTATTCTGTGATTCTCCATGGAAGAAGGTTGGTCCAGTTGCAGAAATTGCCTTATCTGAGATTTGCAGTA---------GAAAATGCATCTACTTTTTC
CAATCCGTTCTCTTCTGCTGCAAGTGATGCTGATGTGAGCCTTGGAGATGGTCGAAAAGGCAAGACTTTCACAGTCTCTTACCTCGTTGATTCAT
TGGGTTTATCGAAAAAGCTCGCAGAATCTATTTCTAGGAAAGTCAGCTTCTCTGGCAAGGGTAATCCTGATTCTGTTCTGAGTCTTTTGAGAAGT
CATGGGTTCACAGATACTCAGATCTCCACCATCATTACGAATTACCCACGATTGCTTACATTAGATGCTGAGAAATCCCTTGGTCCCAAGCTTCAG
TTTCTGCAGTCAAGAGGAGCTTCAAGCTCTGAGCTCACTCAGATTGTTTCTACAGTTCCTAAAATCTTGGGAAAGAGAGGGCACAAAACTATCA
GCAGATACTATGATTTCGTCAAAGTCATTATAGAAGCGGACAAGAGTTCCAAGTACGAGAAGTTATGTCATTCTTTGCCACAGGGTAAGCAGGAG
AACAAAATCCGAAATTTATTGGTTTTGAGAGAACTGGGGGTGCCTCAGAGGTTGTTATTCTCCCTGCTCATCTCCAATCAACATGTCTGCTGTGG
AAAAGAAATTTTTGAAGTATCACTCAGAAAGGTCGTTGATCTGGGTTTTGATCCCACCACCTCAACGTTTGTCGAAGCCCTGTGCACTGTTTACG
GAATGAGCGACAAAACAATAGAAGAAAAAGTCGATGTCTATAAAAGGTTAGGCTTTGCTGTGGAAGATGTATGG------------------------------------------
-----------------------------------------------------------------GCAATGTTCAAGAAGTGGCCACTCTCTCTTGCAAACTCGGAGAAGAAGGTAGCCAACTCCAT
TGAAACATTTCTTGGTCTAGGATTCAGCAGAGATGACTTTGTAAGGATAGTCAAGCGGTTTCCTCAGTGCATTGGATTATCTGCAGAGTTGGTGA
AGAAGAAGACTGAGTTTGTGGTGAAAAAGATGAATTGGCCACTAAAGGCTCTGGTTTCAAACCCTCAGGTACTTGGATTAAGCATGGAGAAGA
GGATTGTCCCAAGGTGTAACGTAATCAAAGCTCTCATCTTAAAAGATCTGCTTAGAAGCAAACTCCCTCCACTCCGGTATGTTTTG---ATTACGGA
TGAGAAGTTCTTGGAAATGTATGTCAGGAAGCACGATGACAAGCAGCTTGTGGCCGAGTTGATGGCTATCTTCACT

Al_894067

ATGAATTCTCTGATAATC------CGTAGGTTCGTGGTGTTGCAGAAATGGCGTAACTTGAGAGTTTCATTG---------CAAAATGGATCTTCTTTTTCCAA
TTCGTTCTCTTCT---GCTAGTGCTGCTGATGTGAGCCCTAAAGATGGTGGAAAAGGGGAGACCTTTAAAGCTTCGTCATTCCTTGATTCACTGCGT
TTA----------------------------------------------------GTCAATGCTGATTCGGTTCTGGATCTTCTTAGAAGTTATGGGTTCACAGATTCTCAGATCTCCAGCAT
CATTAGGAGTGATCCACAAGTGCTTATAGCCAATACTGCGACATCCCTTGGTTCAAAGCTTGAGTTTTTGCAGGCCAGAGGAGCGTCAAGCTCTG
AGCTCACTGAGATTGTTTCTACAGTTCCTAAAATCTTGGGGAAGAGAGAGGGCCAATCTATCAGCAGATACTATGATTTCGTCAAAGTCATTATAG
AAGCAGATAAGAGTTCCAAGTATGTAAAGTTATCTCATTCTTTATCACAGGGT---------AACAAGATCAGAAATGTTTTGGTTTTGAGAGAATTGGG
TGTGCCTCAAAAGCGGTTGTTACCTTTGCTCATCTCCAAAGCGCAGCCCGTGTGTGGAAAAGAAAAATTTGATGCATCACTCAAGAAGGTCGTT
GAGATGGGTTTTGATCCAACCACCTCAACGTTTGTT----------------------------------------------------------------------GTAGGTTTCACTGTGGATGATGTAT
GG-----------------------------------------------------------------------------------------------------------GCAATGGTCAAGAAGTGGCCACGCTCTCTGACACACTCGG
AGAAGAAGGTAGCCAACTCCATTGAAACATTTCTTGGTCTAGGATTCAGCAGAGATGAGTTTTTGATGATGGTCAAGCGGTTTCCTCAATGCATT
GGATTTTCGACAGAGTTAGTGAAGAAGAAGACTGAGTATCTGGTGAAGGAGATGAATTGGCCACTAAAGGCTGTGGCTTCAATCCCTCAAGTAG
TTGGATACAGCCTAGAGAAGAGGACTGTCCCAAGGTGTAATGTAATCAAAGTTCTCATCTCAAAAGGATTGCTCGAAAGTGAACTCCCTGCAAT
ATCGTCTGTTTTGACAAGTACTAGTGAGAAGTTTTTAAATTGTTATGTTAGGAAACATGATGACAAGCAGCTTGTGGCTGAGTTGATGGTTATCTT
CACT

Al_907730

ATGCATTCTCTGTTAATCCATGGAAGAAAGTCTCTTTTGAGAGGTTCAGTGCAAAAACATGTATTC------------------------TCTGCTTTTCCCAGAACG
TTCTCCTCC------------------------GTAAGTGATGGTCGAAAAAGGAGGAATTTGCCAGTC---------GTCGCTTACGAGGGTCTAACTCTAAAACAGACA
GAAAGGTTAACGAGGTTAGTCTACTCC------------AAGCAAGATGCAATTCTGATTCTTCTTAGACGTCATGGGTTCACAGATTCTCAGTTCGGCGA
CATGGTTGAGAGTTACCCACCATTGTTTGACTTAGATGCTCGCAAGTCCATTGCTCCCAAGCTTAAGTTTCTGCGGTCTAGAGGAGCTACAAGTC
TTGAACTCTCTGAGATCCTACCGAAAATTCCTAAAATCTTGGGAATGGAAGGGACTAAAACTGCCGGCTTATACTATCATGTCTTCAAATACATGA

CAACCGCAGATAAGAGTGGAAATTTAGCTCCA------------CTAAAGGGTGGTATGCAGGGGAATGTCATGAGAAATGTATGGGCTTTGAGAGAACT
TGGTGTGCCTCAGAATCTCTTGTTATCCTTGCTTACATCTGATAACAAACTTGTCGGGAAAAGAAGAAGATTTGAAGAAACAGTGAACAAGGTT
GTTGGTAAGGGTCTTGATCCCACGAAGCCAAAGTTTGTCGAGGCTCTAAAAGTTATTTACAAAATGAGTGACAAAACAGAAGAAGAAAAGATC
AATATCTATAAGAGGTTAGGCTTTGCTGTGGGGGATGTATGG------------------------------------------------------------------------------------------------------T
CTCTTTTCAAGAAGTTTCCAAGGATTCTGGCACTCCCGGAGAAGAATATATTAAACTCCAGTGAAACCTTTCTAAGCCTAGGATTCAGCAGAGAT
GAGTTCAAGATGATGATAAAACGACATCCTCCATGCATTGCATATTCTGCAGAGTCGGTGAAGAAGAAGGCTGACTTTCTGATGAAGGAGATGA
AATGGTCTTTGTGC---------------CCTAAGATGCTGTCTTACAGCATGGAGGAGAGGATTCTACCAAGGTGTAACGTAATTAAAGCTTTGATGTCGAA
AGGATTAATTGGAAGTGAATTCCCCTCTGCCGCCACTGTGTTGATATGTACAAATCAGAGTTTCTTGAAAAAATTTGTGAGAAAACACGAAGAC
AAGGAGCTTGTGGCTGAGTTGATGGCTCTTTTCACC

### 2) ML trees of paralogous *mTERF* gene sets (in newick).

#### Osa_1.tree

((((Osa_002:0.09520181,Osa_003:0.13685114):0.03908026,Osa_005:0.18641467):0.03363709,Osa_004:0.18847379):0.12732986,Osa_001:0.34738161);

#### Osa_2.tree

((((((Osa_009:0.06413352,Osa_010:0.08270421):0.01593113,Osa_011:0.10743229):0.00480835,Osa_014:0.11278497):0.06381382,Osa_012:0.17690421):0.02821538,(Osa_007:0.09754833,Osa_008:0.11511010):0.13347973):0.04995343,(Osa_006:0.25324295,Osa_013:0.22221921):0.04530032);

#### Osa_3.tree

(((Osa_015:0.24490095,Osa_016:0.27193806):0.15423114,Osa_017:0.38916404):0.01341544,Osa_018:0.43958464);

#### Ptr_1.tree

((((((((Ptr012:0.00933074,Ptr013:0.03773677):0.26231783,(Ptr014:0.23478801,Ptr015:0.24213606):0.01149338):0.04927254,(Ptr002:0.33156955,(Ptr004:0.23312505,Ptr007:0.24999099):0.02157953):0.01261339):0.01983129,(Ptr005:0.28715420,(Ptr008:0.25422291,Ptr010:0.29301766):0.00662496):0.01399614):0.00534136,(Ptr006:0.26022993,Ptr009:0.27385256):0.00868911):0.00739366,Ptr011:0.29223592):0.02918100,Ptr003:0.25268094):0.04024172,Ptr001:0.45131618);

#### Ptr_2.tree

(((Ptr018:0.07986314,Ptr019:0.10245842):0.17485571,Ptr016:0.23550414):0.00298777,Ptr017:0.28030189);

#### Ptr_3.tree

((((((Ptr020:0.00000000,Ptr027:0.00000000):0.02989333,Ptr023:0.04514186):0.04552859,(Ptr021:0.08248886,Ptr022:0.08073903):0.01469779):0.05775541,Ptr026:0.15475410):0.13729086,Ptr025:0.20710244):0.01384247,Ptr024:0.30607540);

#### Ath.tree

(((((((AT1G61970:0.15081520,AT1G61980:0.13869003):0.05579757,AT1G62490:0.24690816):0.01172598,(AT1G61990:0.19757908,(AT1G62010:0.17898866,AT1G62120:0.12522271):0.03556825):0.01494398):0.01451313,(AT1G61960:0.12248716,(AT3G46950:0.19574166,AT5G23930:0.26881237):0.01922804):0.01539830):0.00812862,AT1G62085:0.12958672):0.01969547,(AT1G62110:0.12044832,AT1G62150:0.17638295):0.01904825):0.07320564,AT1G56380:0.40446843);

#### Aly.tree

((((((((((Al_338172:0.08128361,Al_894067:0.02324975):0.02985001,Al_893283:0.07281196):0.03467747,Al_893300:0.06965633):0.01732949,Al_893270:0.06961425):0.00325917,(Al_315177a:0.07820340,Al_893280:0.07690534):0.00734923):0.00693699,(Al_315177b:0.08706212,Al_893275:0.12637278):0.00482028):0.00018522,((Al_893233:0.08420207,Al_893289:0.08841034):0.00465876,(Al_893298a:0.11140494,(Al_475137:0.08859802,Al_893298b:0.07716045):0.01588142):0.00681500):0.00734234):0.00750375,Al_475136:0.11528353):0.01410683,(Al_315190:0.08889768,(Al_485072:0.14019279,Al_489254:0.20941935):0.01337281):0.00425058):0.02251694,Al_907730:0.24955967);

**3) Control file for PAML analysis.**

```
      seqfile = Osa_2_CDS.phylipi
     treefile = Osa_2.tree

      outfile = Osa_2
        noisy = 3
      verbose = 0
      runmode = 0

      seqtype = 1
    CodonFreq = 2
   aaRatefile = jones.dat
        model = 0
      NSsites = 1 2 7 8
                   * 0:one w;1:NearlyNeutral;2:PositiveSelection; 3:discrete;
                   * 4:freqs;5:gamma;6:2gamma;7:beta;8:beta&w+;9:beta&gamma;
                   * 10:beta&gamma+1; 11:beta&normal>1; 12:0&2normal>1;
                   * 13:3normal>0;
        icode = 0
    fix_kappa = 0
        kappa = 1.6
    fix_omega = 0
        omega = .5

        ncatG = 10
        getSE = 0

   Small_Diff = .1e-6
*    cleandata = 1
   fix_blength = 2  * 0: ignore, -1: random, 1: initial, 2: fixed
```
